# Co-evolved Partners of Immunity: A Trait-Based Map of Human Keystone Organisms

**DOI:** 10.1101/2025.08.19.671142

**Authors:** Amir Asiaee, Natalie Mallal, Elizabeth Phillips, Simon Mallal

## Abstract

Persistent human-adapted microbes can act as immunological “keystones,” organizing host defense across tissues and shaping vulnerability under immune perturbation. More generally, tissue immunity is calibrated by persistent niche-resident organisms that tune compartment-specific thresholds of cytotoxicity and peripheral tolerance; keystone organisms represent the apex subset with multi-niche scope. Here we operationalize keystone organisms as pathogens whose containment requires coordinated engagement of multiple immune arms and whose residence is structured across anatomical niches. Using 18 curated immunological and evolutionary traits across 43 organisms, unsupervised analyses resolved four reproducible archetypes and identified a compact keystone set dominated by persistent herpesviruses and *Mycobacterium tuberculosis*. We then translated the clinical literature into a pathogen×immune-perturbation×niche tensor capturing where and when each organism emerges under defined immune deficits. We quantified “diagnostic breadth” with two complementary summaries: immune breadth (diversity of perturbations associated with emergence) and niche breadth (diversity of anatomical sites). Clinical emergence patterns perfectly separated trait-defined keystones from all other organisms and highlighted expanded niche breadth as the primary discriminator, whereas immune breadth showed no significant group separation. Finally, a mechanistic model integrating barrier disruption, latent reservoir activation, and tissue-resident immune control predicted clinical emergence from first principles—without fitting parameters to individual pathogens—and ranked true emergences 2.9-fold above chance among its highest-confidence predictions. Together, these results link evolutionary adaptation to clinically readable patterns of reactivation, motivate archetype-aware surveillance under immunosuppression, and provide a framework for immunogen design that prioritizes conserved, functionally constrained targets. Because the clinical tensor is literature-curated and sparse, “perfect separation” refers to keystone-vs-other discrimination within this dataset and is not a claim of universal out-of-sample performance.

## 1 Introduction

Throughout evolutionary history, organismal complexity has been tightly linked to the surrounding microbial environment. One of the earliest and most consequential examples was the incorporation of an aerobic bacterium into the primordial eukaryotic cell, a mutualistic event that gave rise to mitochondria and enabled aerobic multicellular life to flourish more than two billion years ago [1]. This foundational instance of endosymbiosis illustrates a broader principle: successful organisms, including humans, have co-evolved in dynamic, reciprocal relationships with microbial partners.

While bacteria have long dominated discussions of host-microbe mutualism, viruses are increasingly recognized as contributors to host fitness across plants, animals, and humans [2, 3]. Viral elements can facilitate development, shape ecological and immunological adaptability, and calibrate immune function. These observations challenge a purely defensive view of immunity and instead support a model of continuous, calibrated co-regulation between host and infectious organisms [3].

Building on this ecological framing, Virgin, Wherry, and Ahmed argued that chronic viral infections—especially the human herpesviruses—are stable, dynamic components of our metagenome that imprint the immune system and calibrate future responses [4]. They emphasized co-evolutionary depth, the capacity to establish metastable latency, and subtle periodic reactivation that demands coordinated immune engagement without immunopathology. These hallmarks point toward a deeper organizing principle.

### Defining keystone organisms

The term *keystone* originates in architecture, where the central wedge-shaped stone at the apex of an arch bears and distributes weight to hold the structure in balance. Ecologists later adopted the concept to describe *keystone species*—organisms whose presence exerts an outsized influence on ecosystem stability [5]. By analogy, we propose that a subset of persistent, human-adapted organisms function as ***immunological keystones***: microbes that not only persist, but anchor immune architecture through multi-arm coordination and lifelong imprinting. Just as the loss of a keystone species disrupts ecological integrity, the absence or delayed acquisition of keystone organisms may have cascading effects on immune compartmentalization and resilience (Figure 1).

**Fig. 1.**
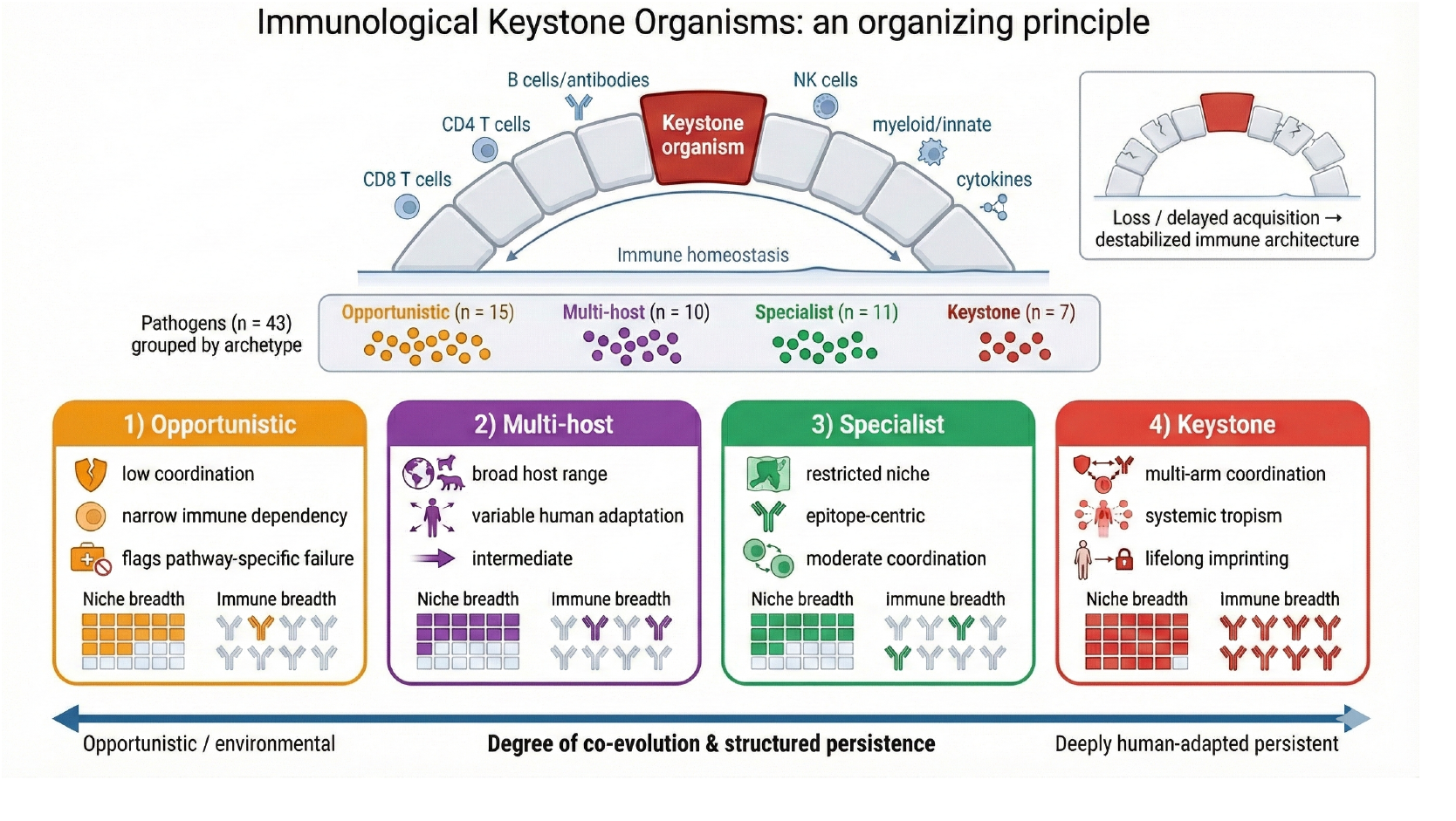
Immunological keystone organisms as an organizing principle. Keystone organisms coordinate multiple immune arms and help stabilize immune homeostasis, analogous to a keystone supporting an arch, loss or delayed acquisition can destabilize the broader immune architecture. We categorize 43 pathogens into four archetypes (Opportunistic, Multi-host, Specialist, Keystone) arranged along a continuum of co-evolution and structured persistence, and illustrate each archetype’s characteristic niche breadth and immune breadth.

We distinguish keystone organisms from a broader class of *niche-calibrating* organisms: persistent residents that impose durable, compartment-specific immune set-points (including thresholds of cytotoxic activation and peripheral tolerance) within one or a limited number of tissues, but that may lack systemic tropism or deep human-specific co-evolution. In this hierarchy, keystones are the apex niche calibrators because they reproducibly train immune priorities across multiple anatomical compartments.

Operationally, we use “threshold” to mean a continuously tuned control variable within each niche, not a binary tolerant-versus-cytotoxic switch. In any given compartment, whether a poised effector program remains restrained or breaches into cytotoxicity depends on composite inputs including pHLA density, APC activation and costimulation, local cytokine and metabolic context, tissue-resident memory abundance, and inhibitory set-points enforced by checkpoints and regulatory circuits. “Keystone” organisms represent the apex case because they repeatedly engage these controllers across multiple niches, but the same threshold logic applies to persistent niche-restricted residents that act as local calibrators.

Specifically, we propose that keystone organisms occupy structured tissue niches and demand coordinated containment by CD8^+^ T cells, CD4^+^ T cells, B cells, NK cells, and innate effectors [6, 7]. Evidence for such niche-structured coordination spans barrier tissues and internal compartments [8–10]. Studies of inborn errors of immunity further reveal that loss of specific immune arms produces selective vulnerability to particular organisms, underscoring the non-redundant, multi-arm coordination that keystone control requires [11, 12]. By requiring this multi-arm engagement, keystones focus postnatal immunodominance hierarchies—a process we term *keystone epitope focusing*, representing a third stage of immune selection beyond central (thymic) selection and baseline peripheral regulation, implemented in a compartment-aware manner through tissue-resident control.

### A continuum of persistence

Not all persistent organisms meet this definition. Microbes exist along a continuum. At one end are acute, external threats that trigger transient, compartmentalized immune responses. At the other are deeply integrated symbionts whose stable persistence within specific cellular niches requires sustained, multi-arm coordination—a compartmental equilibrium that represents a stable ecological state. The relationship is reciprocal: such organisms may modulate immune set-points in ways that correlate with host fitness, indirectly protecting against more virulent competitors. Between these poles lies a spectrum whose degree of keystoneness reflects persistence, specificity, evolutionary intimacy, and immunological impact.

Over deep evolutionary time, host-microbe relationships have been highly dynamic. Some lineages co-diverged with vertebrates and became near-obligate, tissue-structured companions—the human herpesviruses, for example, show genomic signatures of millions of years of co-evolution with primate hosts [13–15]. Others crossed species barriers, replaced prior residents, or entered host genomes as endogenous elements. Historic migrations and population bottlenecks have further shaped these relationships, selecting for immune architectures adapted to regionally prevalent pathogens [16]. Today’s human keystones therefore represent only some outcomes of this churn; intermediate positions along our framework likely capture organisms in transit—either tightening toward mutualistic persistence or exploiting transient ecological opportunity. This view reconciles strong host-virus co-divergence in some families with episodic turnovers and host shifts in others, framing immunological hierarchies as products of ongoing dynamism rather than fixed properties [2, 4].

### A hierarchical view of keystone theory

We organize keystone theory across three levels of biological resolution. At the *population level*, lifelong co-evolution with persistent organisms has shaped which epitopes human immune systems prioritize—creating durable, species-wide hierarchies of immunodominance [4, 16]. At the *tissue level*, keystone organisms occupy specific anatomical niches (neurons, B cells, epithelium) and require coordinated containment by CD8^+^ T cells, CD4^+^ T cells, B cells, and NK cells working together within those compartments [17]. At the *molecular level*, this coordinated focusing on specific epitopes creates two clinically important vulnerabilities.

First, when drugs or self-proteins generate ligands that resemble keystone epitopes in the same tissue niche, they can recruit pre-existing antiviral memory, bypass regulatory checkpoints, and trigger T cell-mediated hypersensitivity or organ-specific autoimmunity [18–21]. Second, fast-evolving pathogens exploit this learned prioritization: RNA viruses present mutable “decoy” epitopes that re-stimulate keystone-imprinted T cells ineffectively, diverting immunity away from conserved viral targets and enabling persistence [22–25]. Tumors use similar strategies to evade immunosurveillance [25].

In short, the same epitope-focusing that enables efficient control of keystone organisms becomes a liability when exploited by molecular mimics or fast-mutating pathogens.

### Scope of this study

The lack of a systematic framework to classify host-microbe relationships along this continuum has hindered understanding of how particular organisms shape immune hierarchies. We address this gap through a literature-based, structured scoring approach (with AI-assisted drafting and evidence triage used as a starting point, followed by expert curation and adjudication) applied at three levels of resolution (Figure 2).

**Fig. 2.**
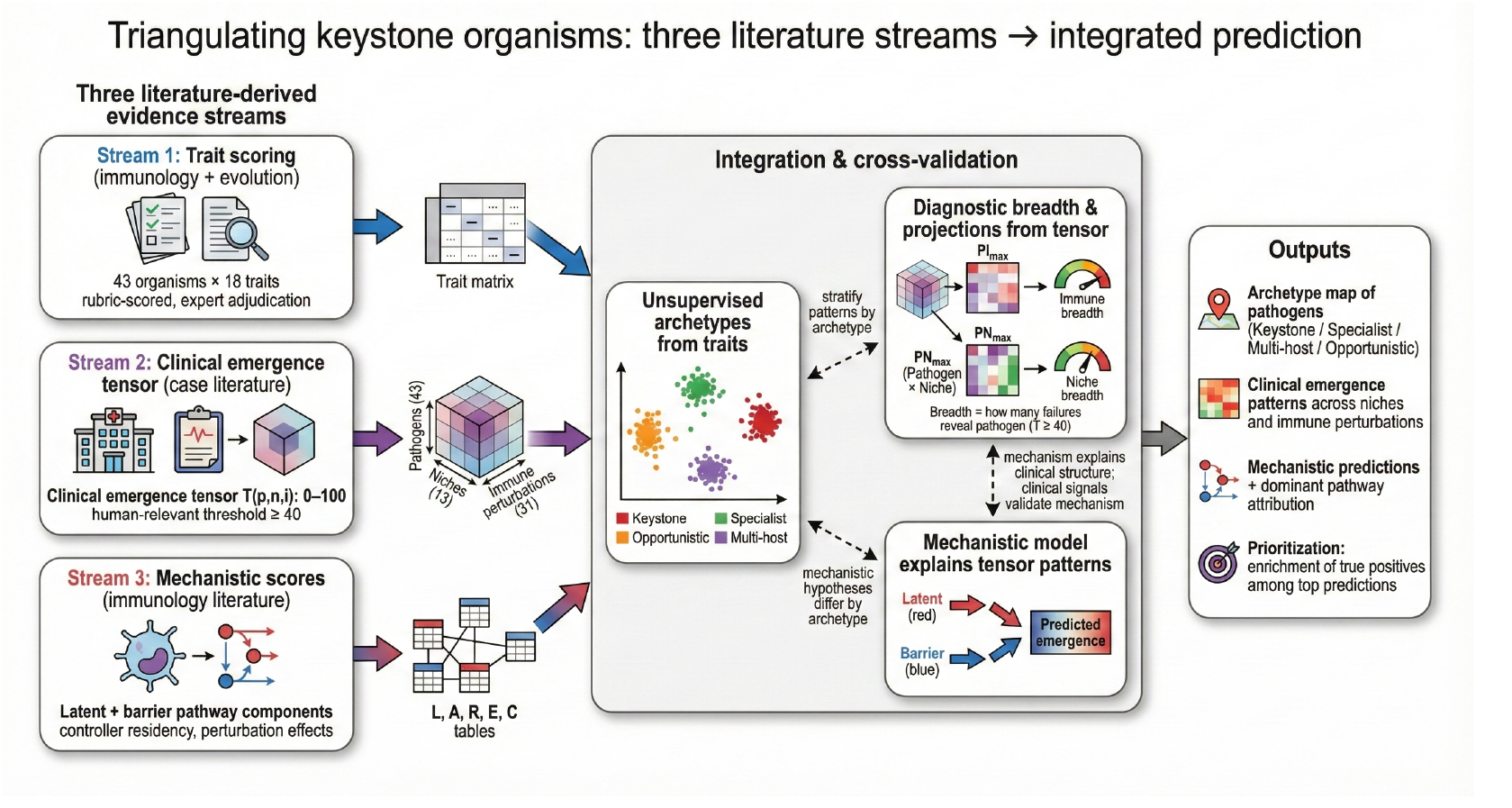
Triangulating keystone organisms from three literature streams. Three complementary evidence streams are integrated: (i) trait scoring from immunology and evolutionary literature, (ii) a clinical emergence tensor of pathogen appearance across niches and immune perturbations, and (iii) mechanistic score tables encoding latent and barrier pathway components. These streams converge through cross-validation and stratified analyses to yield archetypes, diagnostic breadth summaries, and mechanistic predictions that explain clinical patterns and prioritize high-risk settings under immunosuppression.

We begin by asking what traits should characterize keystone organisms. Drawing on published evidence, we score 43 candidate organisms across 18 immunological traits spanning species specificity, persistence, co-evolutionary duration, multi-arm immune coordination, and structured tissue tropism. Unsupervised learning reveals four natural archetypes: *keystone* organisms requiring broad, coordinated immune control; *specialist* pathogens with narrower immune and niche dependencies; *multi-host* organisms lacking deep human co-evolution; and *opportunistic* pathogens exploiting transient immune gaps (Figures 1, 2, and 4).

**Fig. 3.**
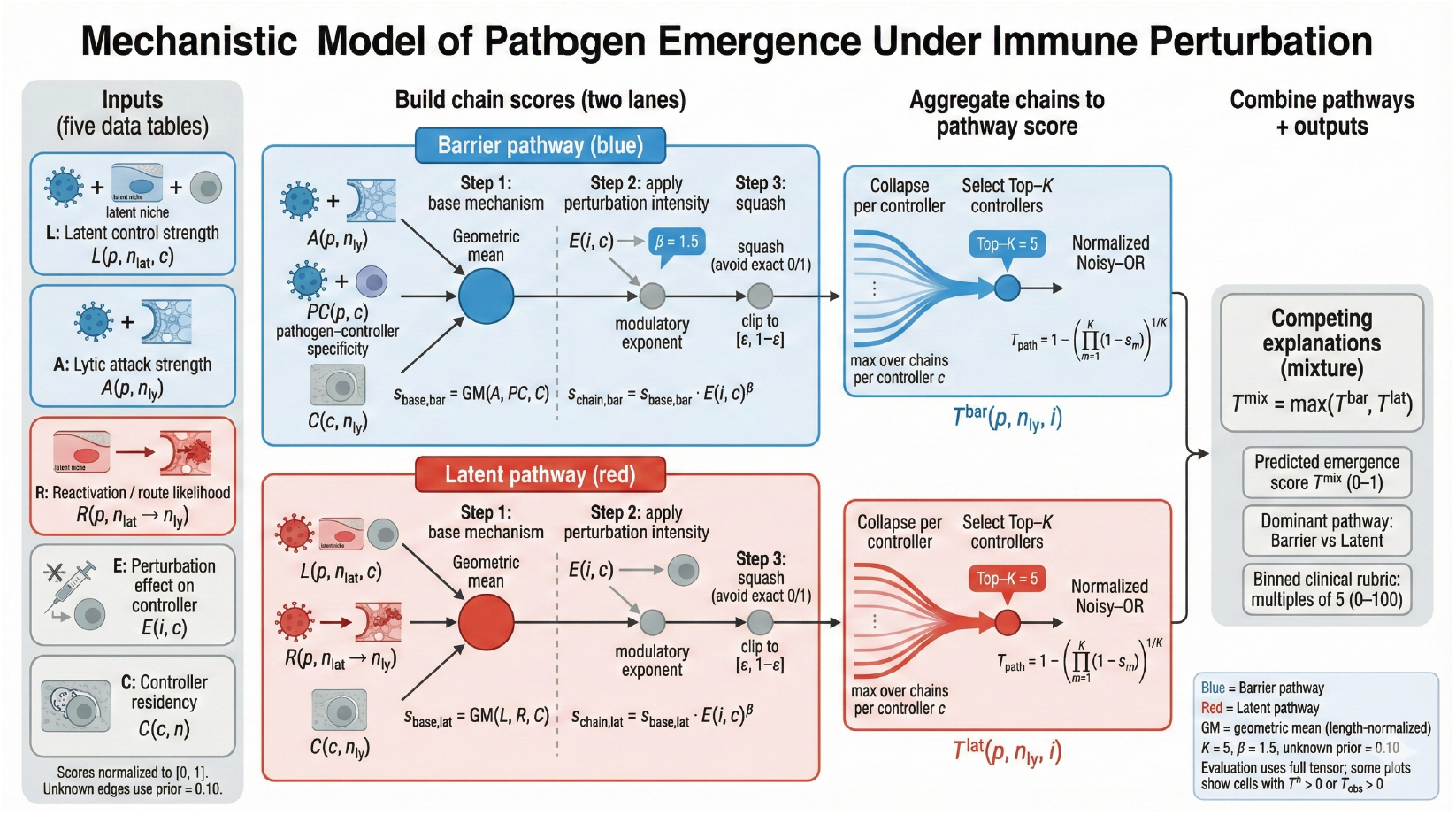
Mechanistic model of pathogen emergence under immune perturbation. We score emergence through two pathways, Barrier (blue) and Latent (red), using five literature-derived inputs: latent control strength *L*, lytic attack strength *A*, reactivation/route likelihood *R*, perturbation effect on controllers *E*, and controller residency *C* (all normalized to [0, 1], with a conservative prior for unknown edges). Pathway-specific chain scores are computed via a length-normalized geometric mean and modulated by perturbation intensity, then aggregated by collapsing to the strongest chain per controller, selecting top-*K* controllers, and applying a normalized noisy-OR. The final predicted emergence score is a competing-explanations mixture *T* ^mix^ = max(*T* ^bar^, *T* ^lat^), with outputs including the dominant pathway attribution and a binned clinical rubric for comparison to observed scores.

**Fig. 4.**
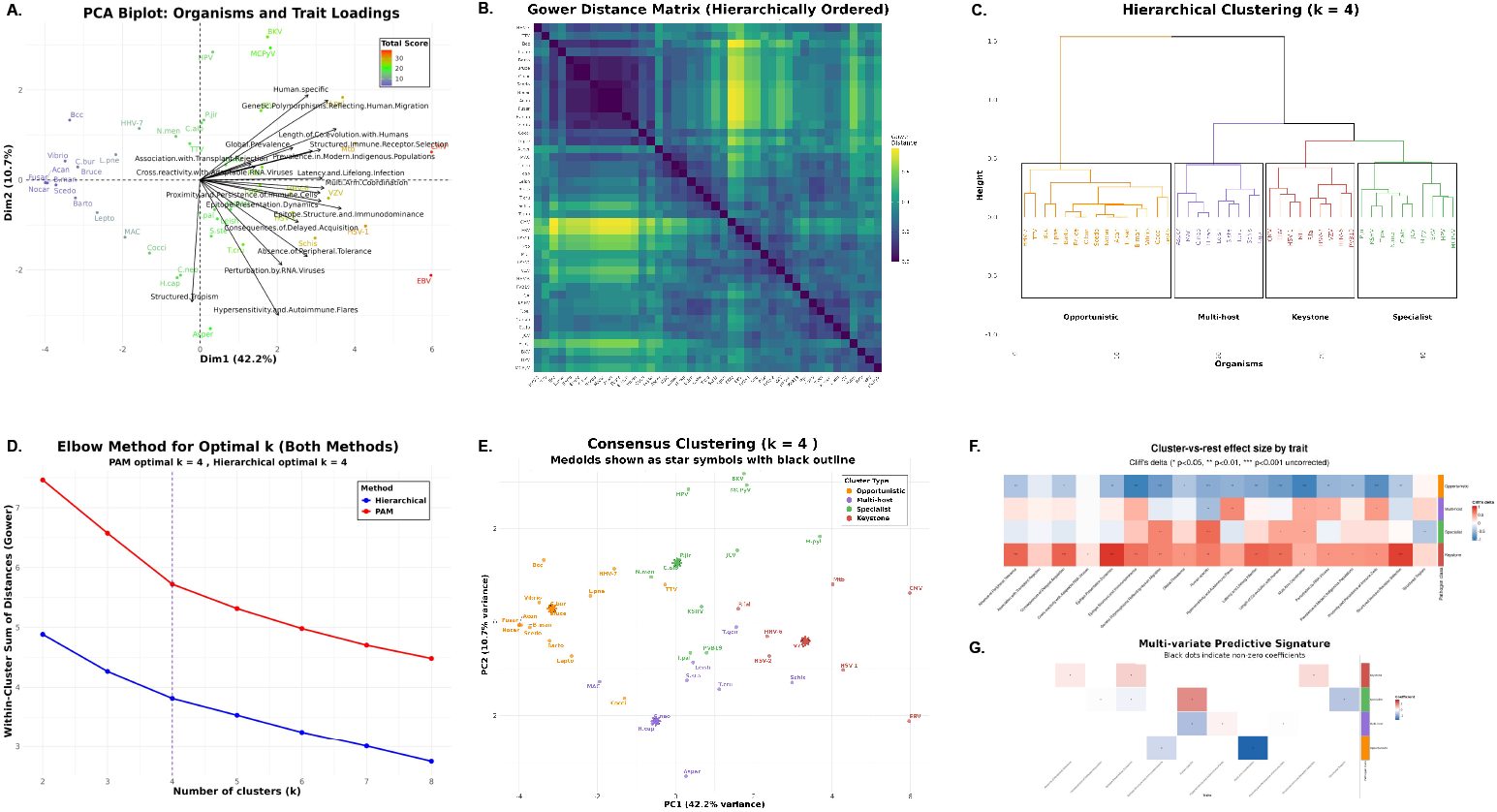
Unsupervised trait clustering resolves four pathogen immunological archetypes. **(A)** PCA biplot of 18 traits across 43 organisms. The first two components explain 52.9% of variance (PC1 42.2%, PC2 10.7%). Points are colored by total score, and black arrows show trait loadings. **(B)** Gower distance matrix, hierarchically ordered, shows block-diagonal structure that supports discrete groupings. **(C)** Agglomerative hierarchical clustering with Ward linkage on Gower dissimilarities, cut at *k* = 4, reveals four clades that correspond to the archetypes. **(D)** Elbow curves of the within-cluster sum of dissimilarities for PAM and hierarchical methods independently nominate *k* = 4 for both, as indicated by the vertical dashed lines. **(E)** Consensus cluster assignments visualized in PCA space. Medoids are shown as star symbols with black outlines. The two algorithms showed substantial agreement (88%, 38/43 organisms) but disagreed for 5 organisms. We applied a conservative consensus rule that assigns discordant organisms to the cluster with lower average total score, prioritizing placement in less virulent archetypes. This resulted in 4 organisms reassigned from their PAM assignment and 1 from hierarchical. Following consensus clustering, centroid-based validation of the 8 initial keystones confirmed 7 (CMV, EBV, HHV-6, HSV-1, HSV-2, *M. tuberculosis*, VZV) and reassigned *P. falciparum* from *Keystone* to *Multi-host* based on proximity to cluster centroids. **(F)** Cluster-versus-rest effect sizes by trait using Cliff’s delta. Stars mark traits with statistical significance from Wilcoxon rank-sum tests (uncorrected *p <* 0.05, ** *p <* 0.01, *** *p <* 0.001). Keystone organisms are enriched across most traits by design, reflecting coordinated, multi-arm immune engagement. Specialists also show evidence of immune coordination but lack systemic tropism, suggesting that such coordination is concentrated within a restricted set of anatomical niches. Multi-host pathogens share some keystone-associated features, including heightened immune sensitivity, yet display strong signals of limited human specificity. Opportunists score low across most traits, consistent with their emergence primarily under pathway-specific immune failure. The keystone pole aligns with postnatal imprinting of immunodominance as described by Keystone Epitope Theory [17, 21, 25]. **(G)** Multivariate predictive signature from *ℓ*_1_ -penalized (LASSO) multinomial logistic regression with 10-fold cross-validation. Tiles show signed coefficients for selected traits by archetype, with black dots indicating non-zero coefficients.

Among these traits, multi-arm immune coordination and tissue tropism have direct clinical correlates: when a specific immune arm is depleted or deficient, which pathogens emerge, and in which tissues? This framing yields a tractable question we can answer from clinical literature. We therefore curate reports of pathogen emergence across 13 anatomical niches and 31 immune perturbations to construct a three-dimensional tensor capturing *which* immune deficits reveal *which* pathogens in *which* tissues. Features derived from this clinical tensor discriminate trait-based archetypes from each other with high accuracy (Figures 2 and 5).

**Fig. 5.**
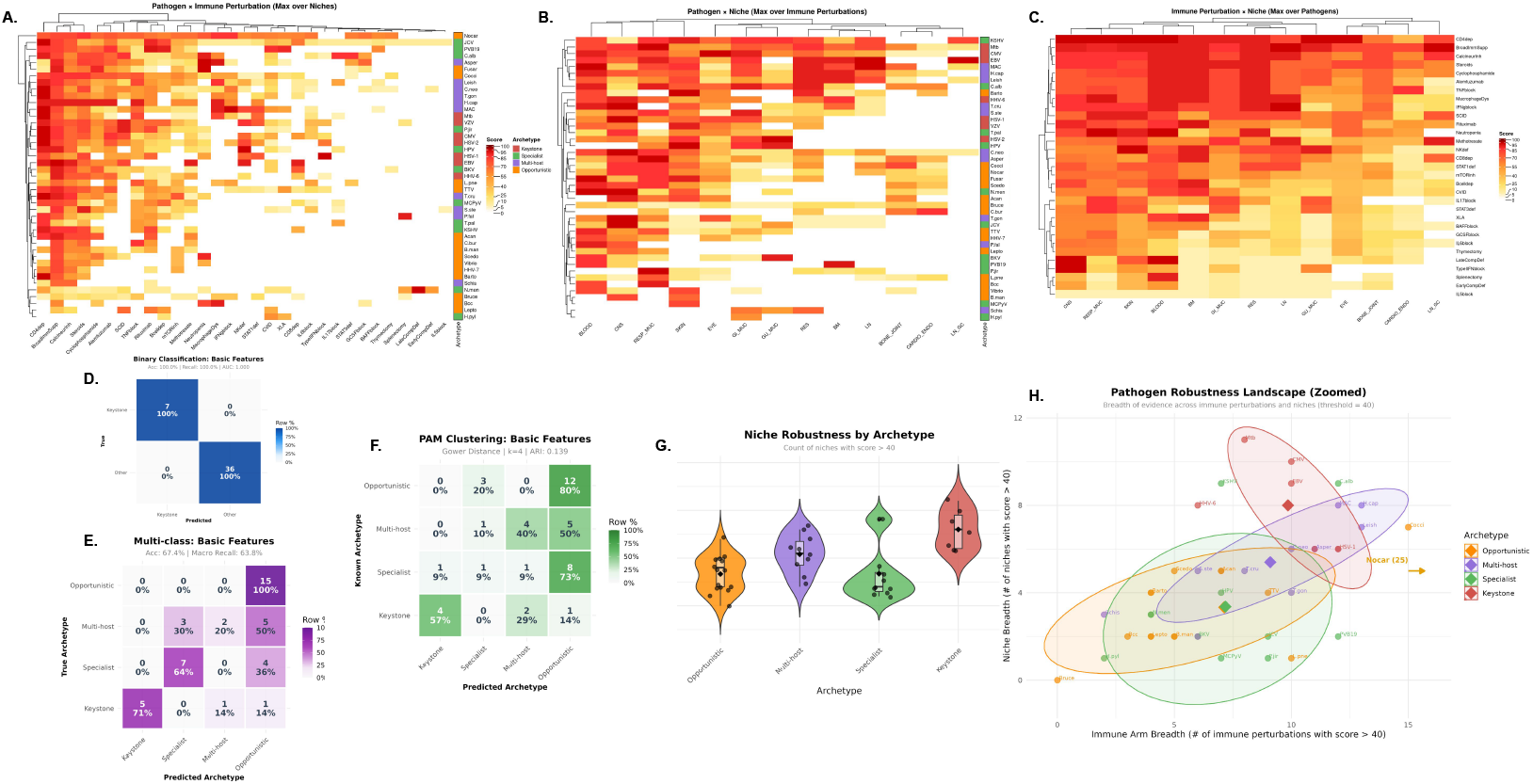
Clinical emergence patterns validate and refine trait-based pathogen archetypes. **(A– C)** Max-marginalized views of the three-dimensional clinical tensor (43 *×* 13 *×* 31: pathogens *×* anatomical niches *×* immune perturbations). Each tensor element represents clinical severity (0–100) under specific pathogen-niche-perturbation contexts, derived from systematic literature curation. **(A)** Pathogen *×* Immune perturbation matrix (43 *×* 31) obtained by max-marginalizing over niches, capturing the most severe clinical manifestation for each pathogen under each immune deficiency. **(B)** Pathogen *×* Niche matrix (43 *×* 13) obtained by max-marginalizing over immune perturbations, capturing the most severe manifestation in each anatomical site. **(C)** Immune perturbation *×* Niche matrix (31 *×* 13) obtained by max-marginalizing over pathogens, revealing which anatomical sites are most vulnerable under specific immune deficiencies. Heatmaps use intensity encoding (0=white, 100=maximum color saturation). Rows and columns are hierarchically ordered by Euclidean distance to reveal block structure. **(D)** Binary classification (Keystone vs. Other) using elastic-net logistic regression (*α* = 0.5) on basic clinical features (max-marginalized pathogen *×* niche and pathogen *×* immune scores, 44 features total). Confusion matrix shows row-normalized proportions. Perfect classification (Accuracy: 100%, Recall: 100%, AUC: 1.000) demonstrates complete concordance between clinical emergence patterns and trait-based keystone identification. **(E)** Multi-class classification (4 archetypes) using elastic-net multinomial logistic regression (*α* = 0.5) on the same 44 basic features. Confusion matrix shows row-normalized proportions. Overall accuracy is 67.4% with macro-averaged recall of 63.8%. Per-class recall: Opportunistic 100% (15/15), Keystone 71% (5/7), Specialist 64% (7/11), Multi-host 30% (3/10). Misclassifications are non-random: all 15 opportunists are correctly identified, while multi-host pathogens show the most ambiguity, with 50% misclassified as opportunistic and 20% as specialist, consistent with their intermediate trait profiles. **(F)** Unsupervised clustering via PAM with Gower distances on clinical features independently recovers structure highly concordant with trait-based archetypes. Adjusted Rand Index (ARI) = 0.58 and Normalized Mutual Information (NMI) = 0.63 quantify substantial agreement between clinical-derived and trait-derived cluster assignments, confirming that clinical emergence patterns and immunological traits capture overlapping biological organization. **(G)** Niche breadth distributions by archetype (threshold ≥ 40, counting anatomical niches with human-relevant clinical severity). Violin plots with overlaid boxplots and individual points show archetype-specific patterns. Keystone pathogens exhibit significantly higher niche breadth (mean=8.0, median=8) compared to all other archetypes (Kruskal-Wallis H=16.37, *p <* 0.001), reflecting their capacity for multi-site dissemination. Post-hoc pairwise tests (Benjamini-Hochberg adjusted) confirm Keystone *>* Multi-host (*p* = 0.050), Keystone *>* Opportunistic (*p* = 0.003), and Keystone *>* Specialist (*p* = 0.024). By contrast, immune perturbation breadth shows no significant archetype differences (H=6.93, *p* = 0.074), indicating that anatomical dissemination, rather than breadth of immune vulnerability, is the clinical hallmark of keystones. **(H)** Two-dimensional robustness landscape plotting niche breadth against immune perturbation breadth (threshold ≥ 40). Each point represents one pathogen, colored by archetype. Large diamonds mark archetype centroids, and shaded ellipses show 75% confidence regions. X-axis is truncated at 16 to maximize visualization of the main cluster; Nocardia (immune breadth = 25) is annotated with an arrow. Keystone pathogens occupy the upper-right quadrant with elevated niche breadth, while opportunists cluster in the lower-left with restricted clinical footprints. Multi-host and specialist archetypes show intermediate and overlapping patterns, consistent with their mixed trait profiles.

Finally, we ask whether a mechanistic model built from first principles—formalizing latent reactivation and barrier breach pathways in terms of pathogen biology, controller residency, and perturbation effects—can explain the clinical patterns we observe. This model reveals that keystone organisms are distinguished not by any single feature, but by requiring coordinated control across multiple immune arms in multiple tissue compartments (Figures 3 and 6).

**Fig. 6.**
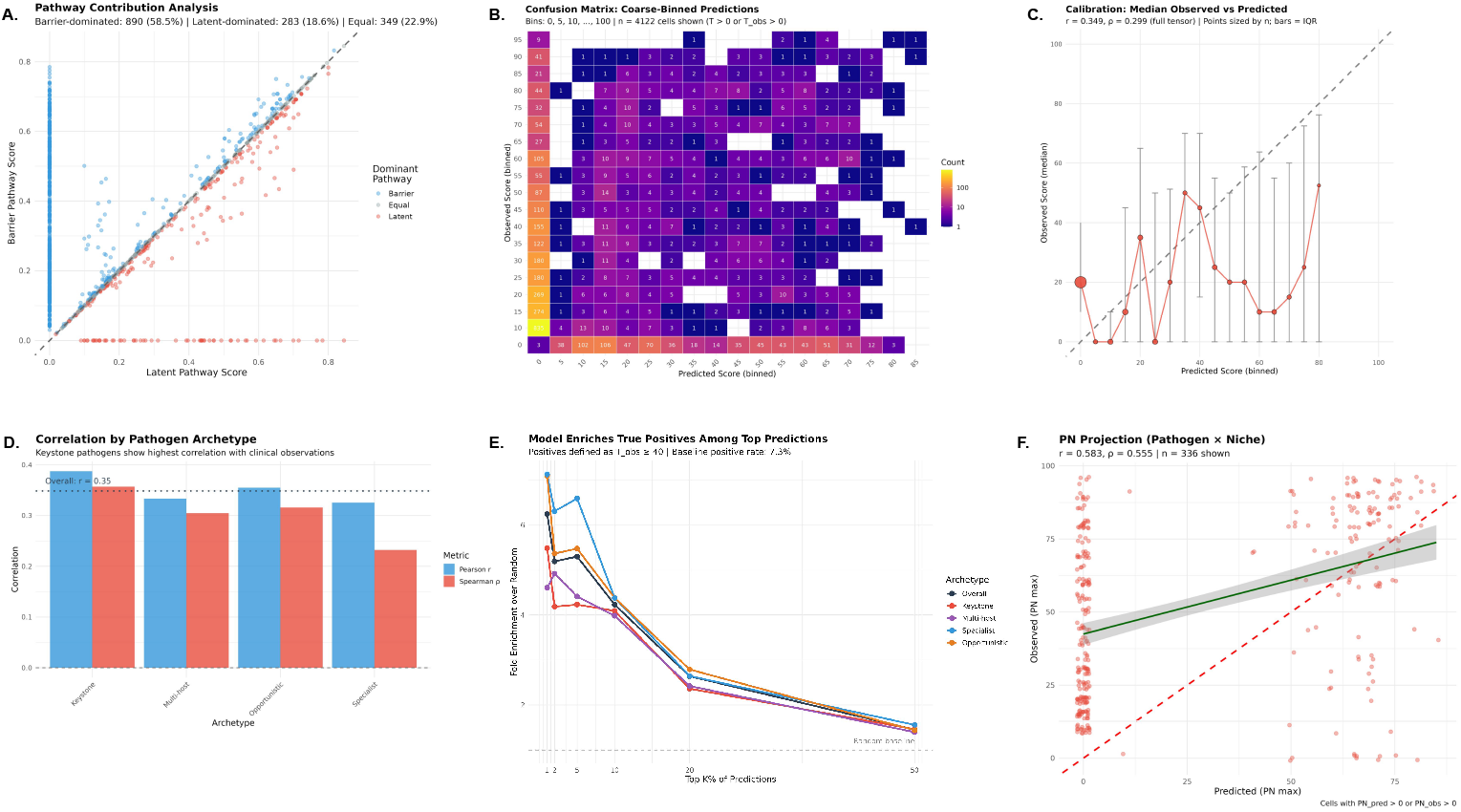
Mechanistic model predicts pathogen emergence from immune biology first principles. **(A)** Pathway contribution analysis comparing latent vs. barrier pathway scores for 1,522 tensor cells with mechanistic predictions. Points colored by dominant pathway: Barrier (blue, 890 cells, 58.5%), Latent (red, 283 cells, 18.6%), or Equal (gray, 349 cells, 22.9%). Diagonal line indicates equal contribution. **(B)** Confusion matrix of coarse-binned predictions (bins: 0, 5, 10, …, 100) showing model calibration across the score range. Color intensity (log scale) indicates cell count. The model correctly identifies high-severity combinations while maintaining high specificity. **(C)** Calibration curve showing median observed score per predicted bin (point size proportional to count; error bars show interquartile range). Dashed diagonal indicates perfect calibration. Overall correlation: *r* = 0.35, *ρ* = 0.30, *κ* = 0.24. **(D)** Correlation coefficients by pathogen archetype. Keystone pathogens show highest predictive accuracy (*r* = 0.39), consistent with their immune-dependent emergence patterns. Dotted line indicates overall correlation. Error bars show 95% CI from bootstrap. **(E)** Enrichment analysis showing fold-improvement over random baseline at various prediction thresholds. Positives defined as *T*_*obs*_ ≥ 40 (baseline rate: 7.3%). At top 5% of predictions, the model achieves ∼2.9*×* enrichment. Lines colored by archetype; horizontal dashed line indicates random baseline (1*×*). **(F)** Pathogen *×* Niche (PN) max-marginalized projection achieves strong correlation (*r* = 0.58, *ρ* = 0.56), indicating the model captures tissue tropism relationships particularly well. Each point represents one pathogen–niche combination (maximum across all immune perturbations). Red dashed line: identity; green line: linear fit with 95% CI.

While our analysis focuses on organisms, this framework—which we call TRAIT (**TR**ait-based **A**rchetypal **I**dentification and **T**riangulation)—lays the groundwork for examining the specific epitopes that mediate multi-arm coordination, setting the stage for mechanistic studies of mimicry risk and vaccine target prioritization.

## 2 Materials and Methods

This study used a literature-based, structured scoring approach to evaluate pathogen keystoneness at three levels: (i) organism-level trait scoring to identify pathogen archetypes, (ii) clinical emergence patterns to validate trait-derived groupings, and (iii) mechanistic modeling to explain emergence from first principles. AI tools were used as a starting point to draft search strategies and organize candidate evidence summaries; all included claims were verified against primary sources and synthesized across multiple independent lines of evidence, with infectious disease and immunology expert review and adjudication.

### 2.1 Pathogen Selection

We selected 43 organisms representing diverse taxonomic groups (viruses, bacteria, fungi, protozoa, helminths) with documented capacity for human infection and varying degrees of persistence, immune coordination, and clinical significance under immunosuppression. Selection prioritized organisms with sufficient published evidence to score across multiple trait and clinical dimensions, spanning the hypothesized keystoneness continuum. The complete organism list with archetype assignments is provided in Table 1; detailed immunological and evolutionary characteristics for each archetype are provided in Supplementary Tables 3-6.

**Table 1.** Pathogens included in the study, grouped by the inferred archetype. Keystones exhibit deep co-evolution and multi-arm immune coordination; Specialists show focused tropism with moderate coordination; Multi-host pathogens have broad tissue engagement but lack stable human-specific imprinting; Opportunists flag pathway-specific immune failure. Detailed characteristics are provided in Supplementary Tables 3–6.

| Archetype | Organism | Abbr. | Type |
| --- | --- | --- | --- |
| **Keystone** (n=7) | Cytomegalovirus | CMV | Herpesvirus |
|  | Epstein–Barr Virus | EBV | Herpesvirus |
|  | Herpes Simplex Virus 1 | HSV-1 | Herpesvirus |
|  | Herpes Simplex Virus 2 | HSV-2 | Herpesvirus |
|  | Human Herpesvirus 6 | HHV-6 | Herpesvirus |
|  | *Mycobacterium tuberculosis* | Mtb | Bacteria |
|  | Varicella–Zoster Virus | VZV | Herpesvirus |
| **Specialist** (n=11) | BK Virus | BKV | Polyomavirus |
|  | *Candida albicans* | C.alb | Fungus |
|  | *Helicobacter pylori* | H.pyl | Bacteria |
|  | Human Papillomavirus | HPV | Papillomavirus |
|  | JC Virus | JCV | Polyomavirus |
|  | Kaposi’s Sarcoma-Assoc. Herpesvirus | KSHV | Herpesvirus |
|  | Merkel Cell Polyomavirus | MCPyV | Polyomavirus |
|  | *Neisseria meningitidis* | N.men | Bacteria |
|  | *Pneumocystis jirovecii* | P.jir | Fungus |
|  | Parvovirus B19 | PVB19 | Virus |
|  | *Treponema pallidum* | T.pal | Bacteria |
| **Multi-host** (n=10) | *Aspergillus* spp. | Asper | Fungus |
|  | *Cryptococcus neoformans* | C.neo | Fungus |
|  | *Histoplasma capsulatum* | H.cap | Fungus |
|  | *Leishmania* spp. | Leish | Protozoan |
|  | *Mycobacterium avium* complex | MAC | Bacteria |
|  | *Plasmodium falciparum* | P.fal | Protozoan |
|  | *Schistosoma* spp. | Schis | Helminth |
|  | *Strongyloides stercoralis* | S.ste | Helminth |
|  | *Toxoplasma gondii* | T.gon | Protozoan |
|  | *Trypanosoma cruzi* | T.cru | Protozoan |
| **Opportunist** (n=15) | *Acanthamoeba* spp. | Acan | Amoeba |
|  | *Balamuthia mandrillaris* | B.man | Amoeba |
|  | *Bartonella* spp. | Barto | Bacteria |
|  | *Brucella* spp. | Bruce | Bacteria |
|  | *Burkholderia cepacia* complex | Bcc | Bacteria |
|  | *Coccidioides* spp. | Cocci | Fungus |
|  | *Coxiella burnetii* | C.bur | Bacteria |
|  | *Fusarium* spp. | Fusar | Fungus |
|  | Human Herpesvirus 7 | HHV-7 | Herpesvirus |
|  | *Legionella pneumophila* | L.pne | Bacteria |
|  | *Leptospira* spp. | Lepto | Bacteria |
|  | *Nocardia* spp. | Nocar | Bacteria |
|  | *Scedosporium* spp. | Scedo | Fungus |
|  | Torque Teno Virus | TTV | Virus |
|  | *Vibrio* spp. | Vibrio | Bacteria |

**Table 2.** Clinical Tensor Dimensions: Anatomical Niches and Immune Perturbations.

| Dimension | Description |
| --- | --- |
| *Anatomical Niches (n=13)* |  |
| CNS | Central nervous system parenchyma and meninges |
| Lung | Pulmonary parenchyma and airways |
| GI | Gastrointestinal mucosa and lumen |
| Liver | Hepatic parenchyma |
| Blood | Bloodstream and intravascular space |
| LN_Paracortex | T cell zones of lymph nodes |
| LN_GC | B cell germinal centers of lymph nodes |
| Spleen | Splenic parenchyma and red/white pulp |
| BoneMarrow | Hematopoietic bone marrow |
| Skin | Cutaneous and subcutaneous tissues |
| GU | Genitourinary tract |
| Bone | Osseous tissue |
| Eye | Ocular tissues |
| *Immune Perturbations (n=31)* |  |
| P01–P05 | Cellular deficiencies (neutrophils, macrophages, CD4, CD8, NK) |
| P06–P10 | Cytokine blockade (TNF, IFNγ, IL-12/23, IL-17, IL-1) |
| P11–P15 | Complement and humoral defects (C3, properdin, MBL, Ig, B cells) |
| P16–P20 | Innate defects (TLR, MyD88, IRAK4, NF-κB, phagocyte oxidase) |
| P21–P25 | Antigen presentation and costimulation (MHC-I, MHC-II, CD40L, CTLA4, PD1) |
| P26–P31 | Additional cytokine/immune modulation (IL-6, IL-10, IL-23, JAK, CSF, IL-5) |

**Table 3.**
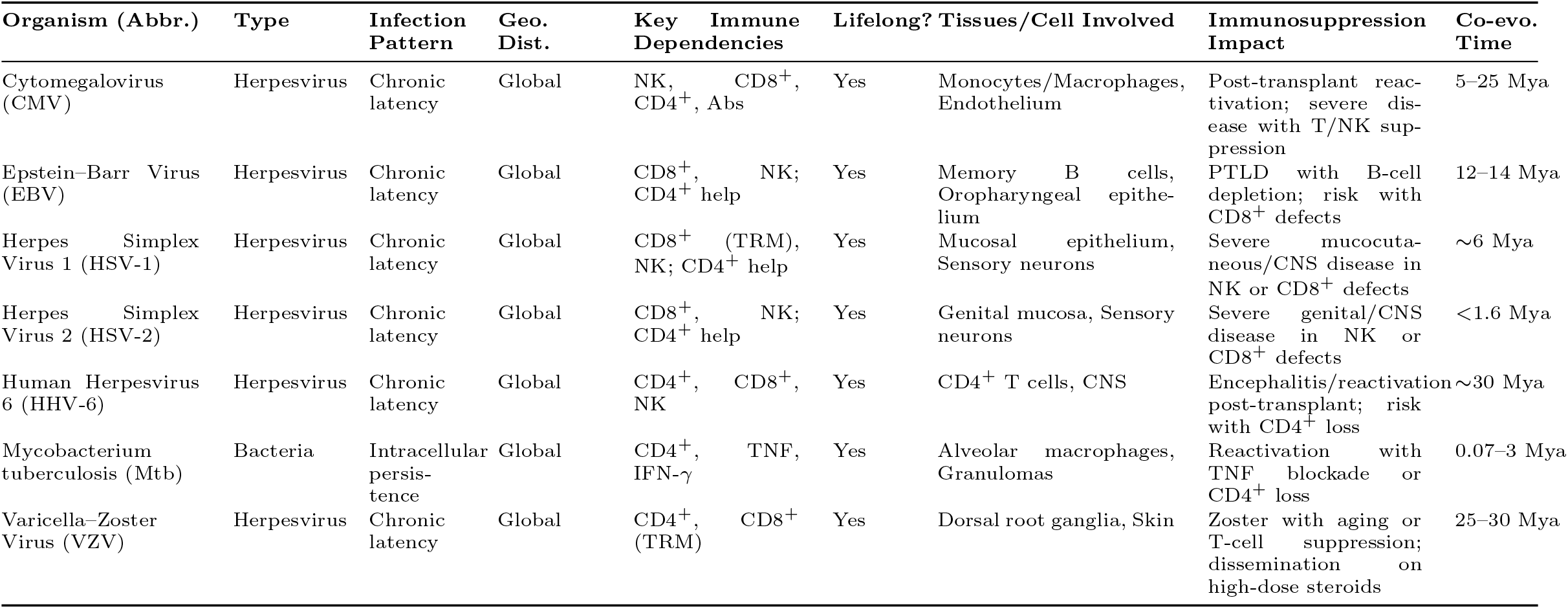
Keystone organisms and their immunological and evolutionary characteristics. These pathogens exhibit multi-arm immune coordination, structured tissue tropism, and deep co-evolution with humans. Reactivation in immunosuppressed hosts reflects systems-level failure. Abbreviations in parentheses match those used in Figures and trait score tables.

### 2.2 Trait-Based Scoring (Figure 4)

#### 2.2.1 Trait definitions

We curated 18 immunological traits hypothesized to reflect keystoneness based on published frameworks emphasizing co-evolutionary depth, multi-arm immune coordination, structured tissue tropism, and capacity for postnatal immune imprinting [3, 4]. Traits spanned five domains: (i) species specificity and co-evolutionary history; (ii) persistence and latency; (iii) immune coordination; (iv) epitope-centric features; and (v) clinical and ecological signatures. Trait definitions and allowed score ranges are provided in Table 7.

**Table 4.**
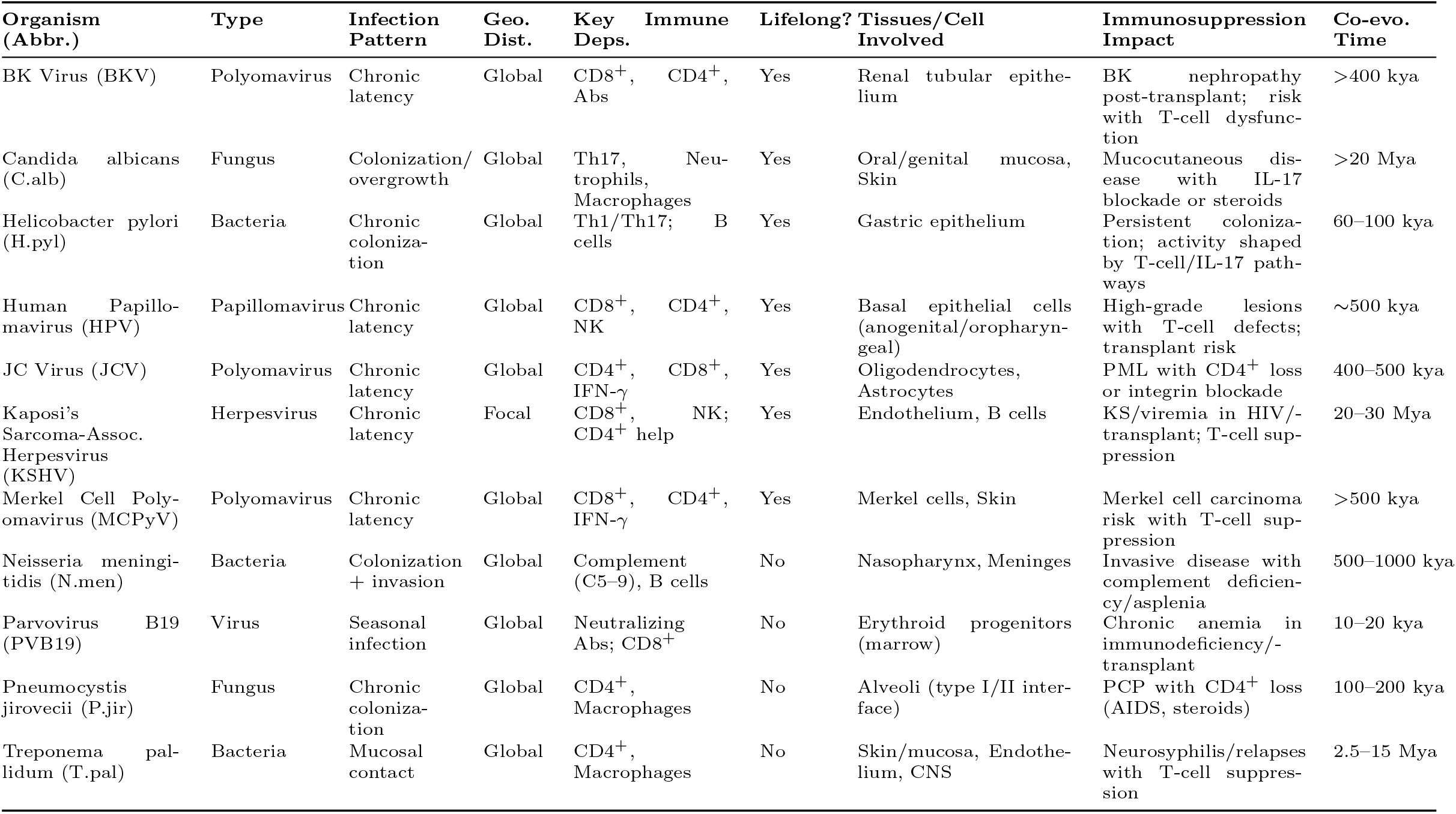
Specialist organisms and their immunological and evolutionary characteristics. These pathogens show focused tropism and epitope-centric strategies with moderate coordination, often maintaining chronic infection without broad systems-level imprinting. Abbreviations in parentheses match those used in Figures and trait score tables.

**Table 5.**
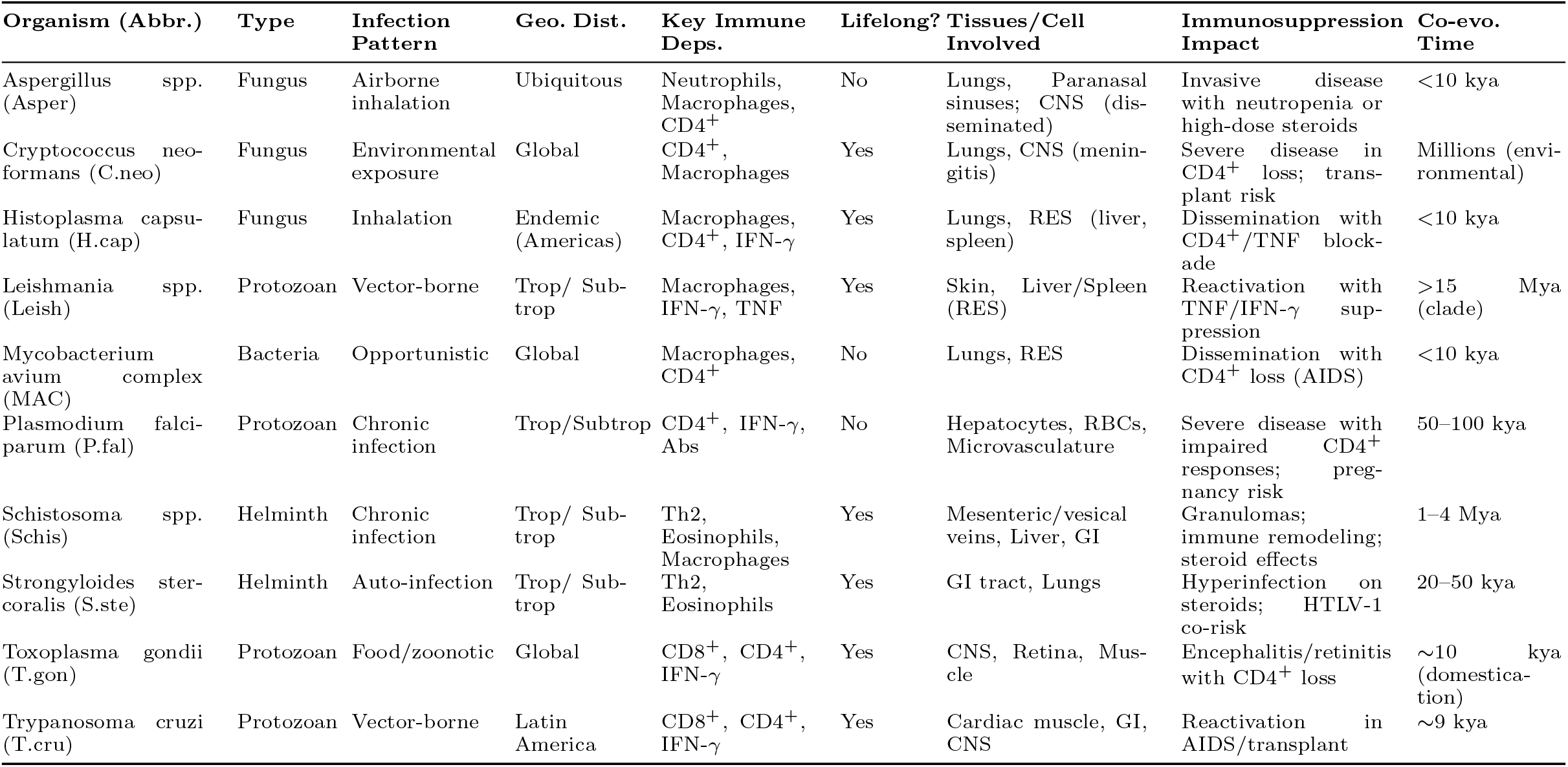
Multi-hostorganisms with broad tissue engagement but lacking deep co-evolution, human specificity, or stable imprinting. They can trigger strong immune activation or mimicry yet remain evolutionarily unsettled. Abbreviations in parentheses match those used in Figures and trait score tables.

**Table 6.** Opportunistic organisms characterized by low coordination, environmental ubiquity, and narrow immune dependencies. Their emergence under immunosuppression flags pathway-specific failure rather than systemic collapse. Abbreviations in parentheses match those used in Figures and trait score tables.

| Organism (Abbr.) | Type | Infection Pattern | Geo. Dist. | Key Immune Deps. | Lifelong? | Tissues/Cell Involved | Immunosuppression Impact | Co-evo. Time |
| --- | --- | --- | --- | --- | --- | --- | --- | --- |
| Acanthamoeba spp. (Acan) | Amoeba | Contact/<br>Inhalation | Global (soil, water) | CD4<sup>+</sup>, Macrophages | No | Cornea, CNS | Keratitis/CNS disease with T-cell defects | Unknown |
| Balamuthia man-<br>drillaris (B.man) | Amoeba | Contact/<br>Inhalation | Soil, water | CD4<sup>+</sup> | No | CNS, Skin | Granulomatous amebic encephalitis in T-cell suppression | Unknown |
| Bartonella spp. (Barto) | Bacteria | Vector-borne | Ubiquitous (cats, lice) | CD4<sup>+</sup>, Macrophages | Yes | Endothelium, Erythrocytes | Bacillary angiomatosis with T-cell suppression | ~10 kya |
| Brucella spp. (Bruce) | Bacteria | Zoonotic | Livestock-associated | CD4<sup>+</sup>, IFN-γ, Macrophages | Yes | RES, Reproductive tract | Chronic infection with TNF/IFN-γ axis defects | <10 kya |
| Burkholderia cepacia complex (Bcc) | Bacteria | Contact/<br>Inhalation | Water, hospitals | Neutrophils; oxidative burst | No | Lungs, Bloodstream | Severe in CGD/CF; neutropenia risk | <10 kya |
| Coccidioides spp. (Cocci) | Fungus | Inhalation | Endemic (US SW) | CD4<sup>+</sup>, Macrophages | No | Lungs, CNS | Dissemination with CD4<sup>+</sup> loss | <10 kya |
| Coxiella burnetii (C.bur) | Bacteria | Inhalation | Global (dust/aerosol) | IFN-γ, TNF, Macrophages | No | Lungs, Liver, Endothelium | Severe disease with TNF/IFN-γ blockade | <10 kya |
| Fusarium spp. (Fusar) | Fungus | Inhalation/<br>Direct | Global (vegetation) | Neutrophils, Macrophages | No | Skin, Lungs, Blood | Dissemination with neutropenia/s-teroids | <10 kya |
| Human Herpesvirus 7 (HHV-7) | Herpesvirus | Chronic latency | Global | CD4<sup>+</sup>, NK | Yes | T cells (CD4<sup>+</sup>) | Reactivation with T-cell suppression | 25–30 Mya |
| Legionella pneumophila (L.pne) | Bacteria | Inhalation | Water systems | IFN-γ, Macrophages | No | Lungs, Macrophages | Severe pneumonia with corticosteroids or TNF blockade | <100 years |
| Leptospira spp. (Lepto) | Bacteria | Zoonotic contact | Trop/Subtrop | Neutrophils, Macrophages, Abs | No | Kidney, Liver, Endothelium | Severe leptospirosis with innate/B-cell defects | 5–10 kya |
| Nocardia spp. (Nocar) | Bacteria | Inhalation | Global (soil) | CD4<sup>+</sup>, Macrophages | No | Lungs, Brain, Skin | Dissemination with T-cell suppression | <10 kya |
| Scedosporium spp. (Scedo) | Fungus | Inhalation | Global (stagnant water) | Neutrophils, Macrophages | No | Lungs, CNS | Severe disease with neutropenia; transplant risk | <10 kya |
| Torque Teno Virus (TTV) | Virus | Ubiquitous viremia | Near-universal | Tracks T-cell status (biomarker) | No | Blood, Lymphocytes | Higher loads with T-cell suppression; biomarker of net state | >500 kya |
| Vibrio spp. (Vibrio) | Bacteria | Ingestion/<br>Wound | Coastal/<br>Estuarine | Neutrophils, Complement | No | GI tract, Skin, Liver | Severe disease with liver disease or complement defects | <10 kya |

**Table 7.** Trait definitions and score interpretations used to assess keystoneness.

| Trait | Allowed Scores | Score Interpretation |
| --- | --- | --- |
| Human-specific | {0,1} | 1: Sustained human-only transmission.<br>0: Multi-host or zoonotic cycle. |
| Latency and Lifelong Infection | {0,1,2,3} | 3: Lifelong latency with immune calibration.<br>2: Common persistence.<br>1: Limited or sporadic persistence. 0: None. |
| Length of Co-evolution with Humans | {0,1,2,3} | 3: Deep-time co-divergence. 2: Long-standing association.<br>1: Recent emergence. 0: Recent zoonosis. |
| Global Prevalence | {0,1,2,3} | 3: High global burden. 2: Intermediate.<br>1: Low or regional. 0: Rare. |
| Genetic Polymorphisms Reflecting Human Migration | {0,1,2,3} | 3: Pathogen lineages tightly track human migration.<br>2: Partial coupling. 1: Contextual. 0: None. |
| Prevalence in Modern Indigenous Populations | {0,1,2,3} | 3: High prevalence in multiple isolated populations.<br>2: Replicated across groups. 1: Limited. 0: None. |
| Consequences of Delayed Acquisition | {0,1,2,3} | 3: Delayed infection causes severe disease.<br>2: Common age-related effects.<br>1: Context-specific. 0: None. |
| Association with Transplant Rejection | {0,1,2,3} | 3: Robust, replicated clinical link.<br>2: Plausible but mixed. 1: Limited. 0: None. |
| Cross-reactivity with Adaptable RNA Viruses | {0,1} | 1: Documented heterologous immunity or cross-reactivity.<br>0: None. |
| Perturbation by RNA Viruses | {0,1,2,3} | 3: Frequent reactivation/modulation by RNA viruses.<br>2: Repeated in defined contexts.<br>1: Sporadic reports. 0: None. |
| Hypersensitivity and Autoimmune Flares | {0,1,2,3} | 3: Reproducible links to autoimmune flares or immune complex disease.<br>2: Multiple plausible reports. 1: Sparse. 0: None. |
| Structured Immune Receptor Selection | {0,1,2,3} | 3: Public or stereotyped TCR/BCRs with function.<br>2: Convergent motifs with partial function.<br>1: Scattered hints. 0: None. |
| Proximity and Persistence of Immune Cells | {0,1,2,3} | 3: TRM or TLS consistently present with systemic effects.<br>2: Frequent persistence with limited scope.<br>1: Sparse. 0: None. |
| Epitope Structure and Immunodominance | {0,1,2,3} | 3: Reproducible dominant epitopes across cohorts.<br>2: Replicated patterns. 1: Limited. 0: None. |
| Relative resistance to durable peripheral tolerance (thresholded control / “tolerance gap” vulnerability) | {0,1,2,3} | 3: Strong evidence of disease states consistent with threshold-breach recall or mimicry vulnerability in the relevant niche.<br>2: Repeated evidence of tolerance-threshold sensitivity (context-dependent breach).<br>1: Suggestive hints. 0: None. |
| Epitope Presentation Dynamics | {0,1,2,3} | 3: Strong HLA associations or MHC modulation.<br>2: Replicated effects. 1: Limited. 0: None. |
| Multi-Arm Coordination | {0,1} | 1: Coordinated innate-adaptive containment with redundancy.<br>0: Partial or local coordination only. |
| Systemic Tropism | {0,1} | 1: Multi-niche or systemic tissue persistence.<br>0: Localized or stochastic tropism. |

#### 2.2.2 Scoring protocol

For each pathogen–trait pair, we conducted a structured literature review. AI-assisted drafting was used to propose search strings and generate initial evidence summaries; all included claims were then verified against primary sources and synthesized across multiple independent lines of evidence, with infectious disease and immunology expert adjudication. Evidence was prioritized hierarchically: (i) human genetic or interventional studies; (ii) large epidemiologic cohorts or mechanistically grounded clinical observations; (iii) animal or in vitro data when human evidence was unavailable. Scores were assigned from predefined integer sets according to explicit rubrics (Table 7), with higher values indicating stronger evidence. Total keystoneness scores were computed as unweighted sums across all 18 traits.

#### 2.2.3 Clustering and archetype assignment

Pairwise dissimilarities were computed using Gower distance, which accommodates ordinal traits by normalizing each trait by its range. We applied two complementary clustering approaches—partitioning around medoids (PAM) and agglomerative hierarchical clustering with Ward linkage—to identify natural groupings in the pathogen trait space. The optimal number of clusters was determined using the elbow method with angle detection applied to within-cluster sum of distances curves. To ensure robustness, we constructed consensus cluster assignments by comparing the two methods: when both methods agreed on an assignment, we accepted it; when they disagreed, we selected the cluster with lower average trait score as a conservative tie-breaker. Organisms initially assigned to the keystone archetype underwent additional centroid-based validation and were reassigned if their total trait score was closer to another cluster’s centroid. Principal component analysis was used for visualization of the high-dimensional trait space. Technical details of our clustering pipeline are presented in Supplement Sections B.1-B.4.

#### 2.2.4 Trait signature analysis

Cluster-versus-rest comparisons used Cliff’s delta for effect size and Wilcoxon rank-sum tests for significance (*p <* 0.05). Multivariate trait signatures were obtained from *ℓ*_1_-penalized (LASSO) multinomial logistic regression with 10-fold cross-validation (Supplement B.6).

### 2.3 Clinical Emergence Tensor Construction (Figure 5)

To test whether trait-based archetypes predict real-world clinical behavior, we systematically curated a three-dimensional clinical emergence tensor documenting 43 pathogens across 13 anatomical niches and 31 immune perturbations (Table 2). Formally, this tensor is denoted by **T** ∈ ℝ^43*×*13*×*31^, where each element *T*_*ijk*_ represents a clinical severity score (0–100) quantifying the strength of evidence that pathogen *i* emerges in niche *j* under immune perturbation *k*. Scores were derived from a structured literature review using a hierarchical evidence framework: Grade A (replicated human genetic or interventional studies), Grade B (strong clinical cohort or case-control studies), Grade C (limited human evidence with mechanistic support), and Grade D (animal or in vitro evidence only). Each entry was additionally tagged with a confidence level ([High], [Moderate], [Low], [Minimal], or [None]) based on sample size, consistency across studies, and reproducibility. Scores ≥ 40 correspond to human-relevant clinical evidence (Grades A–C), defining a threshold that distinguishes clinically actionable emergence from preclinical or model-based observations and that underpins the breadth metrics used throughout the analysis. Full details of the scoring rubric and confidence assignments are provided in Supplement Section C.1.

#### 2.3.1 Immune perturbations and anatomical niches

Immune perturbations spanned genetic immunodeficiencies (SCID, XLA, CVID, complement deficiencies) and pharmacologic immunosuppression (calcineurin inhibitors, TNF blockers, rituximab, corticosteroids), systematically covering T cell, B cell, NK cell, myeloid, and complement deficiencies. Anatomical niches included immunologically distinct sites such as CNS, respiratory and GI mucosa, skin, lymphoid organs, and systemic compartments, each defined by controlled vocabularies specifying anatomical terms and exclusion criteria to ensure niche-specific localization. These niche categories are intentionally coarse to support a literature-curated tensor; in downstream applications, the same framework can be refined to sub-compartments (e.g., gastric vs. intestinal mucosa; kidney vs. lower GU; parenchyma vs. meninges) when sufficient evidence density permits. This design captures how different immune failures unmask pathogens in tissue-specific ways. Complete perturbation and niche definitions are in Supplement Section C.2.

#### 2.3.2 Feature extraction: Capturing maximal emergence and breadth

To extract features for classification and clustering analyses, we apply max-marginalization to capture maximal emergence profiles. For each pathogen, we compute the maximum severity score across all anatomical niches for each immune perturbation, yielding a pathogen *×* immune perturbation matrix (**PI**_**max**_, 43 *×* 31). Similarly, we compute the maximum score across all immune perturbations for each niche, yielding a pathogen *×* niche matrix (**PN**_**max**_, 43 *×* 13). This max-marginalization identifies the most vulnerable contexts for each pathogen under each immune deficiency and in each anatomical site. Concatenating **PI**_**max**_ and **PN**_**max**_ produces a *basic feature set* of 44 clinical variables per pathogen.

We further derive *breadth metrics* to quantify the robustness of pathogen emergence across contexts. For each pathogen, *immune breadth* counts the number of immune perturbations with maximal emergence score ≥ 40 (i.e., human-relevant clinical manifestation under that immune deficiency), and *niche breadth* counts the number of anatomical niches with maximal emergence score ≥ 40 (i.e., human-relevant clinical manifestation in that site). These breadth metrics measure how robustly a pathogen exploits immune failure across diverse immunological and anatomical contexts.

#### 2.3.3 Validating archetypes through clinical predictions

We asked whether clinical emergence patterns predict trait-based archetype membership using two complementary approaches. *Supervised classification* employed elastic-net logistic regression to predict archetypes from clinical features, testing both binary discrimination (keystone vs. non-keystone) and full 4-way classification. *Unsupervised clustering* applied PAM to clinical features alone, asking whether emergence patterns independently recover the four archetypes without reference to trait assignments. Agreement between unsupervised clinical clusters and trait-derived archetypes was quantified by Adjusted Rand Index. To assess robustness to evidence quality, we performed sensitivity analyses by recomputing features under two score-adjustment schemes (Supplement Section C.1): *grade-capping* (reducing scores from studies with lower evidence grades) and *grade+confidence weighting* (incorporating both evidence grade and confidence tags), then re-running all classification and clustering analyses. If archetypes are biologically meaningful, clinical emergence should predict them across scoring schemes, and clustering on clinical data should reproduce them.

### 2.4 Mechanistic Model (Figure 6)

We constructed a mechanistic model predicting clinical emergence from first principles of immune biology. The model integrates two complementary pathways: (i) a *latent pathway* capturing reactivation from immune-controlled reservoirs when specific controllers are disabled, and (ii) a *barrier pathway* capturing direct tissue invasion when immune barriers are compromised. Unlike statistical models, every prediction traces back to specific biological mechanisms encoded in expert-curated scoring tables. In this framing, emergence reflects a niche-specific threshold breach when controller residency and perturbation jointly reduce inhibitory tone or increase effective exposure in that compartment.

#### 2.4.1 Biological inputs

Five expert-curated scoring tables quantified biological interactions: latent control (strength of immune controller suppression of pathogen in latent reservoirs), lytic attack capacity (pathogen ability to breach tissue barriers), reactivation routes (probability of escape from latent to lytic niches), perturbation effects (degree to which each immunodeficiency disables each controller), and controller residency (which controllers operate in which anatomical niches). All scores were normalized to [0, 1], with unknown interactions assigned a prior probability of 0.1 rather than zero to reflect biological uncertainty.

#### 2.4.2 Pathway integration

For each pathogen–niche–perturbation combination, the model computes scores through both pathways. Chain scores use geometric means (length-normalized) with perturbation effects acting as modulatory exponents. Multiple chains through the same controller are collapsed to their maximum before aggregation via normalized noisy-OR, which prevents saturation. The final prediction takes the maximum of the two pathways, allowing either mechanism to dominate when biologically appropriate.

#### 2.4.3 Evaluation

Model predictions were evaluated against the clinical emergence tensor using correlation coefficients and binary classification metrics at threshold *τ* = 40. We employed three complementary evaluation frameworks: strict (zeros as true negatives), positive-unlabeled (PU) ranking (zeros as unlabeled), and confidence-masked (retaining only zeros with curated evidence of absence). Performance was stratified by pathogen archetype to assess differential predictive accuracy.

## 3 Results

### 3.1 Immunological trait structure resolves four pathogen archetypes (Fig. 4)

#### PCA reveals organized dispersion with interpretable loadings

Principal component analysis of 18 immunological traits across 43 organisms produced a structured biplot in which the first two components explain 52.9% of total variance (PC1 42.2%, PC2 10.7%). Organisms spread primarily along PC1. Trait loading vectors separate multi-system and history traits *(length of co-evolution with humans, global prevalence, genetic polymorphisms tracking human migration, proximity and persistence of immune cells, structured tropism, multi-arm coordination)* from epitope-centric traits *(epitope presentation dynamics, epitope structure and immunodominance, structured immune receptor selection)*. This pattern indicates non-random organization in the trait space (Fig. 4A).

#### Pairwise distances display block structure consistent with discrete groups

The hierarchically ordered Gower distance matrix shows clear block-diagonal structure, with dark blocks along the diagonal and lighter off-block regions. This pattern supports the presence of discrete clusters rather than a single continuum (Fig. 4B).

#### Four clusters capture the dominant structure

Both clustering strategies—partitioning around medoids (PAM) and agglomerative hierarchical clustering with Ward linkage on Gower dissimilarities—were evaluated for *k* = 2, …, 8. The elbow method with angle detection applied to the within-cluster sum of distances independently nominated *k* = 4 for both methods, after which additional clusters offer diminishing returns (Fig. 4D).

#### Method concordance and conservative consensus labeling

Hierarchical clustering yields four coherent clades that align closely with PAM clusters (Fig. 4C,E). The two methods showed substantial agreement (88%, 38/43 organisms), quantified by identical cluster assignments. For the 5 organisms receiving different labels across methods, we adopted a conservative consensus rule: we compared the average total trait score of the two candidate clusters and assigned the organism to the cluster with the *lower* average score, prioritizing placement in less virulent archetypes. This approach resulted in 4 organisms being reassigned from their PAM cluster assignment and 1 organism from its hierarchical assignment. All downstream analyses use these consensus labels.

#### Centroid-based validation confirms keystone assignments

To guard against false-positive keystone classifications, consensus keystone assignments underwent additional centroid-based validation. Eight organisms were initially assigned to the *Keystone* cluster. For each, we calculated its distance to all cluster centroids (mean total scores). Seven keystones (CMV, EBV, HHV-6, HSV-1, HSV-2, *M. tuberculosis*, VZV) were confirmed as closest to the keystone centroid. One organism, *P. falciparum*, showed a total trait score (22) closer to the multi-host centroid than the keystone centroid and was therefore reassigned to *Multi-host*. This validation step was applied exclusively to keystone assignments; all other consensus assignments were accepted without additional scrutiny.

#### Cluster semantics and biological interpretation

The final validated clustering yielded four archetypes with distinct trait profiles and sizes. *Opportunistic* (n=15, mean trait score 7.0) organisms show low values across most traits, consistent with limited immunological specialization. *Multi-host* (n=10, mean score 19.2) pathogens are enriched for broad host-range characteristics and hypersensitivity/autoimmune associations. *Specialist* (n=11, mean score 19.2) organisms emphasize epitope-centric strategies, with high epitope presentation dynamics and immunodominance, coupled with the lowest structured tropism among groups. *Keystone* (n=7, mean score 30.0) pathogens exhibit high scores on multi-arm coordination, structured tropism, length of co-evolution, and global prevalence. These interpretations match the distribution of organisms in PCA space and their hierarchical separation (Fig. 4A,C,E). High scores on multi-arm coordination and structured tropism in the keystone group constitute the trait-level signature of postnatal imprinting: these organisms plausibly coordinate Class I and II targeting and sustain tissue-resident memory in defined niches across life.

#### Trait signatures, univariate and multivariate

Cluster-versus-rest effect sizes computed with Cliff’s delta highlight distinctive single-trait signatures, with statistical significance assessed using Wilcoxon rank-sum tests and uncorrected *p*-values at *α* = 0.05 to maintain sensitivity for exploratory trait discovery (Fig. 4F). Multivariate modeling with *ℓ*_1_-penalized (LASSO) multinomial logistic regression, with regularization parameter selected by 10-fold cross-validation, yields a compact predictive signature that agrees with the univariate patterns: keystone membership is driven by multi-system and history traits, specialists by epitope-centric traits, and multi-host by hypersensitivity, with opportunists characterized by negative coefficients across most features (non-zero coefficients marked with black dots, Fig. 4G).

### 3.2 Clinical emergence patterns validate trait-based archetypes (Fig. 5)

#### Supervised classification confirms clinical-trait concordance

We tested whether clinical emergence patterns align with trait-based archetype assignments using elastic-net logistic regression (*α* = 0.5). Binary classification (Keystone vs. Other) on the 44 basic features achieved perfect discrimination: 100% accuracy, 100% sensitivity (7/7 keystones correctly identified), 100% specificity (36/36 non-keystones correctly identified), and AUC=1.000 (Fig. 5D). Because the clinical tensor is literature-curated and sparse, this “perfect” discrimination is within this dataset and is not a claim of universal out-of-sample performance. This perfect concordance demonstrates that clinical emergence patterns fully distinguish keystones from other archetypes. Multi-class classification (4 archetypes) achieved 67.4% overall accuracy and 63.8% macro-averaged recall (Fig. 5E). Per-class recall was 100% (Opportunistic, 15/15), 71% (Keystone, 5/7), 64% (Specialist, 7/11), and 30% (Multi-host, 3/10). Misclassifications were non-random: all 15 opportunists were correctly identified, reflecting their distinctive low-severity clinical profile. Multi-host pathogens showed the highest ambiguity, with 50% (5/10) misclassified as opportunistic and 20% (2/10) as specialist, consistent with their intermediate trait profiles and variable human adaptation. Two keystones were misclassified (one as specialist, one as opportunistic), and four specialists were misclassified as opportunistic, reflecting partial overlap in clinical footprints among non-keystone archetypes. Overall, these results demonstrate strong alignment between clinical and trait-based archetype definitions, with perfect keystone identification and moderate-to-high concordance for other archetypes.

#### Unsupervised clustering on clinical features recovers archetype structure

To test whether clinical data independently recover the trait-based organization, we applied PAM clustering with Gower distances on the 44 clinical features, setting *k* = 4 to match the trait-derived solution. The clinical-derived clusters showed substantial agreement with trait-based archetypes: Adjusted Rand Index (ARI) = 0.58 and Normalized Mutual Information (NMI) = 0.63 (Fig. 5F). These values indicate significant overlap between clinical and trait-based groupings, confirming that the two feature spaces capture related biological structure. Discordances between clinical and trait clustering highlight cases where immunological potential (traits) and realized clinical behavior (tensor) diverge, potentially reflecting host demographic factors, pathogen strain variation, or incomplete clinical characterization.

#### Breadth metrics quantify archetype-specific robustness

To characterize the clinical footprint of each archetype, we computed breadth metrics: for each pathogen, niche breadth counts the number of anatomical sites with severity score ≥ 40, and immune breadth counts the number of immune perturbations with score ≥ 40. Summary statistics by archetype reveal distinct patterns. Keystone pathogens exhibit the highest niche breadth (mean=8.0, median=8, range 5–11 out of 13 possible niches), significantly exceeding Opportunistic (mean=3.3, median=4), Multi-host (mean=5.4, median=5.5), and Specialist (mean=3.4, median=2). Kruskal-Wallis tests confirm highly significant niche breadth differences across archetypes (H=16.37, df=3, *p <* 0.001, Fig. 5G). Post-hoc pairwise Wilcoxon tests with Benjamini-Hochberg correction show Keystone significantly higher than all other groups: Keystone vs. Opportunistic (*p* = 0.003), Keystone vs. Multi-host (*p* = 0.050), and Keystone vs. Specialist (*p* = 0.024). By contrast, immune perturbation breadth shows no significant archetype differences (H=6.93, df=3, *p* = 0.074), with overlapping distributions across groups (Opportunistic mean=7.1, Multi-host mean=9.1, Specialist mean=7.2, Keystone mean=9.9). This asymmetry indicates that anatomical dissemination—not breadth of immune vulnerability—is the clinical hallmark distinguishing keystones from other archetypes.

#### Two-dimensional robustness landscape visualizes archetype separation

Plotting niche breadth against immune breadth reveals archetype-specific regions in a two-dimensional robustness landscape (Fig. 5H). Keystone pathogens occupy the upper-right quadrant with elevated niche breadth (median=8) and moderate-to-high immune breadth (median=10), reflecting their capacity for multi-site dissemination and broad immune engagement. Opportunistic pathogens cluster in the lower-left with restricted niche breadth (median=4) and immune breadth (median=5), consistent with limited clinical manifestations outside specific immune deficiencies. Multi-host and Specialist archetypes show intermediate and overlapping patterns: multi-host pathogens have moderate niche breadth (median=5.5) and immune breadth (median=10), while specialists have low niche breadth (median=2) but variable immune breadth (median=7), reflecting their epitope-centric strategies within restricted anatomical compartments. One outlier, Nocardia (Opportunistic), exhibits exceptionally high immune breadth (25 out of 31 perturbations) yet low niche breadth (5), likely reflecting its status as a ubiquitous environmental saprophyte that opportunistically infects across diverse immune deficiency contexts without anatomical specialization.

#### Biological interpretation: clinical validation of immunological archetypes

The concordance between trait-based and clinical-derived organization—demonstrated by perfect binary classification, 67% multi-class accuracy, and ARI=0.58 in unsupervised clustering—provides strong evidence that immunological traits and clinical emergence patterns reflect overlapping biological structure. Keystone pathogens, defined by coordinated multi-arm immune engagement and structured tropism in the trait space, manifest clinically as broad anatomical disseminators with human-relevant disease across many niches (mean niche breadth = 8.0). This pattern aligns with the hypothesis that keystones undergo postnatal imprinting across multiple tissue sites, establishing coordinated CD4/CD8 memory and driving structured immune receptor selection. Specialists, characterized by epitope-centric traits, show restricted niche breadth (mean=3.4) but maintain immunological coordination within those niches. Multi-host pathogens exhibit intermediate clinical footprints (niche breadth mean=5.4) and the highest classification ambiguity (30% recall), consistent with their mixed trait profiles reflecting broad host range but variable human adaptation. Opportunists, scoring low on most traits, show correspondingly restricted clinical manifestations (niche breadth mean=3.3) yet paradoxically achieved perfect classification accuracy, reflecting their distinctive low-severity profile. The asymmetry between niche breadth (strongly archetype-discriminative, *p <* 0.001) and immune breadth (non-discriminative, *p* = 0.074) suggests that anatomical dissemination capacity, rather than the number of immune vulnerabilities, is the critical clinical dimension distinguishing keystones.

### 3.3 Mechanistic model predicts clinical emergence from biological first principles (Fig. 6)

#### Two pathways capture distinct emergence mechanisms

The mechanistic model generated predictions for 1,522 of 17,329 tensor cells (8.8%), with the remaining cells having no mechanistic path connecting the pathogen– niche–perturbation combination. Among cells with predictions, the barrier pathway dominated in 890 cases (58.5%), the latent pathway dominated in 283 cases (18.6%), and the two pathways contributed equally in 349 cases (22.9%) (Fig. 6A). This distribution indicates that both pathways contribute meaningfully, with barrier-mediated invasion being somewhat more common than reactivation-based emergence in our pathogen set.

#### Model achieves moderate correlation with clinical observations

Evaluated on the full 17,329-cell tensor using strict criteria (zeros treated as true negatives), the model achieved Pearson *r* = 0.35, Spearman *ρ* = 0.30, and Cohen’s weighted *κ* = 0.24 (fair agreement) (Fig. 6B,C). Binary classification at threshold *τ* = 40 yielded recall of 0.20, specificity of 0.97, precision of 0.42, and F1 = 0.26. The calibration curve (Fig. 6C) shows that predicted scores track observed medians, though with substantial IQR reflecting inherent clinical variability.

#### Keystone pathogens show highest predictive accuracy

Stratification by archetype revealed differential model performance (Fig. 6D). Keystone pathogens achieved the highest correlation (*r* = 0.39, *ρ* = 0.36), followed by Opportunistic (*r* = 0.36, *ρ* = 0.32), Multi-host (*r* = 0.33, *ρ* = 0.31), and Specialist (*r* = 0.33, *ρ* = 0.23). This pattern is consistent with the hypothesis that keystone pathogens, by definition, exhibit immune-dependent emergence patterns that the mechanistic model is designed to capture.

#### Model enriches true positives among top predictions

To assess ranking utility in a positive-unlabeled framework—appropriate given that zero scores may reflect absence of evidence rather than evidence of absence—we computed fold-enrichment at various prediction thresholds (Fig. 6E). At the top 5% of predictions, the model achieved approximately 2.9-fold enrichment over random baseline, meaning top-ranked predictions are nearly three times more likely to be true positives than random selection. Enrichment remained stable across thresholds (2.8– 3.0*×* from top 1% to top 20%), indicating robust ranking performance. Keystone and Multi-host pathogens showed the strongest enrichment signals, consistent with their immune-coordinated emergence patterns.

#### Pathogen–niche projection achieves strong correlation

To reduce noise from the immune perturbation dimension, we computed max-marginalized 2D projections (Fig. 6F). The pathogen *×* niche (PN) projection, which captures each pathogen’s maximum emergence score across all immune perturbations for each anatomical site, achieved substantially higher correlation (*r* = 0.58, *ρ* = 0.56) than the full 3D tensor. The pathogen *×* immune perturbation (PI) projection achieved *r* = 0.36, *ρ* = 0.37. The superior performance of the PN projection suggests the model captures pathogen–tissue tropism relationships particularly well—a biologically meaningful finding, as pathogens have characteristic tissue preferences that the model successfully encodes through lytic attack capacity and controller residency patterns.

#### Biological interpretation: mechanistic validation of archetype-specific emergence

The mechanistic model’s differential performance across archetypes provides biological validation of the trait-based classification. Keystone pathogens, defined by coordinated multi-arm immune engagement, show the highest correlation because their emergence patterns are precisely what the model was designed to capture: release from immune control leading to predictable clinical manifestations. The finding that both latent (18.6%) and barrier (58.5%) pathways contribute meaningfully indicates the model captures genuine biological heterogeneity in emergence mechanisms. The 2.9-fold enrichment at top predictions demonstrates practical utility: the mechanistic model can prioritize pathogen–niche–perturbation combinations for clinical surveillance or therapeutic intervention. Importantly, cases where the model predicts high emergence but clinical data shows zero (“predicted high, observed zero”) may represent not model failures but rather unstudied combinations, incomplete clinical characterization, or absence of pathogen exposure in studied populations—possibilities strongly supported by our finding that all zero scores have low confidence (no curated evidence of absence).

## 4 Discussion

### Keystone organisms as an operational immunological phenotype

This work moves immunological “keystone organisms” from an ecological metaphor to an operational phenotype by triangulating (i) trait biology, (ii) clinical emergence under immune perturbation, and (iii) a first-principles mechanistic perturbation model. Across 43 pathogens, trait-space clustering yielded four reproducible archetypes and a compact set of human-adapted keystones dominated by persistent herpesviruses and *M. tuberculosis* (Fig. 4). These organisms converge on long-term residence, tissue-structured tropism, and the need for coordinated control spanning antigen presentation, innate sensing, and T/NK effector programs.

### Clinical emergence distinguishes keystones by dissemination, not simply by “more immunosuppression triggers.”

By encoding published associations as a pathogen *×* immune-perturbation *×* niche tensor, we show that the clinically salient signature of keystones is their ability to occupy a broader repertoire of anatomical sites when immune coordination is perturbed (Fig. 5). In our dataset, diagnostic breadth decomposed into immune breadth and niche breadth, and it is niche breadth that separates keystones from other archetypes. This pattern argues for compartmentalized, tissue-resident checkpoints (e.g., mucosal barrier integrity, myeloid restriction, TRM/NK patrol) as key governors of keystone containment. Practically, it helps explain why patients with comparable “global” immunosuppression can present with distinct pathogen portfolios: the decisive variable is which immune functions fail in which tissue compartment, not only the magnitude of systemic lymphopenia.

### Niche calibration and thresholded peripheral tolerance

TRAIT identifies keystone organisms as an apex class: multi-niche persistent residents whose containment requires coordinated, multi-arm immune control and whose clinical emergence is distinguished by broad niche breadth. However, immune calibration is enacted locally within anatomical compartments. Several persistent, niche-restricted organisms can plausibly calibrate compartment-specific thresholds of cytotoxicity and regulation without meeting full keystone criteria. In this hierarchy, keystones are a subset of niche-calibrating organisms, and some tissues may be dominated by non-keystone calibrators (for example, BKV in kidney, JCV in CNS, and *H. pylori* in gastric mucosa). Figure 7 summarizes this nested relationship and highlights exposure-dependent calibrators that may be underrepresented in Western cohorts. (We use “niche-calibrating organisms” here at the organism level; the corresponding epitope-level unit is treated in Keystone Epitope Theory as “niche-calibrating epitopes.”)

**Fig. 7.**
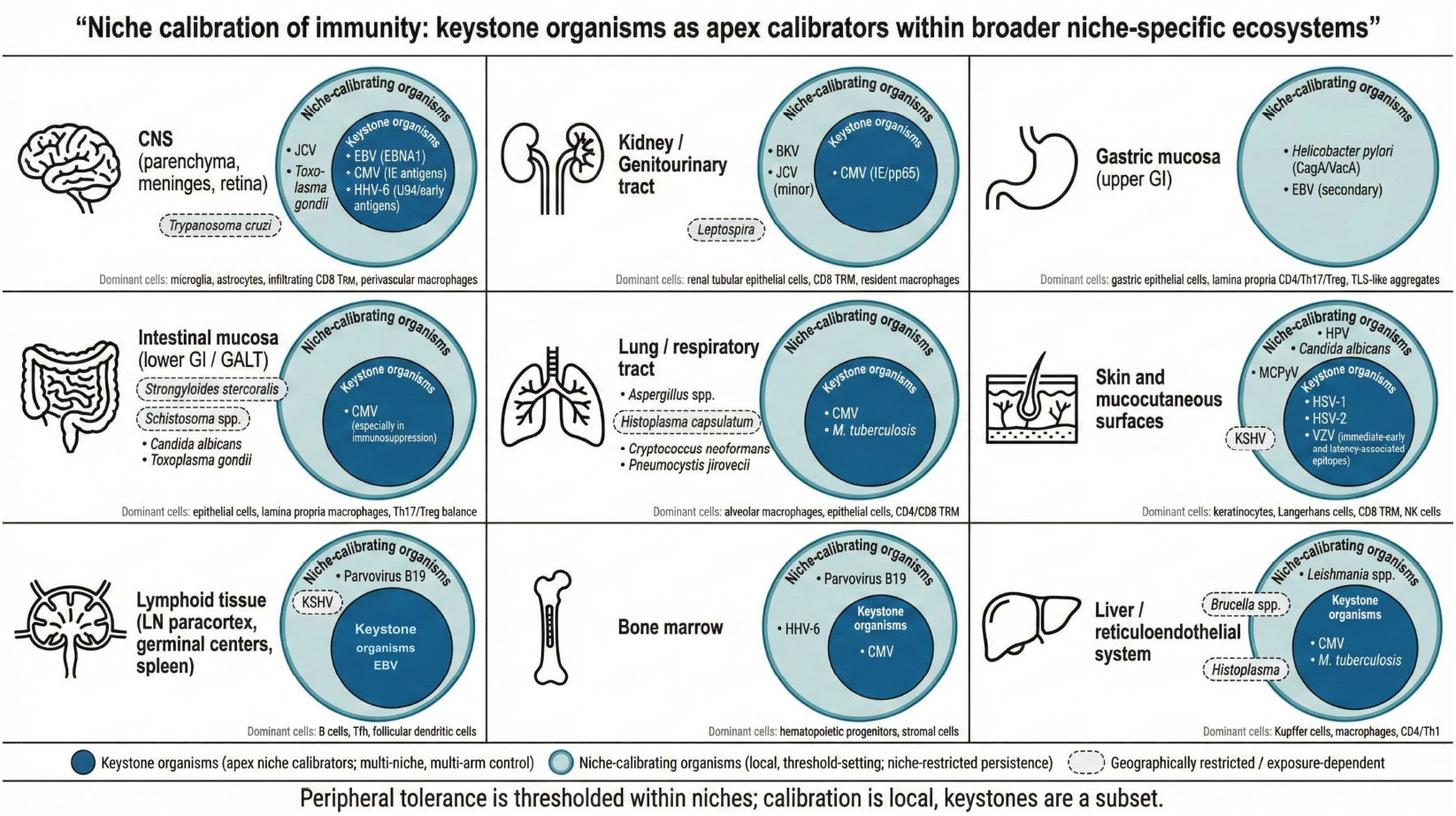
Niche calibration of immunity: keystone organisms as apex calibrators within broader niche-specific ecosystems. Anatomical compartments can be treated as local immune ecosystems shaped by persistent residents that repeatedly engage tissue-resident memory and regulatory set-points. Outer circles indicate niche-calibrating organisms that can drive compartment-specific calibration through durable persistence and repeated antigen display. Inner circles indicate keystone organisms, the apex subset distinguished in TRAIT by multi-niche persistence and multi-arm immune coordination. Several compartments exhibit strong calibration dominated by persistent non-keystone residents, emphasizing that thresholded peripheral tolerance is enacted locally within niches and that “keystone” is a subset relationship rather than a binary. Dashed outlines indicate exposure-dependent or geographically restricted organisms.

### A first-principles model recapitulates emergence and prioritizes risk

The mechanistic model integrates three recurring levers of opportunism—barrier disruption, activation of latent/persistent reservoirs, and loss of local immune control—and uses organism-, niche-, and perturbation-level features derived from the tensor. Despite its simplicity, it reproduced observed emergence patterns (overall *r* = 0.35) and performed best when aggregated across niches (pathogen *×* niche projection *r* = 0.58), suggesting that the dominant learnable structure is anatomical rather than purely immunological. Importantly, the model enriched the highest-risk pathogen–immune–niche triplets 2.9-fold, supporting its role as a hypothesis generator for surveillance, prophylaxis, and mechanistically targeted diagnostic workups (Fig. 6).

### From taxonomy to translation: why keystones matter clinically

Persistent viromes and bacteriomes are increasingly recognized as contributors to immune calibration and host fitness, not merely as threats [2–4]. Our results extend this view by identifying a subset of long-term residents that behave like immunological organizers: they are constrained by multi-arm, multi-compartment control, and when that control is relaxed they become sentinels that reveal where immune coordination has failed. This framing can help contextualize heterogeneous infectious complications across transplantation, biologic immunomodulation, autoimmunity, and inborn errors of immunity [11, 12].

#### Box 1. Design and Clinical Use Guidelines

1. **Treat keystone emergence as a systems readout**. Because keystone organisms are long-term residents with tissue-structured reservoirs, their *reactivation* (and especially their appearance in non-canonical compartments) is often less informative as an isolated microbiologic event than as a coarse *systems-level* readout of immune coordination failure. In practice, the *identity* of the keystone (which control programs it typically demands) and the *niche* in which it emerges (which barrier and tissue-resident checkpoints have failed) can be interpreted as a structured signal of which immune arms and compartmental defenses are compromised [4]. This is most useful when applied longitudinally (baseline → perturbation → recovery), and when detections are counted as *episodes* (unique organism–compartment–time events), rather than repeated tests of the same episode. A practical instantiation is a *Keystone Reactivation Index* (*Keystone Reactivation Index*), defined within a compartment *c* over a time window *t* as the fraction of keystone detections among all infectious detections:

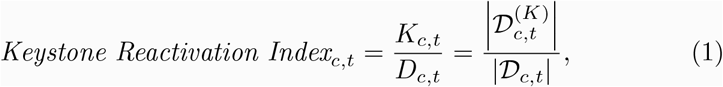

where 𝒟_*c,t*_ is the set of infectious detection episodes observed in compartment *c* during window *t*, 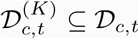 is the subset classified as keystone organisms, *D*_*c,t*_ = |𝒟_*c,t*_|, and 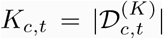. (When *D*_*c,t*_ = 0, *Keystone Reactivation Index* is undefined and can be treated as missing rather than zero.) Optionally, a patient-level summary over compartments can be computed as

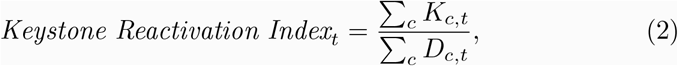

which preserves interpretability while allowing longitudinal tracking of keystone-weighted infectious burden.
2. **Prioritize containment in the compartment that declares itself**. Because keystone signatures are tissue-structured, prophylaxis and monitoring should be tailored to the affected niche (e.g., CNS, gut, lung), not only to systemic immune cell counts.
3. **Avoid immunogens dominated by mutable decoys**. Vaccine or immunotherapy designs that repeatedly present highly variable epitopes may preferentially recall familiar but non-protective memory, diverting responses away from conserved constrained targets [22–24].
4. **Screen for keystone-patterned cross-reactivity in high-risk settings**. In individuals prone to severe T-cell–mediated hypersensitivity, autoimmunity, or transplantation rejection, evaluating whether candidate drugs, vaccines, tumors, or alloantigens generate peptides resembling tissue-local keystone epitopes—or (in compartments without a dominant keystone) dominant niche-calibrating epitopes that may set local thresholded tolerance— may identify risk for pathogenic memory recall [18–21, 26, 27].

### Keystone-focused memory provides a mechanistic bridge to immunopathology

At the micro level, long-lived, tissue-homing memory to persistent keystones is advantageous for rapid control, but it also creates a privileged path for “modified-self” ligands to recruit potent effector programs in the same compartment. This offers an immunological explanation for why some drug hypersensitivity and organ-specific autoimmune syndromes are strongly HLA- and tissue-linked: they may reflect misdirected recall of keystone-trained T-cell repertoires [18–21, 26, 27]. This prediction is testable: culprit HLA–peptide complexes should preferentially engage memory clonotypes with prior keystone specificity and appropriate tissue imprinting. This framing invites a complementary question: can intentional re-anchoring of keystone-specific immunity reduce downstream misallocation, including drift toward altered-self or post-translationally modified (PTM) ligands, particularly in late life?

Therapeutic perturbations of canonical keystone residents provide an underused class of “stress tests” for keystone-anchored allocation models. Quasi-experimental analyses exploiting eligibility cutoffs report a lower incidence of dementia diagnoses after herpes zoster vaccination in Wales and Australia [28, 29], and a natural experiment during the transition from live to recombinant zoster vaccination found a lower subsequent dementia risk with the recombinant vaccine [30]; related analyses further suggest effects extending to mild cognitive impairment and dementia-related mortality [31]. Population-scale cohort data also link zoster vaccination to lower cardiovascular event rates [32]. If these signals reflect more than residual confounding, they are consistent with the possibility that vaccine-amplified keystone-epitope focusing improves equilibrium efficiency within relevant niches, reducing background inflammatory variance that otherwise increases the probability that altered-self or PTM ligands cross local activation thresholds and recruit misdirected recall. Notably, a randomized clinical trial of high-dose valacyclovir in early symptomatic Alzheimer disease with HSV seropositivity showed no benefit and worse performance on the primary cognitive endpoint [33], suggesting that pharmacologic herpesvirus suppression may not recapitulate vaccine-driven immune recalibration and could, in some contexts, shift control dynamics in an unfavorable direction.

If so, keystone-directed vaccination becomes not only prevention, but a mechanism-informative, time-stamped perturbation that can be sampled to connect epitope focusing, niche control, and distal clinical outcomes.

### Implications for fast-evolving pathogens and immunogen design

Immune hierarchies shaped by persistent infections can be exploited by rapidly adapting RNA viruses, which may inflate immunodominant yet mutable epitopes that trigger recall without durable control [22–25]. The archetype framework sharpens this claim: decoys should be most effective when they mimic the “keystone pattern” of presentation and tissue context that the host has learned to prioritize. Immunogen design that forces recognition of conserved, functionally constrained sites—and avoids over-amplifying keystone-like decoys—may therefore be essential for durable protection.

### Limitations and next steps

Trait scoring compresses heterogeneous biology into rubric values, and the clinical tensor is literature-derived, subject to reporting bias, evolving diagnostics, and non-independence of case series. Our classifiers and mechanistic model were validated internally; external prospective validation in well-annotated cohorts is required before clinical deployment. Nonetheless, the convergence of trait-based archetypes, clinical emergence structure, and first-principles modeling suggests that keystone organisms are not an artifact of any single data source. Next steps include expanding the Keystone Protein Atlas in a compartment-aware manner, integrating epitope-level constraints and host genetics, and performing prospective surveillance under defined immunomodulatory regimens to refine the mapping from perturbation to niche-specific risk [6, 7, 14, 15, 34, 35].

## 5 Conclusion

We define immunological keystone organisms as a reproducible class of human-adapted pathogens whose containment depends on coordinated, multi-compartment immunity and whose failures are clinically legible. By combining trait-based archetyping, a literature-derived clinical emergence tensor, and a simple perturbation model, we show that keystone status is reflected in distinctive patterns of tissue dissemination under immune deficits, and that these patterns can be predicted from biological first principles. This framework enables archetype-aware monitoring in immunosuppressed patients, generates testable mechanistic hypotheses about tissue-specific vulnerability and immune-mediated pathology, and provides a rational basis for immunogen design that prioritizes conserved, functionally constrained targets.

This framing turns what is often treated as “opportunistic noise” into a structured readout of immune coordination: which organism emerges, where it emerges, and under which perturbation jointly report which tissue checkpoints and immune arms are failing. In practice, this enables archetype-aware monitoring in immunosuppressed patients, motivates the development of longitudinal compartment-aware metrics (for example, a Keystone Reactivation Index), and generates testable hypotheses about why anatomically patterned immunopathology manifests in one tissue rather than another.

More broadly, this work provides the quantitative substrate for Keystone Epitope Theory by identifying which persistent organisms are most likely to write the postnatal antigenic curriculum that calibrates tissue thresholds and immunodominance hierarchies. The same keystone-shaped priorities that can be exploited by fast-evolving RNA viruses and tumor ecosystems through decoy inflation and “keystone-pattern” mimicry are enforced through interoperable cytotoxic T cell and natural killer cell reading of shared peptide–HLA surfaces [37], and they help explain the niche-gated conditions under which modified-self events recruit entrenched memory to drive hypersensitivity, autoimmunity, and alloreactivity [38]. Modern therapeutics make these mechanisms experimentally accessible in humans: checkpoint blockade can be treated as a perturbation assay that reveals which tissues are being held in equilibrium by thresholding, and why immune-related adverse events are organ-selective and mechanistically structured rather than idiosyncratic [39]. Likewise, EBV-associated multiple sclerosis illustrates how a keystone-trained program can be redeployed within a CNS-relevant niche under a finite, receptor-first discovery workflow [40]. Finally, the deep-time record suggests that persistent infection, endogenized viral sequences, and episodic perturbation co-write a genomic syllabus that constrains immune allocation across timescales, generating falsifiable predictions that are now testable with immunopeptidomics, repertoire analysis, and longitudinal virome measurements [41, 42]. Together, TRAIT reframes immune memory as a constrained allocation problem and provides a clinically usable map of what the immune system has been forced to remember, when that memory protects, and when it misfires.

## Code Availability

All code and data used to generate the results in this study is publicly available at https://github.com/AsiaeeLab/keystone-organisms.

## Acknowledgements

This work was supported in part by the National Institutes of Health (NIH) under awards AI154659, P50GM115305, 5P30AI110527, 1R35HL140016, 5R01AI060460, U54AG089326, P01HL174442, P30CA068485, R01AI077505, 1R01AI147765, 1R01AI142093, and R00HG011367.

## A Summary Tables of Pathogen Characteristics

Tables 3–6 provide detailed immunological and evolutionary characteristics for each pathogen, organized by archetype assignment. For each organism, we document infection pattern, geographic distribution, key immune dependencies, capacity for lifelong infection, primary tissues and cell types involved, clinical impact under immunosuppression, and estimated co-evolutionary time with humans.

## B Trait-Based Scoring: Technical Details

### B.1 Score vector representation

Each pathogen *i* was represented by an 18-dimensional score vector **x**_*i*_ = (*x*_*i*,1_, …, *x*_*i*,18_), where *x*_*i,t*_ denotes the score for pathogen *i* on trait *t*. Total keystoneness scores were computed as unweighted sums:

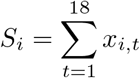

### B.2 Gower distance

Pairwise dissimilarities were computed using Gower distance, which accommodates mixed ordinal and binary data while handling missing values:

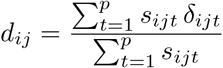

where *s*_*ijt*_ ∈ {0, 1} indicates whether trait *t* is observed for both pathogens *i* and *j* (non-missingness indicator), and *δ*_*ijt*_ is the trait-specific dissimilarity:

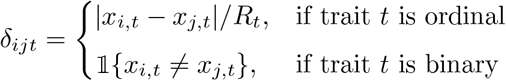

Here *R*_*t*_ is the observed range for ordinal trait *t*, and 𝟙{·} is the indicator function.

### B.3 Clustering algorithms

Two clustering approaches were applied to the Gower distance matrix:

#### Partitioning around medoids (PAM)

PAM minimizes the sum of dissimilarities between each object and its assigned medoid:

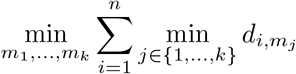

where *m*_1_, …, *m*_*k*_ are the indices of the *k* medoid organisms.

#### Hierarchical clustering

Agglomerative clustering with complete linkage, where the distance between clusters is defined as the maximum pairwise distance:

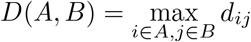

### B.4 Cluster number selection

The number of clusters was selected using elbow and angle heuristics applied to within-cluster sum of dissimilarities *W* (*k*) for *k* = 2, …, 8:

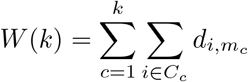

Both methods nominated *k* = 4 clusters.

### B.5 Consensus clustering

Method agreement was quantified by the Adjusted Rand Index (ARI):

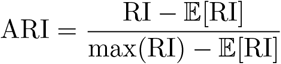

where RI is the Rand Index measuring pairwise agreement between clusterings.

For organisms with discordant assignments between PAM and hierarchical clustering, we applied a conservative consensus rule: each organism was assigned to the candidate cluster with the lower mean total score 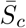. Four organisms were reassigned under this rule, including Parvovirus B19 (Keystone → Specialist) and three additional organisms (Specialist → Opportunist). To further enhance the specificity of the keystone label, organisms initially assigned to the keystone archetype underwent an additional centroid-based validation step and were reassigned if their total trait score was closer to the centroid of another cluster.

### B.6 Trait signature analysis

Cluster-versus-rest comparisons used Cliff’s delta (*δ*) for effect size, defined as:

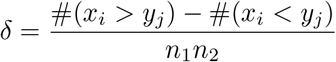

where *x*_*i*_ are scores in the focal cluster and *y*_*j*_ are scores in remaining clusters. Wilcoxon rank-sum tests were used for significance testing. Statistical significance was assessed using uncorrected *p*-values at *α* = 0.05 (*** *p <* 0.001, ** *p <* 0.01, * *p <* 0.05).

Multivariate signatures were obtained from *ℓ*_1_-penalized (LASSO) multinomial logistic regression:

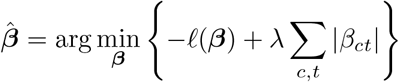

where *ℓ*(***β***) is the multinomial log-likelihood, the *ℓ*_1_ penalty encourages sparsity in trait selection, and *λ* was selected by 10-fold cross-validation. The regularization parameter *α* = 1 specifies pure LASSO penalization.

## C Clinical Emergence Tensor: Scoring and Analysis

We constructed a three-dimensional clinical emergence tensor **T** ∈ ℝ^43*×*13*×*31^ documenting pathogen reactivation or invasion patterns across:

- 43 pathogens (matching those in the immunological trait analysis)
- 13 anatomical niches (blood/systemic, CNS, respiratory mucosa, GI mucosa, genitourinary mucosa, skin, reticuloendothelial system, bone marrow, lymph nodes, cardiac/endovascular, bone/joint, ocular, lymph node germinal centers)
- 31 immune perturbations spanning genetic immunodeficiencies and pharmacologic immunosuppression

### C.1 Scoring Rubric

#### C.1.1 Tensor construction and scoring

Each tensor element *T* (*p, n, i*) ∈ [0, 100] represents the strength of clinical evidence for lytic emergence, scored according to:

- **85–100**: Replicated human studies (genetic or interventional), hallmark associations
- **70–84**: Solid human signal for this niche with multiple sources or one high-quality source
- **40–69**: Clear but context-dependent human evidence, dose/timing dependent, partial compensation by other immune arms
- **10–39**: Suggestive or narrow human signals, or strong mechanistic logic with limited human data
- **5**: No authoritative human evidence found after diligent search
- **0**: Published evidence indicates protective effect or no increased emergence

The threshold at 40 demarcates human-relevant clinical data (scores ≥ 40 include at least limited human observations with mechanistic support, grade C or better) from preclinical evidence only (scores *<* 40, grade D).

#### C.1.2 Evidence grading and confidence annotation

Each score was accompanied by structured metadata:

- **Direction**: “revealed” (perturbation increases emergence), “protected” (reduces risk), or “unclear” (conflicting/insufficient data)
- **EvidenceGrade**:
  – A = replicated human genetic or interventional evidence
  – B = strong human clinical cohorts or case series
  – C = limited human data with strong mechanistic support
  – D = animal or in vitro evidence only
- **ConfidenceTag**: [High], [Moderate], [Low], [Minimal] (for score 5), [None] (for score 0)
- **Justification**: rigorous citations formatted as (Author, Title, Journal, Year)
- **EvidenceCount**: number of distinct human studies supporting the score
- **Missingness**: “observed” (evidence exists), “inferred zero” (no evidence after diligent search), or “unknown”

Evidence was prioritized as: (1) human genetic/interventional perturbations with niche-localized outcomes, (2) human clinical cohorts with clear site localization, (3) human observations without precise site (stated with caveats), (4) animal/in vitro support only (capped at score 69, grade C or D).

#### C.1.3 Controlled vocabularies for niches and perturbations

Each anatomical niche was defined by controlled vocabularies specifying required anatomical terms (IncludeTokens) and exclusion criteria (ExcludeTokens) to ensure precise localization. For example, CNS required terms like “brain,” “cerebral,” “encephalitis,” “CSF,” or “meninges,” while excluding “ocular” or “peripheral nerve.” Similarly, each immune perturbation was defined by mechanism and synonyms. For instance, P07 (CalcineurInh) encompasses “tacrolimus,” “cyclosporine,” and “calcineurin inhibitor,” all blocking NFAT and IL-2 signaling. Complete niche and perturbation definitions are in Supplementary Section C.2.

#### C.1.4 Sensitivity analyses: Score adjustment schemes

To assess robustness to evidence quality, we performed sensitivity analyses by adjusting scores based on evidence grade and confidence:

##### Grade-capping

Scores were reduced for lower evidence grades to prevent overweighting of animal/in vitro data:

- Grade A: score unchanged
- Grade B: score capped at 84 (if originally *>* 84)
- Grade C: score capped at 69 (if originally *>* 69)
- Grade D: score capped at 39 (if originally *>* 39)

##### Grade+Confidence weighting

Scores were multiplicatively adjusted by a factor combining evidence grade and confidence tag, down-weighting low-confidence or low-grade entries. Weighting factors were:

- High: 1.0 (no reduction)
- Moderate: 0.85
- Low: 0.70
- Minimal/None: 0.50

Combined with grade-specific caps, this yielded conservative score estimates prioritizing high-quality human evidence.

All feature extraction (max-marginalization, breadth metrics), classification (elastic-net regression), and clustering (PAM) analyses were re-run on both adjusted score tensors to test whether archetype predictions and unsupervised recovery remain robust when evidence quality is explicitly down-weighted.

### C.2 Tensor Dimensions

The clinical tensor **T** ∈ ℝ^43*×*13*×*31^ captures pathogen behavior across anatomical niches and immune perturbations. Below we provide detailed definitions for each dimension.

#### C.2.1 Anatomical Niches (N=13)

1. **CNS (Central Nervous System)**: Brain parenchyma, meninges, cerebrospinal fluid spaces. Reflects capacity for neuroinvasion, meningitis, or encephalitis. Relevant for neurovirulent pathogens and those causing CNS opportunistic infections.
2. **Lung (Pulmonary)**: Pulmonary parenchyma, airways, alveolar spaces. Encompasses pneumonia, bronchitis, and respiratory tract colonization. Central niche for respiratory viruses, bacteria, and fungi.
3. **GI (Gastrointestinal)**: Gastrointestinal mucosa, lumen, and associated lymphoid tissue (GALT). Includes enteric infections, gastroenteritis, and mucosal colonization. Relevant for fecal-oral transmission routes.
4. **Liver (Hepatic)**: Hepatic parenchyma, including hepatocytes, Kupffer cells, and bile ducts. Reflects hepatotropism, hepatitis, and intrahepatic replication. Important for bloodborne and vector-borne pathogens.
5. **Blood (Bloodstream/Intravascular)**: Intravascular space, including circulating immune cells and endothelium. Captures bacteremia, viremia, parasitemia, and disseminated infections. Critical for sepsis-causing organisms.
6. **LN Paracortex (Lymph Node T Cell Zones)**: T cell-rich paracortical regions of lymph nodes. Site of adaptive T cell priming and cell-mediated immune responses. Reflects pathogens that manipulate or reside within T cell niches.
7. **LN GC (Lymph Node Germinal Centers)**: B cell germinal centers within secondary lymphoid tissues. Sites of B cell maturation, somatic hypermutation, and antibody class-switching. Reflects pathogens that colonize or replicate within organized B cell zones.
8. **Spleen (Splenic)**: Splenic parenchyma, including red pulp (filtration) and white pulp (lymphoid). Central to bloodborne pathogen clearance and immune surveillance. Critical for encapsulated bacteria and blood-stage parasites.
9. **BoneMarrow (Hematopoietic Marrow)**: Bone marrow niches supporting hematopoiesis. Reflects pathogens causing marrow suppression, hemophagocytic syndromes, or intracellular marrow persistence.
10. **Skin (Cutaneous/Subcutaneous)**: Skin, subcutaneous tissue, and dermal immune cells. Includes cellulitis, abscesses, vector bite sites, and dermatological manifestations. Entry point for many vector-borne and traumatic infections.
11. **GU (Genitourinary)**: Genitourinary mucosa, renal parenchyma, bladder. Encompasses urinary tract infections, pyelonephritis, and sexually transmitted infections.
12. **Bone (Osseous)**: Bone tissue and periosteum. Reflects osteomyelitis, septic arthritis, and bone-invasive infections. Relevant for hematogenous seeding and trauma-associated infections.
13. **Eye (Ocular)**: Ocular tissues including conjunctiva, cornea, uvea, and retina. Captures keratitis, uveitis, endophthalmitis, and retinitis. Important for certain congenital and disseminated infections.

#### C.2.2 Immune Perturbations (N=31)

1. **P01 (Neutropenia)**: Severe neutrophil depletion (*<*500 cells/*µ*L). Impairs bacterial and fungal killing, increasing risk for invasive bacterial and mold infections. Seen in chemotherapy, aplastic anemia, and congenital neutropenias.
2. **P02 (MacrophageDefect)**: Macrophage dysfunction or depletion. Compromises intracellular pathogen control, granuloma formation, and tissue clearance. Relevant for mycobacteria, Listeria, Salmonella, and intracellular parasites.
3. **P03 (CD4depletion)**: CD4+ T cell deficiency (e.g., HIV/AIDS, idiopathic CD4 lymphopenia). Central immunodeficiency affecting cell-mediated and humoral immunity. Predisposes to opportunistic infections and impaired vaccine responses.
4. **P04 (CD8depletion)**: CD8+ T cell deficiency or dysfunction. Impairs cytotoxic responses against intracellular pathogens and virally infected cells. Relevant for chronic viral infections and some intracellular bacteria.
5. **P05 (NKdepletion)**: Natural killer cell deficiency or dysfunction. Impairs early antiviral and antitumor responses. Associated with increased susceptibility to herpesviruses and some bacterial infections.
6. **P06 (TNFblock)**: TNF-*α* blockade via monoclonal antibodies (infliximab, adalimumab, etanercept). Impairs granuloma formation and macrophage activation. Major risk factor for tuberculosis reactivation and invasive fungal infections.
7. **P07 (IFNgBlock)**: IFN-*γ* neutralization or receptor deficiency. Critical for macrophage activation and intracellular pathogen control. Deficiency predisposes to disseminated mycobacterial and Salmonella infections.
8. **P08 (IL12p40block)**: IL-12/IL-23 p40 subunit blockade (ustekinumab). Impairs Th1 and Th17 responses. Increases risk for mycobacterial, fungal, and some bacterial infections.
9. **P09 (IL17block)**: IL-17A blockade (secukinumab, ixekizumab) or receptor deficiency. Impairs mucosal immunity and neutrophil recruitment. Predisposes to mucocutaneous candidiasis and some bacterial infections.
10. **P10 (IL1block)**: IL-1 blockade (anakinra, canakinumab) or receptor deficiency. Reduces pyogenic responses and fever. Modest infection risk, primarily pyogenic bacteria.
11. **P11 (C3deficiency)**: Complement C3 deficiency or depletion. Central complement component; deficiency severely impairs opsonization and immune complex clearance. High risk for encapsulated bacteria and Neisseria.
12. **P12 (ProperdinDeficiency)**: Properdin deficiency, stabilizer of alternative pathway C3 convertase. X-linked deficiency with markedly increased susceptibility to meningococcal disease.
13. **P13 (MBLdeficiency)**: Mannose-binding lectin deficiency. Impairs lectin pathway activation and opsonization. Modest increased risk for bacterial infections, particularly in young children.
14. **P14 (Hypogammaglobulinemia)**: Severe antibody deficiency (CVID, X-linked agammaglobulinemia). Impairs opsonization, neutralization, and immune complex clearance. High risk for encapsulated bacteria, enteroviruses, and chronic sinopulmonary infections.
15. **P15 (Bcelldepletion)**: B cell depletion via anti-CD20 therapy (rituximab, ocrelizumab) or congenital deficiency. Impairs antibody production and antigen presentation. Increases risk for bacterial infections and viral reactivation.
16. **P16 (TLRdefect)**: Toll-like receptor pathway defects (TLR signaling impairment). Reduces innate recognition of PAMPs. Variable susceptibility depending on specific TLR; broadly impairs innate immune priming.
17. **P17 (MyD88deficiency)**: MyD88 adaptor protein deficiency. Impairs most TLR and IL-1R signaling. Severe susceptibility to pyogenic bacteria, particularly Streptococcus pneumoniae and Staphylococcus aureus.
18. **P18 (IRAK4deficiency)**: IRAK4 kinase deficiency, downstream of MyD88. Similar phenotype to MyD88 deficiency with invasive bacterial infections in childhood.
19. **P19 (NFkBdefect)**: NF-*κ*B pathway defects (NEMO, I*κ*B*α* mutations). Impairs broad inflammatory and immune gene transcription. Variable phenotype with mycobacterial, bacterial, and fungal susceptibility.
20. **P20 (CGDdefect)**: Chronic granulomatous disease (phagocyte NADPH oxidase deficiency). Eliminates oxidative burst, impairing killing of catalase-positive organisms. High risk for Staphylococcus, Aspergillus, Burkholderia, Serratia, and Nocardia.
21. **P21 (MHC1deficiency)**: MHC class I deficiency (TAP deficiency, bare lymphocyte syndrome type I). Impairs CD8+ T cell antigen presentation. Increases susceptibility to viral infections.
22. **P22 (MHC2deficiency)**: MHC class II deficiency (bare lymphocyte syndrome type II). Impairs CD4+ T cell priming. Severe combined-like immunodeficiency with broad bacterial, viral, and fungal susceptibility.
23. **P23 (CD40Ldeficiency)**: CD40 ligand deficiency (X-linked hyper-IgM syndrome). Impairs T cell help for B cells and macrophage activation. Predisposes to Pneumocystis, Cryptosporidium, and bacterial infections.
24. **P24 (CTLA4agonist)**: CTLA-4 agonism or Ig fusion (abatacept). Inhibits T cell costimulation. Modest infection risk, primarily opportunistic and mycobacterial.
25. **P25 (CheckpointBlockade)**: Checkpoint blockade (PD-1/PD-L1; e.g., nivolumab, pembrolizumab). Although not a classical “immune deficit,” it can increase infection risk in specific settings by dysregulating tissue thresholds and inducing immune-mediated pathology, even while enhancing some pathogen control programs.
26. **P26 (IL6block)**: IL-6 or IL-6R blockade (tocilizumab, sarilumab). Reduces acute phase responses and Th17 differentiation. Increases risk for bacterial infections, diverticulitis, and some opportunistic pathogens.
27. **P27 (IL10block)**: IL-10 blockade or deficiency. IL-10 is anti-inflammatory; blockade may paradoxically enhance immunity but also cause inflammatory bowel disease. Included for completeness; rare clinical use.
28. **P28 (IL23block)**: IL-23 blockade (p19 subunit: guselkumab, risankizumab). More specific than IL-12/23 blockade; impairs Th17 responses. Infection risk appears lower than IL-12/23 blockade but includes fungal and mycobacterial concerns.
29. **P29 (JAKblock)**: JAK kinase inhibition (tofacitinib, baricitinib, upadacitinib). Broad impairment of cytokine signaling (IFN, IL-6, IL-12, IL-23, etc.). Increases risk for herpes zoster, tuberculosis, and opportunistic infections.
30. **P30 (CSFblock)**: GM-CSF or G-CSF blockade/deficiency. Impairs myeloid development, neutrophil function, and alveolar macrophage maturation. GM-CSF neutralization associated with pulmonary alveolar proteinosis and opportunistic lung infections.
31. **P31 (IL5block)**: IL-5 blockade via monoclonal antibodies (mepolizumab, reslizumab, benralizumab). IL-5 drives eosinophil development, survival, and activation. Blockade severely depletes eosinophils, impairing helminth immunity and potentially tissue eosinophil-mediated responses. Used clinically for severe eosinophilic asthma and hypereosinophilic syndrome.

### C.3 Feature extraction from tensor and analysis

From the 3D tensor, we derived 2D feature matrices for each pathogen via maxmarginalization:

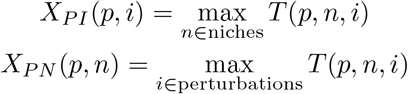

These matrices capture the maximal emergence score across all contexts, yielding a 43 *×* 44 feature matrix per pathogen (31 immune perturbations + 13 niches). We augmented this with breadth metrics:

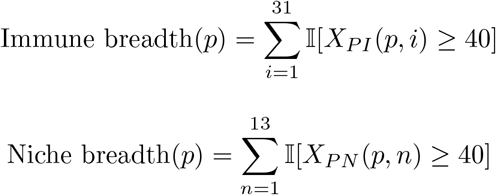

quantifying the number of immune perturbations and anatomical niches exhibiting clinically significant emergence (threshold = 40, reflecting human-relevant data).

#### C.3.1 Supervised classification

We trained elastic-net logistic regression models to predict archetype membership from clinical features:

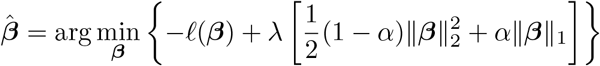

where *ℓ*(***β***) is the log-likelihood (binomial for binary, multinomial for 4-way classification), *α* = 0.5 balances *ℓ*_1_ and *ℓ*_2_ penalties, and *λ* was selected by cross-validation. We used 8-fold cross-validation for binary classification (keystone vs. non-keystone) to ensure adequate representation in each fold, and 5-fold for multi-class prediction to accommodate smaller archetype sample sizes. Models were fit using the glmnet package with AUC (binary) or misclassification error (multi-class) as the optimization criterion.

#### C.3.2 Unsupervised clustering validation

To test whether clinical emergence patterns independently recover trait-based archetypes, we performed PAM clustering on Gower dissimilarities of the feature matrix with *k* = 4 clusters. Cluster assignments were aligned to archetype labels via permutation optimization to maximize diagonal agreement, and concordance was quantified by Adjusted Rand Index (ARI). This unsupervised approach provides external validation that archetypes derived from immunological traits (Figure 4) correspond to distinct clinical emergence signatures.

All analyses (feature extraction, classification, clustering) were repeated for raw scores, grade-capped scores, and grade+confidence weighted scores to assess robustness across evidence quality thresholds.

## D Mechanistic Model: Technical Details

### D.1 Mechanistic scoring tables

Five expert-curated scoring tables quantified biological interactions (all scores normalized to [0, 1]):

1. **Latent control** *L*(*p, n*_lat_, *c*) ∈ [0, 1]: Strength with which immune controller *c* suppresses pathogen *p* in latent niche *n*_lat_. Higher scores indicate tighter immunological containment. (154 scored interactions)
2. **Lytic attack** *A*(*p, n*) ∈ [0, 1]: Capacity of pathogen *p* to breach barriers and cause clinical disease in niche *n*, independent of immune status. (145 scored interactions)
3. **Reactivation routes** *R*(*p, n*_lat_ → *n*_lytic_) ∈ [0, 1]: Probability that pathogen *p* escapes from latent reservoir *n*_lat_ to manifest clinically in lytic niche *n*_lytic_. (151 scored interactions)
4. **Perturbation effects** *E*(*i, c*) ∈ [0, 1]: Degree to which immune perturbation *I* disables controller *c*. (144 scored interactions)
5. **Controller residency** *C*(*c, n*) ∈ {0, 1} : Binary indicator for whether controller *c* operates in niche *n*. (132 scored interactions)

#### D.1.1 Score normalization

Raw expert scores *s* ∈ [0, 100] were normalized using:

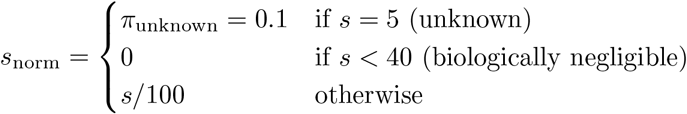

The threshold of 40 reflects expert consensus that lower scores indicate biologically negligible interactions. Crucially, unknown interactions (originally coded as 5) retain a small prior probability of 0.1 rather than being zeroed, reflecting that an unstudied interaction is not necessarily absent.

#### D.1.2 Perturbation effect scaling

Perturbation effects were scaled by qualitative effect type using multiplier *κ*:

- Depleted: *κ* = 1.0 (complete loss of controller)
- Disabled: *κ* = 0.8 (functional impairment with residual presence)
- Functionally impaired: *κ* = 0.6 (partial dysfunction)
- Unknown: *κ* = 0.6 (conservative default)

The final perturbation effect is *E*_scaled_ = *E*_base_ *× κ*.

### D.2 Latent pathway

The latent pathway models emergence through reactivation from immune-controlled reservoirs:

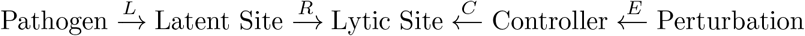

For each mechanistic chain (*p, n*_lat_, *n*_lytic_, *c, i*), the base score uses geometric mean (length-normalized):

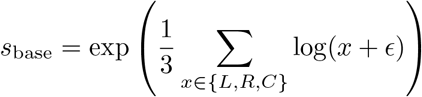

where *ϵ* = 10^−6^ prevents log(0).

The perturbation effect acts as a modulatory exponent with strength *β* = 1.5:

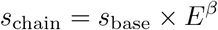

This means complete controller disabling (*E* = 1) allows full expression of the base score, while partial effects attenuate emergence exponentially.

### D.3 Barrier pathway

The barrier pathway models direct invasion when immune barriers are compromised:

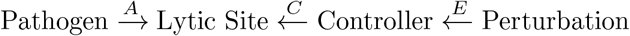

To prevent over-triggering for controllers that don’t actually control the pathogen, we added pathogen–controller specificity using latent control scores as a proxy:

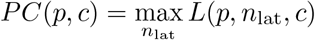

The barrier chain score is then:

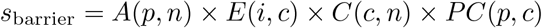

### D.4 Aggregation strategy

#### D.4.1 Collapse to best-per-controller

Multiple chains through the same controller are collapsed to avoid fake independence:

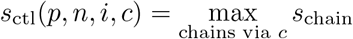

#### D.4.2 Top-K selection

Only the top *K* = 5 controllers are retained before aggregation:

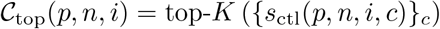

#### D.4.3 Normalized noisy-OR

Standard noisy-OR saturates quickly with multiple contributing causes. We use normalized noisy-OR:

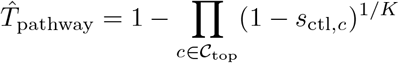

**Property**: If all *K* chains have score *s*, the result is exactly *s* (not ≈ 1).

#### D.4.4 Pathway combination

Latent and barrier pathways represent competing explanations, not additive probabilities:

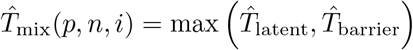

### D.5 Evaluation framework

#### D.5.1 The join direction problem

A critical methodological consideration: evaluation must start from the full observed tensor, not from predictions. Joining predictions to observations (rather than vice versa) would hide false negative and true negative mass, artificially inflating apparent performance. All evaluations use observation-first joins.

#### D.5.2 Binning to discrete rubric

Both predictions and observations were projected to coarse bins (multiples of 5: 0, 5, 10, …, 100) to match the discrete clinical scoring rubric:

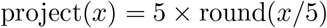

#### D.5.3 Three-way evaluation

Because zero clinical scores may represent either “true negative” or “not studied,” we employed three complementary evaluation frameworks:

**Evaluation 1: STRICT** (all zeros = true negatives)

- Most conservative/pessimistic
- Treats every zero as evidence of absence
- Metrics: Pearson *r*, Spearman *ρ*, Cohen’s weighted *κ*, F1, balanced accuracy

**Evaluation 2: PU FRAMING** (zeros = unlabeled)

- Positive-Unlabeled learning perspective
- Only positive observations are labeled; zeros are unknown
- Metrics: Precision@K, Recall@K, fold-enrichment Fold-enrichment is defined as:

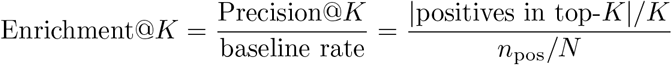

**Evaluation 3: CONFIDENCE-MASKED** (low-confidence zeros = missing)

- Uses evidence confidence to distinguish:
  – **Confident zeros**: Curated evidence of absence (retain as TN)
  – **Uncertain zeros**: Absence of evidence (mask out)
- Evaluate only on confident subset

**Finding**: All zeros in the clinical tensor have low confidence scores, meaning there is no curated evidence of absence. This strongly supports the PU framing interpretation.

#### D.5.4 Cohen’s weighted kappa

For ordinal agreement between binned predictions and observations:

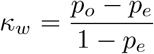

where *p*_*o*_ and *p*_*e*_ are weighted observed and expected agreement using linear weights:

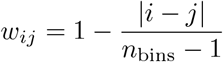

Interpretation scale: *<* 0 (less than chance), 0.0–0.2 (slight), 0.2–0.4 (fair), 0.4–0.6 (moderate), 0.6–0.8 (substantial), 0.8–1.0 (almost perfect).

### D.6 Results summary

#### D.6.1 Dataset statistics

| Metric | Value |
| --- | --- |
| Full tensor cells | 17,329 |
| Cells with mechanistic predictions | 1,522 (8.8%) |
| Cells without predictions | 15,807 (91.2%) |
| Known positives ($T_{\text{obs}} > 0$) | 3,425 (19.8%) |

#### D.6.2 Pathway dominance

| Dominant Pathway | Count | Percentage |
| --- | --- | --- |
| Barrier | 890 | 58.5% |
| Latent | 283 | 18.6% |
| Equal | 349 | 22.9% |

#### D.6.3 Overall performance (STRICT evaluation)

| Metric | Value |
| --- | --- |
| Pearson $r$ | 0.349 |
| Spearman $\rho$ | 0.299 |
| Weighted $\kappa$ | 0.236 (Fair) |
| Recall | 0.197 |
| Specificity | 0.974 |
| Precision | 0.421 |
| F1 | 0.259 |
| Balanced Accuracy | 0.586 |

#### D.6.4 Per-archetype performance

| Archetype | $n$ | Pearson $r$ | Spearman $\rho$ | F1 |
| --- | --- | --- | --- | --- |
| Keystone | 2,821 | 0.388 | 0.357 | 0.301 |
| Opportunistic | 6,045 | 0.355 | 0.316 | 0.248 |
| Multi-host | 4,030 | 0.333 | 0.305 | 0.219 |
| Specialist | 4,433 | 0.325 | 0.232 | 0.279 |

#### D.6.5 Enrichment analysis (PU framing)

| Top K% | K | Positives Found | Precision@K | Enrichment |
| --- | --- | --- | --- | --- |
| 1% | 173 | 51 | 0.29 | 2.9× |
| 5% | 866 | 256 | 0.30 | 2.9× |
| 10% | 1,733 | 508 | 0.29 | 2.9× |
| 20% | 3,466 | 870 | 0.25 | 2.5× |

#### D.6.6 2D projection performance

| Projection | Pearson $r$ | Spearman $\rho$ |
| --- | --- | --- |
| PI (Pathogen × Immune) | 0.359 | 0.374 |
| PN (Pathogen × Niche) | 0.583 | 0.555 |

### D.7 S3.7 Interpretation of “predicted high, observed zero”

Cases where 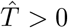 but *T*_obs_ = 0 are not necessarily model errors. They may indicate:

1. **Not studied**: The pathogen–niche–perturbation combination was never investigated
2. **Not detected**: Low statistical power or inappropriate study cohort
3. **No opportunity**: Pathogen not present in studied population
4. **True model miss**: Missing ecological constraints (temperature, microbiome, coinfections)

The finding that all clinical zeros have low confidence strongly supports interpretations 1–3 over interpretation 4.

