## Supplementary Material: Expanded Organism Profiles in the Context of Keystone Theory for "Co-evolved Partners of Immunity: A Trait-Based Map of Human Keystone Organisms" (from another version of this preprint: version 2, 2025-12-30)

**Keystone Organism Selection**

To systematically identify species-specific organisms that may have played a critical role in coordinating human immune function, we applied a structured selection process based on species-specificity, long-term co-evolution with human immunity, infection persistence, and multi-arm immune coordination requirements. Human-adapted organisms with evidence of shaping immune receptor evolution (e.g., HLA, TCR, NK receptors) were prioritized. The final selection included 27 putative keystone organisms spanning viral, bacterial, fungal, protozoan, helminthic, and ectoparasitic infections. However, certain environmentally ubiquitous infections, particularly opportunistic fungi and mycobacteria, may also play a significant role in immune coordination despite not being strictly species-specific. Therefore, a second list of 16 second-order keystone candidates was compiled to capture organisms that are frequently encountered by the immune system due to their environmental ubiquity or intermittent exposure but may lack strict human specificity.

### CMV: Cytomegalovirus (HHV-5)

**Classification and Transmission Ecology:** Human Cytomegalovirus (CMV), a double-stranded DNA virus, belongs to the Herpesviridae family. Classifications within this family are defined based on genomic and biological characteristics, with CMV specifically categorized under β-herpesviruses (Ruan et al., 2023). A hallmark of CMV is its ability to establish lifelong latency after primary infection, with the potential for reactivation during subsequent periods of immunosuppression, which underscores the virus's complex relationship with the host immune system (Zuhair et al., 2019). The transmission routes for CMV are diverse, including saliva, urine, breast milk, blood transfusions, sexual contact, and organ transplantation, thereby facilitating widespread dissemination of the virus within populations (Zuhair et al., 2019) (Nisbet et al., 2018).

**Clinical Manifestations and Latency:** In immunocompetent individuals, CMV infections are often asymptomatic, though some may develop a mononucleosis-like syndrome characterized by fever, fatigue, and lymphadenopathy (Klenerman & Oxenius, 2016). Conversely, congenital CMV infections can lead to severe outcomes such as microcephaly and sensorineural hearing loss in neonates (Cannon et al., 2010). In immunosuppressed populations, such as transplant recipients and those living with HIV/AIDS, CMV can precipitate serious complications, including retinitis, colitis, and pneumonia, significantly affecting morbidity and mortality rates (Griffiths et al., 2014) (Brito et al., 2019). Latency is primarily established within monocytes and endothelial cells, with CMV residing in these cells until reactivation occurs, typically in the context of immune system impairment (Zuhair et al., 2019).

**Global Distribution and Seroprevalence:** CMV is regarded as a ubiquitous virus with a global seroprevalence ranging drastically, influenced by factors such as age, geography, and socioeconomic status. Studies have indicated seroprevalence rates between 40% and 100%, generally exceeding 90% in lower-income countries while remaining at 50–70% in higher-income regions (Zuhair et al., 2019) (Fowler et al., 2022). This variability underscores the epidemiological importance of CMV, particularly in maternal and pediatric health, especially where infection risk is exacerbated by living conditions (Stern-Ginossar et al., 2012).

**Risk Factors for Opportunism:** The most prominent risk groups for CMV-associated complications include organ transplant recipients, individuals with HIV/AIDS, particularly those with significantly depressed CD4+ T-cell counts, and patients undergoing therapy with immunomodulators (Zuhair et al., 2019) (Omosigho et al., 2022). Additional vulnerable populations include infants with congenital infections as well as individuals with congenital or acquired T-cell deficiencies, who present a heightened risk for severe CMV disease (Griffiths et al., 2014) (Omosigho et al., 2022).

**Pathogenesis and Persistence Mechanisms:** CMV exhibits a broad tissue tropism, capable of infecting various cells, including myeloid progenitors, leading to its establishment of latency. The virus employs several immune evasion strategies, such as the interference with Major Histocompatibility Complex (MHC) class I antigen presentation through proteins like US3 and US6 (Ruan et al., 2023). This persistence mechanism allows CMV to avoid robust immune clearance while remaining subject to periodic reactivation controlled primarily by CD8+ T cells and natural killer (NK) cells (Bate et al., 2010) (Fowler et al., 2022).

**Immune Arm Dependencies:** Effective control of CMV infection is significantly reliant on a robust immune response involving CD8+ T cells, CD4+ T cells, NK cells, and humoral immunity (Klenerman & Oxenius, 2016). CD8+ T cells, particularly those targeting immediate-early antigens, play a vital role in mediating antiviral responses. Recent studies suggest that in immunocompromised hosts, diminished CMV-specific T-cell responses correlate with an increased risk of disease onset, reinforcing the importance of these adaptive immune mechanisms (Klenerman & Oxenius, 2016) (Bate et al., 2010).

**Compartmental Tropism and Failure:** In cases of congenital CMV, the central nervous system, cochlea, and liver become primary targets of infection, resulting in significant morbidities (Griffiths et al., 2014). Among immunosuppressed adults, critical organs affected include the lungs, via pneumonitis; the gastrointestinal tract, leading to enterocolitis; and the retina, resulting in retinitis, demonstrating the virus's broad pathogenic potential (Brito et al., 2019) (Nanakaly & Hussen, 2019). Latent CMV chiefly persists within bone marrow tissues and endothelial cells, with reactivation events tightly linked to episodes of myelosuppression or immune modulation (Ruan et al., 2023) (Fowler et al., 2022).

**Immune Modulation and Evasion:** CMV's ability to evade host immune surveillance involves the secretion of multiple immune-modulatory proteins that inhibit effective MHC presentation and alter NK cell functionality (Klenerman & Oxenius, 2016) (Ruan et al., 2023). Such mechanisms of immune evasion permit CMV to maintain a stealth-like presence within the host, often without triggering significant disease manifestations in immunocompetent individuals (Moseley et al., 2022).

**Immune Memory and Coordination:** Infection with CMV has been shown to induce a phenomenon termed "memory inflation," where a substantial population of CMV-specific CD8+ T cells persists in the host over time (Bate et al., 2010). This memory inflation has implications for the overall composition of the adaptive immune response, particularly as it relates to immunosenescence and the balance of naïve versus memory T cell populations in aging individuals (Klenerman & Oxenius, 2016) (Tu & Rao, 2016).

**Role in Immune Architecture:** CMV serves as a model for the keystone epitope theory, structuring both CD8+ T-cell responses and the overall immune dominance hierarchies (Fowler et al., 2022). The persistent antigenic pressure exerted by CMV profoundly influences immune system architecture, shaping the evolutionary dynamics of immune response in various populations (Ruan et al., 2023) (Janković et al., 2023).

**Impact of Immunosuppression:** The immunosuppressive conditions seen in transplant recipients and individuals with AIDS significantly enhance the likelihood of CMV reactivation and associated disease processes. In transplant populations, CMV not only correlates with heightened risks for allograft rejection but also increases the complexity of managing post-transplant infections (Zuhair et al., 2019) (Omosigho et al., 2022). Likewise, AIDS patients exhibit common manifestations such as retinitis and colitis, highlighting the critical intersection of immunocompromised states and CMV pathogenesis (Klenerman & Oxenius, 2016) (Brito et al., 2019).

**Prevalence and Future Considerations:** Given CMV's ongoing status as a major public health concern, particularly in neonates and immune-compromised patients, concerted efforts towards vaccine development are underway. Current research focuses on evaluating various platforms, including subunit, vector-based, and mRNA vaccines, while also considering the implications of CMV on aging, T-cell exhaustion, and immune imprinting strategies to enhance clinical outcomes (Ross et al., 2018) (Omosigho et al., 2022).

#### HSV-1: Herpes Simplex Virus 1 (HHV-1)

**Classification and Transmission Ecology:** Herpes Simplex Virus 1 (HSV-1) is categorized as a double-stranded DNA virus belonging to the Herpesviridae family and primarily the Alphaherpesvirinae subfamily. The virus is prevalent globally, showing a high rate of infection across various populations, with estimates suggesting that 60% to 95% of the world's population is seropositive for HSV-1 (Rafael et al., 2022) (Looker et al., 2015). Transmission predominantly occurs through direct contact with mucosal secretions or infected saliva, which facilitates the virus's ability to spread effortlessly in familial and close-contact settings (Looker et al., 2015). Following initial mucocutaneous infection, HSV-1 travels retrograde along sensory neurons to establish latency in the trigeminal ganglia, with reactivation potentially leading to viral shedding or symptomatic recurrence, underscoring its adaptability (Rafael et al., 2022).

**Clinical Manifestations and Latency:** HSV-1 primarily manifests as orolabial herpes but is also recognized as a significant causative agent of sporadic fatal encephalitis, particularly affecting the temporal lobes of the brain (Ullah et al., 2024). Clinical reactivation can result in cold sores or, in immunocompromised individuals, lead to more severe disseminated diseases, including encephalitis, esophagitis, hepatitis, and pneumonitis (James & Kimberlin, 2015). Encephalitis presents typically with fever, altered mental status, and focal neurological signs, reinforcing the need for vigilance in at-risk populations (Ullah et al., 2024). The reactivation of HSV-1 from latency is influenced by various factors, including stress, immune suppression, and other physiological triggers (Rafael et al., 2022).

**Global Distribution and Seroprevalence:** Globally, it is estimated that approximately 3.7 billion individuals under 50 years are infected with HSV-1, indicating a staggering prevalence of around 60% to 95% in different demographics (Looker et al., 2015). The seroprevalence exceeds 90% in several regions of sub-Saharan Africa and Latin America. In contrast, high-income countries reflect a decline in seroprevalence to below 50% among younger adults, highlighting a shift in transmission dynamics and potential public health implications (Looker et al., 2015) (Nasrallah et al., 2020).

**Risk Factors for Opportunism:** Severe manifestations of HSV-1 are particularly common in vulnerable populations, including neonates, transplant recipients, individuals with HIV/AIDS, and patients undergoing immunomodulatory therapies such as corticosteroids and tumor necrosis factor inhibitors (James & Kimberlin, 2015). These risk factors are crucial for understanding the epidemiology and potential outcomes of HSV-1 infection within different populations, necessitating tailored healthcare strategies (G, 2015).

**Pathogenesis and Persistence Mechanisms:** The pathogenesis of HSV-1 involves the virus infecting epithelial cells and undertaking axonal transport to sensory ganglia, where latency is characterized by the expression of the latency-associated transcript (LAT). Reactivation is facilitated by anterograde viral movement and is modulated by external stressors and immune suppression (Rafael et al., 2022). HSV-1 evades innate immune responses by downregulating major histocompatibility complex (MHC) class I expression and interfering with interferon signaling pathways, maintaining its persistence within the host (Koujah et al., 2018).

**Immune Arm Dependencies:** Control of HSV-1 latency and its reactivation is reliant on tissue-resident CD8+ T cells and natural killer (NK) cells. These immune cells are essential for surveilling sensory ganglia and mucosal surfaces, thus limiting viral replication during both symptomatic and asymptomatic reactivations. Their presence emphasizes the importance of localized immune responses in managing HSV-1 infection.

**Compartmental Tropism and Failure:** HSV-1 is known for its specific tropism towards the orolabial mucosa and the central nervous system (CNS). In cases of herpes simplex encephalitis (HSE), the virus typically localizes in the temporal lobes, resulting in significant neurological damage marked by hemorrhagic necrosis (Ullah et al., 2024). In immunocompromised hosts, HSV-1 can also lead to visceral diseases, further complicating its management (Rafael et al., 2022).

**Immune Modulation and Evasion:** To evade host immune detection, HSV-1 employs mechanisms such as the ICP47 protein, which inhibits the function of the transporter associated with antigen processing (TAP), preventing effective antigen presentation via MHC class I molecules. Additionally, HSV-1 disrupts toll-like receptor signaling pathways and interferon responses, thereby dampening overall antiviral immunity.

**Immune Memory and Coordination:** The establishment of long-lived, tissue-resident CD8+ T-cell populations is critical for HSV-1 immunity. These cells perform local surveillance within the ganglia and mucosa, contributing to the maintenance of immune memory and limiting reactivation events. Continuous subclinical reactivation promotes the reinforcement of epitope-specific memory responses, indicating a complex relationship between HSV-1 and the host immune system.

**Role in Immune Architecture:** While HSV-1 does not induce systemic immune exhaustion, it plays a substantial role in shaping long-term immune memory through persistent engagement with both mucosal and neuronal immune compartments. The dominance of specific viral epitopes, particularly those associated with immediate-early and latency-associated proteins, can significantly influence T-cell repertoire selection and readiness against subsequent infections.

**Impact of Immunosuppression:** In immunocompromised individuals, HSV-1 poses a grave threat, leading to serious disseminated disease presentations, especially in those with primary immunodeficiencies, HIV/AIDS, or those undergoing organ transplants. Increased susceptibility to conditions such as encephalitis and visceral HSV disease has been documented in patients on immunosuppressive therapies (James & Kimberlin, 2015).

**Prevalence and Future Considerations:** Despite its widespread prevalence and generally benign nature in immunocompetent populations, HSV-1 remains a significant cause of viral encephalitis and recurrent mucosal disease globally. Future strategies targeting latency and enhancing immune response through vaccines and immunotherapy hold promise in controlling HSV-1 reactivation and transmission dynamics (Rafael et al., 2022) (Ullah et al., 2024).

#### HSV-2: Herpes Simplex Virus 2 (HHV-2)

**Classification and Transmission Ecology:** Herpes Simplex Virus Type 2 (HSV-2) is classified as a double-stranded DNA virus within the Herpesviridae family, closely related to HSV-1. Its primary mode of transmission is through sexual contact, which leads to mucocutaneous infections. Upon initial infection, HSV-2 establishes latency in sacral sensory ganglia, resulting in a persistent infection characterized by intermittent reactivation and viral shedding, which may occur symptomatically or asymptomatically (Zhu & Viejo-Borbolla, 2021) (Schiffer et al., 2011). Genetic analyses suggest that HSV-1 and HSV-2 diverged from a common ancestor that likely infected early hominins approximately 6 million years ago. This evolutionary pathway indicates that while both viruses are genetically related, their adaptation to different hosts has significantly influenced their pathogenesis and transmission patterns (Looker et al., 2017).

**Clinical Manifestations and Latency:** HSV-2 is recognized as the principal cause of genital herpes, commonly presenting with painful ulcers, dysuria, fever, and lymphadenopathy during initial outbreaks. The virus then enters a latent state, from which reactivation can lead to recurrent genital lesions, varying in frequency from frequent symptomatic episodes to occasional subclinical shedding (Schiffer et al., 2011). Notably, HSV-2 is implicated in severe mucocutaneous ulcerations in immunocompromised patients, particularly those co-infected with HIV/AIDS (Lee et al., 2012). This association underscores the clinical significance of HSV-2 in vulnerable populations; (Looker et al., 2017).

**Global Distribution and Seroprevalence:** The seroprevalence of HSV-2 varies globally, with the highest rates observed in sub-Saharan Africa (over 40%) and lower prevalence in high-income areas (10–25%) (Looker et al., 2017). Women, older adults, and individuals with multiple sexual partners demonstrate higher infection rates, reflecting the disease's opportunistic nature (Zhu & Viejo-Borbolla, 2021) (Lee et al., 2012). Epidemiological data indicate that HSV-2 poses a significant public health concern, highlighting the need for continued surveillance and intervention efforts to contain HSV-2 transmission, especially in regions with high HIV co-infection rates (Looker et al., 2017) (Johnston et al., 2011).

**Risk Factors for Opportunism:** Immunosuppression markedly elevates the risk of severe manifestations of HSV-2, particularly in individuals with HIV/AIDS, notably in those with CD4+ T-cell counts <100/µL (Al-Zubaidi et al., 2024). Immunosuppressive therapies, such as corticosteroids and TNF inhibitors, have been associated with increased HSV-2 reactivation, leading to atypical, severe disease presentations (Schiffer et al., 2011) (Lee et al., 2012). Understanding these risk factors is critical for managing HSV-2 in immunocompromised patients and tailoring therapeutic strategies accordingly (Looker et al., 2017).

**Pathogenesis and Persistence Mechanisms:** HSV-2 initiates infection by penetrating mucosal tissues, replicating within epithelial cells, and subsequently migrating to sensory ganglia where it establishes latency characterized by minimal viral gene expression (Zhu & Viejo-Borbolla, 2021). During latency, transcription of the latency-associated transcript (LAT) plays a crucial role in maintaining persistent infection while evading immune detection (Looker et al., 2017). Upon reactivation, HSV-2 utilizes anterograde transport to reactivate lesions, and mechanisms such as the ICP47 protein assist in evading the host immune response by inhibiting MHC class I peptide presentation (Schiffer et al., 2011).

**Immune Arm Dependencies:** Effective host control of HSV-2 relies on a coordinated immune response, involving tissue-resident CD8+ T cells and CD4+ T helper cells, along with natural killer (NK) cells. These immune components are essential to limit reactivation and control lesion formation (Al-Zubaidi et al., 2024). However, in immunocompromised individuals, particularly those with HIV, the efficacy of HSV-specific CD8+ T-cell responses is significantly impaired, leading to uncontrolled viral shedding and disease exacerbation (Johnston et al., 2011) (Lee et al., 2012).

**Compartmental Tropism and Failure:** While HSV-2 primarily infects the genital and anal mucosa, in immunocompromised hosts, the virus can disseminate to other regions, including the central nervous system (CNS) and lungs, leading to severe clinical manifestations (Al-Zubaidi et al., 2024). The presence of atypical ulcerations can complicate the clinical picture, sometimes mimicking malignancies, thereby necessitating a differential diagnostic approach in affected patients (Lee et al., 2012).

**Immune Modulation and Evasion:** HSV-2 employs multiple strategies to evade the host immune system, including interfering with toll-like receptor signaling and inhibiting type I interferon responses (Schiffer et al., 2011) (Johnston et al., 2011). Immune control during latency is maintained by non-cytolytic mechanisms of CD8+ T cells, which suppress viral gene expression without destroying the neurons harboring latent virus (Zhu & Viejo-Borbolla, 2021) (Al-Zubaidi et al., 2024).

**Immune Memory and Coordination:** The formation of long-lived tissue-resident memory T cells (TRM) is a crucial aspect of HSV-2 immunology, providing rapid local responses to reactivation episodes (Johnston et al., 2011). These memory cells perpetuate immune training within mucosal tissues, forming an essential component of protective immunity against HSV-2 reactivation (Schiffer et al., 2011).

**Role in Immune Architecture:** HSV-2 serves as a critical keystone virus, establishing lifelong latency and prompting compartmentalized immune engagements, which reinforce immunological memory and responses at mucosal barriers (Johnston et al., 2011) (Lee et al., 2012). The sustained presence of viral antigens aids in the development of effective long-term immune responses, underlining the virus's impact on immune system architecture (Looker et al., 2017).

**Impact of Immunosuppression:** Severe immunosuppression, particularly in HIV/AIDS patients, often results in atypical manifestations of HSV-2, such as extensive ulcerations and systemic dissemination (Al-Zubaidi et al., 2024). This association underscores the virus's role as an opportunistic pathogen, significantly complicating the clinical management of co-infected individuals (Looker et al., 2017) (Johnston et al., 2011).

**Prevalence and Future Considerations:** HSV-2 remains a substantial global health issue, being a significant cofactor in HIV transmission (Looker et al., 2017) (Johnston et al., 2011). Its pattern of lifelong latency combined with immune evasion mechanisms presents unique challenges and opportunities for vaccine development. Current research efforts aim to disrupt latency, boost TRM responses, and develop effective antiviral immunotherapies to mitigate the risks associated with HSV-2 (Zhu & Viejo-Borbolla, 2021) (Schiffer et al., 2011).

#### VZV: Varicella-Zoster Virus (HHV-3)

**Classification and Transmission Ecology:** Varicella-Zoster Virus (VZV), a member of the alpha-herpesvirus subfamily, is primarily transmitted via respiratory droplets or direct contact with the fluid from vesicular lesions (Gershon et al., 2015) (Gilden et al., 2003). Following initial infection, which manifests as varicella (chickenpox), the virus establishes lifelong latency within the sensory neurons located in cranial and dorsal root ganglia (Gilden et al., 2003) (Arvin et al.,

2010). This persistence in the nervous system is crucial for the virus's survival, as it remains undetectable to the immune system and can reactivate years later as herpes zoster (shingles) (Gershon et al., 2015) (Gilden et al., 2003). The process of virus reactivation is complex and involves various factors, including the age and immune status of the host, leading to the re-emergence of the pathogen into the skin along the affected dermatome (Arvin et al., 2010) (Haque et al., 2014).

**Clinical Manifestations and Latency:** The clinical presentation of varicella is characterized by a generalized vesicular rash that transitions through macules, papules, vesicles, and eventually crusts (Gilden et al., 2003). Upon reactivation, symptoms include a painful dermatomal rash that can be accompanied by severe pain known as postherpetic neuralgia (Gershon et al., 2015) (Haque et al., 2014). Complications arising from VZV infections, particularly in immunocompromised patients, may include encephalitis, myelitis, and other severe complications (Arvin et al., 2010) (Warren-Gash et al., 2017). Postherpetic neuralgia presents as a chronic pain condition that can significantly impair the quality of life for older adults and those with weakened immune defenses (Gershon et al., 2015) (Arvin et al., 2010).

**Global Distribution and Seroprevalence:** Historically, greater than 95% of children in temperate climates were reported to acquire VZV by adolescence, emphasizing the virus's high seroprevalence prior to the introduction of vaccination programs (Arvin et al., 2010) (Ouwendijk & Georges, 2014). In tropical regions, infections typically occur later, suggesting environmental and demographic influences on the pathology of varicella (Arvin et al., 2010). Since the launch of universal vaccination initiatives, such as those in the United States starting in 1995, there have been notable decreases in the incidence rates of both varicella and herpes zoster, alongside corresponding declines in associated complications (Arvin et al., 2010) (Warren-Gash et al., 2017).

**Risk Factors for Opportunism:** The likelihood of reactivation of VZV significantly rises with advancing age, attributable to immunosenescence (Haque et al., 2014) (Wang et al., 2016). High-risk groups, including individuals with HIV, those undergoing chemotherapy, and patients receiving immunosuppressive therapies (such as corticosteroids and TNF inhibitors), are particularly vulnerable to severe manifestations of herpes zoster (Haque et al., 2014) (Wang et al., 2016) (Tsiouma, 2023). The immunological landscape, especially in aging populations, underscores the necessity for enhanced surveillance and preemptive measures to prevent complications related to reactivation (Wang et al., 2016) (Tsiouma, 2023).

**Pathogenesis and Persistence Mechanisms:** Upon respiratory infection, VZV initially undergoes viremic dissemination, subsequently localizing within neuronal cells (Arvin et al., 2010) (Gerada et al., 2020). The mechanisms allowing VZV to establish latency involve sophisticated evasion tactics against the host immune response, such as the downregulation of MHC class I molecules and the suppression of interferon signaling pathways through specific viral proteins (Arvin et al., 2010) (Gerada et al., 2020). Reactivation is believed to occur via anterograde axonal transport, with the virus traveling along neural pathways to reinfect skin and mucosal surfaces (Arvin et al., 2010) (Gerada et al., 2020).

**Immune Arm Dependencies:** The control of VZV reactivation relies heavily upon the coordinated activity of CD4+ T cells, CD8+ T cells, and natural killer (NK) cells (Wang et al., 2016) (Gilden et al., 2003). In particular, memory CD4+ T cells play a vital role in curtailing reactivation events, with vaccination correlating with enhancement of these protective immune responses, demonstrating a significant interplay between vaccination status and immune resilience against herpes zoster (Gilden et al., 2003) (Haque et al., 2014).

**Compartmental Tropism and Failure:** While VZV predominantly persists in sensory ganglia, instances of reactivation can lead to manifestations in the central nervous system and other areas, sometimes occurring without the classic rash associated with herpes zoster (Wang et al., 2016) (Tsiouma, 2023). Such atypical presentations pose diagnostic challenges and complicate the clinical understanding of VZV reactivation (Wang et al., 2016) (Tsiouma, 2023).

**Immune Modulation and Evasion:** VZV has evolved various strategies to evade immune detection, primarily by disrupting antigen presentation mechanisms and undermining interferon signaling. These adaptations facilitate the virus's ability to maintain latency and reactivate while avoiding significant inflammation (Arvin et al., 2010) (Wang et al., 2016).

**Immune Memory and Coordination:** The viral-specific T-cell memory induced by VZV can persist for decades, implying long-term immune surveillance (Gilden et al., 2003) (Tsiouma, 2023). Vaccination strategies that enhance the numbers and efficacy of these T-cell populations may reduce both the rates of herpes zoster and associated complications (Gilden et al., 2003; Haque et al., 2014). Recent evidence suggests these memory T cells might also confer protective effects against other herpesviruses through cross-reactivity (Wang et al., 2016) (Haque et al., 2014).

**Role in Immune Architecture:** VZV plays a pivotal role in shaping the host's immune architecture due to its ability to engage T-cell memory and coordinate neuroimmune responses over extensive periods (Arvin et al., 2010) (Gilden et al., 2003). Its unique capacity for maintaining a low-profile residency in the nervous system, combined with periodic reactivation, may significantly influence immunological interactions within both mucosal and neuronal tissues (Arvin et al., 2010) (Gerada et al., 2020).

**Impact of Immunosuppression:** In populations with compromised immune systems, the spectrum of VZV-related complications broadens significantly, frequently leading to severe manifestations such as pneumonia and hepatitis. The presentation may also include atypical forms of herpes zoster characterized by the absence of a rash (Wang et al., 2016) (Tsiouma, 2023). These complex disease patterns necessitate meticulous monitoring and tailored therapeutic strategies to manage reactivation in vulnerable patient cohorts (Wang et al., 2016) (Tsiouma, 2023).

**Prevalence and Future Considerations:** Despite the profound impact of vaccination on reducing varicella and herpes zoster incidence, reactivation remains a pressing clinical consideration, particularly in aging and immunosuppressed demographics (Warren-Gash et al., 2017) (Haque et al., 2014). Ongoing investigations into the mechanisms underpinning latency, immune evasion, and neuroimmune interactions hold promise for the next generation of vaccination strategies and antiviral therapies, with an emphasis on enhancing resident memory T-cell functions to mitigate potential complications (Gilden et al., 2003) (Warren-Gash et al., 2017).

#### EBV: Epstein-Barr Virus (HHV-4)

**Classification and Transmission Ecology:** Epstein-Barr Virus (EBV), classified as human gammaherpesvirus (HHV-4), predominantly infects B lymphocytes and epithelial cells. The virus is primarily transmitted through oral secretions, a mode of spread that facilitates infection among adolescents and young adults, often referred to as the "kissing disease" (Barton et al.,

2011). Following primary infection, EBV establishes a state of lifelong latency within memory B cells, which can be reactivated under conditions of immune stress, indicating its evasion of the immune response and potential for periodic clinical manifestations (Zalani et al., 1996). The mechanisms of latency and activation are complex, involving viral genes that transition the virus into a replicative state to maintain long-term infection in the host (Countryman & Miller, 1985) (Linnstaedt et al., 2010).

**Clinical Manifestations and Latency:** Primary EBV infection is often asymptomatic; however, in adolescents and young adults, it may present as infectious mononucleosis characterized by fever, lymphadenopathy, pharyngitis, and fatigue (Virgin et al., 1997). The likelihood of symptomatic infection increases with age, particularly among cases where immune responses are compromised. In immunosuppressed individuals, reactivation of EBV can lead to severe manifestations, including post-transplant lymphoproliferative disorder (PTLD) and various lymphomas. PTLD is frequently a consequence of uncontrolled proliferation of EBV-infected B cells due to ineffective immune surveillance (Zalani et al., 1996) (Lü et al., 2008). The relationship between EBV and various cancers underscores the need for vigilance in high-risk groups affected by immunosuppression (Lü et al., 2008) (Cao et al., 2012).\

**Global Distribution and Seroprevalence:** EBV is highly prevalent, with a seroprevalence exceeding 90% worldwide. In lower-income regions, initial infections occur during early childhood, often presenting as asymptomatic, while in higher-income countries, infections tend to occur later, leading to symptomatic disease, particularly in older adolescents and adults (Virgin et al., 1997). This variation in seroprevalence and symptomology correlates with socio-economic factors, where environmental and healthcare access play crucial roles (Zalani et al., 1996).

**Risk Factors for Opportunism:** Several risk factors are critical for the opportunistic reactivation of EBV. Immunosuppression is the primary risk factor, particularly in transplant recipients, individuals with HIV/AIDS, and those with primary immunodeficiencies. Impaired natural killer (NK) cell function, observed in various genetic conditions, increases susceptibility to severe EBV diseases (Sun et al., 1998) (Zalani et al., 1996). Additionally, specific HLA polymorphisms may impair effective immune surveillance against EBV, indicating the complexity of host immune factors in determining the risk of EBV-related diseases (Barton et al., 2011) (Zalani et al., 1996).

**Pathogenesis and Persistence Mechanisms:** EBV initiates its pathogenesis by infecting naive B cells, inducing their proliferation through latent membrane proteins (LMPs), notably LMP1 and LMP2, which mimic cellular signaling pathways enhancing cell survival and proliferation (Sun et al., 1998) (Linnstaedt et al., 2010). Maintenance of latency is critically dependent on the expression of the EBNA-1 protein, which allows the virus to evade CD8+ T cell-mediated immune responses by inhibiting MHC class I processing (Zalani et al., 1996) (Lü et al., 2008). Additionally, viral microRNAs play significant roles in modulating host cell responses and promoting persistent infection (Sun et al., 1998) (Lü et al., 2008).

**Immune Arm Dependencies:** Effective control of EBV infection heavily relies on the immune system's capacity, particularly the roles of cytotoxic CD8+ T cells, CD4+ T helper cells, and functional NK cells. CD8+ T cells are essential for recognizing lytic and latent EBV antigens, crucial for controlling reactivation of the virus. NK cells also play a vital role during the early phase of infection, preventing unchecked expansion of latently infected B cells (Zalani et al., 1996) (Lü et al., 2008). The interplay among these immune components demonstrates the sophisticated response required to maintain control over EBV and prevent reactivation under stress (Barton et al., 2011) (Linnstaedt et al., 2010).

**Compartmental Tropism and Failure:** EBV has a distinct tropism for B cells located within secondary lymphoid organs. Upon reactivation or under conditions of immunodeficiency, proliferation of EBV-infected cells can extend to extranodal sites, such as the CNS, liver, and gastrointestinal tract. Such manifestations can be severe; for instance, CNS-involved PTLD is particularly aggressive and associated with high EBV loads in cerebrospinal fluid, indicating the virus's potential for widespread effects beyond its traditional sites (Doja et al., 2006) (Cao et al., 2012) (Wu et al., 2000).

**Immune Modulation and Evasion:** EBV employs extensive immune evasion strategies. This includes the function of EBNA-1, which inhibits MHC class I processing, and the mimicry of host signaling pathways by LMP1 and LMP2 (Sun et al., 1998) (Zalani et al., 1996). Viral microRNAs dampen antiviral immune signaling and T cell-mediated clearance, facilitating a favorable environment for EBV persistence (Lü et al., 2008) (Zalani et al., 1996).

**Immune Memory and Coordination:** EBV substantially influences the establishment of long-term memory T-cell responses, predominantly through CD8+ effector-memory T cells that target EBNA3 and LMP antigens. This continuous immune engagement not only maintains tolerance to the virus but also shapes broader T-cell responses to other unrelated antigens, emphasizing the virus's role in modulating the immune landscape (Linnstaedt et al., 2010) (Cao et al., 2012).

**Role in Immune Architecture:** EBV acts as a keystone pathogen, intricately shaping immune architecture. Its diverse antigenic portfolio, combined with its ability to establish latency, enables it to calibrate the immune system's hierarchy. The presence of EBV nuclear and membrane proteins as immunodominant epitope hubs fosters sustained immune surveillance while potentially influencing immune responses against other pathogens (Zalani et al., 1996) (Cao et al., 2012).

**Impact of Immunosuppression:** The impact of immunosuppression on EBV dynamics is profound. Conditions such as T-cell depletion in transplant recipients or advanced HIV severely compromise the host's ability to control EBV. Consequently, unchecked proliferation of EBV-infected B cells can result in severe outcomes like PTLD. Additionally, intrinsic defects in NK cell activity and genetic conditions can predispose individuals to more severe EBV-related complications (Zalani et al., 1996) (Lü et al., 2008) (Cao et al., 2012).

**Prevalence and Future Considerations:** The association of EBV with lymphomas and immune dysregulation is increasingly recognized, particularly in immunocompromised and aging populations. Its ability to manipulate immune memory and evade clearance positions EBV as a significant target for future therapeutic strategies, including vaccine development directed at EBV's envelope and latency proteins. Future research will explore EBV's role in autoimmune pathogenesis and its broader implications in shaping human immune memory across varying life stages (Virgin et al., 1997) (Zalani et al., 1996) (Cao et al., 2012).

#### HHV-6: Human Herpesvirus 6

**Classification and Transmission Ecology:** Human Herpesvirus 6 (HHV-6) is categorized as a betaherpesvirus within the Roseolovirus genus, comprising two distinct species: HHV-6A and HHV-6B. HHV-6 is primarily transmitted through direct person-to-person contact, particularly via maternal saliva to infants (Eliassen et al., 2017) (Hanson et al., 2018). Global epidemiological studies indicate that HHV-6B is the most prevalent species worldwide, while HHV-6A is more frequently isolated from immunocompromised individuals and specific geographical regions, particularly in Africa and Asia (Eliassen et al., 2017) (Nastke et al., 2012). The differences in prevalence and virulence between the two species highlight the complexity of their interactions within varied human populations.

**Clinical Manifestations and Latency:** The primary infection of HHV-6B generally occurs between 6 and 24 months of age, resulting in a common childhood illness known as roseola infantum or exanthem subitum, which typically manifests with high fevers and may lead to rash and febrile seizures (Hanson et al., 2018) (Nefzi et al., 2016). Following this initial infection, HHV-6 establishes a lifelong latent infection, predominantly in immune cells such as CD4+ T cells and monocytes, and can, in rare instances, integrate into human chromosomes—this phenomenon is known as inherited chromosomally integrated HHV-6 (iciHHV-6) (Gulve et al., 2017) (Strenger et al., 2016). The persistence within the host emphasizes the virus's subtleties in affecting long-term health outcomes.

**Global Distribution and Seroprevalence:** Seroprevalence studies demonstrate that greater than 90% of the population shows evidence of HHV-6 infection by the age of two, highlighting its ubiquity (Eliassen et al., 2017) (Hanson et al., 2018). While HHV-6B remains globally dominant, HHV-6A's increased frequency in immunosuppressed individuals and specific populations, especially in sub-Saharan Africa, indicates differing epidemiological dynamics (Nastke et al., 2012). Such insights into seroprevalence are crucial for understanding the long-term impacts of early-life infections and their correlation with later health conditions.

**Risk Factors for Opportunism:** The reactivation of HHV-6 most commonly occurs in patients undergoing allogeneic hematopoietic stem cell transplantation (HCT), particularly in those receiving umbilical cord blood, myeloablative conditioning, or glucocorticoids (Scheurer et al., 2012) (Hanson et al., 2018) (Caselli et al., 2020). Other risk factors include T-cell depletion and delayed recovery of natural killer (NK) cell populations, which are vital for controlling viral reactivations (Hanson et al., 2018) (Nefzi et al., 2016). Although patients receiving CAR-T cell therapy generally show lower rates of HHV-6 reactivation, instances of HHV-6 encephalitis have been documented (Scheurer et al., 2012) (Hanson et al., 2018). Furthermore, in the context of solid organ transplantation, HHV-6 has been associated with serious conditions such as pneumonia and graft rejection (Scheurer et al., 2012) (Hanson et al., 2018) (Caselli et al., 2020).

**Pathogenesis and Persistence Mechanisms:** HHV-6A and HHV-6B utilize different receptors to infiltrate CD4+ T cells; specifically, HHV-6A utilizes CD46, while HHV-6B engages CD134 (Eliassen et al., 2017) (Rizzo & Luca, 2018). The virus exhibits a slow replication cycle, leading to notable cytopathic effects and resulting in a marked downregulation of key immune markers such as CD3, CD4, CD45, and MHC-I, which impairs the host's immune response to the infection (Li et al., 2011) (Rizzo & Luca, 2018). These alterations in T-cell populations are important for understanding virus-induced immunosuppression.

**Immune Arm Dependencies:** Efforts to manage HHV-6 reactivation rely on the coordinated action of CD4+ and CD8+ T cells, NK cells, and the production of interferon-mediated responses. Critical cytokines, such as IL-1β, TNF-α, and IFN-γ, are essential for enhancing innate and adaptive immune responses (Rizzo & Luca, 2018) (Arena et al., 2001). Failures in these immune pathways can lead to compartment-specific reactivations of HHV-6, producing a variety of clinical manifestations depending on which immune arm is compromised (Eliassen et al., 2017) (Rizzo & Luca, 2018).

**Compartmental Tropism and Failure:** Reactivation of HHV-6 is notably prevalent across several compartments: central nervous system (CNS) (notably causing limbic encephalitis), pulmonary (pneumonitis), hematopoietic (bone marrow suppression), and hepatic structures (hepatitis) (Hanson et al., 2018) (Arena et al., 2001). Each compartment's specific immune suppression profile correlates with the distinct immune deficiencies present in those sites, including the depletion of CD4+ and NK cells (Rizzo & Luca, 2018) (Arena et al., 2001).

**Immune Modulation and Evasion:** The mechanisms of immune evasion employed by HHV-6 are multifaceted. The U94 protein is instrumental for chromosomal integration and latency, while the U20 and U24 proteins inhibit NK cell activation and disrupt CD4+ T-cell receptor recycling, respectively (Eliassen et al., 2017) (Ota et al., 2014) (Caselli et al., 2020). Such immune modulation strategies emphasize the virus's ability to persist in the host while undermining the immune surveillance mechanisms.

**Immune Memory and Coordination:** Chronic exposure to HHV-6 has been shown to alter the landscape of T cells, specifically diminishing CD45 expression and affecting TCR (T-cell receptor) responsiveness (Rizzo & Luca, 2018) (Arena et al., 2001). The induction of NK cell activation through IL-15 suggests a complex interplay between innate and adaptive immune compartments, possibly leading to long-term memory responses that result in dysfunctional immunological profiles (Rizzo & Luca, 2018) (Fastenackels et al., 2019).

**Role in Immune Architecture:** HHV-6 may not directly initiate the hierarchies of immune responses; however, its chronic presence and associated latency likely fine-tune and reinforce existing immune architectures. This role may be critical in maintaining the coordination required for effective immune responses against other pathogens (Arena et al., 2001) (Vojdani et al., 2024).

**Impact of Immunosuppression:** In immunocompromised states, the reactivation of HHV-6 can lead to severe complications including encephalitis, pneumonitis, hepatitis, and significant marrow suppression, which collectively contribute to graft rejection and enhance morbidity rates in these populations (Scheurer et al., 2012) (Nefzi et al., 2016) (Caselli et al., 2020). The need for vigilant monitoring of HHV-6 reactivation in such patients is therefore imperative.

**Prevalence and Future Considerations:** HHV-6 remains one of the most widespread herpesviruses affecting humans, with over 90% seroprevalence by early childhood. Given its roles in immune modulation and potential associations with various disease states, there is a crucial need for further research into diagnosis, immune profiling, and targeted therapeutic interventions aimed at managing HHV-6 infections effectively (Scheurer et al., 2012) (Eliassen et al., 2017) (Hanson et al., 2018).

#### HHV-7: Human Herpesvirus 7

**Classification and Transmission Ecology**: Human Herpesvirus 7 (HHV-7) is classified as a ubiquitous betaherpesvirus within the Roseolovirus genus and is closely related to Human Herpesvirus 6 (HHV-6) (Agut et al., 2017). It was first identified and isolated in 1990 from the CD4+ T cells of healthy individuals, marking its classification as a human-adapted virus with no known environmental or zoonotic reservoirs Agut, 2017 #2888} (Denner et al., 2019). Primarily,

HHV-7 is transmitted through saliva, particularly within households, and tends to be acquired early in life, often between the ages of 2 and 5 years (Biganzoli et al., 2010) (Yamamoto et al., 2013) (Denner et al., 2019). Evidence suggests that transmission occurs frequently in familial settings, emphasizing the social aspect of the spread of this virus (Yamamoto et al., 2013).

**Clinical Manifestations and Latency**: The primary infection with HHV-7 typically occurs between the ages of 2 and 5 years (Biganzoli et al., 2010) (Denner et al., 2019). While most infections are asymptomatic, some cases present with febrile illnesses or roseola-like syndromes, which can lead to complications such as febrile seizures in children (Biganzoli et al., 2010). HHV-7 is also associated with severe central nervous system manifestations, including encephalitis and meningitis, particularly in immunocompromised individuals (Ongrádi et al., 2016). The virus demonstrates a propensity for latency in peripheral blood mononuclear cells, where it can remain dormant and may reactivate, especially during T-cell activation (Agut et al., 2017) (Ongrádi et al., 2016).

**Global Distribution and Seroprevalence**: HHV-7 exhibits a near-universal seroprevalence rate surpassing 95% in adults globally, indicating it is widely disseminated throughout human populations (Agut et al., 2017) (Zheng et al., 2021). The constant shedding of HHV-7 via saliva confirms its stability in human populations over time, with frequent viral loads detectable (Yamamoto et al., 2013). This transmission via saliva demonstrates the virus's ability to maintain persistence and reactivation potential within the host (Zheng et al., 2021).

**Risk Factors for Opportunism**: Immunosuppression significantly increases the risk of HHV-7 reactivation, particularly in populations such as hematopoietic stem cell transplant recipients and individuals with HIV/AIDS (Agut et al., 2017). In these immunocompromised patients, HHV-7 DNAemia has been observed, correlating with adverse outcomes such as prolonged neutropenia and a higher likelihood of co-infections, including cytomegalovirus (CMV) (Agut et al., 2017) (Ongrádi et al., 2016). This association highlights the opportunistic nature of HHV-7 in the context of weakened immune defenses.

**Pathogenesis and Persistence Mechanisms**: HHV-7 uniquely infects CD4+ T cells, using CD4 as a receptor during entry (Ongrádi et al., 2016) (Schneider & Hudson, 2011). The virus's immune evasion strategies include the production of the U21 glycoprotein, which binds to MHC class I molecules, redirecting them to lysosomes and impairing antigen presentation (Schneider & Hudson, 2011). Similarly, the U24 protein disrupts endosomal recycling and downregulates important T-cell surface markers, further promoting viral persistence (Schneider & Hudson, 2011). Reactivation of HHV-7 can be triggered by factors such as immune activation and stress, emphasizing the virus's adaptability in the host environment (Tuddenham et al., 2012).

**Immune Arm Dependencies**: Control of HHV-7 infections heavily relies on CD4+ T cell responses, underscoring the virus's tropism and reactivation dynamics (Agut et al., 2017) (Biganzoli et al., 2010). Natural Killer (NK) cells also play a role in combating HHV-7 through IL-15-mediated enhancement of their cytolytic activity, with increased levels of cytokines such as IFN-γ and TNF-α noted in infected individuals (Agut et al., 2017) (Tuddenham et al., 2012). This immune interaction underpins the virus's persistence and highlights the intricate balance between viral evasion and host immune activation.

**Compartmental Tropism and Failure**: Under conditions of immune suppression, HHV-7 can impact several bodily systems, including the central nervous system, manifesting as severe complications such as encephalitis (Ongrádi et al., 2016) (Tuddenham et al., 2012). Evidence suggests that HHV-7's failure to control these manifestations correlates strongly with CD4+ T cell dysfunction and concurrent CMV disease in transplant scenarios (Ongrádi et al., 2016).

**Immune Modulation and Evasion**: HHV-7 employs various mechanisms to modulate immune responses, including the U21 protein's role in diverting MHC class I to lysosomes (Ongrádi et al., 2016). This impairs appropriate immune recognition and response during active and latent phases of infection, illustrating the virus's capability to undermine host defenses (Tuddenham et al., 2012).

**Immune Memory and Coordination**: Persistence of HHV-7 appears to modify the host's cytokine networks, leading to sustained changes in T cell receptor signaling and immune memory profiles, potentially due to downregulation of CD45 (Tuddenham et al., 2012). Such long-term alterations may influence overall immune readiness and cross-reactivity, especially during subsequent infections.

**Role in Immune Architecture**: While less well characterized than HHV-6, HHV-7 may still play a significant role in shaping immune architectures through the expression of latency-associated proteins that enhance antigenic hierarchy and promote T cell activation (Agut et al., 2017) (Tuddenham et al., 2012). Future research should explore whether HHV-7 actively influences these immune responses or merely utilizes pre-existing frameworks.

**Impact of Immunosuppression**: HHV-7 reactivation in immunocompromised individuals has been correlated with severe neurological complications, heightened CMV outcomes, and delayed engraftment in transplant patients (Agut et al., 2017) (Tuddenham et al., 2012). The implications of such reactivation necessitate careful monitoring and management in at-risk populations to mitigate potential adverse effects.

**Prevalence and Future Considerations**: Given its high prevalence and significant immunomodulatory capabilities, HHV-7 warrants further investigation (Agut et al., 2017) (Tuddenham et al., 2012). Future studies should focus on its potential role in reinforcing immune calibration or contributing to immune misdirection, especially under stress. Currently, there are limited diagnostic and therapeutic options available, emphasizing the need for ongoing research, particularly noting its early acquisition and lifetime persistence as a key aspect of human immune interaction models.

#### TB: Mycobacterium tuberculosis

**Classification and Transmission Ecology:** Mycobacterium tuberculosis is classified within the Mycobacterium tuberculosis complex (MTBC), a group of closely related mycobacterial species that include M. bovis, M. africanum, and several animal-adapted lineages. While older models suggested a zoonotic origin from cattle, modern phylogenetic studies show that M. tuberculosis is likely the ancestral human-adapted member of the complex and that M. bovis likely evolved from a human strain (Comas et al., 2013). The bacterium is a slow-growing, acid-fast bacillus predominantly transmitted through airborne droplets when an infected person coughs or sneezes (Sia, 2012). Humans are the primary reservoir, and the lungs serve as the main site of infection. Transmission frequently results in various outcomes: immediate clearance of the pathogen, establishment of latent tuberculosis infection (LTBI), or progression to active disease. This organism has evolved mechanisms conducive to both efficient human-to-human transmission and long-term survival within hosts (Roca et al., 2022), reflecting its deep evolutionary relationship with humans dating back at least 70,000 years (Gagneux, 2012). Notably, the success of M. tuberculosis as a pathogen intensified following the Neolithic transition to agriculture and settled life, when increased population density and indoor crowding allowed for sustained chains of transmission (Hershberg et al., 2008). In contrast, in Paleolithic hunter-gatherer populations, TB likely persisted in a largely latent, low-prevalence state, with reactivation events occurring only under conditions of immunosuppression or seasonal aggregation, which limited its long-term transmission potential (Brites & Gagneux, 2015).

**Clinical Manifestations and Latency:** Clinically, tuberculosis can present as either LTBI or active TB disease, with pulmonary TB being the most prevalent form characterized by symptoms such as persistent cough, weight loss, night sweats, and fever (Sia, 2012). In some cases, TB can disseminate, leading to severe conditions like miliary TB or TB meningitis. Latent TB infection often remains asymptomatic for extended periods, sometimes reactivating years later; this reactivation risk is estimated at 5–10% over a lifetime, varying significantly among individuals, particularly those with compromised immune systems (Quesniaux et al., 2010). Importantly, evolutionary models suggest that this ability to remain latent and reactivate opportunistically in susceptible hosts has been key to the long-term survival of M. tuberculosis in small human populations over time (Gagneux, 2012).

**Global Distribution and Seroprevalence:** According to the World Health Organization (WHO), approximately 1.7 billion people, accounting for about 22% of the global population, are estimated to harbor latent TB infections, with the highest prevalence in regions like sub-Saharan Africa, India, and Southeast Asia (Roca et al., 2022). This staggering figure emphasizes the critical need for robust global health strategies focused on both prevention and treatment, especially in high-burden areas.

**Risk Factors for Opportunism:** High-risk populations for TB include individuals with comorbidities such as HIV/AIDS, certain malignancies, diabetes, chronic kidney disease, and malnutrition. Additionally, TNF inhibitors and corticosteroids contribute significantly to TB reactivation risk in patients treated for rheumatoid arthritis or similar conditions (Toussi et al., 2013) (Kireev et al., 2025) (Olleros et al., 2015). Genetic predisposition, particularly mutations in pathways related to IFN-γ and TNF signaling, has also been identified as enhancing susceptibility to TB infection (Kireev et al., 2025).

**Pathogenesis and Persistence Mechanisms:** The pathogenesis of TB involves a sophisticated interaction between M. tuberculosis and host immune responses. The bacterium employs several strategies to evade immune clearance—most notably, it inhibits phagosomal maturation, resists oxidative stress, and modulates T-cell activation (Li et al., 2015). The formation of granulomas helps contain the infection, serving as both a defensive measure and a niche for bacillary persistence (Quesniaux et al., 2010).

**Immune Arm Dependencies:** Host defense against TB primarily relies on the coordinated signaling mechanisms involving TNF-α, IL-12, and IFN-γ, which are crucial for granuloma formation and stabilization (Kireev et al., 2025). The dysregulation of TNF-α production, either excessive or insufficient, results in compromised containment of M. tuberculosis, leading to increased disease severity or reactivation of latent infections (Toussi et al., 2013) (Kireev et al., 2025).

**Compartmental Tropism and Failure:** Disseminated tuberculosis is more common in vulnerable populations, including young children and individuals with immunocompromising conditions (Olleros et al., 2015). The manifestations of TB may vary according to the anatomical compartments affected, with distinct immune responses delineating the clinical trajectories of the illness (Quesniaux et al., 2010).
**Immune Modulation and Evasion:** M. tuberculosis has evolved to deploy virulence factors that interfere with essential immune pathways, such as Toll-like receptor (TLR) signaling and autophagy, further bolstering its ability to survive and replicate in host cells (Natarajan & Narayanan, 2011). For instance, the pathogen secretes sulfolipids that facilitate immune evasion and promote its transmission (Natarajan & Narayanan, 2011).
**Immune Memory and Coordination:** Specific immune responses to TB are characterized by robust CD4+ and CD8+ T-cell activities. The presence of lung-resident memory T cells and macrophages, likely enhanced by Bacillus Calmette-Guérin (BCG) vaccination, underscores the importance of memory responses in TB control (Kireev et al., 2025).
**Role in Immune Architecture:** TB represents a model of immune coordination, where granuloma formation is a hallmark of integrated immune responses encompassing variously activated lymphocyte and macrophage interactions. This interplay contributes to host immunity and pathogenesis via selective memory and epitope targeting (Kireev et al., 2025).
**Impact of Immunosuppression:** Immunosuppressive therapies, especially TNF antagonists, significantly increase reactivation risks for LTBI (Ha et al., 2022). Specific primary immunodeficiencies that impact IFN-γ or IL-12 signaling also predispose individuals to disseminated forms of TB, reinforcing the complexity of immune interactions in the context of tuberculosis (Kireev et al., 2025) (Ha et al., 2022).
**Prevalence and Future Considerations:** Tuberculosis remains a leading cause of morbidity and mortality globally, with an estimated 10.6 million new cases reported in 2021 and more than 1.5 million deaths attributed to the disease (Roca et al., 2022). The persistence of high numbers of individuals with latent TB reinforces the urgency for novel vaccines and enhanced therapeutic strategies targeting both innate and adaptive immune responses (Kireev et al., 2025).

### KSHV: Kaposi’s Sarcoma-Associated Herpesvirus (HHV-8)

**Classification and Transmission Ecology:** Kaposi's Sarcoma-Associated Herpesvirus (KSHV), also known as human herpesvirus 8 (HHV-8), belongs to the gammaherpesvirus subfamily. It is primarily transmitted through saliva, with significant implications for its epidemiology and transmission dynamics (Ye et al., 2011) (Cai et al., 2010). In addition to person-to-person transmission via saliva, KSHV can also be transmitted through organ transplantation and blood transfusions—although these latter routes are considered rare—and via sexual contact, particularly among men who have sex with men (Ye et al., 2011) (Moore & Chang, 2003). In endemic regions such as sub-Saharan Africa, KSHV transmission commonly occurs in childhood, with maternal pre-mastication of food recognized as a potential transmission route (Poppe et al., 2020).

**Clinical Manifestations and Latency:** While primary infection with KSHV is often asymptomatic, it may manifest with fever, rash, or lymphadenopathy among immunocompetent individuals (Thakker & Verma, 2016). KSHV establishes latency primarily in endothelial cells and B cells, which plays a crucial role in the development of KSHV-associated conditions, such as Kaposi's sarcoma (KS), multicentric Castleman disease (MCD), and primary effusion lymphoma (PEL) (Ye et al., 2011) (Moore & Chang, 2003). The presence of KSHV during latency allows the virus to persist in the host and evade immune surveillance, which is pivotal for its malignant transformation capability (Cai et al., 2010) (Broussard & Damania, 2020).

**Global Distribution and Seroprevalence:** The geographic distribution of KSHV is not uniform, revealing significant seroprevalence variation across different regions. Research indicates that in sub-Saharan Africa, seroprevalence rates can exceed 50%. In contrast, the rates in Mediterranean populations are notably lower, ranging from 10–30%, while in North America, Asia, and Western Europe, seroprevalence usually remains under 10% (Zou et al., 2017) (Lidenge et al., 2018). Certain populations, such as Brazilian Amerindians and individuals in Papua New Guinea, exhibit exceptionally high KSHV seroprevalence rates, indicating localized hotspots of infection (Lidenge et al., 2018) (Ngalamika & Munsaka, 2020).

**Risk Factors for Opportunism:** KSHV is particularly associated with malignancies in individuals experiencing immunosuppression, such as those with HIV/AIDS or those undergoing organ transplantation (Thakker & Verma, 2016) (Tabtieng et al., 2018). The risk factors for the development of KSHV-associated diseases are augmented by genetic predispositions, including perforin gene mutations and OX40 deficiencies, which impair normal immune functions (Tabtieng et al., 2018). This immunosuppressed state creates a permissive environment for the development of KS and related malignancies, emphasizing the critical interplay between host immunity and viral pathogenesis (Thakker & Verma, 2016) (Shimoda et al., 2023).

**Pathogenesis and Persistence Mechanisms:** KSHV exhibits a unique life cycle featuring both latent and lytic phases, akin to other herpesviruses. During latency, it expresses a restricted set of genes including LANA (latency-associated nuclear antigen), vFLIP, and vCyclin, which are essential for maintaining the viral genome within host cells and facilitating angiogenesis and immune evasion (Cai et al., 2010) (Coscoy, 2007). Reactivation of KSHV from latency into the lytic phase can be triggered by various stressors such as hypoxia and immune activation signals (Tabtieng et al., 2018) (Leidal et al., 2012). Understanding these dynamics is vital for devising strategies for therapeutic interventions targeting KSHV-associated malignancies.

**Immune Arm Dependencies:** The control of KSHV infection heavily relies on T-cell-mediated immunity, with CD8+ cytotoxic T cells being particularly crucial. CD4+ T helper cells and NK cells also play supporting roles in the immune response (Damania & Münz, 2019) (Thakker & Verma, 2016). KSHV infection often leads to the generation of antibodies against both lytic antigens and latent proteins; however, it is the cellular immune response that predominantly suppresses viral replication and progression to associated disease states (Coscoy, 2007) (Damania & Münz, 2019). Deficits in T-cell immunity, rather than in humoral responses, strongly correlate with the progression of KSHV-associated diseases, underscoring the importance of effectively managing immune efficacy in these patients (Lagos et al., 2008).

**Compartmental Tropism and Failure:** KSHV exhibits compartmental tropism, with disease manifestations varying significantly by syndrome type: KS typically arises in the skin, mucosa, and various visceral organs; MCD predominantly affects lymphoid tissues; and PEL is observed in serous cavities (Zou et al., 2017) (Lidenge et al., 2018). The affected compartments often depend on the immunological status of the host, as both CD4+ and CD8+ T-cell losses are linked to unmasking and disease emergence (Srivastava et al., 2023) (Tabtieng et al., 2018).

**Immune Modulation and Evasion:** KSHV is adept at evading host immune detection through the expression of numerous viral proteins that mimic or disrupt human immune signals. For example, vFLIP activates NF-κB pathways to inhibit apoptosis, while vIL-6 supports B-cell survival (Coscoy, 2007) (Thakker & Verma, 2016) (Shimoda et al., 2023). Such immune modulation strategies contribute to a challenging landscape for therapeutic interventions, as they allow KSHV to persist and evade host defenses effectively (Lagos et al., 2008) (Shimoda et al., 2023).

**Immune Memory and Coordination:** Post-infection, KSHV can generate broadly specific CD8+ T-cell responses targeting various lytic cycle proteins, contributing to a state of chronic immune calibration. However, the durability of these responses and their coordinated compartmentalization in the face of immune pressure is an area requiring further investigation (Thakker & Verma, 2016) (Tabtieng et al., 2018). The expression of antigens during latent infection continuously engages the immune system, contributing to KSHV's ability to maintain presence in the host for extended durations (Tabtieng et al., 2018) (Cai et al., 2010).

Role in Immune Architecture: KSHV's expression of immunodominant proteins plays a potentially reinforcing role in shaping local immune hierarchies, although definitive proof of this role in initiating epitope hierarchies is still lacking (Thakker & Verma, 2016) (Damania & Münz, 2019). The virus's capacity to modulate CD8+, CD4+ T-cell, and NK cell interactions suggests a contributory role in local immune coordination, affecting overall immune architecture (Thakker & Verma, 2016) (Lingel et al., 2016).

**Impact of Immunosuppression:** In scenarios of profound T-cell dysfunction, such as in HIV/AIDS or post-transplant patients, KSHV replication can become unchecked, leading to more severe manifestations of associated diseases, including KS, MCD, and PEL (Tabtieng et al., 2018) (Poppe et al., 2020). The interaction between host immunity and the viral lifecycle underlines the necessity for concurrent management of both viral and immunological factors in affected individuals (Moore & Chang, 2003) (Lidenge et al., 2018).

**Prevalence and Future Considerations:** Despite the known oncogenic potential of KSHV, there are currently no approved vaccines or specific antiviral therapies available. Moreover, serologic testing protocols for KSHV remain non-standardized, complicating clinical management (Tabtieng et al., 2018) (Poppe et al., 2020). Future directions in KSHV research include the development of improved diagnostics, targeted immunotherapeutics, and prophylactic strategies particularly aimed at high-risk populations (Poppe et al., 2020) (Botto et al., 2017).

#### JCV: John Cunningham Virus

**Classification and Transmission Ecology:** The JC virus (JCV) is classified as a small, non-enveloped double-stranded DNA polyomavirus belonging to the family Polyomaviridae. It establishes asymptomatic infections primarily during childhood, with transmission occurring likely through fecal–oral or respiratory routes. The seroprevalence of JCV is notably high, ranging from 50% to 80% globally, which reflects its widespread nature and emphasizes the significance of early-life exposure (Elia et al., 2016) (Hildreth & Alcendor, 2021). Following primary infection, JCV demonstrates a propensity for latent infection, particularly in the renal epithelium and lymphoid tissues, demonstrating a long-term persistence in human hosts (Hildreth & Alcendor, 2021) (Delbue et al., 2013). This characteristic highlights its adaptation to human hosts and limited interspecies tropism (Delbue et al., 2013).

**Clinical Manifestations and Latency:** After its initial asymptomatic infection, JCV establishes latency in various body tissues, notably in the kidneys, bone marrow, and potentially in lymphoid tissues. In individuals who are immunocompromised, such as those living with HIV or undergoing treatments that suppress the immune system, JCV can reactivate, leading to severe neurological manifestations including progressive multifocal leukoencephalopathy (PML) (Bartsch et al., 2019) (Major, 2011). PML is characterized by the lytic infection of oligodendrocytes and astrocytes, resulting in significant demyelination (Major, 2011). In addition to PML, JCV can lead to other neurological conditions, including granule cell neuronopathy, JCV encephalopathy, and meningitis, highlighting its ability to affect diverse neuronal populations (Major, 2011) (Reoma et al., 2019).

**Global Distribution and Seroprevalence:** JCV exhibits a global distribution, with seroprevalence estimates between 50% and 80%, primarily acquired during childhood. Despite the high rates of infection, JCV remains asymptomatic and clinically silent in the vast majority of immunocompetent individuals, which underscores its capacity for long-term persistence in the human population (Hildreth & Alcendor, 2021) (Kolasa et al., 2015). Previous studies have demonstrated that adults often retain antibodies against JCV, indicating prior exposure (Elia et al., 2016) (Kolasa et al., 2015). These findings accentuate the importance of early-life infection in the virus's epidemiology and serological profile globally.

**Risk Factors for Opportunism:** The reactivation of JCV and subsequent development of PML predominantly affects individuals with profound T-cell dysfunction. This includes patients infected with HIV/AIDS, particularly those with CD4+ T-cell counts below 200 cells/µL, and individuals with multiple sclerosis receiving immunomodulatory therapies like natalizumab (Bartsch et al., 2019) (Warnke et al., 2011). Additionally, other risk factors for opportunistic manifestations include transplant recipients on immunosuppressive regimens and individuals with primary immunodeficiencies (Bartsch et al., 2019) (Major, 2011). There is evidence that JCV can lead to disease even in the absence of notable immune deficiencies, emphasizing the complexity of host-viral interactions in different clinical contexts (Hildreth & Alcendor, 2021) (Kolasa et al., 2015).

**Pathogenesis and Persistence Mechanisms:** JCV's ability to persist and reactivate in immunocompromised individuals is intrinsically linked to specific rearrangements in the viral genome's non-coding control region. Such genomic alterations enhance the virus's replication capacity and its tropism for glial cells, which correlates with significant neurological damage (Major, 2011). Additionally, the persistence of JCV in the nervous system may be due to a combination of latency and localized immune evasion mechanisms, allowing for its reactivation under conditions of diminished immune surveillance (Jiang et al., 2010).

**Immune Arm Dependencies:** The immune response to JCV is crucial, with intact CD4+ and CD8+ T-cell functions being essential for controlling the virus. CD8+ T cells play a particularly significant role in the recognition and containment of JCV infection (Jiang et al., 2010). Individuals exhibiting strong JCV-specific T-cell responses typically show a reduced risk for PML development, underscoring the importance of immune surveillance in limiting JCV-associated pathologies (Bartsch et al., 2019) (Jiang et al., 2010).

**Compartmental Tropism and Failure:** While JCV primarily targets white matter in the brain during PML, it also affects gray matter structures, including the cerebellum and cortex. This affinity for gray matter can lead to distinct clinical manifestations, such as ataxia from granule cell neuronopathy and cortical dysfunction in cases of JCV encephalopathy (Major, 2011) (Reoma et al., 2019). The capacity of JCV to invoke neurological conditions beyond PML demonstrates its broad pathogenic potential within the central nervous system (Reoma et al., 2019).

**Immune Modulation and Evasion:** The large T-antigen of JCV is known to disrupt the p53 tumor suppressor pathway, which aids in viral replication and cellular survival by bypassing normal cellular checkpoints. This mechanism of immune evasion is finely tuned to exploit human cellular pathways, highlighting the virus's adaptation to its human host (Bayliss et al., 2011). In contrast, similar mechanisms do not function in murine models, emphasizing the specificity of JCV to human biology (Bayliss et al., 2011).

**Immune Memory and Coordination:** Long-lived immune memory has been documented for JCV-specific CD8+ T cells, which remain detectable years after initial infection, suggesting that these memory responses play a crucial role in mediating immune control and potentially limiting reactivation of latent infection (Jiang et al., 2010) (Bayliss et al., 2011). The restored immune surveillance following periods of immunosuppression may contribute to shaping the immune repertoire over time, indicating the nuanced interplay between JCV and host immune responses (Warnke et al., 2011) (Jiang et al., 2010).

**Role in Immune Architecture:** JCV may function as an immune engager, primarily preserving memory T-cell responses rather than initiating them. The ability to persist in a latent state and reactivate upon immune loss indicates JCV's potential role in safeguarding the immune architecture, particularly in the central nervous system (Bayliss et al., 2011).

**Impact of Immunosuppression:** The emergence of PML is predominantly associated with severe T-cell deficiencies, particularly involving CD4+ and CD8+ T cells. Therapeutics like natalizumab, which target immune system pathways, are significant risk factors for the development of PML (Major, 2011) (Warnke et al., 2011). The diagnosis of PML typically involves the detection of JCV DNA in cerebrospinal fluid (CSF) through PCR, although negative results do not rule out the diagnosis, particularly in patients undergoing antiretroviral therapy due to reduced viral loads (Bartsch et al., 2019) (Major, 2011).

**Prevalence and Future Considerations:** With the growing popularity of biological and immunosuppressive treatments, the risk of JCV reactivation and PML prevalence is on the rise. Genetic variations in immune regulatory genes may predispose individuals to PML under immunosuppressive conditions (Hildreth & Alcendor, 2021) (Lee et al., 2019). Future research directions may include the development of JCV-specific T-cell therapies and enhanced risk stratification tools aimed at restoring immune response without inducing immune reconstitution inflammatory syndrome (IRIS) (Hildreth & Alcendor, 2021) (Lee et al., 2019).

#### BKV: BK Polyomavirus

**Classification and Transmission Ecology:** BK virus (BKV) is classified as a non-enveloped, double-stranded DNA polyomavirus, first identified in 1971. It is a member of the Polyomaviridae family and is adapted to human hosts, persisting for life, primarily in the renal tubular epithelium and uroepithelium (Hirsch & Snydman, 2005) (Hirsch, 2002). Primary infection usually occurs during childhood, often transmitted via respiratory or fecal–oral routes. This initial infection is generally asymptomatic and leads to viral latency, wherein the virus remains dormant for extended periods without causing clinical illness in healthy individuals (Hirsch & Snydman, 2005) (Hirsch, 2002). Studies indicate that seroprevalence of BKV can be alarmingly high, with over 70% of the adult population exhibiting serological evidence of exposure (Hirsch & Snydman, 2005) (Dharnidharka et al., 2011).

**Clinical Manifestations and Latency:** In immunocompetent individuals, BKV typically remains asymptomatic following the primary infection due to effective host immune responses. However, in immunosuppressed populations, particularly kidney transplant recipients, there is a significant risk of BKV reactivation, which can lead to serious complications such as BK virus-associated nephropathy (BKVAN) (Hirsch, 2002) (Dharnidharka et al., 2011). This condition is characterized by progressive tubulointerstitial nephritis and may ultimately contribute to graft failure in an estimated 1-10% of renal transplant patients (Dall & Hariharan, 2008) (Comoli et al., 2006). Similarly, in hematopoietic stem cell transplant (HCT) recipients, reactivation may manifest as hemorrhagic cystitis (Dropulic & Jones, 2007) (Laskin et al., 2013), posing substantial morbidity (Dropulic & Jones, 2007).

**Global Distribution and Seroprevalence:** BKV exhibits a high global seroprevalence, with estimates indicating that 70% to over 90% of adults worldwide demonstrate antibodies against it. This high prevalence suggests widespread asymptomatic infection, with primary exposures typically occurring in early childhood before the age of 10 years (Hirsch & Snydman, 2005) (Alcendor, 2019). Given that most individuals become seropositive without experiencing noticeable symptoms, BKV is often overlooked despite its prevalence as a commensal virus in the human population (Alcendor, 2019).

**Risk Factors for Opportunism:** The reactivation of BKV and subsequent nephropathy is closely associated with various immunosuppression protocols, particularly in renal transplant patients (Dall & Hariharan, 2008) (Kant et al., 2022). Specifically, recipients treated with calcineurin inhibitors such as tacrolimus or cyclosporine, and mycophenolate mofetil are at increased risk (Comoli et al., 2006) (Kant et al., 2022). Other identified risk factors include the use of anti-thymocyte globulin, the presence of HLA mismatches, and ureteral stenting (Dall & Hariharan, 2008) (Kant et al., 2022). In HCT patients, risk factors for developing hemorrhagic cystitis include specific conditioning regimens, notably those involving cyclophosphamide, combined with delayed immune reconstitution (Dropulic & Jones, 2007) (Laskin et al., 2013).

**Pathogenesis and Persistence Mechanisms:** BKV establishes latency within the renal and uroepithelial cells, with reactivation often triggered by immune suppression (Liacini et al., 2010) (Randhawa et al., 2003). During this phase, mutations may occur in the viral noncoding control region, enhancing replication and altering cellular tropism (Randhawa et al., 2003). The viral large T antigen plays a critical role in this process by disrupting tumor suppressor functions among host cells, such as those mediated by p53 and retinoblastoma (pRb) proteins (Liacini et al., 2010) (Costa, 2012). Such interactions promote cell proliferation and facilitate the persistence of the virus, which can lead to significant pathological consequences when immune surveillance is impaired (Costa, 2012) (Lamarche et al., 2016).

**Immune Arm Dependencies:** Effective control of BKV is dependent on innate and adaptive immunity, particularly the response of CD4+ and CD8+ T cells, which correlate significantly with the clearance of the virus (Belhassen-García et al., 2015) (Šťastná-Marková et al., 2021). Patients who manage to maintain robust BK-specific T-cell responses are less likely to progress to viremia or associated nephropathy (Belhassen-García et al., 2015) (Šťastná-Marková et al., 2021). While NK cells can also contribute to immune control of BKV, their specific role remains less clearly defined within the context of T-cell-mediated immunity (Belhassen-García et al., 2015).

**Compartmental Tropism and Failure:** BKV predominantly affects the renal allograft in kidney recipients and the bladder mucosa in HCT recipients (Liacini et al., 2010) (Decker et al., 2009). Outcomes associated with viral reactivation include conditions such as tubulointerstitial nephritis, ureteral stenosis, and hemorrhagic cystitis (Laskin et al., 2013) (Liacini et al., 2010). These manifestations are directly correlated with the extent of immunosuppressive therapy and the effectiveness of immune surveillance, emphasizing the compartment-specific nature of BKV pathology (Dropulic & Jones, 2007) (Liacini et al., 2010).

**Immune Modulation and Evasion:** The large T antigen of BKV disrupts critical host cell cycle checkpoints by inactivating p53 and pRb, effectively preventing apoptosis and allowing for sustained viral infection (Costa, 2012) (Lamarche et al., 2016). Despite not encoding a broad array of immunoevasins like some herpesviruses, BKV relies significantly on its T-antigen for modulating the host's immune response and enhancing viral persistence (Liacini et al., 2010) (Costa, 2012).

**Immune Memory and Coordination:** Due to the establishment of a virus-specific immune response, particularly involving CD4+ and CD8+ T cells, patients can exhibit long-term immune memory post-infection, which may aid in controlling viral replication timely (Dharnidharka et al., 2011) (Šťastná-Marková et al., 2021). Importantly, the presence of BK-specific T cells following transplantation significantly correlates with a reduced risk of viral reactivation (Dharnidharka et al., 2011) (Šťastná-Marková et al., 2021). This underscores the necessity for ongoing immune surveillance in maintaining health post-transplant.

**Role in Immune Architecture:** While BKV does not act as a cornerstone of immune memory, it critically requires coordinated T-cell responses for effective control. The development of persistent BKV-specific T-cell immunity can contribute to immune readiness, particularly within the urogenital tract, underscoring how BKV could influence the structural integrity of the immune environment (Ambalathingal et al., 2017) (Schachtner et al., 2011). Enhancing this immune architecture through tailored immunomodulation strategies could bolster the host's defense against subsequent reactivations of BKV during immunosuppressive therapy, reiterating the importance of a balanced immune response in managing BKV-related diseases (Ambalathingal et al., 2017) (Schachtner et al., 2011).

**Impact of Immunosuppression:** In kidney transplant recipients, BKV reactivation often occurs within the first year, with viruria detected in up to 40% and viremia in 10-20% of patients, leading to BKVAN in about 1-10% (Dall & Hariharan, 2008) (Alcendor, 2019). This condition poses a serious threat, as graft loss attributed to BKV nephropathy can occur in 10-20% of affected cases (Comoli et al., 2006) (Alcendor, 2019). Similarly, in HCT, hemorrhagic cystitis linked to BKV reactivation occurs in approximately 15% of patients, contributing to substantial morbidity and extended hospitalization (Dropulic & Jones, 2007) (Laskin et al., 2013).

**Prevalence and Future Considerations:** BKV continues to be recognized as a leading cause of viral morbidity in renal transplantation. Currently, there are no specific antiviral therapies approved for BKV; however, ongoing research aims to develop virus-specific T-cell therapies alongside immune modulation approaches to safeguard graft function while minimizing the risk of rejection (Belhassen-García et al., 2015) (Lamarche et al., 2016) (Dharnidharka et al., 2011). Future therapeutic strategies will need to strike a balance between enhancing immune surveillance and preventing the complications from transplant rejection.

#### Treponema pallidum (Syphilis)

**Classification and Transmission Ecology:** Treponema pallidum is classified as a spirochete bacterium and is the sole causative agent of syphilis, an infection exclusive to humans (Radolf et al., 2016). It is characterized as a strictly human pathogen; T. pallidum has no known animal reservoir, reinforcing the efficiency of its transmission primarily through sexual contact, including genital, oral, and anal routes (Radolf et al., 2016) (Thurlow et al., 2022). The bacterium can also be transmitted vertically from mother to fetus, leading to congenital syphilis, a significant public health concern due to its severe fetal and neonatal complications (Radolf et al., 2016) (Fadipe et al., 2024).

**Clinical Manifestations and Latency:** Syphilis progresses through four clinically distinct stages, beginning with primary syphilis that presents as a painless chancre, followed by secondary syphilis denoted by systemic rashes and lymphadenopathy (Chan et al., 2016) (Larsen & Larsen, 2020). If not treated, the disease may enter a latent phase characterized by the absence of symptoms, during which T. pallidum persists within host tissues (Thurlow et al., 2022) (Zheng et al., 2025). Reactivation can occur years later, indicating the potential for a prolonged asymptomatic period despite continuous infection (Chan et al., 2016) (Zheng et al., 2025).

**Global Distribution and Seroprevalence:** The prevalence of syphilis is notable worldwide, with endemic conditions reported especially in sub-Saharan Africa and Southeast Asia, as well as among men who have sex with men (MSM) in high-income countries (Thurlow et al., 2022) (Kaminiów et al., 2024). The United States has observed rising syphilis rates over the past decade, particularly among individuals co-infected with HIV, further exacerbating the public health implications of the infection (Chan et al., 2016) (Kaminiów et al., 2024). The World Health Organization reports millions of new cases annually, underscoring the disease's persistent global burden (Kaminiów et al., 2024).

**Risk Factors for Opportunism:** Several risk factors contribute to T. pallidum infections, including high-risk sexual behaviors associated with MSM status, sex work, and the presence of multiple sexual partners (Chan et al., 2016) (Kaminiów et al., 2024). Co-infection with HIV significantly amplifies the risk of syphilis complications, such as neurosyphilis, as the immunocompromised state may hinder the host's ability to control these infections (Kaminiów et al., 2024) (Fadipe et al., 2024). Insufficient prenatal screening also correlates with increased rates of congenital syphilis, presenting further challenges in controlling its spread (Fadipe et al., 2024).

**Pathogenesis and Persistence Mechanisms:** T. pallidum utilizes skin and mucosal penetration strategies to establish infection, followed by hematogenous dissemination, which facilitates widespread organ involvement, including the CNS (Kojima et al., 2022). The bacterium's outer membrane is devoid of numerous exposed proteins, minimizing its visibility to the immune system, and presenting as an “immunological stealth pathogen” (Radolf et al., 2016) (Kojima et al., 2022). Additionally, T. pallidum employs antigenic variation through the TprK gene to evade immune responses, complicating the host's ability to mount an effective defense (Giacani et al., 2012) (Zheng et al., 2025).

**Immune Arm Dependencies:** Both humoral and cell-mediated immune responses are necessary to control T. pallidum. Specifically, CD4+ T cells, particularly those with Th1-type cytokine profiles, play a pivotal role in lesion resolution and disease control (Kojima et al., 2022) (Kaminiów et al., 2024). However, the antibody responses generated are typically non-protective, and prior infection does not confer sterilizing immunity, leaving individuals vulnerable to reinfection (Radolf et al., 2016) (Kojima et al., 2022).

**Compartmental Tropism and Failure:** T. pallidum demonstrates a wide tissue tropism, with a tendency to invade almost all organ systems intermittently, particularly the CNS even in the nascent stages of the disease (Chan et al., 2016) (Houston et al., 2022). Untreated infections may lead to serious long-term consequences, including neurosyphilis and cardiovascular disease, underscoring the bacterium's capacity for chronic persistence and sequelae (Chan et al., 2016) (Kaminiów et al., 2024).

**Immune Modulation and Evasion:** The organism's ability to evade the host immune response is attributed not only to its sparse outer membrane protein expression and antigenic variation but also to its capability to inhibit processes such as phagocytosis Kojima et al., 2022), Yi et al., 2024) (Kojima et al., 2022) (Yi et al., 2024). This immune modulation strategy enables T. pallidum to sustain persistent infections, even in hosts with intact immune systems (Goetting-Minesky et al., 2022) (Yi et al., 2024).

**Immune Memory and Coordination:** While some degree of immune memory develops post-infection, it remains incomplete and does not confer lasting immunity against reinfection. High-risk populations demonstrate a notable frequency of reinfections, indicating a persistent gap in protective immunity following T. pallidum exposure (Radolf et al., 2016) (Kojima et al., 2022).

**Role in Immune Architecture:** The chronic persistence of T. pallidum, particularly in immune-privileged sites like the CNS and placenta, leads to disturbed immune responses. In cases of congenital syphilis, significant disruptions to fetal development and immune balance arise, often resulting in complex systemic manifestations at birth (Radolf et al., 2016) (Goetting-Minesky et al., 2022).

**Impact of Immunosuppression:** Immunosuppression, particularly due to HIV, fundamentally alters the dynamics of syphilis transmission and progression. It can exacerbate symptoms, accelerate the progression to neurosyphilis, and hinder the efficacy of treatment responses (Kaminiów et al., 2024) (Fadipe et al., 2024), further complicating the epidemiological landscape of T. pallidum infections (Kaminiów et al., 2024).

**Prevalence and Future Considerations:** The global resurgence of syphilis, particularly among MSM and HIV-infected individuals, indicates a pressing need for refined public health strategies, including improved vaccine development and enhanced screening methodologies aimed at interrupting perinatal transmission (Radolf et al., 2016) (Chan et al., 2016) (Kojima et al., 2022). Addressing these challenges continues to be a significant public health priority, given the disease's longstanding impact on reproductive health and overall well-being (Kaminiów et al., 2024) (Fadipe et al., 2024).

#### Helicobacter pylori (Helicobacter)

**Classification and Transmission Ecology:** Helicobacter pylori (H. pylori) is classified as a gram-negative, spiral-shaped bacterium that primarily colonizes the gastric mucosa in humans. It demonstrates adaptations that allow it to thrive in the acidic environment of the stomach, a feature invaluable for its survival and pathogenicity (Frydman et al., 2015). The transmission of H. pylori is chiefly maternal or familial, occurring predominantly in early childhood. This bacterium is typically not acquired environmentally during adulthood, and close living conditions contribute significantly to its spread via fecal–oral and oral–oral routes (Parsonnet, 1999). Moreover, epidemiological studies indicate that the bacterium is not easily transmitted through contaminated drinking water, reinforcing the importance of direct person-to-person contact in its transmission (Frydman et al., 2015).

**Clinical Manifestations and Latency:** In the majority of immunocompetent hosts, H. pylori infection is asymptomatic; however, up to 10% of those infected may develop diseases such as chronic gastritis, peptic ulcer disease, gastric adenocarcinoma, or mucosa-associated lymphoid tissue (MALT) lymphoma due to the chronic inflammation induced by the bacterium (Smith et al., 2019). Interestingly, H. pylori establishes persistent infections within the gastric mucosa but lacks classical states of latency. The chronic inflammation and subsequent pathologies align with the infection's prolonged nature, contributing to an increased risk of gastric malignancies (Sgouras et al., 2015).

**Global Distribution and Seroprevalence:** H. pylori affects more than 50% of the global population, showcasing significant disparities in prevalence based on geographic and socioeconomic factors. In low- and middle-income countries, seroprevalence often exceeds 80%, with higher rates linked to poorer sanitation and crowded living conditions (Smith et al., 2019) (Bruce & Maaroos, 2008). Early childhood acquisition remains prevalent in such regions, underscoring the correlation between H. pylori transmission and environmental factors (Bruce & Maaroos, 2008).

**Risk Factors for Opportunism:** The transition from mere colonization to clinical manifestations of H. pylori infection can be exacerbated by several risk factors. High-virulence strains, such as those that express CagA, are associated with greater pathogenic potential (Sgouras et al., 2015). Other relevant factors include smoking and the use of non-steroidal anti-inflammatory drugs (NSAIDs), which can lead to increased mucosal inflammation and a heightened risk of complications like ulcers and malignancies (Sgouras et al., 2015) (Bruce & Maaroos, 2008). Immunocompromised states, while not significantly altering colonization rates, may lead to an increased risk of complications such as MALT lymphoma (Smith et al., 2019) (Sgouras et al., 2015).

**Pathogenesis and Persistence Mechanisms:** H. pylori employs several mechanisms to persist in the gastric mucosa, primarily through its use of urease, which neutralizes gastric acid and facilitates survival in the harsh environment (Fung et al., 2019). Additionally, its flagella confer motility, and adhesins enable the bacterium to adhere tightly to epithelial surfaces, anchoring its colonization efforts (Hidaka, 2001). The virulence factors CagA and VacA actively disrupt host cellular processes, contributing to tissue inflammation, damage, and long-term colonization (Sgouras et al., 2015).

**Immune Arm Dependencies:** The host's immune response to H. pylori infection relies on both innate and adaptive mechanisms. Initial responses involve macrophages and neutrophils, while the adaptive immune response, characterized by CD4+ Th1 and Th17 cells, ultimately occurs (Frydman et al., 2015). Despite these robust immune responses, H. pylori exhibits remarkable evasion capabilities, allowing it to persist without being cleared (Suganuma et al., 2008). This persistence is further complicated in individuals with immunodeficiencies, which may alter the disease phenotype and exacerbate complications (Suganuma et al., 2008) (Zhou et al., 2023).

**Compartmental Tropism and Failure:** The confinement of H. pylori to the gastric mucosa has significant implications for the associated disease processes. Chronic infection results in ongoing inflammation and epithelial injury, leading to atrophy and an enhanced risk of metaplasia and subsequent neoplastic progression (Smith et al., 2019) (Sgouras et al., 2015). The failure of immune regulation in these settings is a key factor in the pathogenesis of gastric cancer (Bruce & Maaroos, 2008).

**Immune Modulation and Evasion:** H. pylori's ability to modulate host immune responses is critical to its survival strategy. Proteins such as VacA inhibit the activation of T cells and alter autophagic processes, while CagA interferes with key signaling pathways in epithelial cells (Suganuma et al., 2008). This immune evasion is complemented by the downregulation of dendritic cell functions, effectively creating a permissive environment for chronic infection (Suganuma et al., 2008) (Frydman et al., 2015).

**Immune Memory and Coordination:** Although H. pylori infection stimulates the formation of memory T and B cells, these responses are often non-protective. The lack of durable immunity against H. pylori facilitates reinfection, highlighting the inadequacies of the adaptive response in recalibrating systemic immune hierarchies (Bruce & Maaroos, 2008) (Frydman et al., 2015). Thus, while some immunological memory is established, it does not confer long-lasting protection.

**Role in Immune Architecture:** Despite not being a classical keystone pathogen, H. pylori holds the potential to shape the local immune environment of the gastric mucosa by inducing lymphoid aggregates and stimulating chronic antigenic responses, which can enhance the risk of lymphoma through persistent inflammation (Suganuma et al., 2008) (Hidaka, 2001). This chronic immune activation underscores the complex interplay between H. pylori, host immune responses, and the development of gastric diseases.

**Impact of Immunosuppression:** Immunosuppressive therapies generally do not increase the overall colonization burden of H. pylori. However, they can promote disease progression to complications like MALT lymphoma (Suganuma et al., 2008) (Hidaka, 2001). Co-infection with HIV does not markedly affect H. pylori prevalence, although it may influence local inflammatory responses, contributing to more severe disease manifestations (Smith et al., 2019) (Hidaka, 2001).

**Prevalence and Future Considerations:** H. pylori remains a globally influential pathogen, with ongoing research needed to address cancer risk stratification and vaccine development. There is also emerging interest in its potential inverse relationship with conditions such as gastroesophageal reflux disease (GERD), which remains an area ripe for further exploration (Bruce & Maaroos, 2008) (Fujimura et al., 2002) (Hidaka, 2001).

### Human Papillomavirus (HPV)

**Classification and Transmission Ecology:** Human Papillomaviruses (HPVs) are classified as small, non-enveloped, double-stranded DNA viruses that belong to the Papillomaviridae family, specifically within the genus Papillomavirus (Frazer, 2010). These viruses exhibit a high degree of species specificity and are primarily human pathogens (Tiggelaar et al., 2012). Currently, more than 200 distinct HPV genotypes have been documented, categorized based on their tissue tropism into cutaneous and mucosal types (Lu et al., 2011). The primary mode of transmission for HPVs is through direct skin-to-skin or mucosal contact, which includes various forms of sexual interaction, such as genital-genital, oral-genital, and anal contact (Scherer et al., 2016) (Vaccarella et al., 2010).

**Clinical Manifestations and Latency:** Most HPV infections are asymptomatic and typically resolve without intervention within a year (Mooij et al., 2013). However, persistent infections with high-risk HPV types—such as HPV 16 and HPV 18—can lead to significant clinical complications, including cervical, anal, and oropharyngeal cancers (Demir et al., 2012). Research indicates that HPV can enter a latent state within basal epithelial cells, with potential reactivation in older adults or immunosuppressed individuals (Elmi et al., 2017).

**Global Distribution and Seroprevalence:** HPV is recognized as one of the most prevalent sexually transmitted infections worldwide, with nearly all sexually active individuals contracting HPV at some point in their lives (Scherpenisse et al., 2012). The prevalence of cervical HPV is particularly high in regions such as sub-Saharan Africa, which report heightened levels of infection compared to other areas (Dauner et al., 2010). Studies indicate seroprevalence varies significantly based on age, sexual orientation, and geographic region, reflecting the complex epidemiology of HPV (Nicoli et al., 2020).

**Risk Factors for Opportunism:** Individuals with compromised immune systems, including those living with HIV, transplant recipients, or patients undergoing chemotherapy, have an increased vulnerability to HPV infections (Ji et al., 2012). Engaging in high-risk sexual behaviors, having multiple sexual partners, starting sexual activity at a young age, and inconsistent condom use are significant factors that elevate the risk of HPV acquisition (Wilson et al., 2013). Data also suggest that male circumcision may correlate with decreased HPV persistence, presenting a potential avenue for risk reduction (Katzenellenbogen et al., 2015).

**Pathogenesis and Persistence Mechanisms:** Oncogenic HPV types can integrate into the host genome, leading to the expression of early viral proteins E6 and E7, which disrupt tumor suppressor pathways, including the p53 and Rb pathways, thereby facilitating malignant transformation (Aghakhani et al., 2016). This integration process often disrupts the E2 gene, which normally inhibits E6/E7 expression, promoting oncogenesis (Castro et al., 2014).

**Immune Arm Dependencies:** The body's ability to clear HPV infections is markedly influenced by cell-mediated immunity. Individuals with HIV or other forms of systemic immunodeficiency demonstrate a reduced capability to clear HPV, associated with an elevated cancer risk. Furthermore, immunocompromised patients generally experience decreased vaccine efficacy for HPV vaccination (Antonsson et al., 2013) (Sun et al., 2023).

**Compartmental Tropism and Failure:** HPVs specifically target stratified squamous epithelia located in the cervix, anus, oropharynx, vulva, vagina, and penis. Research indicates that oropharyngeal sites, particularly the tonsils and the base of the tongue, exhibit heightened susceptibility to HPV-driven carcinogenesis, prominently associated with the high-risk HPV type 16 (Rahman et al., 2016).

**Immune Modulation and Evasion:** HPV employs various mechanisms to evade the host immune response, including downregulating markers of innate immune activation and restricting viral protein expression to differentiated epithelial cells. This strategy, combined with E6-mediated p53 degradation and E7-mediated Rb inactivation, impairs apoptosis and immune-mediated cell clearance, facilitating persistent infections (Michael et al., 2010) (Kim et al., 2012).

**Immune Memory and Coordination:** Natural HPV infections often induce type-specific immunity; however, this immunity may not always be long-lasting (Lewis et al., 2018). Vaccination strategies have demonstrated robust and durable antibody responses across multiple HPV types, significantly enhancing immune memory (Kricker et al., 2020). The protective effects of vaccination may also extend to herd immunity, providing broader public health benefits (Ortíz et al., 2018).

**Role in Immune Architecture:** In contrast to herpesviruses, which engage in systemic immune coordination, HPV primarily exploits localized immune responses within epithelial layers. Observations of reactivation in older adults and immunosuppressed individuals indicate that HPV utilizes tissue-targeted persistence without establishing systemic immune imprints (Ji et al., 2012) (Wilson et al., 2013).

**Impact of Immunosuppression:** Individuals with HIV, organ transplant recipients, and others with immunosuppression have a higher incidence of HPV-related cancers) (Mooij et al., 2013). Specifically, a low CD4 count (<200) has been significantly associated with persistent anal and oropharyngeal HPV infections, subsequently elevating the risk of malignancy (Antonsson et al., 2013).

**Prevalence and Future Considerations:** Vaccination efforts against HPV, especially in regions with high coverage, have led to substantial reductions in the prevalence of HPV types 6, 11, 16, and 18, along with associated diseases. For instance, in the United States, type-specific HPV prevalence has significantly declined among vaccinated individuals aged 14-24 years (Sun et al., 2023). Ongoing clinical trials further support the durability and cost-effectiveness of both two- and three-dose vaccine regimens, highlighting the necessity for expanding vaccination efforts and promoting earlier administration to enhance global control of HPV (Scherpenisse et al., 2012) (Ortíz et al., 2018).

#### Strongyloides stercoralis (Strongyloides)

**Classification and Transmission Ecology:** Strongyloides stercoralis is a soil-transmitted helminth belonging to the family Strongyloididae that displays a unique autoinfective lifecycle enabling persistence within hosts without the need for external reinfection. Transmission typically occurs through skin contact with contaminated soil, wherein the infective larvae penetrate the skin, migrate through the bloodstream to the lungs, and are eventually swallowed to lodge in the small intestine (Breloer & Abraham, 2016) (Ye et al., 2022). This lifecycle allows the parasite to endure within a host for decades, contributing to chronic infections that often go unnoticed until significant immune suppression occurs (Potter et al., 2003) (Brown et al., 2006).

**Clinical Manifestations and Latency:** In immunocompetent individuals, strongyloidiasis may present as predominantly asymptomatic, though some experience intermittent gastrointestinal issues such as abdominal discomfort or diarrhea. Pulmonary manifestations may include dry cough and wheezing (Rose et al., 2009) (Liu et al., 2009). In stark contrast, individuals with impaired cellular immunity—such as those on corticosteroids or undergoing immunosuppression post-transplant—are at high risk for severe manifestations, including hyperinfection syndrome and disseminated strongyloidiasis. These complications can lead to critical conditions such as sepsis, pneumonia, and even meningitis (Kumar et al., 2020) (Guérin et al., 2021).

**Global Distribution and Seroprevalence:** Strongyloides is endemic to tropical and subtropical regions, with seroprevalence rates reported as high as 50% in certain areas. In high-income countries, cases primarily occur among migrants and individuals with past exposure to endemic regions (Norberg et al., 2023) (Banicioiu et al., 2023). This global distribution emphasizes the necessity for increased awareness and potential screening, particularly for at-risk populations undergoing immunosuppressive treatments (Ye et al., 2022) (Liu et al., 2009).

**Risk Factors for Opportunism:** Significant risk factors for severe disease include chronic corticosteroid use, hematologic malignancies, solid organ transplantation, TNF-alpha inhibitor therapy, and co-infection with HTLV-1 (Carvalho & Porto, 2004) (Banicioiu et al., 2023) (Khaliq et al., 2018). While co-infection with HIV is associated with immunosuppression, it does not appear to carry the same heightened risk for hyperinfection compared to these other conditions (Potter et al., 2003) (Guérin et al., 2021).

**Pathogenesis and Persistence Mechanisms:** The ability of Strongyloides stercoralis to complete its lifecycle entirely within humans is crucial for its persistence, enabling the infective larvae to reinfect the intestinal mucosa or perianal skin directly (Breloer & Abraham, 2016) (Kim et al., 2010). Immunosuppressive states, particularly those compromising Th2-mediated immunity, facilitate unchecked replication and systemic spread of the larvae, exploiting the host's impaired immune response (Guérin et al., 2021) (Nnaoma et al., 2019).

**Immune Arm Dependencies:** Control of Strongyloides is chiefly mediated through immune mechanisms centered around Th2-type responses, eosinophils, and macrophages. Eosinophilia driven by IL-5 is characteristic during the early stages of infection but may diminish in chronic cases. While CD4+ T cells play a vital role, the adaptive immune response tends to be localized rather than systemic, increasing the risk of dissemination when both innate and mucosal defenses are compromised (Kumar et al., 2020) (Potter et al., 2003).

**Compartmental Tropism and Failure:** Despite initial localized infections in the gastrointestinal tract, hyperinfection can lead to hematogenous dissemination to the lungs and central nervous system, indicating a broader clinical failure as immunity falters (Nnaoma et al., 2019) (Kirkham et al., 2021). This widespread compromise can allow enteric bacteria to translocate, potentially resulting in severe secondary infections (Rose et al., 2009) (Guérin et al., 2021).

**Immune Modulation and Evasion:** Strongyloides employs various strategies for immune evasion, including secretion of protease inhibitors and modulation of the host's cytokine profiles. The larval stage is particularly skilled at evading neutrophil responses, complicating the diagnosis of hyperinfection, which is often characterized by absent eosinophilia (Hays et al., 2015) (Rose et al., 2009).

**Immune Memory and Coordination:** Unlike many viral infections, Strongyloides stercoralis does not foster enduring systemic adaptive memory. However, its persistence within mucosal

niches—particularly in chronic autoinfection—may contribute to ongoing innate immune training via elevated localized Th2 cytokine responses (Potter et al., 2003) (Khaliq et al., 2018).
**Role in Immune Architecture:** Positioning Strongyloides uniquely in host-parasite interactions, its lifecycle strategy does not invoke continuous systemic immune imprinting as seen with persistent viruses. Nonetheless, it reinforces local immune responses at epithelial surfaces, potentially shaping immune architecture at these critical barriers (Guérin et al., 2021) (Kirkham et al., 2021).
**Impact of Immunosuppression:** Corticosteroids stand out as a primary risk factor for hyperinfection, as they impair eosinophil function, suppress Th2 cytokines, and accelerate the natural autoinfective cycle (Shorman & Al-Tawfiq, 2009) (Banicioiu et al., 2023). The rapid onset of severe outcomes such as multi-organ failure and sepsis in untreated hyperinfection cases can exceed 80% mortality rates (Kumar et al., 2020) (Rose et al., 2009). Evidence regarding the risks associated with newer biologics, such as TNF inhibitors, is still emerging (Khaliq et al., 2018).
**Prevalence and Future Considerations:** Despite its prevalence, Strongyloides stercoralis infections remain largely underdiagnosed, especially in immunosuppressed patients from endemic regions. There is an urgent need to develop comprehensive screening protocols before commencing immunosuppressive therapies. Future research should focus on elucidating mucosal immune thresholds that could prevent the progression to severe disease states and consider the safety of immune checkpoint modulation (Shorman & Al-Tawfiq, 2009) (Liu et al., 2009).

#### Leishmania species (Leishmania)

**Classification and Transmission Ecology:** Leishmania are obligate intracellular protozoan parasites responsible for leishmaniasis, transmitted by infected female sandflies of the Phlebotomus genus in the Old World and Lutzomyia in the New World. This genus includes several species that dominate various forms of human disease, classified primarily as promastigotes in the vector (sandflies) and amastigotes in the mammalian host such as humans or animals (Smith et al., 2007) (Kaye et al., 2020). The diversity of species leads to varied clinical manifestations, ranging from cutaneous lesions to more severe forms such as visceral leishmaniasis, which can be lethal without intervention (Kaye et al., 2020) (Pissarra et al., 2021).
**Clinical Manifestations and Latency:** Clinical manifestations of leishmaniasis are influenced by the infecting species and the host's immune response. Cutaneous leishmaniasis presents as localized ulcerative lesions, while mucocutaneous leishmaniasis is characterized by destructive mucosal damage affecting the oral and nasal cavities (Kaye et al., 2020) (Pissarra et al., 2021). Visceral leishmaniasis, or kala-azar, is indicated by symptoms such as prolonged fever, hepatosplenomegaly, and pancytopenia, and is often linked with latent infections that can reactivate under conditions of immunosuppression (Kaye et al., 2020) (Freitas et al., 2016). The ability of Leishmania to persist latently poses significant challenges in clinical management, particularly among immunocompromised individuals (Freitas et al., 2016) (Okwor & Uzonna, 2013).
**Global Distribution and Seroprevalence:** Leishmaniasis is endemic in more than 90 countries, with cutaneous forms being prevalent in regions such as the Middle East and North Africa, while visceral leishmaniasis is concentrated in South Asia, East Africa, and Brazil (Pissarra et al., 2021)

(Okwor & Uzonna, 2013). The World Health Organization estimates that there are between 700,000 to 1 million new cases of cutaneous leishmaniasis and between 50,000 to 90,000 new cases of visceral leishmaniasis annually, highlighting its global health significance as a neglected tropical disease (Pissarra et al., 2021) (Okwor & Uzonna, 2013).

**Risk Factors for Opportunism:** Risk factors for leishmaniasis include malnutrition, HIV infection, corticosteroid usage, and general immunosuppression. Specifically, co-infection with HIV substantially increases the risk of severe visceral leishmaniasis and can contribute to disease relapse (Okwor & Uzonna, 2013) (Olivier et al., 2003). Environmental factors and poor socioeconomic conditions further exacerbate transmission risks, making high-risk populations more vulnerable to the severe consequences of the disease (Kaye et al., 2020) (Valigurová & Kolářová, 2023).

**Pathogenesis and Persistence Mechanisms:** Following sandfly transmission, Leishmania promastigotes enter the host's macrophages and transform into amastigotes, a critical phase that enables their replication within the harsh phagolysosomal environment. These parasites possess mechanisms to inhibit phagosome maturation, evade oxidative stress, and suppress effective antigen presentation (Freitas et al., 2016) (Valigurová & Kolářová, 2023) (Santos & Brodskyn, 2014). The ability of Leishmania to persist in host macrophages without being eliminated is a key factor in its pathogenicity and the chronic nature of leishmaniasis (Freitas et al., 2016) (Santos & Brodskyn, 2014).

**Immune Arm Dependencies:** The host's immune response plays a crucial role in controlling leishmaniasis. The development of effective Th1-mediated immunity, characterized by the production of interferon-gamma (IFN-γ) and interleukin-12 (IL-12), is essential for activating macrophages to eliminate intracellular amastigotes (Okwor & Uzonna, 2013) (Donovan et al., 2009). Conversely, a Th2-biased response can lead to disease persistence and progression, emphasizing the delicate balance of the immune system in managing Leishmania infections (Freitas et al., 2016; Okwor & Uzonna, 2013) (Freitas et al., 2016) (Okwor & Uzonna, 2013).

**Compartmental Tropism and Failure:** Leishmania exhibits compartmental tropism, primarily targeting the spleen, liver, and bone marrow in cases of visceral leishmaniasis, while cutaneous and mucocutaneous diseases affect the skin and mucosal tissues (Colmenares, Kar, et al., 2002) (Kaye et al., 2020) (Freitas et al., 2016). The pathology in these organs can result in severe immune dysregulation, leading to further complications. The immune failure in these compartments contributes significantly to the ulceration in cutaneous leishmaniasis and systemic disease manifestations in visceral forms (Okwor & Uzonna, 2013) (Olivier et al., 2003) (Valigurová & Kolářová, 2023).

**Immune Modulation and Evasion:** Leishmania has developed intricate strategies to evade host immune responses, including the suppression of IL-12 and the induction of IL-10 production. This immune modulation assists in creating a favorable environment for the parasite's survival and replication within macrophages, ultimately leading to chronic infection (Santos & Brodskyn, 2014) (Pessenda & Silva, 2020). By manipulating macrophage signaling pathways, Leishmania effectively skirt the immune system's elimination strategies (Freitas et al., 2016) (Santos & Brodskyn, 2014).

**Immune Memory and Coordination:** The establishment of memory T-cell responses is crucial following leishmaniasis infection; however, these responses often fail to confer lasting immunity, particularly in endemic areas where transmission is ongoing (Pissarra et al., 2021)

(Okwor & Uzonna, 2013). In individuals with visceral leishmaniasis, impaired cellular immunity can lead to frequent relapses, especially under immunocompromised conditions (Santos & Brodskyn, 2014) (Valigurová & Kolářová, 2023).

**Role in Immune Architecture:** Chronic leishmaniasis can significantly reshape local immune responses, driving granulomatous inflammation or fibrotic remodeling within tissues. In visceral leishmaniasis, the architectural disruption within the spleen and liver contributes to a persistent state of immunosuppression, making patients more vulnerable to secondary infections (Pissarra et al., 2021) (Santos & Brodskyn, 2014) (Moradimotlagh et al., 2023).

**Impact of Immunosuppression:** The intersection of immunosuppression and leishmaniasis, particularly through HIV co-infection, results in aggravated disease manifestations and poor treatment outcomes. Patients with neutropenia or those undergoing immunosuppressive therapies (e.g., organ transplant recipients) are particularly at risk for severe leishmaniasis complications and reactivation of latent infections (Olivier et al., 2003) (Moradimotlagh et al., 2023) (Colmenares, Constant, et al., 2002).

**Prevalence and Future Considerations:** Despite advances in understanding leishmaniasis, it remains a neglected tropical disease with significant challenges such as drug resistance and the persistence of the disease in immunocompromised individuals. Efforts are being directed toward vaccine development, vector control strategies, and novel therapies aimed at enhancing the host's immune responses to effectively combat this complex parasitic infection (Pissarra et al., 2021) (Santos & Brodskyn, 2014) (Donovan et al., 2009) (Pessenda & Silva, 2020).

#### Neisseria meningitidis

**Classification and Transmission Ecology:** Neisseria meningitidis, a significant human pathogen, is classified as a gram-negative diplococcus. It predominantly colonizes the nasopharyngeal epithelium, making it an obligate human pathogen associated with respiratory transmission methods, such as respiratory droplets and direct contact with respiratory secretions. Asymptomatic carrier states are particularly prevalent among adolescents and young adults, who serve as the primary reservoirs facilitating community spread of the pathogen (Parikh et al., 2020) (Siena et al., 2018). For instance, the carriage prevalence has been reported to be 8-25% globally in the nasopharynx, with peak rates observed in adolescents, where up to 23.7% may carry the organism (Kristinsdóttir et al., 2023). This aspect underscores the opportunistic nature of N. meningitidis and its capacity for transmission within crowded living conditions (Parikh et al., 2020).

**Clinical Manifestations and Latency:** Infections by Neisseria meningitidis can lead to severe invasive disease characterized by acute onset and rapid progression. The primary clinical manifestations include meningitis, meningococcemia, purpura fulminans, and septic shock. While N. meningitidis can persist asymptomatically in the nasopharynx, its transition to invasive disease is marked by an accelerated and acute clinical presentation (Jafri et al., 2013). The pathogen's quick progression highlights the urgent need for timely diagnosis and intervention to mitigate morbidity and mortality associated with invasive meningococcal disease (Jafri et al., 2013).

**Global Distribution and Seroprevalence:** N. meningitidis displays considerable serogroup diversity, with serogroups A, B, C, W, and Y accounting for nearly all reported cases of meningococcal disease worldwide. Notably, the incidence is particularly high in the African meningitis belt, where historical outbreaks of serogroup A have been frequent, although ongoing vaccination programs have significantly reduced its prevalence (Mwenda et al., 2019). Carriage rates are highest in adolescents and young adults, emphasizing the public health significance of this age group in the epidemiology of the disease. Furthermore, environmental factors, such as crowded settings, enhance the likelihood of transmission and disease outbreaks (Kristinsdóttir et al., 2023).

**Risk Factors for Opportunism:** Several risk factors contribute to the susceptibility to invasive N. meningitidis disease. Individuals with complement deficiencies, especially affecting terminal components such as C5–C9, as well as asplenic patients or those living with HIV, are at markedly increased risk (Jafri et al., 2013). Furthermore, environmental factors, such as close contact with infected individuals, exposure to cigarette smoke, and concurrent viral upper respiratory infections, further elevate the likelihood of both carriage and progression to invasive disease (Jafri et al., 2013) (Parikh et al., 2020).

**Pathogenesis and Persistence Mechanisms:** N. meningitidis employs specific virulence factors for adhesion and invasion, including type IV pili and opacity-associated proteins, facilitating its interaction with epithelial cells. The polysaccharide capsule is crucial for evasion of the host immune response, inhibiting phagocytosis and complement-mediated opsonization. Additionally, the pathogen's lipooligosaccharide (LOS) is instrumental in immune modulation, contributing to cytokine release and endothelial cell injury, thus facilitating invasive pathogenesis (Jafri et al., 2013).

**Immune Arm Dependencies:** The complement system plays a pivotal role in host defense against N. meningitidis bacteremia, with complement-mediated lysis being a primary defense mechanism. Individuals with complement deficiencies or those undergoing complement inhibition therapies (e.g., eculizumab) are at significantly elevated risk of invasive disease despite vaccination (Jafri et al., 2013). The immune response to capsular polysaccharides and surface proteins is critical for effective protection, underscoring the basis for existing vaccines (Jafri et al., 2013).

**Compartmental Tropism and Failure:** The primary locus of N. meningitidis colonization is the nasopharynx. Upon translocating to the bloodstream during invasive disease episodes, the organism can breach the blood-brain barrier, leading to severe complications such as meningitis. The subsequent endothelial injury from systemic infection can progress to purpura fulminans and multiorgan failure (Jafri et al., 2013).

**Immune Modulation and Evasion:** The polysaccharide capsule represents a key virulence determinant that protects N. meningitidis from phagocytosis and complement-mediated clearance. Mechanisms such as phase variation in surface protein expression and phenotypic mimicry—where elements of the serogroup B capsule resemble human neural cell adhesion molecules—facilitate immune evasion (Jafri et al., 2013).

**Immune Memory and Coordination:** Antibody-mediated immune memory directed against specific serogroups is critical for maintaining effective immunity against N. meningitidis. While natural exposure and vaccinations successfully stimulate protective immune responses, the necessity for booster doses arises from waning immunity in younger populations.

**Role in Immune Architecture:** While N. meningitidis does not conform to the definitions of a keystone pathogen, its capacity to exploit complement pathway deficiencies and navigate mucosal barriers illustrates a highly specialized adaptation to human immune niches (Jafri et al., 2013). Its interaction with the complement system is a central aspect of its immunopathogenesis.

**Impact of Immunosuppression:** Immunosuppressed patients, particularly those receiving eculizumab and individuals with HIV, exhibit a dramatically heightened risk of invasive meningococcal disease, revealing the significant intersection of immune status and disease susceptibility (Jafri et al., 2013).

**Prevalence and Future Considerations:** Global efforts to reduce the incidence of invasive meningococcal disease through vaccination strategies, particularly with conjugate vaccines targeting serogroups A, C, W, and Y, have proven effective; however, serogroup B remains a challenging area due to its complex antigenic structure. Continued research is essential to develop innovative, broad-spectrum vaccines aimed at enhancing mucosal immunity and preventing widespread incidence.

#### Torque Teno Virus (TTV)

**Classification and Transmission Ecology:** Torque Teno Virus (TTV) is classified as a small, non-enveloped, single-stranded circular DNA virus approximately 3.8 kb in size. It falls within the genus Alfatorquevirus of the Anelloviridae family, first described in the 1990s (Okamoto, 2009). TTV is notably a human-adapted virus, found extensively in various human tissues and bodily fluids including blood, saliva, stool, urine, and nasal secretions, with no confirmed zoonotic reservoirs (Maggi & Bendinelli, 2010). Transmission of TTV is thought to occur through multiple routes, including parenteral (e.g., blood transfusion), vertical transmission from mother to child, and potentially respiratory or fecal-oral routes (Maggi & Bendinelli, 2010) (Tyschik et al., 2017). Studies have shown TTV's widespread presence in the population, emphasizing its ubiquity, with high prevalence rates observed across different demographics (Maggi & Bendinelli, 2010).

**Clinical Manifestations and Latency:** In immunocompetent individuals, TTV generally does not produce specific or consistent clinical symptoms. Initial hypotheses suggested a relationship between TTV and hepatitis, but substantial research has refuted this link (Koohi et al., 2012) (Blois et al., 2014). Longitudinal studies have indicated that TTV serves predominantly as a commensal organism within the human virome, characteristic of lifelong persistence and limited immunopathological effects (Giménez et al., 2020). Thus, TTV has been repositioned as a virus with low pathogenicity, primarily existing alongside other microbial inhabitants of the human body without contributing significantly to disease processes (Giménez et al., 2020).

**Global Distribution and Seroprevalence:** Prevalence studies indicate that TTV is nearly universal, with seroprevalence exceeding 80%-90% among healthy adults globally (Maggi & Bendinelli, 2010) (Koohi et al., 2012). Research highlights that TTV infection typically occurs early in life, evidenced by the detection of TTV DNA in infants and toddlers. This early acquisition underscores the virus's prevalence within human populations and lends to the notion of its role as an endemic component of the human virome (Mazzola et al., 2015) (Ikuta et al., 2019).

**Risk Factors for Opportunism:** While TTV is largely non-pathogenic, significant viral loads have been documented in immunosuppressed patients, such as organ and stem cell transplant recipients, indicating potential opportunistic behavior in compromised immune systems (Tian et al., 2020). Elevated levels of TTV viremia have been proposed as a marker for immune dysfunction, suggesting a role for TTV replication as an indicator of the immunocompromised state (Tian et al., 2020) (Tyschik et al., 2017) and reinforcing the idea that while TTV may not directly cause disease, it can reflect immune status.

**Pathogenesis and Persistence Mechanisms:** TTV replication is hypothesized to utilize a rolling-circle mechanism, with the open reading frame 1 (ORF1) believed to encode a capsid-like protein (Okamoto, 2009). The virus demonstrates remarkable genetic diversity, which contributes to its persistence in the human host, with lymphocytes and other hematopoietic cells likely serving as primary replication sites (Giménez et al., 2020). This viral diversity complicates detection and treatment modalities, as various strains can engage differently with the immune system.

**Immune Arm Dependencies:** The immune response to TTV involves the production of antibodies, specifically IgM and IgG. However, these antibodies are not effective in clearing the infection, and their roles in immune dynamics remain poorly understood (Spandole-Dinu et al., 2013). Suppression of T lymphocytes, especially via chemotherapy or immunosuppressive treatments, has been linked to increased TTV viremia, thereby highlighting the critical dependence on cellular immunity in the control of the virus (Tian et al., 2020) (Tyschik et al., 2017).

**Compartmental Tropism and Failure:** TTV is detectable in nearly all human tissues and body fluids except for cerebrospinal fluid, indicating a wide tissue tropism (Giménez et al., 2020). High levels of TTV have been associated with various conditions, although the correlation with specific respiratory diseases like asthma requires further evidence (Spandole-Dinu et al., 2013). These associations suggest a complex interplay between viral presence and health outcomes, warranting further investigation into the viral impacts on respiratory diseases.

**Immune Modulation and Evasion:** Recent studies have uncovered complex interactions between TTV and the host immune system. Notably, TTV has been shown to encode several peptides that stabilize HLA-E, fostering the expansion of regulatory immune cells, including NKG2A-expressing CD8+ T cells and NK cells (Novosel et al., 2012). This immune modulation may represent TTV's strategy for persistence, where T cell responses are regulated rather than fully activated, leading to a phenomenon of immune exhaustion that could facilitate the chronic presence of the virus while preventing tissue damage (Spandole-Dinu et al., 2013) (Novosel et al., 2012).

**Immune Memory and Coordination:** Despite the presence of anti-TTV IgG that can last for years, these responses do not confer sterilizing immunity, allowing for potential reinfections by different TTV genotypes (Spandole-Dinu et al., 2013) (Novosel et al., 2012). TTV does not seem to evoke strong epitope hierarchies similar to more immunogenic viruses, reinforcing its role as a non-disruptive component of immune architecture rather than a central player in orchestrating immune responses against other pathogens (Tian et al., 2020).

**Role in Immune Architecture:** TTV's high prevalence and its reactions to immune suppression render it a potential indicator of immune status rather than a coordinator of immune hierarchies or tolerance systems (Tian et al., 2020) (Tyschik et al., 2017). It could serve practical applications in monitoring immune competence and evaluating the risks of infections and complications following immunosuppressive therapies (Tian et al., 2020).

**Impact of Immunosuppression:** Studies demonstrate that TTV replication intensifies in response to therapies targeting T cell populations, indicating significant implications for monitoring viral load in immunocompromised patients, particularly in transplant settings (Tian et al., 2020)

(Tyschik et al., 2017). This monitoring could aid in assessing the risk of opportunistic infections, providing a non-invasive metric for immune status (Tian et al., 2020) (Tyschik et al., 2017).

**Prevalence and Future Considerations:** Despite being widely recognized, TTV remains under-characterized in the scientific literature, lacking commercial testing platforms (Maggi & Bendinelli, 2010) (Spandole-Dinu et al., 2013). Future research is crucial to better understand the immune mechanisms controlling TTV infection, dynamics of co-infection with other viruses, and its potential role as a biomarker for immune suppression (Spandole-Dinu et al., 2013).

#### Cryptococcus neoformans (Cryptococcus)

**Classification and Transmission Ecology:** C. neoformans is classified as an encapsulated, yeast-like fungus belonging to the Basidiomycota phylum, recognized for its production of a thick polysaccharide capsule that serves as a crucial virulence factor (Lin & Heitman, 2006) (Kantardjiev et al., 2006). This pathogen is primarily acquired from environmental sources, such as soil and avian excreta, particularly pigeon droppings, which facilitate its aerosolization and subsequent inhalation by humans and animals (Nielsen et al., 2007) (Esakkiammal & Rajakumari, 2025). Humans serve as incidental hosts and do not transmit the infection to others, indicating a unique ecological niche where C. neoformans is linked mainly to environmental reservoirs rather than direct human-to-human transmission (Boekhout et al., 2001).

**Clinical Manifestations and Latency:** Infections by C. neoformans often localize in the lungs, leading to asymptomatic cases or mild respiratory symptoms that may resolve spontaneously (Lin & Heitman, 2006). However, in individuals with compromised immune systems, particularly those with advanced HIV infection, the fungus can reactivate from latent pulmonary foci, allowing for dissemination to the central nervous system (CNS), which can precipitate severe conditions like cryptococcal meningitis (Baddley et al., 2011) (Zhang et al., 2022). This progression from lung infection to CNS involvement underscores the critical importance of immune status in disease latency and reactivation (Vanhove et al., 2016).

**Global Distribution and Seroprevalence:** Globally, C. neoformans is ubiquitous, with higher prevalence rates observed in tropical and subtropical regions, especially in areas heavily affected by HIV/AIDS (Fortes et al., 2001) (Guillot et al., 2008). Sub-Saharan Africa is particularly burdened by cryptococcal disease, significantly contributing to mortality in late-stage AIDS patients, often ranking as a leading cause of HIV-related deaths following tuberculosis (Mseddi et al., 2011) (Pedroso et al., 2009). In resource-limited settings, inadequate diagnostics and care amplify the risk of cryptococcal meningitis, necessitating global health initiatives aimed at improving access to antifungal therapies and diagnostic tools like the Cryptococcal Antigen (CrAg) lateral flow assay ((Esakkiammal & Rajakumari, 2025) (Yang et al., 2021).

**Risk Factors for Opportunism:** The primary risk factors for C. neoformans infection include profound immunosuppression, particularly in individuals with advanced HIV/AIDS (CD4 <100 cells/μL), as well as those undergoing solid organ transplantation or immunosuppressive therapy (Fortes et al., 2001) (Mseddi et al., 2011). Conditions such as hematologic malignancies and the presence of anti-GM-CSF autoantibodies further predispose individuals to infection, emphasizing the critical interaction between host immune status and the opportunistic nature of this pathogen (Mak et al., 2015) (Kwon-Chung & Varma, 2006).

**Pathogenesis and Persistence Mechanisms:** Once inhaled, C. neoformans establishes a niche in the pulmonary alveoli, where it can remain dormant for extended periods (Lin & Heitman, 2006) (Fortes et al., 2001). The virulence of this pathogen is linked to its polysaccharide capsule, which inhibits phagocytosis, production of melanin to resist oxidative stress, and urease activity that alters the local microenvironment (Zhang et al., 2022) (Hamasha et al., 2004). Additionally, C. neoformans can survive and replicate within host macrophages, potentially facilitating its dissemination into the CNS (Koh et al., 2002).

**Immune Arm Dependencies:** The immune response against C. neoformans is heavily reliant on T-cell mediated mechanisms, particularly Th1 responses characterized by the action of cytokines such as interferon-gamma (IFN-γ) and tumor necrosis factor-alpha (TNF-α) (Fortes et al., 2001) (Guillot et al., 2008). These cytokines promote macrophage activation, which is essential for pathogen clearance. While antibodies can contribute to opsonization, they are generally insufficient by themselves to eradicate the infection in immunocompromised individuals, revealing a gap in protective immunity that may lead to severe disease (Mseddi et al., 2011) (Pedroso et al., 2009).

**Compartmental Tropism and Failure:** C. neoformans displays a particular tropism for the CNS, where failure of cellular immunity can lead to significant complications such as meningoencephalitis, characterized by increased intracranial pressure and neurological impairment (Mak et al., 2015) (Kwon-Chung & Varma, 2006). The initial pulmonary disease often remains undetected, allowing for progressive and severe forms of the disease, particularly in immunocompromised individuals (Koh et al., 2002).

**Immune Modulation and Evasion:** The intricate immune evasion strategies utilized by C. neoformans include the suppression of inflammatory cytokines through the shedding of capsular components like glucuronoxylomannan (GXM), which can impair the host's immune response (Vanhove et al., 2016) (Fortes et al., 2001) (Guillot et al., 2008). The organism's ability to persist within macrophages facilitates its survival and complicates treatment efforts in affected individuals (Zhang et al., 2022) (Hamasha et al., 2004).

**Immune Memory and Coordination:** Immunity to C. neoformans is often insufficiently protective, particularly in individuals with dysfunctional T-cell responses, leading to occurrences such as Immune Reconstitution Inflammatory Syndrome (IRIS) in those undergoing antiretroviral therapy (Fortes et al., 2001) (Guillot et al., 2008). This disordered immune response reflects challenges in maintaining effective long-term immunity against cryptococcosis (Pedroso et al., 2009) (Kwon-Chung & Varma, 2006).

**Role in Immune Architecture:** C. neoformans modifies local immune architecture, particularly within the lungs and CNS, by disrupting macrophage polarization and antigen processing (Fortes et al., 2001) (Castro et al., 2006). In chronic infections, it can reshape granulomatous responses, altering the host's ability to mount effective immune defenses against subsequent infections (Koh et al., 2002) (Kantardjiev et al., 2006).

**Impact of Immunosuppression:** Immunosuppressive conditions, especially those involving T-cell depletion, are primary factors driving disease progression and poor outcomes in C. neoformans infections. Untreated cases exhibit high mortality rates, particularly due to cryptococcal meningitis, indicating an urgent need for prompt therapeutic interventions (Mseddi et al., 2011) (Pedroso et al., 2009) (Yang et al., 2021).

**Prevalence and Future Considerations:** Despite the availability of antifungal treatment, cryptococcal meningitis remains a leading cause of AIDS-related mortality, especially in low-resource settings. Greater access to screening, preemptive therapy, and deeper immunological understanding are necessary to curtail the global burden of this fungal pathogen (Esakkiammal & Rajakumari, 2025) (Yang et al., 2021). Ongoing research focused on improving treatment modalities and preventive strategies remains critical in decreasing the prevalence of C. neoformans-related diseases.

#### Histo: Histoplasma capsulatum

**Classification and Transmission Ecology:** Histoplasma capsulatum is a thermally dimorphic fungus, existing in a mold form in soil and transforming into a yeast when inhaled into the human pulmonary system (Wheat, 1997). This fungus is primarily associated with environments enriched in organic matter, notably areas populated by birds and bats, as their droppings provide a nutrient source (Lyon et al., 2004) (Perio et al., 2021). Transmission typically occurs through inhalation of aerosolized microconidia during activities that disturb the soil, such as construction work, demolition, and exploration of caves (McKinsey & McKinsey, 2011) (Diaz, 2018). Individuals, especially those who explore caves or engage in occupations that expose them to affected soil, face increased risks of contracting histoplasmosis from environments where H. capsulatum thrives (Hajjeh, 1995) (Perio et al., 2021).

**Clinical Manifestations and Latency:** Histoplasmosis can present a wide spectrum of clinical manifestations, ranging from asymptomatic illness to life-threatening disseminated disease, particularly prevalent in immunocompromised patients (McKinsey et al., 1997) (McKinsey, 2021). In its acute form, pulmonary histoplasmosis may manifest with symptoms such as fever, cough, and chest pain, while chronic infections may mimic tuberculosis, leading to apical lung fibrous lesions and cavitation (Wheat, 1997) (Niknam et al., 2017). Disseminated histoplasmosis can affect multiple organs, particularly impacting the liver, spleen, bone marrow, and central nervous system (CNS) in severely immunocompromised individuals (McKinsey et al., 1997) (Hajjeh, 1995).

**Global Distribution and Seroprevalence:** H. capsulatum has defined endemicity, particularly in the Ohio and Mississippi River valleys of the United States, along with certain regions of Central and South America, parts of Africa, and Southeast Asia (Park et al., 2020). The emergence of cases from non-endemic regions has increased, largely attributed to travel-related exposure and donor-derived infections among organ transplant recipients (Wheat, 1982) (Pulgarin et al., 2021). This underscores the need for improved awareness and surveillance in both endemic and non-endemic areas regarding histoplasmosis (Miller et al., 2022).

**Risk Factors for Opportunism:** Immunocompromised states significantly heighten the risk of developing disseminated forms of histoplasmosis. Conditions such as AIDS, characterized by CD4+ T-cell deficiency (with counts below 150 cells/µL), the use of immunosuppressive medications including TNF-α inhibitors and corticosteroids, and congenital mutations affecting IFN-γ pathways are critical risk factors for dissemination (McKinsey, 2021) (Ferreira et al., 2019). Additionally, individuals at the extremes of age are particularly susceptible, emphasizing the importance of identifying and managing these populations (Wheat, 1982) (Benedict, 2023).

**Pathogenesis and Persistence Mechanisms:** Upon inhalation, H. capsulatum's conidia convert to yeast forms within the alveolar environment and are phagocytosed by macrophages (McKinsey & McKinsey, 2011) (Arango et al., 2011). In healthy hosts, activated CD4+ T-cells enhance macrophage capacity to eliminate the fungus through essential cytokine signaling involving TNF-α and IFN-γ (McKinsey, 2021) (Chai et al., 2015). Conversely, in immunosuppressed individuals, the inability to adequately activate macrophages leads to excessive intracellular proliferation of the fungus, resulting in hematogenous dissemination and severe manifestations of illness (Hajjeh, 1995) (Chai et al., 2015).

**Immune Arm Dependencies:** Successful defense against histoplasmosis depends on the interplay of CD4+ and CD8+ T-cell-mediated immune responses alongside macrophage activation, primarily through IL-12 and IFN-γ signaling pathways (Bento et al., 2025). Disruptions in these immune responses can precipitate uncontrolled proliferation of H. capsulatum, leading to extensive infection dissemination (Wheat, 1997) (Benedict, 2023).

**Compartmental Tropism and Failure:** Disseminated histoplasmosis frequently involves vital organs such as the liver, spleen, adrenal glands, gastrointestinal tract, CNS, and bone marrow (Park et al., 2020). Particularly concerning is adrenal involvement, which can lead to adrenal insufficiency, alongside CNS manifestations that can be dire, often fatal, in severely immunocompromised patients, such as those with AIDS (Diaz, 2018) (Pulgarin et al., 2021).

**Immune Modulation and Evasion:** The persistence of H. capsulatum yeast cells within macrophages is partly due to their ability to inhibit lysosomal acidification and phagolysosomal fusion (McKinsey & McKinsey, 2011) (Arango et al., 2011). Certain fungal proteins, such as CBP1, are instrumental in impairing macrophage-mediated killing and compromising complement signaling pathways, illustrating effective immune evasion tactics employed by this pathogen (Khairunnisa et al., 2023) (Chai et al., 2015).

**Immune Memory and Coordination:** Chronic exposure to H. capsulatum may evoke robust memory T-cell responses, contributing to "trained immunity" within innate immune cells (Pulgarin et al., 2021). This regional immune imprinting can mitigate disease severity in individuals with prior exposure, highlighting the significance of immunoregulation in endemic populations (Benedict, 2023) (Bento et al., 2025).

**Role in Immune Architecture:** Histoplasma antigens promote granuloma formation, reinforcing the coordination between CD4+ T-cells and macrophages during the immune response (Khairunnisa et al., 2023) (Chai et al., 2015). Persistent antigenic stimulation helps establish immunodominance hierarchies, essential for managing chronic infections (Chai et al., 2015).

**Impact of Immunosuppression:** The incidence of reactivation or severe histoplasmosis is significantly elevated among patients undergoing treatment with TNF-α inhibitors or corticosteroids and those with genetic deficits in IFN-γ or IL-12 signaling pathways (Lyon et al., 2004) (Ferreira et al., 2019). Disseminated histoplasmosis is common and often fatal in patients with untreated AIDS, underscoring the relationship between immunosuppression and the disease's severity (McKinsey, 2021) (Pulgarin et al., 2021).

**Prevalence and Future Directions:** Histoplasmosis often goes underdiagnosed in non-endemic regions, delaying timely therapeutic intervention (Benedict et al., 2021) (Pulgarin et al., 2021). Advancing diagnostic capabilities, developing antifungal vaccines, and enhancing immune-targeted therapies against H. capsulatum are critical future goals in combating this pervasive fungal pathogen (Lyon et al., 2004) (Pulgarin et al., 2021) (Bento et al., 2025). Focused inquiries into the role of histoplasmosis in shaping immune memory may yield insights beneficial to endemic populations (Khairunnisa et al., 2023) (Bento et al., 2025).

#### Toxoplasma gondii

**Classification and Transmission Ecology:** Toxoplasma gondii is an obligate intracellular protozoan parasite belonging to the phylum Apicomplexa, which primarily exploits felines as its definitive hosts while infecting nearly all warm-blooded animals, including humans, as intermediate hosts. This broad host range facilitates various transmission routes. Human infection predominantly occurs through the ingestion of tissue cysts in undercooked meat, oocysts from contaminated water or soil, and, in some cases, congenitally from mother to fetus (Hill et al., 2005) (Harker et al., 2015). The complexity of its life cycle involves multiple stages; oocysts shed by felines can persist in the environment, further contributing to transmission dynamics (Hunter & Sibley, 2012).

**Clinical Manifestations and Latency:** In immunocompetent individuals, T. gondii infection is often asymptomatic or manifests as mild febrile lymphadenopathy. The parasite's life cycle includes a stage where it encysts in neural and muscular tissues, promoting a latent infection that can last for the host's lifetime (Yamamoto et al., 2012). In contrast, immunocompromised individuals, particularly those with HIV/AIDS, face significant risks of reactivation, leading to severe clinical conditions such as encephalitis (Yamamoto et al., 2012) (Niedelman et al., 2012). This reactivation occurs due to a breakdown in the host's immune defenses, with T. gondii exploiting these vulnerabilities to cause disease (Lima & Lodoen, 2019).

**Global Distribution and Seroprevalence:** T. gondii exhibits a widespread global distribution, with seroprevalence rates varying significantly across regions, ranging from 10% to 80%. Factors influencing these rates include geographic location, climatic conditions, food preparation practices, and exposure to cats (Degrandi et al., 2012). Notably, high seroprevalence rates are observed in Latin America, parts of Europe, and sub-Saharan Africa, indicative of environmental and sociocultural determinants of exposure (Catchpole et al., 2023).

**Risk Factors for Opportunism:** Individuals with profound T-cell dysfunction, such as patients with AIDS, organ transplant recipients, and those undergoing immunosuppressive therapies, are particularly at increased risk of severe disease due to T. gondii reactivation (Niedelman et al., 2012) (Kim et al., 2016). Moreover, the risk of congenital infection is notably heightened if a mother acquires T. gondii during pregnancy, emphasizing the need for vigilance in prenatal care (Dong et al., 2022).

**Pathogenesis and Persistence Mechanisms:** T. gondii exhibits a remarkable ability to invade a range of nucleated cells, where it rapidly replicates as tachyzoites during acute infection before transitioning to bradyzoites in cysts for chronic infection. This life cycle stage enables the parasite to evade the host's immune clearance (Harker et al., 2015) (Boothroyd & Dubremetz, 2008). Noteworthy is its encapsulation within a parasitophorous vacuole, which prevents lysosomal fusion and enhances its survival within host cells (Harker et al., 2015) (Chen et al., 2020).

**Immune Arm Dependencies:** Effective control of T. gondii replication heavily relies on IFN-γ-mediated mechanisms, along with robust CD8+ and CD4+ T-cell responses (Denkers, 2003). Interleukin-12 (IL-12), tumor necrosis factor-alpha (TNF-α), and nitric oxide are pivotal in orchestrating host immune responses against T. gondii. While antibodies can limit the dissemination of tachyzoites, they are inadequate for clearing latent cysts, emphasizing the complexity of immune evasion strategies employed by the parasite (Alday & Doggett, 2017).

**Compartmental Tropism and Failure:** The CNS and muscular tissues are primary sites of T. gondii persistence, with reactivation in immunocompromised patients frequently leading to complications like necrotizing encephalitis (Harker et al., 2015). Ocular toxoplasmosis is also a significant manifestation, causing posterior uveitis associated with both congenital and postnatally acquired infections (Yamamoto et al., 2012) (Dong et al., 2022).

**Immune Modulation and Evasion:** To evade immune surveillance, T. gondii employs a variety of strategies that include secreting effector proteins that interfere with host immune signaling pathways. By modulating cytokine production and obstructing MHC (Major Histocompatibility Complex) expression, the parasite promotes long-term persistence within its host (Kim et al., 2016) (Denkers, 2003). This manipulation underscores the importance of understanding host-pathogen interactions in developing therapeutic strategies.

**Immune Memory and Coordination:** Memory T-cell responses are crucial for controlling T. gondii latency; however, they do not prevent reactivation in immunocompromised individuals. The intricacies of vaccine development remain complicated by the parasite's adeptness at evading sterilizing immunity, showcasing significant hurdles in the quest for effective preventive measures (Alday & Doggett, 2017).

**Role in Immune Architecture:** T. gondii appears to influence broader immune architecture by provoking a strong type 1 cytokine response during acute infection. In chronic stages, it shifts responses towards immune regulation to maintain its latent state. Persistent CNS infections may also lead to subtle neuroinflammatory responses and have been connected with altered behaviors in animal models (Catchpole et al., 2023).

**Impact of Immunosuppression:** The risks associated with T. gondii infection escalate significantly within populations subject to immunosuppression, such as those with HIV/AIDS, individuals undergoing organ transplantation, or those with hematological malignancies. In pregnant women, primary infections can result in adverse outcomes, including miscarriages or congenital anomalies like hydrocephalus and chorioretinitis (Yamamoto et al., 2012) (Lima & Lodoen, 2019).

**Prevalence and Future Considerations:** T. gondii remains a prominent cause of opportunistic CNS infections and congenital diseases. Currently, there is no effective vaccine for humans. Future preventative strategies may focus on early maternal screening and prophylactic measures for at-risk populations, particularly in enhancing management protocols for transplant recipients to mitigate the risk of reactivation (Catchpole et al., 2023) (Rommereim et al., 2016).

#### Trypanosoma cruzi

**Classification and Transmission Ecology**: Trypanosoma cruzi, the causative agent of Chagas disease, is classified as a flagellated protozoan parasite belonging to the family Trypanosomatidae. This parasite primarily relies on triatomine insects, commonly referred to as kissing bugs, for its transmission. The transmission occurs when these insects defecate on or near the skin of a human host, allowing the parasite to enter through mucosal surfaces or skin abrasions (Gurgel-Gonçalves et al., 2012). Although vector transmission is the predominant mode, T. cruzi can also be transmitted via congenital routes, blood transfusions, organ transplants, and ingestion of contaminated food or beverages (Oliveira et al., 2010) (Rengifo-Correa et al., 2017). The ecological dynamics of this transmission are influenced by various factors, including biodiversity in both sylvatic and domestic environments, where certain triatomine species show adaptability to urban settings, thereby increasing human exposure (Busselman & Hamer, 2022).

**Clinical Manifestations and Latency**: T. cruzi infection manifests acutely with symptoms such as fever, lymphadenopathy, myocarditis, or notably, the Romaña sign, which is characterized by periorbital edema (Macaluso et al., 2023). However, many cases remain asymptomatic, and the acute phase can resolve within weeks to months, leading into a latent or chronic phase. Upon reaching this chronic phase, approximately 20–30% of individuals may eventually develop serious complications, primarily involving the heart and gastrointestinal system, which may take decades to surface (Viana et al., 2022).

**Global Distribution and Seroprevalence**: Chagas disease has a significant presence in Latin America, where it affects more than 6 million individuals. Due to increased migration and travel, cases have been documented in non-endemic regions such as the United States, Europe, and Australia, highlighting the disease's expanding geographic footprint. In rural areas of endemic countries, seroprevalence can exceed 10%, indicating high levels of exposure and infection (Dorn et al., 2022).

**Risk Factors for Opportunism**: The principal risk factors for T. cruzi infection include being born in endemic areas, receiving blood transfusions or organ transplants from infected individuals, and congenital transmission (Boukaabar et al., 2024). Immunosuppressed individuals, such as organ transplant recipients or HIV-positive patients, are particularly susceptible to severe outcomes resulting from T. cruzi reactivation (Shikanai-Yasuda, 2016).

**Pathogenesis and Persistence Mechanisms**: Upon entering the host, T. cruzi trypomastigotes invade host cells, differentiating into intracellular amastigotes where they replicate extensively (Macaluso et al., 2023). The parasite demonstrates a predilection for cardiac and smooth muscle cells, evading immune responses through various mechanisms, including subversion of antigen presentation and taking advantage of immune-privileged sites (Macaluso et al., 2023).

**Immune Arm Dependencies**: The success of immune control over T. cruzi relies heavily on robust responses from CD8+ and CD4+ T-cells, coupled with the activation of macrophages by IFN-γ (Shikanai-Yasuda, 2016). Antibody responses contribute to limiting parasite burden in the bloodstream, yet the presence of immunosuppressive therapies can precipitate reactivation of dormant infections.

**Compartmental Tropism and Failure**: In the chronic phase of Chagas disease, T. cruzi predominantly affects the heart and enteric nervous system, leading to chronic Chagas cardiomyopathy, characterized by arrhythmias, heart failure, and potential sudden death. Gastrointestinal consequences manifest as megaesophagus or megacolon, representing severe morbidity among chronic patients (Gurgel-Gonçalves et al., 2012).

**Immune Modulation and Evasion**: T. cruzi has developed sophisticated strategies to modulate host immunity, including interference with antigen presentation mechanisms, inducing T-cell exhaustion, and the promotion of anti-inflammatory cytokines such as IL-10, ultimately enabling its persistence despite an active immune response (Shikanai-Yasuda, 2016).

**Immune Memory and Coordination**: Despite the formation of memory T-cells, immunity against T. cruzi is not sterilizing, leading to potential re-infection or reactivation under conditions of immune compromise (Macaluso et al., 2023). The long-term immune response becomes increasingly complex due to chronic antigen stimulation, which may lead to a decline in the functionality of these memory cells.

**Role in Immune Architecture**: The presence of T. cruzi can significantly alter the immune landscape, driving changes in T-cell populations and tissue architecture within the heart and enteric tissues over prolonged infection, thereby influencing local and systemic immune responses (Macaluso et al., 2023).

**Impact of Immunosuppression**: Reactivation of Chagas disease poses significant risks, especially for organ transplant recipients and HIV/AIDS patients, manifesting in serious clinical complications such as myocarditis, meningoencephalitis, or cutaneous lesions (Shikanai-Yasuda, 2016). The immune dynamics in these groups highlight the importance of early detection and treatment.

**Prevalence and Future Considerations**: Despite advances in treatment and screening, Chagas disease remains significantly underdiagnosed. The current therapeutic options, such as benznidazole and nifurtimox, face limitations including poor tolerance, particularly among adults. Future research priorities focus on developing improved diagnostics, safer and more effective treatments, and potential vaccines to combat the disease (Macaluso et al., 2023).

### MAC: Mycobacterium avium complex

**Classification and Transmission Ecology:** Mycobacterium avium complex (MAC) refers primarily to two species: Mycobacterium avium and Mycobacterium intracellulare, both of which are classified as non-tuberculous mycobacteria (NTM). These bacteria are commonly found in various environmental reservoirs, including soil, freshwater, and drinking water systems, which serve as their ecological niches (Falkinham et al., 2001) (To et al., 2020). Unlike Mycobacterium tuberculosis, which is primarily transmitted through person-to-person contact, MAC is acquired through environmental exposure, predominantly via inhalation or ingestion. Researchers have identified biofilms in water distribution systems as key factors contributing to increased exposure to MAC, particularly in immunocompromised individuals (Falkinham et al., 2001) (To et al., 2020).

**Clinical Manifestations and Latency:** MAC infections present a broad spectrum of clinical manifestations. Patients may experience asymptomatic colonization, chronic pulmonary infections, or disseminated disease, especially in those with significant immunocompromise, such as individuals with untreated HIV/AIDS or on immunosuppressive therapy (Amin et al., 2023) (Khan et al., 2005). Symptoms for pulmonary infections often resemble those seen in tuberculosis, including cough, fatigue, weight loss, and specific imaging findings such as nodules or bronchiectasis (Amin et al., 2023) (Khan et al., 2005). In extreme cases, disseminated MAC disease can occur, leading to serious complications, particularly in patients with very low CD4+ T-cell counts (Amin et al., 2023).

**Global Distribution and Seroprevalence:** MAC organisms have a worldwide presence, with documented higher prevalence rates in countries like Japan, North America, and parts of Europe. This increased prevalence is frequently observed in aging populations or those with pre-existing lung conditions, emphasizing the need for enhanced surveillance and recognition of the disease (Goepfert et al., 2014) (Nukaga et al., 2021). Epidemiological studies reveal significant rates of seropositivity among wildlife, such as wild boars and carnivores in the Iberian Peninsula, indicating a wider ecological distribution (Matos et al., 2018).

**Risk Factors for Opportunism:** Key risk factors for opportunistic MAC infection include structural lung disease (such as COPD, bronchiectasis, and cystic fibrosis), severe immunosuppression (notably in patients with CD4+ T-cell counts below 50 cells/μL), and the use of tumor necrosis factor (TNF) inhibitors (Amin et al., 2023) (Goepfert et al., 2014). Notably, the condition termed "Lady Windermere syndrome" highlights the occurrence of pulmonary MAC disease in elderly women lacking apparent immunosuppressive factors (Baylor & Larson, 2008), suggesting complex interactions between the pathogen and host susceptibility.

**Pathogenesis and Persistence Mechanisms:** Distinct from tuberculosis, MAC does not create latent infections but relies on continuous environmental acquisition. Studies indicate variability in virulence among different MAC strains, with some demonstrating enhanced capabilities for macrophage invasion and survival (To et al., 2020) (Amin et al., 2023). There is also in vitro evidence suggesting that MAC can augment HIV replication in infected individuals by increasing CCR5 expression on macrophages, thus linking the two pathogens (Crandall et al., 2012) (Babu et al., 2013).

**Immune Arm Dependencies:** Successful control of MAC infections necessitates a robust immune response, predominantly involving CD4+ T cells, macrophages, and effective TNF signaling pathways. Immunocompromised patients who lack these critical components face heightened vulnerability to severe pulmonary or disseminated MAC diseases (Crandall et al., 2012) (Amin et al., 2023). Research emphasizes the importance of TNF in granuloma formation—an essential immune response aimed at containing mycobacterial infections (Crandall et al., 2012).

**Compartmental Tropism and Failure:** The anatomical distribution of MAC infections often leads to pulmonary involvement primarily affecting the middle lobe and lingula, especially in patients with underlying bronchiectasis. Conversely, disseminated MAC can compromise various organs, including the bone marrow, liver, lymph nodes, and gastrointestinal tract, often leading to specific manifestations such as hepatosplenomegaly and enteropathy (Amin et al., 2023) (Horné & Skerrett, 2019) (Goepfert et al., 2014).

**Immune Modulation and Evasion:** While MAC may not utilize classical immune evasion tactics like antigenic variation, it can indirectly modulate host immune responses, particularly in HIV-coinfected patients by enhancing macrophage CCR5 expression (Crandall et al., 2012) (Babu et al., 2013). Moreover, the disruption of granuloma architecture is a significant pathogenic strategy that allows MAC to persist and thrive in the host (Crandall et al., 2012).

**Immune Memory and Coordination:** Although MAC does not persist in the body as latent TB does, chronic exposure may lead to granuloma formation, fostering a form of immune imprinting. This response suggests that prolonged interactions with MAC can stimulate a refined immune memory (Crandall et al., 2012).

**Role in Immune Architecture:** MAC is not considered a classical keystone pathogen; however, it plays a critical role in maintaining immune architecture through its necessity for multi-arm immune coordination. Its reliance on TNF, IFN-γ, and T-cell activity enforces a structured immune response, which can be detrimental when underlying immunosuppression occurs (Crandall et al., 2012) (Amin et al., 2023).

**Impact of Immunosuppression:** The dissemination of MAC is notably pronounced in untreated AIDS, particularly when patients' CD4 counts fall below 50 cells/µL. Furthermore, therapies involving TNF inhibitors dramatically increase risks of MAC infections, particularly extrapulmonary cases (Amin et al., 2023) (Goepfert et al., 2014) and are also seen in transplant recipients and those with idiopathic CD4+ lymphocytopenia (Amin et al., 2023).
**Prevalence and Future Considerations**: MAC remains a salient pathogen, causing recurrent infections in both immunocompetent and HIV-infected individuals. There is a growing recognition of atypical presentations like Lady Windermere syndrome, notably in elderly female demographics (Nukaga et al., 2021) (Crandall et al., 2012). The evolution of diagnostics and a deeper understanding of genetic susceptibility—particularly towards bronchiectasis—are pivotal for managing MAC infections going forward (Goepfert et al., 2014) (Nukaga et al., 2021).

#### Pneumocystis jirovecii (PJP)

**Classification and Transmission Ecology:** Pneumocystis jirovecii is classified as an atypical fungus due to its unique characteristics and behavior within host organisms. Its life cycle includes both trophic forms and cystic forms, which are crucial for its survival and transmission. The organism primarily spreads through airborne particles, and there is evidence suggesting potential interhuman transmission via respiratory droplets, particularly in immunocompromised populations (Nevez et al., 2008). Most humans are exposed to Pneumocystis jirovecii by the age of four, as indicated by serologic studies, underscoring its widespread nature (Nevez et al., 2008) (Eddens & Kolls, 2014). While colonization is typically asymptomatic in immunocompetent individuals, it may serve as a reservoir for future infections, particularly in patients with compromised immune systems, such as those undergoing immunosuppressive therapies (Eddens & Kolls, 2014).
**Clinical Manifestations and Latency:** Pneumocystis pneumonia (PJP) often presents with an insidious onset marked by symptoms such as fever, dry cough, and dyspnea, worsening over days to weeks (Eddens & Kolls, 2014). Hypoxemia frequently complicates these symptoms, accompanied by specific radiographic findings, including bilateral infiltrates and ground-glass opacities, which are indicative of infection (Walzer et al., 2009). Histologically, Pneumocystis can be detected within alveolar exudates, confirming the diagnosis during clinical assessments (Tesini et al., 2017). The notion of latency in Pneumocystis jirovecii infection remains debated, with some studies arguing that most clinical cases arise from reexposure rather than reactivation of a dormant infection (Eddens & Kolls, 2014).
**Global Distribution and Seroprevalence:** Pneumocystis jirovecii has a global presence, colonizing the lungs of many healthy individuals. Studies indicate considerable variability in seroprevalence, with colonization rates up to 20% in immunocompetent adults and higher among hospitalized patients with pulmonary infections (Shimizu et al., 2009). This widespread colonization suggests that while many are asymptomatic carriers, effective disease management remains critical, especially during periods of immune compromise (Walzer et al., 2009).
**Risk Factors for Opportunism:** The primary risk factor for the development of PJP is significant T-cell immunosuppression, particularly in individuals with CD4+ counts below 200 cells/µL, such as those with HIV/AIDS (Walzer et al., 2009) (Marimuthu, 2019). Other notable at-risk groups

include solid organ transplant recipients and individuals undergoing treatment with corticosteroids or biologic agents, including TNF-α inhibitors. These immunosuppressed populations are at heightened risk for opportunistic infections, particularly during invasive procedures or in the presence of additional infections (Walzer et al., 2009) (Kottom et al., 2021).

**Pathogenesis and Persistence Mechanisms:** Pneumocystis jirovecii initiates infection through adhesion to alveolar epithelial cells, eliciting immune responses from the host. In conditions such as HIV infection, impairments in macrophage function and cytokine signaling hinder the clearance of the organism from the lungs, contributing to pathogenesis (Tesini et al., 2017) (Kottom et al., 2021). The organism's cell wall components, notably β-glucans, play a significant role in activating immune responses, while antigenic variation in the major surface glycoprotein (MSG) allows for evasion of host immune detection (Kottom et al., 2021).

**Immune Arm Dependencies:** Effective clearance of PJP heavily relies on the presence and function of CD4+ T-cells, which are essential for activating and recruiting alveolar macrophages to combat the infection (Eddens & Kolls, 2014). Although CD8+ T-cells may offer ancillary protection, they can also inadvertently contribute to lung injury in chronic cases where CD4+ counts are low (Hadfield et al., 2019). Neutrophils additionally play a critical role in mediating inflammatory responses (Walzer et al., 2009).

**Compartmental Tropism and Failure:** Pneumocystis jirovecii is localized within the alveolar compartments of the lungs. Its pathogenic mechanisms primarily affect pulmonary tissue, leading to significant disease states, including diffuse alveolar damage and associated respiratory failure (Hadfield et al., 2019). Instances of disseminated infection are rare and are largely confined to severely immunocompromised hosts (Walzer et al., 2009).

**Immune Modulation and Evasion:** The capacity of Pneumocystis to express surface glycoproteins that undergo antigenic variation is crucial for immune evasion. The predominant surface antigen, MSG, facilitates this variation, allowing persistence even in previously immunocompetent hosts (Tesini et al., 2017). This feature underscores the challenges faced in developing effective vaccines and therapeutic strategies against PJP (Grønseth et al., 2021).

**Immune Memory and Coordination:** Unlike many viral pathogens, Pneumocystis does not seem to induce robust systemic immune memory, though cases of repeated colonization may enhance innate immune responses without generating long-lasting adaptive immunity (Walzer et al., 2009) (Grønseth et al., 2021). This subtle interplay could influence disease dynamics, particularly in endemic regions with a high burden of exposure (Grønseth et al., 2021).

**Role in Immune Architecture:** The emergence of PJP in immunosuppressed patients underscores the critical role of CD4+ T-cell surveillance in maintaining pulmonary immune homeostasis. The inability of the immune system to control this opportunistic pathogen marks a failure point, often preceding secondary infections by other opportunistic pathogens, including cytomegalovirus (CMV) and Mycobacterium avium complex (MAC) (Kottom et al., 2021).

**Impact of Immunosuppression:** Significant immunosuppression, particularly through therapies involving corticosteroids or agents that significantly lower T-cell counts, correlates with an increased incidence of PJP. This relationship emphasizes the need for vigilant surveillance and prophylactic measures in at-risk populations, particularly those with chronic conditions or who are subjected to potent immunosuppressive treatments (Marimuthu, 2019) (Grønseth et al., 2021).

**Prevalence and Future Considerations:** Despite advances in antiretroviral therapy and prophylactic approaches, PJP continues to be a leading cause of morbidity and mortality among immunocompromised individuals, especially those with untreated HIV or late-presenting cases (Eddens & Kolls, 2014) (Kottom et al., 2021). Future research directions may involve refining clinical guidelines for prophylaxis based on immune profile assessments and exploring innovative therapeutic avenues, including vaccine development (Grønseth et al., 2021).

#### Candida albicans (Candida)

**Classification and Transmission Ecology:** Candida albicans is a eukaryotic organism classified as a yeast, being a significant member of the fungal kingdom and part of the normal human microbiota. It resides primarily on mucosal surfaces including the oral cavity, gastrointestinal tract, and genital mucosa in healthy individuals (Silva et al., 2012). Transmission predominantly occurs through the endogenous overgrowth of genetically similar strains as opposed to external acquisition, which emphasizes its nature as a commensal organism (Guinea, 2014). Studies indicate that about 50-70% of healthy adults carry C. albicans without clinical disease manifestations, highlighting its role as a normal flora constituent (Deorukhkar et al., 2014).
**Clinical Manifestations and Latency:** C. albicans is responsible for a variety of clinical manifestations ranging from superficial mucosal infections, such as oropharyngeal candidiasis (thrush), esophageal candidiasis, and vulvovaginal candidiasis, to invasive infections in immunocompromised patients, leading to complications like candidemia and endophthalmitis (Chow et al., 2008). The underlying mechanisms triggering these conditions often relate to the host's immune status; infections are predominantly seen in severely immunodeficient patients, such as those with advanced HIV/AIDS, where C. albicans may lead to systemic disease (Chow et al., 2008) (Sarıgüzel et al., 2015).
**Global Distribution and Seroprevalence:** Globally, C. albicans remains the most prevalent species responsible for candidiasis, accounting for a significant percentage of infections reported in various studies, particularly regarding candidemia (Deorukhkar et al., 2014) (Pfaller et al., 2000). C. albicans typically colonizes a significant percentage of the population asymptomatically, with colonization rates estimated at around 50-70% (Guinea, 2014). Its widespread nature reflects the favorable conditions in various environments of the human body, promoting its establishment as a common pathogen.
**Risk Factors for Opportunism:** The development of mucocutaneous candidiasis is notably linked with impaired cellular immunity, particularly characterized by Th17 dysfunction. Noteworthy risk factors include HIV/AIDS, diabetes, chronic corticosteroid use, previous antibiotic exposure, extended use of dental appliances such as dentures, and radiation treatment impacting head and neck areas (Fan et al., 2008). Consequently, immune compromise enables overgrowth of C. albicans, increasing the risk for opportunistic infections (Deorukhkar et al., 2014) (Mohandas & Ballal, 2011).
**Pathogenesis and Persistence Mechanisms:** The pathogenicity of C. albicans is linked intricately to both the immune system's integrity and mucosal barrier integrity. The organism employs a range of virulence mechanisms, including the expression of adhesins and hydrolytic enzymes, allowing it to invade epithelial tissues. Chronic infections, particularly in cases of mucocutaneous candidiasis, are often associated with defects in immune signaling pathways, especially those

involving IL-17 (Richter et al., 2005) (Tavanti et al., 2001). Such immune evasion tactics are crucial for its persistence in the host environment.

**Immune Arm Dependencies:** Robust immune responses characterized by Th17 pathways play a vital role in controlling C. albicans infections at mucosal surfaces, necessitating neutrophil recruitment for effective containment. Conversely, disseminated infections require the activation of macrophages and neutrophils to clear the pathogen (Negi et al., 2019). The IL-17 signaling pathway is critical for the host's defense by enhancing mucosal immunity, illustrating the necessity for a well-orchestrated immune response (Negi et al., 2019) (Akbar & Tahawi, 2001) (St-Germain et al., 2001).

**Compartmental Tropism and Failure:** Mucosal candidiasis typically reflects a failure in Th17 immune responses, as C. albicans invades the bloodstream and disseminates to vital organs such as the kidneys, liver, spleen, and central nervous system, particularly in patients with neutropenia or critical illness. This trend of dissemination indicates failures in localized immune responses and underscores the vulnerability of immunocompromised patients to invasive candidiasis (Deorukhkar et al., 2014) (Akbar & Tahawi, 2001).

**Immune Modulation and Evasion:** C. albicans utilizes several mechanisms to evade the host immune system, including phenotypic switching, protective shielding of β-glucans, and production of secreted aspartyl proteinases (Saps), which interfere with epithelial barriers and immune signaling pathways (Arendrup et al., 2013) (Xin et al., 2008). These mechanisms not only promote colonization but also complicate therapeutic interventions.

**Immune Memory and Coordination:** While C. albicans does not induce the same sustained immune memory as classical pathogens such as herpesviruses, it plays a crucial role in shaping mucosal immunity. Exposure to C. albicans can stimulate protective Th17 immune responses, potentially generalizing to offer protection against other fungal pathogens, indicating a capacity for cross-reactive immunity (Rodrigues et al., 2017) (Werner et al., 2020).

**Role in Immune Architecture:** C. albicans acts as an "immunological rheostat," modulating mucosal immune responses and maintaining homeostasis at barrier sites. The establishment of Th17 responses is significant for setting immune surveillance thresholds and controlling pathogen invasion (Lupetti et al., 2002) (Borman et al., 2013).

**Impact of Immunosuppression:** Candidiasis is frequently observed as a complication during immunodeficiency stages, notably in HIV/AIDS when CD4+ T cell counts drop below critical thresholds (Hedderwick et al., 2000). This opportunistic nature of C. albicans highlights its role as a sentinel marker for monitoring mucosal immune decline in patients undergoing immunosuppressive therapies (Cutler et al., 2011) (Sobel et al., 2003).

**Prevalence and Future Considerations:** The prevalence of Candida infections is expected to rise with the increased use of systemic immunosuppressive therapies. Targeting therapeutic strategies focused on Th17 immune pathways could potentially reduce susceptibility to C. albicans infections. Continued research into the fungus's role in mucosal immunity, along with exploring cross-fungal immune training, may pave the way for innovative vaccine designs and improved therapeutic strategies (Shigemura et al., 2014) (King et al., 2011).

#### Plasmodium species (Malaria)

**Classification and Transmission Ecology:** Plasmodium falciparum is a protozoan parasite classified within the Apicomplexa phylum, specifically falling under the genus Plasmodium. It is one of the five species of Plasmodium that cause human malaria, alongside P. vivax, P. ovale, P. malariae, and P. knowlesi. Transmission of these species occurs primarily through the bite of infected Anopheles mosquitoes, which inject sporozoites during their blood meal. After reaching the human bloodstream, the sporozoites migrate to the liver where they undergo asexual reproduction, initiating the malaria infection (Wipasa et al., 2010) (Weiss et al., 2010). Human malaria caused by P. falciparum is particularly severe and accounts for the majority of malaria-related morbidity and mortality globally (Tse et al., 2014).

**Clinical Manifestations and Latency:** Infection with P. falciparum typically presents with fever, chills, headache, myalgia, and malaise. However, severe cases can escalate to cerebral malaria, acute respiratory distress syndrome (ARDS), hemolytic anemia, and multi-organ failure (Bhumiratana et al., 2013). Notably, while P. vivax and P. ovale can remain latent in the liver as hypnozoites and can reactivate to cause relapses weeks or months after the initial infection (Gray et al., 2018), P. falciparum does not exhibit this latency.

**Global Distribution and Seroprevalence:** Malaria, particularly from P. falciparum, is endemic in over 90 countries, with the highest burden observed in sub-Saharan Africa. Southeast Asia and certain areas of Latin America also contribute significantly to the global incidence. Seroprevalence, reflecting past exposure, varies widely among populations and is influenced by factors such as age and the level of endemicity. Individuals exposed repeatedly to malaria infections exhibit higher frequencies of antibodies; however, these antibodies indicate past exposure rather than effective sterilizing immunity (Nogaro et al., 2011) (Fowkes et al., 2016).

**Risk Factors for Opportunism:** Certain demographic and health factors markedly increase the risk of severe P. falciparum infection. Non-immune travelers, young children (especially under 5 years), pregnant women, individuals with HIV/AIDS, and those with asplenia or congenital hemoglobinopathies face significantly heightened vulnerability. Additional risks are associated with immunosuppressive therapies and underlying malnutrition, which may impair the host's ability to combat malaria effectively (Cook et al., 2012) (Tripet, 2009).

**Pathogenesis and Persistence Mechanisms:** The lifecycle of P. falciparum involves complex interactions with the human host. After hepatic replication, merozoites enter the bloodstream and infect red blood cells, leading to cyclic ruptures that release more merozoites, causing an inflammatory response. The expression of PfEMP1 on the surface of infected erythrocytes facilitates their adhesion in vascular tissues, enabling the parasite to evade host immune detection through antigenic variation, a mechanism that allows P. falciparum to maintain chronic infections (Águas et al., 2012) (Krzych et al., 2014).

**Immune Arm Dependencies:** The immune response to P. falciparum is multifaceted, involving both innate and adaptive immunity. Toll-like receptors (TLRs) recognize the parasite and initiate inflammatory cytokine responses, which help limit the spread of the infection. CD4+ and CD8+ T cells, along with B cells, play critical roles in generating immune memory; however, immunity remains incomplete without continuous exposure to the parasite (Ly & Hansen, 2019) (Stevenson et al., 2011).

**Compartmental Tropism and Failure:** P. falciparum has a propensity for targeting both the hepatic and erythrocytic compartments of the human host. Complications arise when parasitized erythrocytes sequester in microvasculature of organs such as the brain, lungs, and kidneys, leading to severe outcomes. In individuals without a functioning spleen, the ability to clear infected erythrocytes diminishes, heightening the severity of malaria infections (Lefebvre & Harty, 2020).
**Immune Modulation and Evasion:** P. falciparum effectively modulates the immune response to promote its survival. Strategies include the antigenic variation of surface proteins, such as PfEMP1, and inhibiting dendritic cell maturation, which reduces the host's ability to mount an effective immune response. Chronic infections also lead to immune exhaustion, wherein continual antigenic stimulation diminishes the functional capacity of T cells ((Moormann, 2009) (Rönnberg et al., 2021).
**Immune Memory and Coordination:** The formation of immune memory against malaria is contingent upon repeated infections, as the immunity generated tends to be non-sterilizing and short-lived. Antibody levels wane considerably in the absence of further infections, highlighting the incomplete immune response characteristic of malaria, as particularly observed in both adult and pediatric populations after leaving endemic areas (Mooney et al., 2015) (Furtado et al., 2023).
**Role in Immune Architecture:** The chronic presence of P. falciparum in endemic areas has significant implications for the restructuring of immune function. In contrast to persistent viral infections, which modulate immune architecture systematically, malaria, due to its reliance on insect vectors for transmission, results in fluctuating immune responses across populations, particularly affecting long-term immunity and adaptive immune responses (Carter & Mendis, 1992) (Obeagu & Obeagu, 2024).
**Impact of Immunosuppression:** Individuals who are immunosuppressed due to conditions such as HIV/AIDS or therapies like corticosteroids exhibit higher parasitemia, prolonged infection duration, and increased severity of malaria symptoms. Co-infection with HIV has been shown to compromise the adaptive immune response to malaria, worsening clinical outcomes (Li et al., 2023) (Mura et al., 2019).
**Prevalence and Future Considerations:** Despite advancements in malaria control, the disease continues to impose a significant burden, with an estimated 247 million infections and 619,000 deaths reported in 2021. Ongoing research focuses on understanding the complexities of immune responses, particularly concerning antigenic variation and the mechanisms behind incomplete immune memory. Insights from this research are crucial for the development of more effective malaria vaccines and treatment strategies (Chen et al., 2023) (Tartz et al., 2013).

#### Aspergillus fumigatus (Aspergillus)

**Classification and Transmission Ecology:** Aspergillus fumigatus is classified as a filamentous fungus within the genus Aspergillus, notable for its ubiquitous environmental presence, especially in soil and decaying organic matter. Transmission occurs primarily through inhalation of airborne conidia, which are capable of surviving in varying environmental conditions, allowing them to thrive in suboptimal habitats for other microorganisms (Latgé & Chamilos, 2019) (Park & Mehrad, 2009). Infection in humans predominantly occurs in immunocompromised conditions, as the fungus is not adapted to person-to-person transmission (Latgé & Chamilos, 2019) (Kolwijck & Frank, 2014). Such ecological adaptability supports a significant environmental reservoir while limiting its capacity for persistent human transmission (Kosmidis & Denning, 2014).

**Clinical Manifestations and Latency:** Aspergillus leads to a range of pulmonary conditions varying from benign colonization to aggressive invasive disease. In immunocompetent individuals with underlying lung conditions, such as chronic obstructive pulmonary disease (COPD), chronic pulmonary aspergillosis (CPA) may manifest as a slow-progressing cavitary disease, sometimes with the formation of aspergillomas (Kosmidis & Denning, 2014) (Dogra et al., 2016). For immunocompromised patients, invasive aspergillosis (IA) emerges as an acute and often lethal pulmonary infection, characterized by its angio-invasive nature, leading to high mortality rates due to rapid disease progression and potential dissemination to various organs (Chotirmall et al., 2013). Additionally, allergic bronchopulmonary aspergillosis (ABPA) may occur, particularly in patients with asthma and cystic fibrosis, linked to hypersensitivity to Aspergillus antigens that exacerbate respiratory symptoms (Speth et al., 2019) (Goh et al., 2017).

**Global Distribution and Seroprevalence:** Globally, A. fumigatus is recognized as the most frequent species linked to chronic pulmonary aspergillosis (CPA) and invasive aspergillosis (IA). Its distribution extends across various climates and geographic regions, flourishing particularly in humid environments (Chotirmall et al., 2013) (Goh et al., 2017). Estimates suggest that over three million individuals may be affected by CPA globally, with significant prevalence in regions burdened by tuberculosis, highlighting the intersection of infectious diseases and fungal pathogenesis (Shao et al., 2005). This relationship underscores the necessity for better surveillance and public health strategies to manage the impact of Aspergillus globally.

**Risk Factors for Opportunism:** The principal risk factor for invasive aspergillosis is severe immunosuppression, commonly manifested as neutropenia and exacerbated by high-dose corticosteroid therapies, chemotherapeutic regimens, and biologic agents such as TNF inhibitors (Kosmidis & Denning, 2014) (Shikha Bhatia et al., 2011). CPA, by contrast, is often associated with structural lung diseases, particularly post-tuberculosis and chronic pulmonary conditions, indicating a need for focused research to identify patients at heightened risk for both conditions (Lass-Flörl et al., 2013) (Rosowski et al., 2018).

**Pathogenesis and Persistence Mechanisms:** Upon inhalation, Aspergillus spores germinate in the pulmonary airways, evading immune responses such as phagocytosis. This is achieved through various mechanisms, including the secretion of proteases and polysaccharides that mask immunogenic structures like β-glucans (Netea et al., 2003) (Kosmidis & Denning, 2014). Neutrophils and macrophages play decisive roles in recognizing and combating the fungus, with β-glucan detection being crucial for immune activation (Latgé & Chamilos, 2019; (Rosowski et al., 2018; (Latgé & Chamilos, 2019) (Rosowski et al., 2018). Invasive forms penetrate tissues, leading to necrosis and potential systemic spread, complicating the clinical picture (Netea et al., 2003) (Speth et al., 2019).

**Immune Arm Dependencies:** Neutrophils and macrophages are essential for the effective clearance of Aspergillus. In neutropenic patients, IA develops rapidly, highlighting the critical role of these immune cells (Kolwijck & Frank, 2014) (Knox et al., 2014). Genetic vulnerabilities, such as defects in Th1 immune pathways linked to reduced interferon-gamma production, further underscore individual susceptibility to infection (Lass-Flörl et al., 2013) (Rosowski et al.,

2018). Conditions like chronic granulomatous disease and hyper-IgE syndrome exemplify how genetic and immunologic deficiencies can heighten infection risk, necessitating preventive strategies in at-risk populations (Speth et al., 2019) (Shikha Bhatia et al., 2011).
**Compartmental Tropism and Failure:** Aspergillus primarily targets the lungs, but disseminated infections can affect other sites, notably the central nervous system and skin, particularly in those with profound immunosuppression (Chotirmall et al., 2013) (Shikha Bhatia et al., 2011). Chronic pulmonary aspergillosis is typically localized to pre-existing cavitary lesions, where the formation of aspergillomas can occur, leading to complications such as hemorrhage (Hughes & Boomsma, 2004) (Shao et al., 2005). Dissemination becomes a critical concern in severely immunosuppressed patients due to the angio-invasive properties of the fungus (Ordaya et al., 2021).
**Immune Modulation and Evasion:** A. fumigatus employs several strategies to evade host immune responses, including the secretion of secondary metabolites and proteases that hinder phagocytic activity (Latgé & Chamilos, 2019) (Netea et al., 2003). While this species may lack dedicated immune evasion genes, its ability to alter immune responses represents a sophisticated form of pathogenicity that complicates typical immunological defense mechanisms (M. J. Lee et al., 2015) (Gravelat et al., 2013). This immune suppression can result in rapid infection progression among susceptible populations, illustrating the challenge posed by this environmental pathogen (Rosowski et al., 2018) (Shao et al., 2005).
**Immune Memory and Coordination:** Current evidence suggests a lack of long-term adaptive immune memory to A. fumigatus in most hosts. However, repeated exposure may lead to trained immunity, particularly involving macrophages and neutrophils, and in certain contexts may stimulate adaptive responses, as seen in ABPA (Shikha Bhatia et al., 2011) (Lass-Flörl et al., 2013). This raises questions about the potential for immune conditioning as a therapeutic avenue for recurrent infections in high-risk groups (Beck et al., 2006) (Shikha Bhatia et al., 2011).
**Role in Immune Architecture:** Aspergillus serves predominantly as an intermittent environmental challenge rather than establishing persistent immunological imprinting like chronic viruses. Its interactions with the immune system are generally transient and elicit robust responses primarily during active infections (Cramer et al., 2011). While it influences local immune polarization, especially in allergic responses, it does not centralize systemic immune architecture (Shao et al., 2005) (Kolwijck & Frank, 2014).
**Impact of Immunosuppression:** Factors such as neutropenia, prolonged corticosteroid use, and targeted biologics critically increase the risk for invasive fungal diseases (Kosmidis & Denning, 2014) (Shikha Bhatia et al., 2011). Treatments with TNF inhibitors and IL-6 blockers are correlated with heightened susceptibility to opportunistic infections, emphasizing the balance required in managing immunosuppression and preventing Aspergillus infections (Rosowski et al., 2018) (Lass-Flörl et al., 2013).
**Prevalence and Future Considerations:** Chronic pulmonary aspergillosis remains underdiagnosed worldwide, posing substantial challenges for healthcare providers (Shao et al., 2005) (Speth et al., 2019). Future research is imperative to improve serological diagnostics, monitor antifungal resistance patterns, and refine immune risk stratification tools. Establishing more precise thresholds of immune impairment could facilitate the transition from fungal colonization to symptomatic disease—a critical area for targeted intervention in

immunocompromised patients (Latgé & Chamilos, 2019; Kolwijck & Veerdonk, 2014) (Latgé & Chamilos, 2019) (Kolwijck & Frank, 2014).

#### Cocci: Coccidioides spp.

**Classification and Transmission Ecology:** Coccidioides spp., comprising C. immitis and C. posadasii, are recognized as dimorphic fungi endemic to arid regions of the southwestern United States, parts of northern Mexico, and select areas of Central and South America. Infection occurs primarily through the inhalation of airborne arthroconidia released from disturbed soils, underscoring the environmental and ecological niches these fungi occupy (Cox & Magee, 2004). The transition from a mycelial phase in soil to a pathogenic spherule phase upon inhalation is critical in the pathogenesis of coccidioidomycosis, as highlighted by (Cox & Magee, 2004). Studies suggest that increasing human interaction with these environments, coupled with climate changes, may augment the risks of exposure and consequently infection rates (Engelthaler et al., 2016) (Stewart & Thompson, 2016).

**Clinical Manifestations and Latency:** The clinical spectrum of coccidioidomycosis usually includes a self-limited pulmonary illness presenting with fever, cough, fatigue, and night sweats (Parish & Blair, 2008) (Thompson, 2011). While many individuals exhibit mild or asymptomatic forms, immunologic responses can lead to distinctive conditions such as erythema nodosum or joint symptoms often referred to as "desert rheumatism" (Galgiani et al., 2000). The disease's potential for dissemination to various organs, notably skin, bones, and the central nervous system (CNS), is heightened in immunocompromised populations (Parish & Blair, 2008) (Blair, 2007). For instance, individuals with HIV/AIDS and other immunosuppressive conditions experience a higher incidence of severe manifestations, including widespread dissemination (Singh et al., 1996) (Stewart & Thompson, 2016). The latent capacity for Coccidioides spp. to persist after initial exposure further complicates clinical management, as reactivation may occur following future immunosuppression (Thompson, 2011) (Glenn et al., 2005).

**Global Distribution and Seroprevalence:** Coccidioides spp. have a well-established endemic range, particularly in the San Joaquin Valley of California and parts of Arizona, as well as in Mexico and Central America (Laniado-Laborín, 2007) (Stewart & Thompson, 2016). Recent epidemiological data suggest a rising incidence and geographic expansion of coccidioidomycosis linked to climate changes and demographic shifts, such as increased tourism and migration to endemic regions (Parish & Blair, 2008) (Engelthaler et al., 2016) (Freedman et al., 2018). Improved clinical awareness and diagnostic capabilities have also contributed to higher reported case numbers, broadening the recognition of the disease in non-endemic areas (Stewart & Thompson, 2016).

**Risk Factors for Opportunism:** Disseminated coccidioidomycosis is particularly prevalent among individuals with compromised immune systems, including those with HIV (specifically with CD4+ counts below 250 cells/µL), organ transplant recipients, and individuals undergoing corticosteroid therapy or using TNF inhibitors (Miller et al., 2004) (Saling et al., 2021). Genetic mutations affecting the immune response, such as defects in the pathways of interferon-gamma (IFN-γ) and interleukin-12 (IL-12), further elevate the risk of severe disease manifestations (Miller et al., 2004) (Singh et al., 1996). Thus, demographic and health-related factors critically influence disease outcomes in infected populations.

**Pathogenesis and Persistence Mechanisms:** After inhalation, arthroconidia convert into spherules, generating endospores that facilitate further infection (Cox & Magee, 2004) (Bronnimann et al., 1987). The immune response, particularly the activation of T-cells and macrophages mediated by IFN-γ, is pivotal in controlling infection; robust granuloma formation signifies an adequate immune defense against the pathogen (Saling et al., 2021). The role of immunomodulatory proteins such as spherule outer wall glycoprotein (SOWgp) highlights the fungi's ability to manipulate host immune responses, contributing to its persistence and pathogenicity (C.-Y. Lee et al., 2015) (Miller et al., 2004).

**Immune Arm Dependencies:** A coordinated cellular immune response is essential for successful containment of coccidioidomycosis; CD4+ T cells and macrophages are critical in orchestrating this response, while cytokines like IFN-γ and TNF-α are necessary for effective fungal clearance (Singh et al., 1996) (Stewart & Thompson, 2016). Deficiencies in any of these immune mechanisms can lead to poorly formed granulomas, allowing for expansive fungal growth and dissemination (Parish & Blair, 2008) (Saling et al., 2021) (Stewart & Thompson, 2016).

**Compartmental Tropism and Failure:** The primary manifestations of coccidioidomycosis are pulmonary, but there is a notable risk for dissemination, particularly towards the CNS, bones, and skin (Thompson, 2011) (Singh et al., 1996). Importantly, the propensity for dissemination is markedly amplified in individuals with depressed immune systems, where granulomatous responses are often inadequate, leading to increased fungal burden (Mendoza et al., 2015) (Kokseng & Blair, 2010).

**Immune Modulation and Evasion:** Coccidioides spp. exhibit several mechanisms for immune evasion, such as the skewing of dendritic cell activation by SOWgp and inhibition of neutrophil action by CFEM domain-containing proteins (C.-Y. Lee et al., 2015) (Bronnimann et al., 1987). These adaptations allow the fungus to persist even under active immune conditions, complicating the host response further (C.-Y. Lee et al., 2015) (Glenn et al., 2005).

**Immune Memory and Coordination:** Individuals previously exposed to Coccidioides spp. may develop a form of trained immunity, characterized by enhanced responsiveness upon re-exposure due to epigenetic modifications within innate immune cells (Miller et al., 2004) (Mendoza et al., 2015). Persistent exposure to low antigen levels may aid in fine-tuning the immune memory, bolstering defense against reinfection (Miller et al., 2004).

**Role in Immune Architecture:** Coccidioides antigens are crucial for robust T-cell priming, fostering long-term coordination among various immune cells, including CD4+ and CD8+ T cells and macrophages (Thompson, 2011) (Kokseng & Blair, 2010). This continuous engagement helps maintain tissue-specific immune readiness, particularly in the lungs and CNS, an essential factor in managing subsequent exposures or infections (Miller et al., 2004) (Mendoza et al., 2015).

**Impact of Immunosuppression:** Immunosuppressed individuals face a significantly heightened risk of severe manifestations of coccidioidomycosis, including high rates of dissemination in populations such as those with HIV or transplant recipients (Mendoza et al., 2015) (Singh et al., 1996). The interplay between immunosuppression and coccidioidal infection contributes to increased morbidity and mortality (Galgiani et al., 2000) (Singh et al., 1996).

**Prevalence and Future Considerations:** The incidence of coccidioidomycosis is on the rise, driven by factors such as climate change and population dynamics that extend beyond traditional endemic regions (C.-Y. Lee et al., 2015) (Stewart & Thompson, 2016). Continued research is imperative for the development of effective vaccines and improved diagnostic strategies to combat this increasingly pertinent public health issue (C.-Y. Lee et al., 2015) (Freedman et al., 2018).

#### Brucella Species (Bruc)

**Classification and Transmission Ecology:** Brucella spp. are classified as small, gram-negative, nonmotile, facultative intracellular coccobacilli. The most clinically significant species include Brucella melitensis, prevalent in small ruminants like sheep and goats, B. abortus, primarily found in cattle, B. suis, which affects pigs, and B. canis, associated with dogs (Ko & Splitter, 2003) (Poester et al., 2013). Notably, B. melitensis is recognized as the most virulent strain for humans (Ko & Splitter, 2003). Human infection routes primarily include the consumption of unpasteurized dairy products, direct contact with infected animal tissues or fluids, and inhalation of aerosols. Person-to-person transmission is extremely rare (Xavier et al., 2010).

**Clinical Manifestations and Latency:** Infection by Brucella can lead to acute brucellosis, characterized by symptoms such as fever, night sweats, malaise, and arthralgia. Chronic brucellosis can present with prolonged symptoms, and unresolved cases may lead to a relapsing form marked by persistent fever and fatigue (Haag et al., 2010) (Eskra et al., 2001).

**Global Distribution and Seroprevalence:** Brucellosis is endemic in various regions, including the Mediterranean, the Middle East, Central and South Asia, sub-Saharan Africa, and parts of Latin America. It is estimated that around 500,000 new cases of human brucellosis are reported annually, indicating significant public health concerns in endemic zones (Jong & Tsolis, 2011). However, the actual incidence is believed to be much higher due to underreporting, which underscores the need for enhanced surveillance and awareness (Jong & Tsolis, 2011).

**Risk Factors for Opportunism:** Occupational exposure significantly contributes to the risk of brucellosis, especially among abattoir workers, veterinarians, and shepherds. Additionally, the consumption of raw milk products is a crucial risk factor for the general population (Jiao et al., 2021). Pregnant individuals are particularly vulnerable, as brucellosis can lead to severe outcomes, including spontaneous abortion and other complications (Jiao et al., 2021) (Elfaki et al., 2015).

**Pathogenesis and Persistence Mechanisms:** Brucella species exhibit complex pathogenesis and use immune evasion strategies to establish chronic infections by inhibiting phagolysosome fusion within macrophages (Baldi & Giambartolomei, 2013) (Letesson et al., 2002). They form a Brucella-containing vacuole (BCV), which allows them to evade host defenses and provides a suitable environment for survival and replication (Amraei et al., 2020). Chronic infections are often characterized by granuloma formation and a slow replication rate, complicating treatment (Amraei et al., 2020).

**Immune Arm Dependencies:** Control of Brucella infection predominantly relies on a Th1 immune response, particularly involving interferon-gamma (IFN-γ), which activates macrophages to eliminate bacteria. While antibody formation is crucial for diagnosis, it plays a lesser role in actual immunity (Splitter & Banai, 2006) (Adetunji et al., 2020).

**Compartmental Tropism and Failure:** Brucella has a particular tropism for the reticuloendothelial system and osteoarticular tissues, often resulting in complications such as spondylitis, sacroiliitis, and peripheral arthritis (Gao & Xu, 2013). The bacteria can also invade the central nervous system and cardiovascular system, leading to serious complications (Xiao et al., 2022) (Lin et al., 2015).

**Immune Modulation and Evasion:** Brucella can modulate host immune responses by inhibiting apoptosis and downregulating antigen presentation, which helps reduce inflammatory signals and supports persistence (Qin et al., 2024) (Yi, 2019). These mechanisms contribute to chronicity and complicate treatment efforts, allowing Brucella to thrive in a less inflammatory microenvironment (Yi, 2019) (João Marcelo Azevedo de Paula & Megid, 2013).

**Immune Memory and Coordination:** Serum IgG antibodies can remain elevated for extended periods after successful treatment. However, declining antibody titers may not reliably indicate a cure; hence, serologic monitoring should be approached cautiously (Pakzad et al., 2023) (Yang et al., 2020).

**Role in Immune Architecture:** Unlike many pathogens, Brucella exploits tolerogenic niches devoid of robust systemic immune responses, which aids in its long-term survival within the host (Yang et al., 2020).

**Impact of Immunosuppression:** In immunocompromised patients, brucellosis can manifest similarly to immunocompetent individuals; however, these patients may face higher rates of relapses and complications like neurobrucellosis, emphasizing the need for vigilant monitoring.

**Prevalence and Future Considerations:** The ongoing under-recognition of human brucellosis in endemic regions, coupled with the absence of an effective human vaccine, presents substantial challenges. Public health priorities must shift towards aggressive control measures, improved diagnostics, and the development of more effective treatment regimens.

### Coxiella burnetii (Coxiella)

**Classification and Transmission Ecology:** Coxiella burnetii is classified as a gram-negative, obligate intracellular bacterium and is recognized as the etiological agent of Q fever, characterized by its environmental resilience and high infectiousness. The primary transmission route is through inhalation of aerosolized particles from infected animals, especially livestock, or contaminated airborne dust, primarily in rural settings where livestock density is high (Celina & Černý, 2022) (Hogerwerf et al., 2012). Evidence supports the notion that wind can enhance the dissemination of C. burnetii, thereby facilitating its inhalation and potential outbreak scenarios, particularly noted in regions such as southern France (Tissot-Dupont et al., 2004). Therefore, understanding its ecological transmission dynamics is crucial for controlling its spread.

**Clinical Manifestations and Latency:** The clinical spectrum of Q fever is divided into acute and chronic manifestations. Acute Q fever commonly presents as high fever, severe fatigue, symptoms resembling pneumonia, or hepatic involvement, while chronic Q fever has serious complications such as endocarditis, vascular infections, or osteomyelitis, particularly in immunocompromised patients or those with preexisting valvular heart conditions (Roest et al., 2012) (González-Barrio et al., 2022). The latency period can extend significantly, sometimes lasting years before the onset of chronic disease, adding complexity to disease management and prevention strategies (Seo et al., 2016) (Shannon & Heinzen, 2008).

**Global Distribution and Seroprevalence:** C. burnetii exhibits a global presence, particularly in rural geographical areas with substantial livestock farming operations. The seroprevalence of Q fever varies widely, influenced by geographical factors and occupational hazards—higher seroprevalence is observed among individuals closely working with livestock, such as farmers and veterinarians (Celina & Černý, 2022) (Njeru et al., 2016) (Potter et al., 2011). Significant outbreaks have been documented in various regions, including large-scale cases in the Netherlands, indicative of the public health implications tied to farming practices and environmental conditions conducive to bacterial proliferation (Sprong et al., 2011) (González-Barrio et al., 2022).

**Risk Factors for Opportunism:** Specific risk factors significantly increase susceptibility to chronic Q fever, including occupational exposure to potentially infected animals, pregnancy, and underlying conditions such as valvular heart disease or presence of vascular prostheses (Potter et al., 2011) (González et al., 2019). Immunosuppressive therapies like TNF inhibitors and corticosteroids further increase the likelihood of developing severe manifestations such as endocarditis (Roest et al., 2012) (Voth & Heinzen, 2009). These insights underline the importance of targeted education and preventive measures for high-risk populations to mitigate the progression of the disease.

**Pathogenesis and Persistence Mechanisms:** C. burnetii pathogenesis is characterized by its ability to survive and proliferate within the hostile environment of macrophage phagolysosomes. The bacterium adapts by modulating immune response pathways and utilizing cellular mechanisms to evade host defenses, ensuring its persistence and replication inside host cells (Voth & Heinzen, 2009) (Guatteo et al., 2007). It employs a specialized type IV secretion system that allows for effective intracellular manipulation and establishment of replication niches, facilitating its obligate intracellular lifestyle (Shannon & Heinzen, 2008) (Burette & Bonazzi, 2020).

**Immune Arm Dependencies:** The host immune defense against C. burnetii relies heavily on macrophage activation, with TNF-α and IFN-γ playing pivotal roles in controlling intracellular replication, emphasizing the necessity for robust cell-mediated immunity for effective pathogen clearance (Voth & Heinzen, 2009) (Guatteo et al., 2007). Antibody-driven mechanisms alone are insufficient for protection, highlighting the integral contributions of T-cell responses to infection resolution (Chisu et al., 2021) (Howard & Omsland, 2020).

**Compartmental Tropism and Failure:** During acute infections, C. burnetii predominantly affects the lungs and liver. In chronic cases, its tropism shifts towards the endocardium and vascular endothelium, with complications affecting the central nervous system and skeletal structures in immunocompromised patients or those with specific risk profiles) (Roest et al., 2012) (Guatteo et al., 2007) (Shannon & Heinzen, 2008). This compartmentality highlights varying clinical manifestations associated with different phases of infection and necessitates an approach that considers the specific tissues involved.

**Immune Modulation and Evasion:** The bacterium employs mechanisms to evade immune responses, such as antigenic phase variation, particularly between phase I and II lipopolysaccharides, which affects recognition by the host immune system, allowing for chronic inflammation without robust immunological memory formation (Seo et al., 2016) (Shannon & Heinzen, 2008). This evasive strategy enables C. burnetii to persist within host-endothelial cells, underscoring the need for ongoing research into its adaptive defenses (Njeru et al., 2016) (Hogerwerf et al., 2012).

**Immune Memory and Coordination:** Despite its capacity to provoke antibody responses, C. burnetii does not consistently lead to durable immunological memory across all hosts. Studies indicate that protective immunity is predominantly driven by robust TNF and IFN-γ responses rather than purely adaptive memory, suggesting intrinsic limitations in host responses that may contribute to chronic disease outcomes (Voth & Heinzen, 2009) (González-Barrio et al., 2022).
**Role in Immune Architecture:** Within the context of the human immune system, C. burnetii can be categorized as a keystone pathogen, revealing immune thresholds within macrophage and TNF signaling pathways. It fails to maintain persistent antigenic engagement (Njeru et al., 2016) (González-Barrio et al., 2022). Chronic disease represents a failure of innate immune coordination rather than an overt dysregulation of memory, highlighting the pathological complexities of C. burnetii infections.
**Impact of Immunosuppression:** There is an increased risk of chronic Q fever among individuals on immunosuppressive therapies. Such treatments compromise the integrity of TNF-mediated immune containment, allowing C. burnetii to bypass macrophage-mediated host defenses, leading to enhanced replication and severe disease manifestations (Potter et al., 2011) (González et al., 2019). Awareness within the clinical community regarding these risks is essential for patient management and intervention.
**Prevalence and Future Considerations:** Q fever remains an under-reported zoonosis in many regions worldwide, exacerbated by the rising prevalence of immunosuppressive therapies (Hogerwerf et al., 2012) (Potter et al., 2011). Future research initiatives should prioritize identifying immune response thresholds conducive to effective control, improving screening modalities within at-risk cohorts, and exploring vaccine options tailored to high-exposure populations (Roest et al., 2012) (Sprong et al., 2011).

#### Merkel Cell Polyomavirus (MCPyV)

**Classification and Transmission Ecology:** Merkel cell polyomavirus (MCPyV) is classified as a non-enveloped, double-stranded DNA virus belonging to the Polyomaviridae family. Discovered in 2008, it is significantly associated with Merkel cell carcinoma (MCC), a neuroendocrine malignancy of the skin (Coursaget et al., 2013) (Shailender Bhatia et al., 2011). MCPyV is considered human-adapted and is prevalent in the skin virome, typically acquired during childhood via skin contact or respiratory secretions (Liu et al., 2016). Evidence suggests that there is no known animal reservoir, thereby supporting its classification strictly as a human pathogen (Coursaget et al., 2013). As a prevalent virus within the general population, MCPyV can be detected asymptomatically in healthy adults who shed it from the skin (Liu et al., 2016).
**Clinical Manifestations and Latency:** MCPyV generally remains asymptomatic in the majority of the population, but it can contribute to oncogenesis in certain individuals by integrating into the host genome and perpetuating the expression of its viral oncoproteins (Chang & Moore, 2012) (Kuwamoto et al., 2011). This integration has been demonstrated to be a crucial step in the pathogenesis of MCC, where MCPyV is present in approximately 80% of tumors (Chang & Moore, 2012) (Cimino et al., 2014). The oncogenic potential of MCPyV is primarily linked to the continuous expression of the large T antigen, which is necessary for tumor cell viability (Chang & Moore, 2012).

**Global Distribution and Seroprevalence:** Seroprevalence studies have shown that MCPyV infection rises significantly with age, ranging from approximately 40% in children to over 80% in older adults, indicating early and consistent exposure throughout life (Liu et al., 2016) (Harms et al., 2016). In North America, up to 80% of MCC cases are found to be positive for MCPyV, while lower percentages are observed in Australia and other regions with high ultraviolet (UV) exposure (Santos-Juanes et al., 2015). This geographical variance highlights the influence of environmental factors on virus prevalence and associated tumor pathology (Santos-Juanes et al., 2015).

**Risk Factors for Opportunism:** The incidence of MCC is markedly elevated in individuals with compromised T cell immunity, such as solid organ transplant recipients, individuals living with HIV, and those suffering from chronic lymphocytic leukemia (Harms et al., 2016) (Cimino et al., 2014). In these populations, the immunosuppressive state significantly contributes to the risk and progression of MCC, as the innate immune response fails to effectively control MCPyV (Liu et al., 2016).

**Pathogenesis and Persistence Mechanisms:** MCPyV persists within the skin microbiome, yet its oncogenic transformation necessitates clonal integration into the host genome and the continual expression of its oncoproteins, particularly the large T and small T antigens (Chang & Moore, 2012). Mutations in the large T antigen are often truncating, which negates its capacity for viral replication while maintaining its tumorigenic functions, thereby instigating uncontrolled cell proliferation (Kuwamoto et al., 2011).

**Immune Arm Dependencies:** T cell-mediated immunity appears instrumental in controlling MCPyV infections. However, immunosuppression, particularly impacts on T cell function, predisposes individuals to the development of MCC (Harms et al., 2016). Virus-positive tumors may present viral antigens that trigger immune responses, yet many tumors exploit evasion tactics to escape immune clearance (Liu et al., 2016).

**Compartmental Tropism and Failure:** MCPyV DNA has been detected across various anatomic sites, highlighting its distribution within the skin virome (Liu et al., 2016). Virus-positive tumors often arise in UV-exposed areas, such as the face and extremities, although atypical presentations may occur in immunocompromised individuals (Santos-Juanes et al., 2015). This demonstrates the adaptability and persistence of MCPyV in diverse skin environments.

**Immune Modulation and Evasion:** MCPyV has developed mechanisms to evade immune detection, particularly by modulating the expression of its large T antigen to retain critical interactions with the retinoblastoma protein while circumventing signals necessary for viral replication (Cimino et al., 2014). Notably, virus-negative MCCs also exist and are characterized by UV-induced mutations and divergence in mutation burdens, suggesting distinct pathways of oncogenesis (Nakamura et al., 2010).

**Immune Memory and Coordination:** Both humoral and cellular immune responses to MCPyV are common in the population. Notably, antibody titers against MCPyV oncoproteins correlate with tumor burden and may decline following effective treatment – with rising titers during monitoring serving as potential early indicators of recurrence (Harms et al., 2016). This dynamic demonstrates the intricate balance of immune memory in the context of persistent viral infections.

**Role in Immune Architecture:** Unlike other oncogenic viruses, MCPyV does not seem to foster long-term immune coordination across different compartments; rather, its impact remains localized to contributing towards oncogenic processes without shaping broader systemic immune responses (Cimino et al., 2014).

**Impact of Immunosuppression:** Chronic T-cell suppression, notably via agents like calcineurin inhibitors and azathioprine, significantly increases MCC risk and negatively influences outcomes. Immunosuppressed patients are frequently diagnosed with advanced-stage disease, correlating with higher rates of recurrence (Nakamura et al., 2010) (Santos-Juanes et al., 2015).

**Prevalence and Future Considerations:** While generally benign, MCPyV is largely ubiquitous and only oncogenic in selected cases. The advent of MCPyV serology shows promise for monitoring and potentially treating malignancy associated with this virus (Harms et al., 2016). Ongoing investigations are scrutinizing circulating tumor DNA (ctDNA) and novel immune checkpoint inhibitors as additional avenues for therapeutic intervention and surveillance (Bofill-Mas et al., 2010) (Santos-Juanes et al., 2015).

#### Parvovirus B19

**Classification and Transmission Ecology:** Parvovirus B19, a member of the Erythroparvovirus genus within the family Parvoviridae, is a small, non-enveloped, single-stranded DNA virus recognized for its pathogenicity in humans. It is classified uniquely due to its lack of known animal reservoirs, being exclusively human-adapted (Qiu et al., 2017) (Corcoran & Doyle, 2004). The primary transmission routes for Parvovirus B19 include respiratory droplets, particularly prevalent among children, as well as hematogenous routes, such as blood transfusions, and vertical transmission from mother to fetus during pregnancy (Qiu et al., 2017) (Heegaard & Brown, 2002). The history of B19's discovery further highlights its significance; Cossart et al.'s initial identification in 1975 paved the way for extensive research into its epidemiology and impacts on human health (Qiu et al., 2017) (Corcoran & Doyle, 2004).

**Clinical Manifestations and Latency:** In immunocompetent individuals, Parvovirus B19 infection often presents asymptomatically or manifests as erythema infectiosum or arthropathy, with approximately 8% of teenagers and up to 60% of affected adults developing arthralgias (Corcoran & Doyle, 2004) (Albayrak et al., 2019). Among patients with hemoglobinopathies, notably sickle cell disease, infection can precipitate a transient aplastic crisis, due to the virus's targeting of erythroid progenitor cells (Qiu et al., 2017) (Raju et al., 2013). Conversely, in immunocompromised hosts, the virus can lead to chronic anemia due to persistent infection, with documented cases of the virus's DNA remaining in tissues for years, particularly in bone marrow (Schenk et al., 2009) (Erol & Alehan, 2006).

**Global Distribution and Seroprevalence:** Parvovirus B19 demonstrates a global distribution, with seroprevalence rates increasing substantially with age—around 30–60% in adult populations (Hino et al., 2000) (Pattabiraman et al., 2022). Seasonal outbreaks are common in temperate climates, and recent data indicated renewed spikes in cases within the United States and Europe as of 2024 (Hino et al., 2000) (Pattabiraman et al., 2022). This ubiquity emphasizes the virus's significant public health impact.

**Risk Factors for Opportunism:** Individuals with underlying hemolytic anemias and those who are immunosuppressed are at a heightened risk for severe manifestations of Parvovirus B19 infection (Qiu et al., 2017) (Broliden, 2001). Pregnancy poses additional risks, as the infection can lead to fetal complications such as hydrops fetalis and spontaneous abortions, warranting careful monitoring in affected populations (Enders et al., 2004) (Jong et al., 2006).

**Pathogenesis and Persistence Mechanisms:** The pathogenesis of Parvovirus B19 includes its binding to the P antigen on erythroid progenitor cells, leading to cell apoptosis and inhibition of normal erythropoiesis (Qiu et al., 2017) (Chisaka et al., 2003). Notably, the non-structural protein NS1 plays a crucial role in inducing apoptosis and promoting inflammatory cytokine expression, further contributing to cytotoxicity (Corcoran & Doyle, 2004) (O'Sullivan et al., 1994). The virus's ability to persist in solid tissues, including bone marrow, has serious implications for individuals with compromised immune systems, often leading to a chronic state of infection and associated anemia (Schenk et al., 2009) (Boer et al., 2014).

**Immune Arm Dependencies:** The clearance of Parvovirus B19 relies heavily on the host's immune response, specifically the generation of neutralizing IgG antibodies targeting the VP1u region (Qiu et al., 2017) (Heegaard & Brown, 2002). Immunocompromised patients, who cannot mount an effective immune response, frequently develop chronic anemia requiring interventions such as intravenous immunoglobulin (IVIG) treatment (Dembinski, 2003).

**Compartmental Tropism and Failure:** Primary replication of the virus occurs in erythroid progenitor cells found in the bone marrow and fetal liver, while secondary nonproductive infections can manifest in other tissues such as the myocardium and placenta (Schenk et al., 2009) (Erol & Alehan, 2006). The persistence of B19 DNA in these organs, even in the absence of active infection, presents challenges in clinical management and diagnosis (Pattabiraman et al., 2022) (Chisaka et al., 2003).

**Immune Modulation and Evasion:** The NS1 protein of Parvovirus B19 is known to transactivate pro-inflammatory cytokines, including TNF-α and IL-6, which can facilitate immune evasion (Qiu et al., 2017) (Pattabiraman et al., 2022). The VP1u protein further aids in this process by exhibiting phospholipase A2 activity, allowing the virus to survive and evade detection by the immune system (Corcoran & Doyle, 2004) (Enders et al., 2004).

**Immune Memory and Coordination:** Long-term immunity is primarily mediated by IgG antibodies, which confer protection against subsequent infections (Qiu et al., 2017) (Corcoran & Doyle, 2004). Persistent CD8+ T cell responses have been observed even after viral clearance, suggesting a robust immune memory following infection (Raju et al., 2013) (Dembinski, 2003).

**Role in Immune Architecture:** Although Parvovirus B19 is not classified as a keystone virus, its specific tropism for erythroid progenitor cells and significant reliance on the immune response render it a sensitive marker for assessing host hematologic health and potential immune deficiencies (Broliden, 2001) (Pattabiraman et al., 2022). This aspect is particularly relevant in high-risk populations, such as pregnant women and those with HIV or following organ transplants, where vulnerabilities in immune regulation can be revealed (Chisaka et al., 2003) (Knöll et al., 2002).

**Impact of Immunosuppression:** In populations undergoing organ transplantation, particularly solid organ recipients, Parvovirus B19 can instigate persistent viremia and anemia, resulting in significant morbidity (Broliden, 2001) (Lundqvist et al., 2005). Effective treatment options such as IVIG are often employed, though recurrent anemia may necessitate additional therapies, including antiviral agents like foscarnet in dire cases (Qiu et al., 2017) (Dembinski, 2003).

**Prevalence and Future Considerations:** Despite its relatively high seroprevalence and significant clinical impacts, Parvovirus B19 remains underrecognized in cases involving fetal loss and post-transplant anemia (Raju et al., 2013) (Corcoran & Doyle, 2004). Previous vaccine trials utilizing VP1/VP2 virus-like particles have shown promise but faced termination due to adverse events (Jordan et al., 2001). Consequently, innovative vaccine strategies, like a yeast-expressed VLP vaccine combined with inactivated phospholipase A2, are being investigated and may provide future avenues for prevention (Knöll et al., 2002) (Tu et al., 2015).

#### Nocardia species (Nocardia)

**Classification and Transmission Ecology:** Nocardia species are classified within the phylum Actinobacteria and are recognized as aerobic, filamentous, gram-positive bacteria. More than 50 species of Nocardia have been identified through both classical phenotypic methods and advanced molecular techniques, which continue to evolve (Brown-Elliott et al., 2006) (Wilson, 2012). These microorganisms predominantly inhabit soil, decaying organic matter, and water, thus making them environmental organisms. As opportunistic pathogens, Nocardia infections typically occur following exposure via inhalation of contaminated aerosols or direct cutaneous inoculation, especially in individuals with compromised immune systems (Wilson, 2012) (Viscuse & Mohabbat, 2019).

**Clinical Manifestations and Latency:** Pulmonary nocardiosis is the most common clinical manifestation, characterized by symptoms such as fever, cough, chest pain, and hemoptysis; radiological findings may show nodules, cavitation, or consolidation (Wilson, 2012) (Viscuse & Mohabbat, 2019). Extrapulmonary dissemination is noted in up to one-third of cases, typically affecting the central nervous system (CNS) and skin, especially in severely immunocompromised patients (Wilson, 2012) (Viscuse & Mohabbat, 2019). Unlike many viral infections, Nocardia does not induce latent infections or persist chronically within hosts, thereby distinguishing its pathogenesis from that of herpesviruses (King et al., 2008) (Minero et al., 2009).

**Global Distribution and Seroprevalence:** While Nocardia species are distributed worldwide, the incidence of human nocardiosis appears to be low and is likely underreported, primarily due to difficulties in clinical recognition and diagnosis (Yu et al., 2016) (Wang et al., 2023). Currently, there is no dedicated serological test available for the screening or assessment of seroprevalence; colonization of Nocardia is infrequent among immunocompetent individuals (Yu et al., 2016).

**Risk Factors for Opportunism:** Individuals with compromised cell-mediated immunity are at the highest risk for nocardiosis. Specific risk factors include advanced HIV/AIDS, prolonged corticosteroid therapy, hematologic malignancies, solid organ transplantation, and the use of TNF inhibitors, all of which can lead to a significant reduction in the host's ability to fight off infectious agents like Nocardia (Sakyi et al., 2018) (Onaiwu et al., 2017). In fact, studies indicate that a significant majority of Nocardia infections occur in patients with these immunocompromising conditions (Sakyi et al., 2018) (Onaiwu et al., 2017).

**Pathogenesis and Persistence Mechanisms:** Nocardia is characterized as a facultative intracellular pathogen capable of surviving within macrophages by preventing phagosome-lysosome fusion and resisting oxidative stress mechanisms, including the production of catalase and superoxide dismutase (Brown-Elliott et al., 2006). Despite its ability to persist locally during immune suppression, Nocardia does not establish latency or long-term tissue residency, factors that are crucial for understanding its pathogenicity (Brown-Elliott et al., 2006) (Cooper et al., 2014).

**Immune Arm Dependencies:** The immune response to Nocardia relies heavily on CD4+ T cells, which are vital for macrophage activation and the containment of the infection. Studies have shown that depletion of CD4+ T cells or disruption of the IL-12/IFN-γ signaling pathways leads to uncontrolled nocardiosis, underscoring the significance of the Th1 immune response in managing this infection (King et al., 2008) (Wang et al., 2024). While CD8+ T cells and antibodies contribute to the immune response, their role appears to be secondary in comparison to the critical importance of CD4+ T cells (King et al., 2008).

**Compartmental Tropism and Failure:** Pulmonary disease is the primary manifestation; however, CNS involvement is often more severe and associated with increased mortality, particularly when diagnosis is delayed in immunocompromised patients (Wilson, 2012) (Viscuse & Mohabbat, 2019). Cutaneous infections can occur through direct inoculation, particularly in patients who have undergone solid organ transplantation or are receiving high doses of immunosuppressive therapy (Wilson, 2012) (Cooper et al., 2014).

**Immune Modulation and Evasion:** Nocardia does not produce known immune evasion genes; instead, it circumvents phagocytic killing through enzymatic defenses and inhibiting phagolysosomal fusion. The absence of mechanisms to downregulate antigen presentation or interfere with T-cell signaling suggests a unique strategy focused primarily on resistance to immediate immune challenges rather than long-term immune evasion (Brown-Elliott et al., 2006) (Viscuse & Mohabbat, 2019).

**Immune Memory and Coordination:** Nocardia does not incite durable adaptive immune memory as seen with viral infections; however, prior exposure may enhance the responsiveness of local macrophages to subsequent infections. In specific immunological contexts, allergic bronchopulmonary aspergillosis (ABPA)-like hypersensitivity reactions indicate that some form of T-cell memory might develop, albeit in a more limited capacity compared to typical viral infections (Wang et al., 2024) (Wang et al., 2022).

**Role in Immune Architecture:** Nocardia serves as a model organism to study the interplay between macrophages and CD4+ T-cell coordination during an infection. Its unique intracellular lifestyle emphasizes the need for robust Th1 activation for effective containment of pathogens, thereby highlighting the complexities of immune architecture during infection (Brown-Elliott et al., 2006) (Viscuse & Mohabbat, 2019).

**Impact of Immunosuppression:** Immunosuppression—due to conditions such as HIV/AIDS, corticosteroid treatment, or immunosuppressive drugs—significantly increases the risk of disseminated nocardiosis, particularly involving the CNS, and often leads to poor outcomes, including death (Sakyi et al., 2018) (Onaiwu et al., 2017). This underscores the critical role of intact immune function in managing infections caused by Nocardia.

**Prevalence and Future Considerations:** Nocardia remains underrecognized, especially with increasing use of immunosuppressive therapies. Future directions in research should focus on improving diagnostic methods such as molecular diagnostics and rapid culture techniques, which can enhance detection rates. There is a pressing need to define immune thresholds for preventing compartmental failure and to explore potential immunoprophylactic strategies for high-risk populations (Sakyi et al., 2018) (Cooper et al., 2014).

#### Burkholderia species (Burkholderia)

**Classification and Transmission Ecology:** The Burkholderia cepacia complex (BCC) includes a group of genetically related, gram-negative bacteria that are primarily environmental pathogens. BCC predominantly infects individuals with cystic fibrosis (CF) or chronic granulomatous disease (CGD), often through inhalation or direct contact with contaminated surfaces (Chaparro et al., 2001) (LiPuma, 2005). Transmission routes can include percutaneous inoculation or exposure to contaminated environments, rather than direct consumption of resources like soil or water (Boussaud et al., 2008). Notably, Burkholderia pseudomallei, the causative agent of melioidosis, is endemic in specific regions, particularly known for its severe outbreaks during monsoonal downpours (Boussaud et al., 2008).

**Clinical Manifestations and Latency:** Patients with CF typically experience chronic respiratory infections characterized by persistent airway colonization and deterioration of lung function. In contrast, individuals with CGD may suffer from acute life-threatening pneumonia when infected with BCC. Strains such as B. cenocepacia are associated with significantly high mortality rates following lung transplantation (Aris et al., 2001) (Nash et al., 2010). B. pseudomallei also has a documented capacity for latency, with reactivation occurring years post-exposure, highlighting its complicated interaction with the host immune system (Caraher et al., 2007) (LiPuma, 2001).

**Global Distribution and Seroprevalence:** Melioidosis, caused by B. pseudomallei, is mainly recognized in Southeast Asia and northern Australia, with increasing reports in Africa and the Americas linked to historical trade routes that may have facilitated its dispersion (Coenye, 2010) (Langley et al., 2003). Recent studies indicate that B. pseudomallei has also been identified in Puerto Rico and along the U.S. Gulf Coast, suggesting a need for enhanced surveillance to address its expanding geographical presence (Nash et al., 2010) (Daccò et al., 2019).

**Risk Factors for Opportunism:** Individuals with chronic granulomatous disease are particularly vulnerable to BCC infections due to their impaired immune responses, contributing to the severity of the disease (Hauser & Orsini, 2015). Other notable risk factors for melioidosis include diabetes, chronic renal failure, and hazardous alcohol use (Langley et al., 2003) (Soyza & Corris, 2003). This underscores the significance of continuous clinical vigilance in at-risk populations (Mitchell & Glanville, 2021).

**Pathogenesis and Persistence Mechanisms:** B. pseudomallei displays remarkable adaptability as a facultative intracellular pathogen, able to survive within macrophages and evade immune mechanisms by obstructing lysosomal fusion (Ramos et al., 2016). Concurrently, BCC's biofilm-forming ability in the pulmonary environment contributes to chronic infections in CF patients, complicating treatment due to antibiotic resistance (Coenye, 2010) (Canning et al., 2024).

**Immune Arm Dependencies:** Effective clearance of both BCC and B. pseudomallei relies heavily on the functionality of phagocyte NADPH oxidase, with individuals having CGD showing significant susceptibility due to the lack of reactive oxygen species crucial for combating these pathogens (Vermis et al., 2002) (George et al., 2006). Although adaptive immunity plays a lesser role, T cell responses have been observed to aid in managing intracellular infections caused by B. pseudomallei, indicating a complex immune landscape (Goodlet et al., 2019) (Carraro et al., 2018).

**Compartmental Tropism and Failure:** BCC primarily affects respiratory health but can also lead to systemic infections in cases involving CGD or lung transplants. B. pseudomallei presents a broader spectrum of potential infections, impacting various organs, including the central nervous system (Chaparro et al., 2001) (Ho et al., 2019).

**Immune Modulation and Evasion:** B. pseudomallei possesses mechanisms to evade host defenses, such as downregulating toll-like receptor (TLR) signaling and forming multinucleated giant cells that escape detection (Blower & Keyt, 2016) (Vermis, 2002). Similarly, BCC's capabilities of quorum sensing and biofilm formation enhance its resilience against host immune strategies (Vazirani et al., 2021) (Bogaart & Manuel, 2022).

**Immune Memory and Coordination:** Both BCC and B. pseudomallei do not typically induce a robust, enduring protective immune response, with recurrent infections and disease reactivation noted, suggesting an ability to circumvent traditional markers of adaptive immunity and demonstrating limited immune imprinting (Vazirani et al., 2021) (Bogaart & Manuel, 2022).

**Role in Immune Architecture:** Rather than being mere organizers of immune responses, Burkholderia species expose vulnerabilities in the innate immune system. Their presence can serve as diagnostic markers for assessing immune competence rather than establishing hierarchies in adaptive immunity (Vazirani et al., 2021).

**Impact of Immunosuppression:** In lung transplant recipients, B. cenocepacia infection correlates with poor postoperative outcomes, where the risk of pneumonia and sepsis from B. pseudomallei is particularly pronounced among immunocompromised individuals. This necessitates stringent preventive measures in high-risk settings (Cho et al., 2013) (Vazirani et al., 2021).

**Prevalence and Future Considerations:** The inherent multidrug resistance of BCC, particularly for B. cenocepacia, complicates treatment and management in CF patients. Additionally, the often underdiagnosed nature of melioidosis in various regions highlights an urgent need for improved diagnostic strategies and regional surveillance to effectively address these infections (Vazirani et al., 2021) (Bogaart & Manuel, 2022).

#### Legionella pneumophila (Legionella)

**Classification and Transmission Ecology:** Legionella pneumophila, an aerobic, gram-negative bacillus, is part of the genus Legionella, which was established following an outbreak of pneumonia among American Legion members in 1976 (Fields et al., 2002). This bacterium thrives in warm aquatic environments, such as cooling towers and hot water systems, where it can be aerosolized into tiny droplets that are inhaled by humans. Transmission occurs not through direct person-to-person contact but primarily through inhalation of these contaminated aerosols (Fields et al., 2002) (Yu et al., 2002) (Moran-Gilad et al., 2014). L. pneumophila is frequently associated with intracellular parasitism within protozoa in freshwater ecosystems, which facilitate its survival and proliferation in aquatic environments (Yu et al., 2002) (Atlas, 1999).

**Clinical Manifestations and Latency:** The most significant clinical disease caused by L. pneumophila is Legionnaires' disease, a severe form of pneumonia that is often accompanied by multisystem complications, including gastrointestinal and neurological symptoms (Fields et al., 2002) (Niu & Zhao, 2022). The onset of this disease is typically acute, reflecting a rapid progression to critical illness without a noted latency period (Fields et al., 2002) (Moran-Gilad et al., 2014). In immunocompromised individuals, the disease may present variably, but the acute nature remains a defining characteristic of L. pneumophila infections (Niu & Zhao, 2022) (Kümpers et al., 2008).

**Global Distribution and Seroprevalence:** L. pneumophila is commonly found in freshwater sources around the globe, particularly thriving in environments like warm water supplies and complex systems such as plumbing (Fields et al., 2002) (Yu et al., 2002). Among its serogroups, serogroup 1 is responsible for the majority of cases of human legionellosis, accounting for approximately 80-85% of infections (Yu et al., 2002) (Moran-Gilad et al., 2014). Despite the widespread exposure to this pathogen, seroprevalence studies indicate that the population-level immunity against L. pneumophila is relatively low, which aligns with its episodic nature of outbreaks (Fields et al., 2002) (Yu et al., 2002).

**Risk Factors for Opportunism:** Individuals at the highest risk for developing infections due to L. pneumophila include the elderly, smokers, and those with chronic lung diseases or undergoing immunosuppressive therapies (Fields et al., 2002) (Yu et al., 2002) (Kümpers et al., 2008). These treatments, particularly with glucocorticoids and TNF-α blockers, significantly impair essential immune functions such as macrophage activation, which is crucial for controlling Legionella infections (Yu et al., 2002) (Zhu et al., 2024). Such demographic vulnerabilities highlight the need for specific preventative measures in high-risk groups (Fields et al., 2002) (Kümpers et al., 2008).

**Pathogenesis and Persistence Mechanisms:** L. pneumophila has evolved sophisticated mechanisms for survival within host alveolar macrophages. It utilizes a type IV secretion system to evade phagolysosomal fusion, thereby avoiding destruction and creating a niche for replication (Fields et al., 2002) (Yu et al., 2002). The bacterium's ability to manipulate host cellular processes and prevent oxidative killing is closely linked to its evolutionary adaptation to thrive not only in human hosts but also in protozoan hosts, which confer important survival traits (Yu et al., 2002) (Atlas, 1999).

**Immune Arm Dependencies:** The effectiveness of immune responses to L. pneumophila heavily relies on the innate immune system, particularly through the activation of macrophages via IFN-γ from natural killer (NK) and CD4+ T-cells (Fields et al., 2002) (Yu et al., 2002). Furthermore, TNF-α is crucial in managing the intracellular replication of the bacterium, emphasizing the role of a robust cellular immune response in the body's defense against this pathogen (Fields et al., 2002) (Yu et al., 2002). Interestingly, antibodies generated against L. pneumophila play a lesser role in offering significant protection compared to these cellular pathways (Fields et al., 2002) (Yu et al., 2002).

**Compartmental Tropism and Failure:** While the primary infection site for L. pneumophila is the lungs, there is potential for extrapulmonary dissemination, especially in immunocompromised patients. This systemic involvement may lead to organ dysfunction, including hepatic and renal disturbances, stemming from an overwhelming systemic inflammatory response rather than direct bacterial invasion (Fields et al., 2002) (Niu & Zhao, 2022) (Kümpers et al., 2008).

**Immune Modulation and Evasion:** Legionella employs over 300 effector proteins through the Dot/Icm type IV secretion system, facilitating evasion of host defenses by inhibiting key processes such as phagosome maturation and apoptosis in macrophages (Fields et al., 2002) (Atlas, 1999). This complex machinery underscores the bacterium's adeptness at manipulating host immune responses, allowing for its persistence within immune cells (Yu et al., 2002) (Atlas, 1999).

**Immune Memory and Coordination:** Host immunity following infection by L. pneumophila tends to be serogroup-specific and short-lived, indicating that while some adaptive responses are activated, they do not confer long-term protection across different serogroups (Fields et al., 2002) (Yu et al., 2002). The pathogen primarily acts as an acute stressor to innate immune pathways rather than eliciting a robust immune recalibration (Fields et al., 2002) (Yu et al., 2002).
**Role in Immune Architecture:** Although L. pneumophila does not fit neatly into the category of classic keystone pathogens, it does expose deficiencies in macrophage activation and cell-mediated immunity, thereby serving as a stress-test for these immune mechanisms (Yu et al., 2002) (Atlas, 1999). Its intracellular living strategy positions it as an intermediate category of pathogens that challenge the immune system to adapt (Fields et al., 2002) (Atlas, 1999).
**Impact of Immunosuppression:** Immunosuppressive therapies markedly increase susceptibility to L. pneumophila, leading to a higher mortality rate from pneumonia caused by this pathogen, particularly in transplant recipients and individuals with hematological malignancies (Zhu et al., 2024) (Kümpers et al., 2008). The association of high mortality rates underscores the vulnerability of these populations to opportunistic infections such as those caused by L. pneumophila (Fields et al., 2002) (Yu et al., 2002).
**Prevalence and Future Considerations:** While outbreaks of Legionella are relatively infrequent, L. pneumophila remains a significant concern for immunocompromised populations, as well as individuals exposed to contaminated water systems (Fields et al., 2002) (Niu & Zhao, 2022). A comprehensive understanding of IFN-γ and TNF-α pathways may inform the development of future host-directed therapies, while better surveillance and enhanced management of water systems are crucial strategies for preventing outbreaks (Fields et al., 2002) (Yu et al., 2002) (Kümpers et al., 2008).

#### Balamuthia mandrillaris (Balamuthia)

**Classification and Transmission Ecology:** Balamuthia mandrillaris is classified within the genus Balamuthia, family Acanthamoebidae, and is recognized as a free-living amoeba found primarily in soil and freshwater environments worldwide. It is well-documented to be an opportunistic pathogen, capable of causing granulomatous amoebic encephalitis (GAE), a severe central nervous system infection that is often fatal (Visvesvara et al., 2007) (Matin et al., 2008). Transmission of B. mandrillaris typically occurs via inhalation of aerosolized particles or through skin contact with contaminated soil or water (Visvesvara et al., 2007) (Paudel et al., 2021). Its ecological presence has led to environmental isolation from various habitats, underscoring its ubiquitous distribution (Niyyati et al., 2009).
**Clinical Manifestations and Latency:** The clinical manifestations of GAE include a protracted and insidious onset of neurological symptoms such as headache, confusion, seizures, and focal neurological deficits (Bakardjiev et al., 2003) (Matin et al., 2008). These symptoms often appear weeks to even months after exposure to B. mandrillaris, complicating diagnosis (Mani & Hudgins, 2021). Latency has been postulated in some cases, but the evidence remains insufficient (Yohannan & Feldman, 2019). Additionally, initial skin lesions have been noted in some patients, often preceding the onset of CNS symptoms, which may reflect an earlier stage of infection (Mani & Hudgins, 2021) (Matin et al., 2008).

**Global Distribution and Seroprevalence:** Balamuthia mandrillaris is reported globally, with notable occurrences in the Americas, particularly in the United States (Zheng et al., 2024) (Matin et al., 2008). Environmental studies have isolated the amoeba from various soil and freshwater samples, underscoring its widespread presence (Niyyati et al., 2009) (Paudel et al., 2021). Furthermore, seroprevalence studies indicate that exposure to Balamuthia may be far more common than reported clinical cases suggest, indicating a potential underdiagnosis of subclinical or mild forms of infection (Visvesvara et al., 2007) (Zheng et al., 2024) (Silva-Vergara et al., 2007).
**Risk Factors for Opportunism:** Individuals who are immunocompromised due to conditions such as corticosteroid therapy, HIV/AIDS, or organ transplantation represent significant risk groups for GAE; however, cases have also been documented in immunocompetent individuals exposed to high doses of the amoeba (Bakardjiev et al., 2003) (Mani & Hudgins, 2021). Epidemiological analyses suggest that environmental exposure, particularly through activities like gardening or wounds from soil (Paudel et al., 2021) (Mani & Hudgins, 2021) (Niyyati et al., 2009), plays a critical role in infection among both immunocompromised and healthy populations.
**Pathogenesis and Persistence Mechanisms:** Upon entry through the skin or respiratory system, Balamuthia can disseminate hematogenously to the central nervous system, instigating granulomatous inflammation and necrotizing vasculitis, primarily affecting neuronal tissues (Visvesvara et al., 2007) (Bakardjiev et al., 2003) (Matin et al., 2008). Unlike other free-living amoebae such as Acanthamoeba, B. mandrillaris does not thrive on standard culture media, which complicates diagnostic efforts and our understanding of its persistence mechanisms in host organisms and environments (Visvesvara et al., 2007) (Matin et al., 2008).
**Immune Arm Dependencies:** Control of B. mandrillaris infection heavily relies on the host's intact immune system, particularly functions mediated by macrophages and CD4+ T cells. Recent findings posit that neutrophils play a secondary but still important role (Visvesvara et al., 2007) (Bakardjiev et al., 2003). Patients with compromised cellular immunity exhibit markedly worse outcomes, indicating the critical nature of cell-mediated responses in the pathogenesis of GAE (Visvesvara et al., 2007) (Schuster et al., 2006).
**Compartmental Tropism and Failure:** Balamuthia's primary site of invasion and pathology is the central nervous system, specifically targeting the frontal and temporal lobes. Moreover, the amoeba has been associated with lesions on the skin and in the lungs, suggesting a multifocal pathogenic spread before it reaches the CNS (Bakardjiev et al., 2003) (Matin et al., 2008). Such findings elevate concerns about the systemic effects of B. mandrillaris and the timing of clinical intervention (Mani & Hudgins, 2021) (Paudel et al., 2021).
**Immune Modulation and Evasion:** Balamuthia mandrillaris seemingly lacks classical immune evasion mechanisms, such as antigenic variation. Instead, it appears to evade clearance through the development of slow, localized inflammation that hinders immune detection and response (Visvesvara et al., 2007) (Mani & Hudgins, 2021). This adaptive progression could illuminate avenues for therapeutic strategies aimed at enhancing immune response against the pathogen (Schuster et al., 2006) (Matin et al., 2008).
**Immune Memory and Coordination:** Current evidence does not support the existence of robust adaptive immune memory following Balamuthia infection, rendering reinfection risk indeterminate (Yohannan & Feldman, 2019) (Matin et al., 2008). The immune response is typically characterized by delayed activation, lacking the organized epitope hierarchy seen with

other pathogens, as mortality rates remain high in the absence of effective responses (Mani & Hudgins, 2021) (Matin et al., 2008).

**Role in Immune Architecture:** Rather than contributing positively to immune system architecture, Balamuthia mandrillaris exploits weaknesses, particularly within the cell-mediated immunity that encompasses macrophage and T-cell coordination (Visvesvara et al., 2007) (Matin et al., 2008). This interaction underscores the need for an integrated understanding of how such pathogens could impact long-term immune health.

**Impact of Immunosuppression:** Immunosuppressive therapies, including corticosteroids and anti-humoral agents, considerably augment the risks associated with B. mandrillaris infections, often culminating in severe GAE with mortality rates approaching 100% in documented cases (Silva-Vergara et al., 2007) (Paudel et al., 2021). This highlights the infectious burden that Balamuthia poses particularly to vulnerable populations (Visvesvara et al., 2007) (Matin et al., 2008).

**Prevalence and Future Considerations:** The rarity of clinically reported Balamuthia infections belies the high fatality rate associated with GAE. Awareness of underdiagnosis is crucial, with improved diagnostics like PCR and immunohistochemistry being pivotal for managing and containing outbreaks (Visvesvara et al., 2007) (Schuster et al., 2006). Future research should pivot towards early detection methods, understanding host susceptibility, and exploring combination therapies as promising avenues to mitigate the severe impacts of this pathogen (Matin et al., 2008) (Silva-Vergara et al., 2007).

#### Schistosoma species (Schistosoma)

**Classification and Transmission Ecology:** Schistosoma species, classified under the trematode class of flatworms, are parasitic organisms that play a significant role in global public health. The primary modes of transmission involve skin penetration by cercariae, which are released from freshwater snails, the intermediate hosts (Pearce & MacDonald, 2002) (McManus et al., 2018). Major human pathogens in this genus include *S. mansoni*, *S. haematobium*, and *S. japonicum*. These adult parasites typically reside in the mesenteric venous plexus (for *S. mansoni* and *S. japonicum*) or the pelvic venous plexus (for *S. haematobium*), exposing their eggs to the host's immune system, which triggers a granulomatous inflammatory response (Pearce & MacDonald, 2002) (McManus et al., 2018) (Marascio et al., 2023). The pathophysiology associated with egg release is vital for understanding the disease's clinical impact and its resultant morbidity (Lamb et al., 2007).

**Clinical Manifestations and Latency:** Acute schistosomiasis, colloquially known as Katayama fever, can present with a constellation of symptoms, including fever, eosinophilia, rashes, and respiratory issues, particularly in individuals lacking prior exposure (McManus & Loukas, 2008) (Schwartz et al., 2014). Chronic infection can lead to severe hepatosplenic disease for *S. mansoni* and *S. japonicum*, resulting in complications such as portal hypertension. *S. haematobium* primarily affects the urinary system and is associated with hematuria, bladder wall fibrosis, and an increased risk for squamous cell carcinoma of the bladder (McManus & Loukas, 2008) (Schwartz et al., 2014). The understanding of these clinical manifestations is underscored by varied immune responses observed during infections, emphasizing the complex interplay between the parasite and host immunity (Haeberlein et al., 2017).

**Global Distribution and Seroprevalence:** The reach of schistosomiasis is extensive, infecting over 200 million individuals worldwide, predominantly in endemic regions of sub-Saharan Africa, parts of the Middle East, South America (notably Brazil), and Southeast Asia (Morales-Suárez-Varela et al., 2019) (Schwartz et al., 2014). The prevalence of schistosomiasis is inherently tied to environmental factors such as exposure to freshwater sources conducive to snail habitation and the resultant cercarial release (Pearce & MacDonald, 2002) (Wilson et al., 2006). As such, seropositivity often correlates with the incidence of contact with contaminated water bodies, highlighting the importance of environmental monitoring in controlling the disease (Schwartz et al., 2014).

**Risk Factors for Opportunism:** Individuals engaging in activities like bathing, swimming, or occupational exposure in contaminated freshwater ecosystems are at heightened risk for schistosomiasis (Morenikeji et al., 2016). Poor sanitation practices and migration to endemic areas further exacerbate vulnerability, particularly in children and previously unexposed populations who often develop symptomatic disease more readily (Maghraby et al., 2010). Historical data emphasize the socio-economic disparities that influence the dynamics of exposure and infection rates (Maghraby et al., 2010) (Colombe et al., 2018).

**Pathogenesis and Persistence Mechanisms:** The pathogenesis of schistosomiasis is notably dependent on the host immune response against the eggs laid by adult worms, prompting granulomatous reactions that lead to fibrosis in various organs, primarily the liver and urogenital tract (Duan et al., 2015). Schistosomes have evolved mechanisms to evade immune clearance, including dynamic changes in their tegument and the mimicking of host antigens, enabling them to persist in the host for extended periods (Ji et al., 2008). These immune evasion strategies are fundamental to understanding the chronic nature of the infection and its resultant pathology (Isabella Maria Monteiro de et al., 2022).

**Immune Arm Dependencies:** Infected individuals typically exhibit a Th2-skewed immune response characterized by heightened eosinophilia and elevated IgE levels. Regulatory T cells play a critical role in modulating inflammation and preventing extensive host tissue damage (Gao et al., 2015). The initial immune responses involve Th1 dominance, which later transitions to Th2 predominance, informing the chronic stages of infection (Pinho et al., 2009). Thus, the immune response to Schistosoma spp. is multifaceted and reflects a complex balance between efforts to control the parasite and the parasite's strategies to manipulate host immunity (Lei et al., 2019).

**Compartmental Tropism and Failure:** Schistosoma spp. are known for their compartmentalized residency; *S. mansoni* and *S. japonicum* predominantly localize in mesenteric veins, while *S. haematobium* targets the pelvic venous plexus. This compartmentalization is critical for understanding the distinct pathologies associated with each species, including liver fibrosis and urogenital complications (Colombe et al., 2018) (Melo et al., 2010). The localized nature of infection can lead to significant morbidity, particularly in high-burden areas, where chronic disease processes are prevalent (Colombe et al., 2018).

**Immune Modulation and Evasion:** The schistosome's capacity to evade host immunity is significantly attributable to strategies such as antigen mimicry and the alteration of surface protein expression. These tactics skew the immune response toward Th2-related pathways, effectively dampening Th1-mediated immune clearance mechanisms and promoting a persistent state of infection (Kumar et al., 2019). This immunomodulation underpins the chronic nature of schistosomiasis and complicates vaccine development (Jiang et al., 2011).

**Immune Memory and Coordination:** Long-term immune responses to Schistosoma are complex and require continuous exposure to antigens from the parasites. Despite inducing some level of immunological memory, achieving sterilizing immunity remains rare, complicating the pursuit of effective vaccines against schistosomiasis (Das et al., 2010). Vaccination strategies have faced significant hurdles primarily due to the parasite's evolved mechanisms for immune evasion and the intricate dynamics of the immune response that favor regulation over clearance (Jiang et al., 2011).

**Role in Immune Architecture:** Schistosoma spp. significantly influence the immune architecture of host organisms, often leading to systemic immune modulation. Chronic infection can result in the expansion of regulatory T cells and a generalized dampening of inflammatory reactions to co-infective pathogens or vaccines. The implications of such immune alterations extend beyond schistosomiasis, suggesting that these parasites may influence broader public health immunological outcomes in endemic regions.

**Impact of Immunosuppression:** Immunosuppressive conditions like HIV/AIDS alter the dynamics of schistosomiasis, notably through the reduction of CD4+ T cell responses, ultimately increasing parasite loads and exacerbating liver pathologies. Data suggests that co-infection with schistosomes modulates the immune landscape, which may lead to poorer clinical outcomes particularly in immunocompromised populations. This relationship underscores the interaction between various pathogens and their impact on host disease trajectories (Colombe et al., 2018).

**Prevalence and Future Considerations:** Schistosomiasis continues to be a leading cause of chronic morbidity in populations globally, reinforcing the need for effective preventative measures such as praziquantel mass drug administration (Pearce & MacDonald, 2002). Future research necessitates a focused investigation into immune-modulating vaccine candidates, strategies for defining regulatory thresholds, and the dynamics of co-infection scenarios to enhance control efforts. Understanding these factors will be vital for tailoring public health interventions and limiting the socio-economic burden of schistosomiasis in endemic areas.

#### Bartonella species (Bartonella)

**Classification and Transmission Ecology:** Bartonella species are classified as fastidious gram-negative intracellular bacteria that rely on arthropod vectors for transmission or direct contact with infected animals. Notably, B. henselae is primarily transmitted to humans through scratches or bites from infected cats, as these felines are significant reservoirs for this pathogen (Chomel & Kasten, 2010) (Shikoska et al., 2023). Alternatively, B. quintana is primarily spread by body lice, demonstrating the diverse transmission pathways for Bartonella species (Harms & Dehio, 2012). These microorganisms are adept at infecting erythrocytes and endothelial cells, leading to chronic bacteremia, a condition characterized by the presence of bacteria in the bloodstream (Harms & Dehio, 2012) (Xi et al., 2024).

**Clinical Manifestations and Latency:** In humans, B. henselae is known to cause cat-scratch disease (CSD), which typically manifests as fever and localized lymphadenopathy. In immunocompromised individuals, severe complications may arise, including bacillary angiomatosis and peliosis hepatis, conditions marked by abnormal blood vessel formation and blood-filled cysts in the liver, respectively (Maggi et al., 2013) (Guptill, 2010). Conversely, B. quintana is responsible for trench fever, which presents with relapsing symptoms, including fever and bone pain, as well as more severe manifestations such as endocarditis (Boonjakuakul et al., 2007). These infections have the potential for asymptomatic persistence or recurrence, particularly in patients with compromised immune systems, due to the pathogen's ability to reside intracellularly (Harms & Dehio, 2012) (Siewert et al., 2022).

**Global Distribution and Seroprevalence:** Bartonella species have a wide global distribution, with B. henselae showing high seroprevalence rates among domestic cats, especially in warmer climates where flea populations are abundant (Chomel & Kasten, 2010) (Guptill, 2010). B. quintana, on the other hand, tends to be endemic in overcrowded and unsanitary conditions, often affecting homeless populations (Chomel & Kasten, 2010) (Pitassi et al., 2015). Human seroprevalence varies greatly based on geography and exposure history, indicating the importance of environmental and social factors in the epidemiology of Bartonella infections (Pitassi et al., 2015) (Breitschwerdt et al., 2010).

**Risk Factors for Opportunism:** Certain populations, particularly those affected by HIV/AIDS, are at increased risk for severe Bartonella infections (Breitschwerdt et al., 2010) (Siewert et al., 2022). Other key risk factors include homelessness, immunosuppressive therapies, exposure to flea-infested cats, and professions that involve close contact with animals, such as veterinary work (Boonjakuakul et al., 2007) (Guptill, 2010). In immunocompromised hosts, the incidence of conditions like endocarditis and disseminated disease is markedly higher, highlighting the opportunistic nature of these infections (Breitschwerdt et al., 2010) (Maggi et al., 2013).

**Pathogenesis and Persistence Mechanisms:** The pathogenicity of Bartonella species is largely attributed to their ability to invade endothelial cells and erythrocytes. This intracellular lifestyle enables them to evade the host's immune responses and maintain chronic bacteremia (Harms & Dehio, 2012) (Xi et al., 2024). They utilize type IV secretion systems to manipulate host cell functions, facilitating their survival and chronic infection (Harms & Dehio, 2012). Intracellular localization not only provides a sanctuary from antibodies but also promotes persistent infections that can lead to significant clinical manifestations despite host immune responses (Xi et al., 2024) (Breitschwerdt et al., 2010).

**Immune Arm Dependencies:** Effective control of Bartonella infections necessitates robust activation of macrophages and CD4+ T-cell responses (Breitschwerdt et al., 2010) (Xi et al., 2024). Although antibody-mediated clearance plays a role, it is often inefficient due to the bacteria's ability to hide within host cells (Harms & Dehio, 2012) (Breitschwerdt et al., 2010). This hidden lifestyle is particularly perilous for immunosuppressed individuals, leading to unchecked bacterial proliferation and subsequent complications such as bacillary angiomatosis and peliosis hepatis (Breitschwerdt et al., 2010) (Maggi et al., 2013).

**Compartmental Tropism and Failure:** Bartonella species exhibit a distinct tropism for particular tissues, preferring to infect endothelial cells, liver sinusoids, cardiac valves, and skin (Breitschwerdt et al., 2010) (Siewert et al., 2022). In compromised hosts, the development of angiogenic lesions across various organs is a common complication, complicating the diagnosis of conditions like endocarditis, which might occur without typical vegetative structures (Harms & Dehio, 2012) (Maggi et al., 2013).

**Immune Modulation and Evasion:** Bartonella species are known to secrete effector proteins that impede apoptosis, inhibit antigen presentation, and promote angiogenesis, all of which delay immune recognition and clearance (Harms & Dehio, 2012) (Xi et al., 2024). This strategic immune evasion allows for the establishment of chronic infections without the overt suppression of the immune system, complicating treatment and management efforts (Harms & Dehio, 2012) (Breitschwerdt et al., 2010).

**Immune Memory and Coordination:** Adaptive immune responses to Bartonella infections tend to be incomplete and short-lived. Immune memory, particularly in immunocompromised populations, is often insufficient, leading to relapses (Pitassi et al., 2015) (Breitschwerdt et al., 2010). Consequently, individuals may experience reinfections or recrudescence, emphasizing the need for ongoing vigilance and possible intervention strategies (Breitschwerdt et al., 2010) (Xi et al., 2024).

**Role in Immune Architecture:** Bartonella species occupy a complex position on the 'keystone pathogen spectrum'. They induce persistent infections that engage specific immune responses without providing strong, systemic immunity (Breitschwerdt et al., 2010) (Xi et al., 2024). This localized immune modulation allows Bartonella to coexist with the host while preserving the ability to spread, indicating a specialized approach to immune interaction that differs from traditional pathogens (Harms & Dehio, 2012) (Breitschwerdt et al., 2010).

**Impact of Immunosuppression:** Immunocompromised states, particularly in individuals with HIV/AIDS or those who are post-transplant, significantly amplify the severity and dissemination of Bartonella infections (Breitschwerdt et al., 2010; Siewert et al., 2022). The depletion of CD4+ T-cells is a significant factor that contributes to complications like bacillary angiomatosis and peliosis hepatis, reinforcing the importance of immune status in infection outcomes (Breitschwerdt et al., 2010) (Siewert et al., 2022).

**Prevalence and Future Considerations:** Despite their potential to be underdiagnosed, Bartonella infections pose significant clinical challenges, especially among immunocompromised patients (Chomel & Kasten, 2010) (Pitassi et al., 2015). Ongoing research should focus on elucidating the mechanisms underlying intracellular persistence, the nature of immune memory, and the interactions between co-infections. Additionally, enhanced diagnostic techniques and effective vector control measures are essential for improving public health outcomes related to Bartonella spp (Breitschwerdt et al., 2010) (Siewert et al., 2022).

#### Fusarium species (Fusarium)

**Classification and Transmission Ecology:** Fusarium is classified as a genus of filamentous fungi belonging to the class Sordariomycetes. This genus is widely distributed in diverse environments such as soil, water, and decomposing plant material. Human infections mainly occur through the inhalation of airborne conidia or through direct contact with contaminated surfaces, and these fungi are considered environmental pathogens rather than being specifically adapted to humans. Notably, Fusarium spp. exhibit persistence in various environmental settings, including plumbing systems and on surfaces, which is critical for their role as opportunistic pathogens (Nucci & Anaissie, 2007).

**Clinical Manifestations and Latency:** Fusarium spp. are etiological agents for a range of clinical presentations, including opportunistic infections in immunocompromised patients. Superficial infections like keratitis, onychomycosis, and cellulitis can occur, while more severe manifestations include locally invasive disease (e.g., sinusitis) and disseminated fusariosis, which often features fungemia and multi-organ involvement (Nucci & Anaissie, 2007) (Garnica et al., 2014). Importantly, Fusarium spp. do not establish latency, making timely intervention critical in at-risk populations (Nucci & Anaissie, 2007) (Hope et al., 2005).

**Global Distribution and Seroprevalence:** Globally, Fusarium is found predominantly in tropical and subtropical regions, and cases of fusariosis have been reported worldwide. Despite nearly universal environmental exposure, invasive Fusarium infections are primarily observed in severely immunocompromised individuals. Consequently, seroprevalence studies have indicated limited predictive value, as they do not effectively correlate with actual risk for invasive disease (Nucci & Anaissie, 2007) (Cafarchia et al., 2019).

**Risk Factors for Opportunism:** The primary risk factor for developing serious Fusarium infections is profound and prolonged neutropenia, which often accompanies hematologic malignancies or the effects of intensive treatments such as chemotherapy and stem cell transplants (Nucci & Anaissie, 2007) (Yao et al., 2021). Other significant risk factors include solid organ transplantation, immunosuppression from corticosteroid therapy, and underlying skin conditions that may predispose individuals to external infections. A study has shown that cutaneous portals of entry are prevalent in patients with hematologic disorders, thus increasing the likelihood of invasive fusariosis in such contexts (Garnica et al., 2014) (Yao et al., 2021).

**Pathogenesis and Persistence Mechanisms:** Fusarium species exhibit a range of virulence mechanisms that contribute to their pathogenicity. They produce microconidia and hyphal forms that can resist neutrophil-mediated effector functions, enabling them to colonize and invade epithelial tissues (Nucci & Anaissie, 2007) (Al-Hatmi et al., 2016). Additionally, Fusarium's ability to persist in water systems facilitates their transmission and the potential for hematogenous spread, distinguishing them from other mold pathogens (Nucci & Anaissie, 2007) (Al-Hatmi et al., 2016).

**Immune Arm Dependencies:** The host's innate immune response, particularly the actions of neutrophils, is crucial for controlling Fusarium infections. Studies have shown that reactive oxygen species (ROS) generated by neutrophils significantly contribute to the containment of these fungal pathogens. The adaptive immune system appears less effective against Fusarium, particularly in patients with neutropenia, leading to increased incidences of disseminated infections and serious clinical outcomes (Nucci & Anaissie, 2007) (Goyal et al., 2018).

**Compartmental Tropism and Failure:** Fusarium primarily targets the skin, sinopulmonary system, and bloodstream, with disseminated infections often characterized by painful, erythematous lesions that can become necrotic (Nucci & Anaissie, 2007) (Garnica et al., 2014). The lack of typical radiological signs, such as halo signs, in pulmonary presentations further complicates diagnosis and management, especially in neutropenic patients where prompt intervention is critical (Nucci & Anaissie, 2007) (Garnica et al., 2014).

**Immune Modulation and Evasion:** Fusarium spp. rely on rapid growth and angioinvasion rather than sophisticated immune modulation strategies found in other pathogens. Their ability to thrive in environments devoid of neutrophils marks a significant aspect of their pathogenesis, leading to severe infections in vulnerable populations (Nucci & Anaissie, 2007) (Garnica et al., 2014).

**Immune Memory and Coordination:** Current evidence suggests that there is insufficient adaptive immune memory post-Fusarium infection; memory T-cell responses do not seem to play a significant role, indicating that long-lasting immunity is not established. The clinical course of infections is primarily determined by the recovery of neutrophil function rather than any adaptive immune response (Nucci & Anaissie, 2007) (Hope et al., 2005).

Role in Immune Architecture

Fusarium does not significantly alter immune architecture but highlights the limitations of the innate immune system, particularly through the lens of neutrophil activity. It serves as a critical marker of neutrophil integrity and function in immunocompromised individuals, exhibiting high mortality in cases of profound immunosuppression (Nucci & Anaissie, 2007) (Garnica et al., 2014).

**Impact of Immunosuppression:** The burden of Fusarium infections weighs heavily on immunosuppressed demographics, especially those with neutropenia resulting from antineoplastic therapies or stem cell transplants, where mortality rates can exceed 50% despite antifungal treatment (Nucci & Anaissie, 2007) (Yao et al., 2021). This underscores the need for enhanced therapeutic strategies and monitoring in these high-risk groups (Nucci & Anaissie, 2007) (Garnica et al., 2014).

**Prevalence and Future Considerations:** While Fusarium infections are relatively rare, they pose a serious threat to immunocompromised individuals. Their inherent resistance to antifungal agents, particularly azoles, alongside their environmental resilience, poses challenges for prevention and treatment. The need for newer antifungal agents, improved diagnostic measures, and potential prophylaxis strategies in at-risk populations is critical (Nucci & Anaissie, 2007) (Garnica et al., 2014) (Yao et al., 2021).

#### Scedosporium species (Scedosporium)

**Classification and Transmission Ecology:** Scedosporium spp., including Scedosporium apiospermum and Scedosporium boydii, are filamentous fungi primarily found in environmental niches such as soil, polluted water, and sewage systems. Their classification within the fungal kingdom has become increasingly refined due to molecular studies that elucidate their phylogenetic relationships. Recent research has emphasized the phylogenetic separation of Scedosporium species from other related fungi such as Pseudallescheria, thus enhancing our understanding of their ecological roles and taxonomic classification (Gilgado et al., 2005) (Rougeron et al., 2017) (Lackner & Hoog, 2011). Transmission of Scedosporium occurs primarily through the inhalation of airborne conidia or via traumatic inoculation through skin contact with contaminated materials (Heath et al., 2009) (S. H. Kim et al., 2015). Environmental studies indicate that these fungi thrive particularly in human-modified habitats, such as wastewater treatment facilities and areas with high human activity, increasing the likelihood of exposure to humans (Mouhajir et al., 2020) (Luplertlop et al., 2019).

**Clinical Manifestations and Latency:** In healthy individuals, Scedosporium can manifest as localized infections, primarily affecting the skin, eyes (keratitis), or leading to the development of mycetoma, a chronic granulomatous infection (S. H. Kim et al., 2015) (Mouhajir et al., 2020). In contrast, immunocompromised individuals are at high risk for severe outcomes, including invasive pulmonary disease, central nervous system abscesses, fungemia, and disseminated infections, as observed in various clinical studies (Tamm et al., 2001) (Blyth et al., 2010) (Reinhold et al., 2024). The infections typically present acutely in high-risk patients without a notable latency period (Tamm et al., 2001) (Reinhold et al., 2024).

**Global Distribution and Seroprevalence:** Scedosporium spp. exhibit a global distribution but are more frequently isolated in temperate and tropical climates. Investigations into environmental samples have shown that these fungi are commonly identified in hospital drainage systems and stagnant water bodies, highlighting their prevalence in areas with compromised sanitation (Heath et al., 2009) (Mouhajir et al., 2020) (Luplertlop et al., 2019). However, there is a noted lack of significant seroprevalence data indicating long-term adaptive exposure in human populations, suggesting that while the fungi are widely present, they do not establish lasting immune relationships with their hosts (Rougeron et al., 2017) (Castelli et al., 2008).

**Risk Factors for Opportunism:** Certain clinical conditions significantly heighten the risk of Scedosporium infections. These include severe neutropenia, solid organ or hematopoietic stem cell transplantation, prolonged corticosteroid therapies, and chronic granulomatous disease (Reinhold et al., 2024) (Blyth et al., 2010). Furthermore, breakthrough infections are observed in patients undergoing prophylactic antifungal treatments with triazoles, highlighting the intrinsic resistance of Scedosporium spp. to conventional antifungal therapies, particularly amphotericin B and to some extent voriconazole (O'Bryan, 2005) (Reinhold et al., 2024).

**Pathogenesis and Persistence Mechanisms:** The pathogenesis of Scedosporium species involves angioinvasive growth patterns, particularly in immunocompromised hosts, leading to significant tissue damage and systemic dissemination (Blyth et al., 2010) (Reinhold et al., 2024). These fungi possess mechanisms that confer resistance against oxidative stress and antifungal drugs, including efficiency in managing oxidative agents through specific superoxide dismutase enzymes and the formation of biofilms (Staerck et al., 2021) (Reinhold et al., 2024). Such mechanisms complicate the clinical management of infections and underscore the urgent need for novel therapeutic strategies.

**Immune Arm Dependencies:** Scedosporium infections predominantly engage the innate immune system, particularly relying on neutrophils and macrophages for fungal clearance. Neutrophil-derived reactive oxygen species play a crucial role in this defense; however, evidence points to a limited activation of adaptive immunity in cases of rapid disease progression, complicating the patient's response to infection (Williamson et al., 2001) (Castelli et al., 2008). This suggests a need for further exploration into host-pathogen interactions and the role of adaptive immunity, especially in the context of severe fungal infections.

**Compartmental Tropism and Failure:** Scedosporium can affect multiple body compartments, including the skin, lungs, central nervous system, and vasculature. The involvement of the CNS, especially in cases following near-drowning incidents, is particularly concerning as it can lead not only to localized infections but also to systemic spread, especially in immunocompromised individuals (S. H. Kim et al., 2015) (Blyth et al., 2010) (Reinhold et al., 2024). The tendency for compartmental tropism is indicative of the fungal adaptation to various host tissues and underscores the need for vigilant monitoring in at-risk populations.

**Immune Modulation and Evasion:** Unlike classical pathogens, Scedosporium spp. do not actively modulate immune pathways. Instead, they employ strategies to evade phagocytic clearance. Their survival within host tissue is primarily attributed to their hardiness and immune evasion tactics, which include the development of structures that confer antifungal resistance

(Hoog et al., 1994) (Mouhajir et al., 2020) (Staerck et al., 2021). Understanding these mechanisms may inform future therapeutic approaches and guide vaccine development.

**Immune Memory and Coordination:** Current evidence indicates that Scedosporium infections do not generate structured adaptive immune memory. The likelihood of reinfection is more closely related to the host's immune status rather than prior exposure, suggesting a lack of durable immune imprinting or modulation of epitope hierarchy (Rougeron et al., 2017) (Castelli et al., 2008). This presents significant challenges for long-term management of infections, especially in populations with fluctuating immune competence.

**Role in Immune Architecture:** Scedosporium occupies a unique niche within the spectrum of opportunistic pathogens, demonstrating the limitations of innate immunity in response to invasive fungal infections. It does not influence systemic immune balance but rather takes advantage of phagocyte function, highlighting the episodic nature of its pathogenic impact (Castelli et al., 2008) (Reinhold et al., 2024).

**Impact of Immunosuppression:** Levels of immunosuppression critically dictate susceptibility to Scedosporium infections. Neutropenia and corticosteroid use significantly increase the risk, leading to severe outcomes, particularly in patients undergoing triazole therapy due to the inherent resistance of Scedosporium to conventional antifungals (Markowska-Szczupak et al., 2015) (Reinhold et al., 2024). The emergence of resistant strains complicates the treatment landscape, necessitating further research into host-pathogen dynamics and improved antifungal protocols.

**Prevalence and Future Considerations:** Infections with Scedosporium spp. remain a pressing concern within increasingly immunocompromised populations globally. The rise of antifungal resistance, particularly to triazoles, coupled with limited therapeutic options, underscores the urgent need for enhanced diagnostic measures and innovative antifungal strategies (Heath et al., 2009) (Castelli et al., 2008) (Reinhold et al., 2024). Continued research should prioritize the mechanisms of neutrophil-mediated clearance and host-pathogen interactions to inform future therapeutic and preventative approaches.

#### Acanthamoeba species (Acanthamoeba)

**Classification and Transmission Ecology:** Acanthamoeba spp. is classified within the phylum Amoebozoa and are characterized as free-living, opportunistic amoebae often found in diverse environments such as soil, freshwater, dust, and air conditioning systems. They exist in two primary forms: trophozoites (active feeding stage) and cysts (dormant and resistant forms). This dual capability allows them to thrive in various ecological niches, facilitating widespread geographical distribution (Kotton & Lattes, 2009) (Magnet et al., 2012) (Orosz & Posta, 2020). Transmission primarily occurs through direct contact with contaminated water or soil, where the amoebae can enter the human body via corneal microtrauma, often exacerbated by contact lens use, as well as through inhalation or breaks in the skin (Kotton & Lattes, 2009) (Magnet et al., 2012) (Libbey et al., 2013). The ubiquitous nature of Acanthamoeba in public infrastructures such as water treatment facilities underscores the constant risk of exposure in modern living environments (Magnet et al., 2012).

**Clinical Manifestations and Latency:** Acanthamoeba infections primarily lead to two significant clinical syndromes: keratitis and granulomatous amoebic encephalitis (GAE). Keratitis, particularly prevalent among contact lens users, is marked by symptoms including severe ocular pain, redness, and photophobia, often leading to vision loss if not timely addressed (Libbey et al., 2013) (Lee et al., 2006). GAE is noted for its devastating impact on immunocompromised individuals, resulting in neurological symptoms such as seizures and altered mental status, thus showcasing the dire potential of Acanthamoeba infections in vulnerable populations (Libbey et al., 2013) (Kotton & Lattes, 2009). Latency has been suggested in cases of cutaneous or central nervous system infections, indicating that some individuals may harbor the pathogen without active disease presentation (Libbey et al., 2013).

**Global Distribution and Seroprevalence:** The global distribution of Acanthamoeba is noteworthy, with studies indicating its presence across a broad spectrum of natural and man-made environments. Investigations reveal high seroprevalence rates of anti-Acanthamoeba antibodies among healthy populations, often exceeding 80%, which implies widespread asymptomatic exposure to these pathogens (Kotton & Lattes, 2009) (Magnet et al., 2012). Despite this high rate of exposure, clinical manifestations remain relatively rare, highlighting a significant gap between environmental presence and disease incidence (Magnet et al., 2012) (Kotton & Lattes, 2009). This contrast emphasizes the need to understand the underlying factors contributing to infection susceptibility (Schwartz & Mawhorter, 2013) (Kotton & Lattes, 2009).

**Risk Factors for Opportunism:** Key risk factors for opportunistic infections include immunosuppression attributable to conditions like HIV/AIDS and hematologic malignancies or immunosuppressive therapies such as corticosteroids (Schwartz & Mawhorter, 2013) (Kotton & Lattes, 2009). In terms of keratitis, improper contact lens hygiene and exposure to contaminated water while wearing lenses are substantial risk contributors (Kotton & Lattes, 2009). The interplay between individual health status and specific environmental exposures creates a complex risk landscape that must be navigated to mitigate the likelihood of infection (Magnet et al., 2012) (Kotton & Lattes, 2009).

**Pathogenesis and Persistence Mechanisms:** The pathogenicity of Acanthamoeba is heightened by its capacity to exist as both trophozoites and cysts. Trophozoites actively invade host tissues, where they deploy proteolytic enzymes that facilitate cytolysis, leading to tissue necrosis and inflammation, particularly in GAE cases where they cross the blood-brain barrier (Kotton & Lattes, 2009) (Libbey et al., 2013). Cysts exhibit exceptional resistance to standard disinfection methods and environmental stresses, thereby enhancing their survival and transmissibility (Kotton & Lattes, 2009) (Veugen et al., 2025). This cystic phase is critical for survival in unfavorable conditions, contributing to the persistence of the organism in diverse settings (Kotton & Lattes, 2009).

**Immune Arm Dependencies:** The immune response against Acanthamoeba is predominantly reliant on innate defenses, particularly macrophage activity and cytotoxic T-cell responses (Kotton & Lattes, 2009) (Magnet et al., 2012). Neutrophils and complement pathways also play a role in early containment of keratitis. However, deficiencies in both innate and adaptive immunity can predispose individuals to severe forms of Acanthamoeba disease (Schwartz & Mawhorter, 2013) (Libbey et al., 2013) (Kotton & Lattes, 2009). Understanding immune interaction profiles is essential for developing targeted therapeutic strategies and prophylactic measures against infections (Kotton & Lattes, 2009) (Khan & Siddiqui, 2014).

**Compartmental Tropism and Failure:** In immunocompetent individuals, Acanthamoeba primarily targets ocular tissues, manifesting as keratitis. In contrast, in immunocompromised hosts, CNS involvement becomes more prevalent, often resulting in GAE. Cutaneous lesions may also be observed concomitant to GAE, revealing the amoeba's multifaceted tropism within the host (Libbey et al., 2013) (Kotton & Lattes, 2009). Cases of multiorgan dissemination remain uncommon but have been documented, reinforcing the need for vigilant monitoring in high-risk scenarios (Schwartz & Mawhorter, 2013).

**Immune Modulation and Evasion:** Acanthamoeba exhibits several strategies for evading immune responses, including the secretion of proteolytic enzymes that degrade complement proteins and immunoglobulins. These actions hinder effective immune clearance, while its cystic form further resists disinfection efforts (Kotton & Lattes, 2009) (Libbey et al., 2013). Despite this active evasion, it is noted that Acanthamoeba does not possess mechanisms for sustained immune modulation or antigenic variation, highlighting the unique challenges it presents to host defenses (Magnet et al., 2012) (Kotton & Lattes, 2009).

**Immune Memory and Coordination:** Studies indicate that individuals recovering from Acanthamoeba infections do not garner protective immunity against subsequent infections. This phenomenon underscores a limitation of the adaptive immune response, where minimal long-term memory or epitope hierarchy development occurs, thereby hindering the potential for immunological memory against reinfection (Schwartz & Mawhorter, 2013) (Kotton & Lattes, 2009) (Khan & Siddiqui, 2014). This aspect complicates the clinical landscape surrounding management and prevention of Acanthamoeba-related diseases.

**Role in Immune Architecture:** In broader contexts, Acanthamoeba interactions with human immune systems often reveal deficiencies in macrophage function and cellular immunity rather than shaping overall immune priorities. This propensity positions Acanthamoeba more as a "stress-test" pathogen that intermittently exposes weaknesses in innate immune systems, emphasizing the necessity for comprehensive immunological assessments in affected individuals (Libbey et al., 2013) (Khan & Siddiqui, 2014).

**Impact of Immunosuppression:** Immunocompromised patients, particularly those undergoing corticosteroid therapy or solid organ transplantation, exhibit a higher susceptibility to GAE, with case outcomes tending to be rapid and frequently fatal (Schwartz & Mawhorter, 2013) (Libbey et al., 2013) (Kotton & Lattes, 2009). Topical corticosteroids used in contact lens wearers can further contribute to the exacerbation of keratitis, complicating treatment regimens and necessitating stringent guidance for at-risk populations (Kotton & Lattes, 2009) (Khan & Siddiqui, 2014).

**Prevalence and Future Considerations:** The awareness of Acanthamoeba infections has risen, closely tied to the increased prevalence of contact lens use and enhanced diagnostic capabilities. Future research endeavors should delve into elucidating the macrophage and T-cell response thresholds critical for infection containment while exploring disinfection strategies to mitigate keratitis risks (Schwartz & Mawhorter, 2013) (Libbey et al., 2013) (Kotton & Lattes, 2009). Understanding these dynamics is paramount as they inform public health strategies and clinical management paradigms.

#### Vibrio species (Vibrio)

**Classification and Transmission Ecology:** Vibrio species, which are gram-negative, halophilic bacteria, predominantly inhabit marine and estuarine environments where they thrive in warm, saline waters. Key pathogenic species include *Vibrio vulnificus*, *V. parahaemolyticus*, and non-O1/non-O139 strains of *V. cholerae*. These bacteria are significant due to their association with diseases following the ingestion of contaminated seafood, particularly raw oysters, or through direct exposure of open wounds to warm coastal waters (Finkelstein & Oren, 2011). Environmental factors such as salinity and water temperature impact the abundance and pathogenic potential of Vibrio species (Wong et al., 2024).

**Clinical Manifestations and Latency:** Upon infection, symptoms typically manifest acutely and can range markedly in severity. *V. parahaemolyticus* primarily leads to self-limiting gastroenteritis, characterized by diarrhea and abdominal cramps, whereas *V. vulnificus* infections can escalate to severe outcomes, including necrotizing fasciitis, septicemia, and potential multiorgan failure (Mariya et al., 2021). Additionally, non-O1/non-O139 strains of *V. cholerae* can cause a spectrum of conditions from diarrhea to wound infections and septicemia, particularly in immunocompromised individuals. Importantly, infections by these bacteria do not exhibit latency, maintaining the acute nature of clinical presentations (Azwai et al., 2016).

**Global Distribution and Seroprevalence:** Globally, Vibrio species display a wide geographic distribution, particularly in the coastal waters of tropical and subtropical regions. Their prevalence is notably higher during the warmer months, correlating with increased water temperatures that favor their proliferation (Wang et al., 2021). Regions such as the Gulf of Mexico and Southeast Asia report higher incidences of infections, underlining the impact of local environmental conditions on seroprevalence (Brumfield et al., 2023) (Karatuna et al., 2023). The ongoing trend of rising sea surface temperatures poses significant concerns for public health, as it expands the habitat range of these pathogens, potentially leading to increased infection rates in newer locales (Banerjee & Farber, 2017) (Karatuna et al., 2023).

**Risk Factors for Opportunism:** Individuals with existing health conditions such as liver disease (including cirrhosis and hemochromatosis), diabetes, and immunosuppression (including those undergoing chemotherapy or with neutropenia) are at heightened risk of severe *Vibrio* infections (Chen et al., 2011) (Charles et al., 2017). Environmental risk factors, particularly the consumption of raw or undercooked shellfish and exposure of wounds to contaminated seawater, significantly elevate the likelihood of infection (Mariya et al., 2021) (Mancini et al., 2023).

**Pathogenesis and Persistence Mechanisms:** The pathogenicity of *V. vulnificus* can be attributed to various virulence factors, including capsular polysaccharides, which enhance tissue invasion, alongside cytolysins and metalloproteases that facilitate immune evasion (Hsiao et al., 2006). Following entry, *V. vulnificus* can rapidly disseminate through the bloodstream, resulting in severe systemic illness, particularly septic shock. Although the pathogen does not establish chronic infections, it can survive in the environment by entering a viable but nonculturable state, permitting its persistence in aquatic habitats (Li et al., 2012).

**Immune Arm Dependencies:** The host's defense against Vibrio infections primarily relies on innate immunity, with neutrophils playing a pivotal role in controlling the initial response to infection (H. J. Kim et al., 2015). In individuals with impairments such as neutropenia or liver dysfunction, this innate defense is compromised, leading to increased susceptibility to severe infections (Mancini et al., 2023). While TNF-α and complement pathways contribute to pathogen clearance, adaptive immunity has less impact on the outcome of Vibrio infections, indicating a critical reliance on innate immune functions (Karatuna et al., 2023).

**Compartmental Tropism and Failure:** Vibrio species predominantly target the gastrointestinal tract and skin/soft tissue, leading to varied clinical outcomes depending on the patient's immune status. For instance, gastrointestinal infections caused by *V. parahaemolyticus* are usually self-limited, while infections from *V. vulnificus* can swiftly progress from localized cellulitis to systemic sepsis (Finkelstein & Oren, 2011) (Mizuno et al., 2020). Understanding this compartmental tropism is essential for identifying patients who may be at higher risk of severe infections, particularly in immunocompromised populations (Mariya et al., 2021) (Chen et al., 2011).

**Immune Modulation and Evasion:** Vibrio species evade host immune responses through rapid replication and tissue invasion mechanisms. They employ antiphagocytic strategies, including the expression of capsules and proteases that hinder opsonization and clearance by immune cells, thereby facilitating disease progression in susceptible individuals (Aguinaga et al., 2009).

**Immune Memory and Coordination:** Infections with Vibrio species are characterized by the absence of durable or broadly protective adaptive immunity. Prior exposure does not confer robust immunity, as immune responses are typically strain-specific and show limited persistence (Richards et al., 2005). Thus, recurrent infections remain possible, and the outcome is significantly influenced by the host's innate immune capacity at the time of new exposure (Richards et al., 2005).

**Role in Immune Architecture:** Vibrio species do not function as classical keystone organisms in the microbial ecosystem; however, they can reveal vulnerabilities within the host's innate immune responses, particularly affecting neutrophil functions (Karatuna et al., 2023). This opportunistic behavior underscores the importance of maintaining a healthy immune surveillance mechanism, especially in vulnerable populations (Richards et al., 2005).

**Impact of Immunosuppression:** The burden of severe infections caused by *V. vulnificus* is disproportionately high among immunocompromised patients, particularly those with liver disease or receiving immunosuppressive therapies, often leading to critical health outcomes and high mortality rates (Mizuno et al., 2020). Overall, timely recognition and treatment of these infections are pivotal to improving clinical outcomes (Mariya et al., 2021).

**Prevalence and Future Considerations:** With the projected impacts of climate change leading to increased seawater temperatures and expanding coastal populations, severe infections from *V. vulnificus* are anticipated to rise. There is an urgent need to enhance rapid diagnostic methods, establish early intervention protocols for high-risk populations, and implement environmental monitoring strategies to mitigate the risks of such infections (Navarro-Navarro et al., 2013) (Karatuna et al., 2023).

#### Leptospira species (Leptospira)

**Classification and Transmission Ecology:** Leptospira spp. are classified as aerobic, motile spirochetes that thrive in freshwater and moist soil environments. These bacteria are primarily harbored in the renal tubules of reservoir hosts, particularly rodents, from which they are excreted into the environment through urine, facilitating transmission through contaminated water or soil. Studies have shown that the presence of environmental factors, such as heavy rainfall and flooding, is significantly associated with the incidence of Leptospira infections, confirming the critical role of these conditions in the transmission dynamics of leptospirosis (Lau et al., 2010) (Mwachui et al., 2015) (Chadsuthi et al., 2018).

**Clinical Manifestations and Latency:** The clinical presentation of leptospirosis can be biphasic, beginning with a non-specific acute phase characterized by fever, headache, myalgia, and conjunctival suffusion. In severe cases, known as Weil's disease, patients may experience symptoms such as jaundice, acute kidney injury, and multi-organ failure. Research indicates that while Leptospira can persist in renal tissues of animals, chronic latency in humans remains rare, underscoring the necessity for timely diagnosis and intervention (Wang et al., 2012) (Kurniawati & Nuryati, 2018) (Guerra, 2013).

**Global Distribution and Seroprevalence:** Leptospira spp. exhibit a global distribution, showing a higher incidence in tropical and subtropical regions, particularly where sanitary conditions are inadequate and flooding occurs frequently. Seroprevalence studies reveal significant variation by occupation and exposure risk, with agricultural workers, urban poverty-stricken populations, and individuals engaging in recreational water activities being notably affected (Lau et al., 2016) (Chadsuthi et al., 2017) (Cook et al., 2016). Moreover, increased flooding events exacerbated by climate change have been linked to heightened leptospirosis risk in many regions (Lau et al., 2010) (Lau et al., 2016) (Chadsuthi et al., 2018).

**Risk Factors for Opportunism:** Occupational exposure remains a significant risk factor for leptospirosis, with activities such as farming, sewage work, and animal husbandry identified as high-risk environments. Recreational activities, including swimming and rafting in contaminated waters, also elevate infection risk. Additionally, socioeconomic factors such as urban poverty can amplify vulnerability to the disease, particularly in areas susceptible to environmental stresses, which are often exacerbated by climate change (Mwachui et al., 2015) (Adi et al., 2013). Impaired innate immunity from various health conditions can lead to increased disease severity in susceptible individuals (Lau et al., 2016) (Adi et al., 2013).

**Pathogenesis and Persistence Mechanisms:** Upon entry into the host, Leptospira bacteria disseminate through the bloodstream, adhering to endothelial and epithelial surfaces via outer membrane proteins. The bacteria's ability to resist complement lysis and phagocytosis during the early stages of infection has been well-documented, illustrating their capacity to evade host immune defenses. Furthermore, Leptospira demonstrates a tropism for vital organs such as the liver, kidneys, lungs, and central nervous system (CNS) (Wang et al., 2012) (Lin et al., 2011).

**Immune Arm Dependencies:** The innate immune response, particularly through neutrophil and macrophage activation, is crucial for controlling early Leptospira infection. Conversely, the roles of adaptive immunity (e.g., antibody production and T-cell responses) are less critical, and the bacteria have evolved mechanisms to evade strong immune engagement, allowing for opportunistic dissemination within the host (Lin et al., 2011) (Guerra, 2013) (Adi et al., 2013).

**Compartmental Tropism and Failure:** Severe manifestations of leptospirosis are associated with multiple organ failure, including hepatic dysfunction, renal impairment, and potential pulmonary hemorrhage. The ability of Leptospira to invade the CNS may contribute to cases presenting with aseptic meningitis. Additionally, the role of renal tubules as a chronic reservoir in asymptomatic animal carriers emphasizes the complexity of disease transmission and persistence (Wang et al., 2012) (Guerra, 2013) (Adi et al., 2013).

**Immune Modulation and Evasion:** Leptospira exhibits several strategies to evade host immune responses, including the secretion of proteases that inhibit neutrophil activity and the expression of outer membrane proteins that disrupt complement activation. This immune modulation provides a window during which the bacteria can disseminate before being effectively cleared by the immune system (Lin et al., 2011) (Guerra, 2013) (Adi et al., 2013).

**Immune Memory and Coordination:** Protective immunity following leptospirosis infection appears to be incomplete and often serovar-specific. Studies indicate that memory T-cells may not persist long-term, leading to vulnerability to reinfection in endemic regions. The lack of effective coordination of multi-compartmental immunity further complicates the effective management of recurrent infections (Mwachui et al., 2015) (Adi et al., 2013).

**Role in Immune Architecture:** Leptospira acts as an environmental opportunist rather than systematically structuring host immune responses. While it may trigger systemic inflammation, it does not persistently shape immune prioritization, emphasizing the threshold responses of the host rather than an orchestrated immune response (Guerra, 2013) (Adi et al., 2013).

**Impact of Immunosuppression:** Individuals with compromised immune systems may experience exacerbated leptospirosis, but research indicates that the most severe cases are often tied to intense environmental exposure rather than solely immune deficiency. Factors such as neutropenia and corticosteroid usage are known to heighten risks for adverse disease outcomes (Wang et al., 2012) (Guerra, 2013) (Adi et al., 2013).

**Prevalence and Future Considerations:** Leptospirosis remains an underrecognized but significant public health issue, particularly in tropical and subtropical areas. As climate change and urbanization continue to pose increased risks, further research is warranted to better understand immune thresholds, mechanisms of complement evasion, and the development of vaccines that provide durable, cross-serovar immunity (Lau et al., 2010) (Lau et al., 2016) (Wang et al., 2012).

#### Refezences

Adetunji, S. A., Faustman, D. L., Adams, L. G., González, D. G., Hensel, M. E., Khalaf, O. H., & Arenas-Gamboa, A. M. (2020). Brucella Abortus and Pregnancy in Mice: Impact of Chronic Infection on Fertility and the Role of Regulatory T Cells in Tissue Colonization. *Infect Immun*, *88*(10). https://doi.org/10.1128/iai.00257-20

Adi, M. S., Bertolatti, D., Maycock, B., Spickett, J., & Dhaliwal, S. S. (2013). Risk Factors for Leptospirosis Infection in Humans and Implications for Public Health Intervention in Indonesia and the Asia-Pacific Region. *Asia Pacific Journal of Public Health*, *26*(1), 15-32. https://doi.org/10.1177/1010539513498768

Aghakhani, A., Mamishi, S., Sabeti, S., Bidari-Zerehpoosh, F., Banifazl, M., Bavand, A., & Ramezani, A. (2016). Gender and Age-Specific Seroprevalence of Human Papillomavirus 16 and 18 in General Population in Tehran, Iran. *Medical Microbiology and Immunology*, *206*(2), 105-110. https://doi.org/10.1007/s00430-016-0487-5

Águas, R., Ferreira, M. U., & Gomes, M. G. M. (2012). Modeling the Effects of Relapse in the Transmission Dynamics of Malaria Parasites. *Journal of Parasitology Research*, *2012*, 1-8. https://doi.org/10.1155/2012/921715

Aguinaga, A., Portillo, M. E., Yuste, J. R., Pozo, J. L. d., García-Tutor, E., Pérez-Gracia, J. L., & Leiva, J. (2009). Non-O1 Vibrio Cholerae Inguinal Skin and Soft Tissue Infection With Bullous Skin

Lesions in a Patient With a Penis Squamous Cell Carcinoma. *Annals of Clinical Microbiology and Antimicrobials*, *8*(1), 17. https://doi.org/10.1186/1476-0711-8-17

Agut, H., Bonnafous, P., & Gautheret-Dejean, A. (2017). Update on Infections With Human Herpesviruses 6A, 6B, and 7. *Médecine Et Maladies Infectieuses*, *47*(2), 83-91. https://doi.org/10.1016/j.medmal.2016.09.004

Akbar, D. H., & Tahawi, A. T. (2001). Candidemia at a University Hospital: Epidemiology, Risk Factors and Predictors of Mortality. *Annals of Saudi Medicine*, *21*(3-4), 178-182. https://doi.org/10.5144/0256-4947.2001.178

Al-Hatmi, A. M. S., Meis, J. F., & Hoog, S. d. (2016). Fusarium: Molecular Diversity and Intrinsic Drug Resistance. *PLOS Pathogens*, *12*(4), e1005464. https://doi.org/10.1371/journal.ppat.1005464

Al-Zubaidi, N., Al-Zubaidi, M. S. K., Gholizadeh, N., & Zaresharifi, S. (2024). Cutaneous Perianal Herpes Simplex Virus Infection Mimicking Condylomata Acuminata in a Renal Transplant Recipient: A Case Report and Literature Review. *Clinical Case Reports*, *12*(9). https://doi.org/10.1002/ccr3.9428

Albayrak, H. T., Bakır, A., Güney, M., & Yavuz, M. T. (2019). Investigation of Parvovirus B19 Antibodies in Serum Samples With Pre-Diagnosis of Arthritis-Arthralgia. *The Anatolian Journal of Family Medicine*. https://doi.org/10.5505/anatoljfm.2019.35220

Alcendor, D. J. (2019). BK Polyomavirus Virus Glomerular Tropism: Implications for Virus Reactivation From Latency and Amplification During Immunosuppression. *Journal of Clinical Medicine*, *8*(9), 1477. https://doi.org/10.3390/jcm8091477

Alday, H., & Doggett, J. S. (2017). Drugs in Development for Toxoplasmosis: Advances, Challenges, and Current Status. *Drug Design Development and Therapy*, *Volume11*, 273-293. https://doi.org/10.2147/dddt.s60973

Alston, C. I., & Dix, R. D. (2017). Murine Cytomegalovirus Infection of Mouse Macrophages Stimulates Early Expression of Suppressor of Cytokine Signaling (SOCS)1 and SOCS3. *PLoS One*, *12*(2), e0171812. https://doi.org/10.1371/journal.pone.0171812

Ambalathingal, G., Francis, R. S., Smyth, M. J., Smith, C., & Khanna, R. (2017). BK Polyomavirus: Clinical Aspects, Immune Regulation, and Emerging Therapies. *Clinical Microbiology Reviews*, *30*(2), 503-528. https://doi.org/10.1128/cmr.00074-16

Amin, K., Rosshandler, Y., Norris, J. M., & Arinze, F. (2023). A Case of Mycobacterium Avium Complex Vertebral Osteomyelitis in a Patient With Human Immunodeficiency Virus. *Infectious Diseases in Clinical Practice*, *31*(4). https://doi.org/10.1097/ipc.0000000000001283

Amraei, F., Narimisa, N., Sadeghi, B., Lohrasbi, V., & Jazi, F. M. (2020). Persister Cells Formation and Expression of Type II Toxin-Antitoxin System Genes in Brucella Melitensis (16M) and Brucella Abortus (B19). *Iranian Journal of Pathology*, *15*(2), 127-133. https://doi.org/10.30699/ijp.2020.118902.2294

Antonsson, A., Waterboer, T., Bavinck, J. N. B., Abeni, D., Koning, M. d., Euvrard, S., Feltkamp, M. C., Green, A. C., Harwood, C. A., Naldi, L., Nindl, I., Pfister, H., Proby, C. M., Quint, W., Stockfleth, E., Weißenborn, S. J., Pawlita, M., & Neale, R. E. (2013). Longitudinal Study of Seroprevalence and Serostability of 34 Human Papillomavirus Types in European Organ Transplant Recipients. *Virology*, *436*(1), 91-99. https://doi.org/10.1016/j.virol.2012.10.037

Arango, M., Castañeda, E., Agudelo, C. I., Bedout, C. d., Agudelo, C. A., Tobón, Á. M., Linares, M., Valencia, Y., Restrepo, Á., & The Colombian Histoplasmosis Study, G. (2011). Histoplasmosis en

Colombia: Resultados De La Encuesta Nacional, 1992-2008. *Biomédica*, *31*(3), 344. https://doi.org/10.7705/biomedica.v31i3.348

Arena, A., Iannello, D., Gazzara, D., Speranza, A., Bonina, L., & Mastroeni, P. (2001). Role of Interleukin-18 in Peripheral Blood Mononuclear Cells Infected With Human Herpes Virus Type 6. *Intervirology*, *44*(4), 250-254. https://doi.org/10.1159/000050055

Arendrup, M. C., Dzajic, E., Hare, R. K., Johansen, H. K., Kjaldgaard, P., Knudsen, J. D., Kristensen, L. H., Leitz, C., Lemming, L., Nielsen, L., Olesen, B., Rosenvinge, F. S., Røder, B., & Schønheyder, H. C. (2013). Epidemiological Changes With Potential Implication for Antifungal Prescription Recommendations for Fungaemia: Data From a Nationwide Fungaemia Surveillance Programme. *Clinical Microbiology and Infection*, *19*(8), e343-e353. https://doi.org/10.1111/1469-0691.12212

Aris, R. M., Routh, J. C., LiPuma, J. J., Heath, D. G., & Gilligan, P. H. (2001). Lung Transplantation for Cystic Fibrosis Patients With <i>Burkholderia Cepacia</I> Complex. *American Journal of Respiratory and Critical Care Medicine*, *164*(11), 2102-2106. https://doi.org/10.1164/ajrccm.164.11.2107022

Arvin, A. M., Moffat, J. F., Sommer, M., Oliver, S. L., Che, X., Vleck, S. E., Zerboni, L., & Ku, C. C. (2010). Varicella-Zoster Virus T Cell Tropism and the Pathogenesis of Skin Infection. 189-209. https://doi.org/10.1007/82_2010_29

Atlas, R. (1999). <i>Legionella</I>: From Environmental Habitats to Disease Pathology, Detection and Control. *Environmental Microbiology*, *1*(4), 283-293. https://doi.org/10.1046/j.1462-2920.1999.00046.x

Azwai, S. M., Alfallani, E. A., Abolghait, S. K., Garbaj, A. M., Naas, H. T., Moawad, A. A., Gammoudi, F. T., Rayes, H. M., Barbieri, I., & Eldaghayes, I. (2016). Isolation and Molecular Identification of ≪i>Vibrio</I> Spp. By Sequencing of 16S rDNA From Seafood, Meat and Meat Products in Libya. *Open Veterinary Journal*, *6*(1), 36. https://doi.org/10.4314/ovj.v6i1.6

Babu, T. M., Laguio, M., Luque, A. E., & Larppanichpoonphol, P. (2013). <i>Mycobacterium Avium-</I>complex Pericarditis: A Case of Unmasking Immune Reconstitution Inflammatory Syndrome. *International Journal of STD & Aids*, *25*(2), 148-151. https://doi.org/10.1177/0956462413495821

Baddley, J. W., Schain, D., Gupte, A., Lodhi, S. A., Kayler, L. K., Frade, J. P., Lockhart, S. R., Chiller, T., Bynon, J. S., & Bower, W. A. (2011). Transmission of Cryptococcus Neoformans by Organ Transplantation. *Clinical Infectious Diseases*, *52*(4), e94-e98. https://doi.org/10.1093/cid/ciq216

Bakardjiev, A. I., Azimi, P. H., Ashouri, N., Ascher, D. P., Janner, D., Schuster, F. L., Visvesvara, G. S., & Glaser, C. (2003). Amebic Encephalitis Caused by Balamuthia Mandrillaris: Report of Four Cases. *The Pediatric Infectious Disease Journal*, *22*(5), 447-452. https://doi.org/10.1097/01.inf.0000066540.18671.f8

Baldi, P. C., & Giambartolomei, G. H. (2013). Pathogenesis and Pathobiology of Zoonotic Brucellosis in Humans. *Revue Scientifique Et Technique De L Oie*, *32*(1), 117-125. https://doi.org/10.20506/rst.32.1.2192

Banerjee, S. K., & Farber, J. M. (2017). Detection, Enumeration, and Isolation of Vibrio Parahaemolyticus and v. Vulnificus From Seafood: Development of a Multidisciplinary Protocol. *Journal of Aoac International*, *100*(2), 445-453. https://doi.org/10.5740/jaoacint.16-0290

Banicioiu, F. C., Chirilă, C., Toderan, A., Florescu, S. A., & Popescu, C. P. (2023). Bacterial Meningoencephalitis Secondary to Disseminated Strongyloidiasis in a Pacient With COVID-19. *Romanian Journal of Infectious Diseases*, *26*(3), 116-119. https://doi.org/10.37897/rjid.2023.3.3
Barton, E. S., Mandal, P., & Speck, S. H. (2011). Pathogenesis and Host Control of Gammaherpesviruses: Lessons From the Mouse. *Annual Review of Immunology*, *29*(1), 351-397. https://doi.org/10.1146/annurev-immunol-072710-081639
Bartsch, T., Rempe, T., Leypoldt, F., Riedel, C., Jansen, O., Berg, D., & Deuschl, G. (2019). The Spectrum of Progressive Multifocal Leukoencephalopathy: A Practical Approach. *European Journal of Neurology*, *26*(4), 566. https://doi.org/10.1111/ene.13906
Bate, S. L., Dollard, S. C., & Cannon, M. J. (2010). Cytomegalovirus Seroprevalence in the United States: The National Health and Nutrition Examination Surveys, 1988–2004. *Clinical Infectious Diseases*, *50*(11), 1439-1447. https://doi.org/10.1086/652438
Bayliss, J., Karasoulos, T., & McLean, C. (2011). Frequency and Large T (LT) Sequence of JC Polyomavirus DNA in Oligodendrocytes, Astrocytes and Granular Cells in Non-PML Brain. *Brain Pathology*, *22*(3), 329-336. https://doi.org/10.1111/j.1750-3639.2011.00538.x
Baylor, P., & Larson, R. K. (2008). Mycobacterium Avium Septicemia With ARDS in a Patient With Diabetes Mellitus and No Other Known Immune-Compromising Condition. *Journal of Intensive Care Medicine*, *24*(2), 140-143. https://doi.org/10.1177/0885066608330103
Beck, O., Topp, M. S., Koehl, U., Roilides, E., Simitsopoulou, M., Hanisch, M., Sarfati, J., Latgé, J. P., Klingebiel, T., Einsele, H., & Lehrnbecher, T. (2006). Generation of Highly Purified and Functionally Active Human TH1 Cells Against Aspergillus Fumigatus. *Blood*, *107*(6), 2562-2569. https://doi.org/10.1182/blood-2005-04-1660
Belhassen-García, M., Alberto, R. a. b.-G. e. r., Velasco-Tirado, V., Romero-Alegría, Á., rez-Garcia, M.-L. P. e., & Martín-Oterino, J.-Á. (2015). Atypical Progressive Multifocal Leukoencephalopathy in a Patient With Antisynthetase Syndrome. *Internal Medicine*, *54*(5), 519-524. https://doi.org/10.2169/internalmedicine.54.2748
Benedict, K., Toda, M., & Jackson, B. R. (2021). Revising Conventional Wisdom About Histoplasmosis in the United States. *Open Forum Infectious Diseases*, *8*(7). https://doi.org/10.1093/ofid/ofab306
Benedict, S. (2023). Identification of B- Cell and T- Cell Specific Peptide Vaccine for Histoplasma Capsulatum. *Journal of Advanced Zoology*, *44*(S4), 158-166. https://doi.org/10.17762/jaz.v44is4.2411
Bento, J. T., Coelho, A. C., Rebelo, H., & Mesquita, J. R. (2025). Histoplasma Capsulatum in Bat Species in Portugal. *Veterinary Sciences*, *12*(2), 94. https://doi.org/10.3390/vetsci12020094
Bhatia, S., Afanasiev, O. K., & Nghiem, P. (2011). Immunobiology of Merkel Cell Carcinoma: Implications for Immunotherapy of a Polyomavirus-Associated Cancer. *Current Oncology Reports*, *13*(6), 488-497. https://doi.org/10.1007/s11912-011-0197-5
Bhatia, S., Fei, M., Yarlagadda, M., Qi, Z., Akira, S., Saijo, S., Iwakura, Y., Rooijen, N. v., Gibson, G. A., Croix, C. M. S., Ray, A., & Ray, P. (2011). Rapid Host Defense Against Aspergillus Fumigatus Involves Alveolar Macrophages With a Predominance of Alternatively Activated Phenotype. *PLoS One*, *6*(1), e15943. https://doi.org/10.1371/journal.pone.0015943
Bhumiratana, A., Intarapuk, A., Sorosjinda-Nunthawarasilp, P., Maneekan, P., & Koyadun, S. (2013). Border Malaria Associated With Multidrug Resistance on Thailand-Myanmar and

Thailand-Cambodia Borders: Transmission Dynamic, Vulnerability, and Surveillance. *Biomed Research International*, *2013*, 1-13. https://doi.org/10.1155/2013/363417
Biganzoli, P., Ferreyra, L., Sicilia, P., Carabajal, C. C., Frattari, S. R. d., Littvik, A., Nates, S., & Paván, J. V. (2010). IgG Subclasses and DNA Detection of HHV-6 and HHV-7 in Healthy Individuals. *J Med Virol*, *82*(10), 1679-1683. https://doi.org/10.1002/jmv.21880
Blair, J. E. (2007). Coccidioidomycosis in Patients Who Have Undergone Transplantation. *Annals of the New York Academy of Sciences*, *1111*(1), 365-376. https://doi.org/10.1196/annals.1406.009
Blois, S., Mallus, F., Liciardi, M., Pilo, C., Camboni, T., Macera, L., Maggi, F., & Manzin, A. (2014). High Prevalence of Co-Infection With Multiple Torque Teno Sus Virus Species in Italian Pig Herds. *PLoS One*, *9*(11), e113720. https://doi.org/10.1371/journal.pone.0113720
Blower, K., & Keyt, H. (2016). Don't Let the Burk Holderia You Back! *Chest Journal*, *150*(4), 178A. https://doi.org/10.1016/j.chest.2016.08.187
Blyth, C. C., Middleton, P. G., Harun, A., Sorrell, T. C., Meyer, W., & Chen, S. C. (2010). Clinical Associations and Prevalence Of<i>Scedosporium</I>spp. In Australian Cystic Fibrosis Patients: Identification of Novel Risk Factors? *Medical Mycology*, *48*(O1), S37-S44. https://doi.org/10.3109/13693786.2010.500627
Boekhout, T., Theelen, B., Diaz, M. R., Fell, J. W., Hop, W. C., Abeln, E. C. A., Dromer, F., & Meyer, W. (2001). Hybrid Genotypes in the Pathogenic Yeast Cryptococcus Neoformans. *Microbiology*, *147*(4), 891-907. https://doi.org/10.1099/00221287-147-4-891
Boer, S. L. d., Meijer, R. P., Iperen, G. G. v., Harkel, A. D. t., Gideon, J. d. M. S., Straver, B., Rammeloo, L., Tanke, R. B., Jeroen, J. A. v. K., & Dalinghaus, M. (2014). Evaluation of the Diagnostic Work-Up in Children With Myocarditis and Idiopathic Dilated Cardiomyopathy. *Pediatric Cardiology*, *36*(2), 409-416. https://doi.org/10.1007/s00246-014-1022-3
Bofill-Mas, S. l., Rodríguez-Manzano, J., Calgua, B., Carratalà, A., & Gironés, R. (2010). Newly Described Human Polyomaviruses Merkel Cell, KI and WU Are Present in Urban Sewage and May Represent Potential Environmental Contaminants. *Virology Journal*, *7*(1). https://doi.org/10.1186/1743-422x-7-141
Bogaart, L. v. d., & Manuel, O. (2022). Antibiotic Therapy for Difficult-to-Treat Infections in Lung Transplant Recipients: A Practical Approach. *Antibiotics*, *11*(5), 612. https://doi.org/10.3390/antibiotics11050612
Boonjakuakul, J. K., Gerns, H. L., Chen, Y.-T., Hicks, L. D., Minnick, M. F., Dixon, S. E., Hall, S. C., & Koehler, J. E. (2007). Proteomic and Immunoblot Analyses Of <i>Bartonella Quintana</I> Total Membrane Proteins Identify Antigens Recognized by Sera From Infected Patients. *Infect Immun*, *75*(5), 2548-2561. https://doi.org/10.1128/iai.01974-06
Boothroyd, J. C., & Dubremetz, J. F. (2008). Kiss and Spit: The Dual Roles of Toxoplasma Rhoptries. *Nature Reviews Microbiology*, *6*(1), 79-88. https://doi.org/10.1038/nrmicro1800
Borman, A. M., Székely, A., Linton, C. J., Palmer, M. D., Brown, P., & Johnson, E. M. (2013). Epidemiology, Antifungal Susceptibility, and Pathogenicity of Candida Africana Isolates From the United Kingdom. *Journal of Clinical Microbiology*, *51*(3), 967-972. https://doi.org/10.1128/jcm.02816-12
Botto, S., Gustin, J. K., & Moses, A. V. (2017). The Heme Metabolite Carbon Monoxide Facilitates KSHV Infection by Inhibiting TLR4 Signaling in Endothelial Cells. *Frontiers in Microbiology*, *8*. https://doi.org/10.3389/fmicb.2017.00568

Boukaabar, M., Oduro, B., & Chataa, P. (2024). Congenital Transmission of Chagas Disease: The Role of Newborn Therapy on the Disease's Dynamics. https://doi.org/10.1101/2024.02.18.24302989
Boussaud, V., Guillemain, R., Grenet, D., Coley, N., Souilamas, R., Bonnette, P., & Stern, M. (2008). Clinical Outcome Following Lung Transplantation in Patients With Cystic Fibrosis Colonised With Burkholderia Cepacia Complex: Results From Two French Centres. *Thorax*, *63*(8), 732-737. https://doi.org/10.1136/thx.2007.089458
Breitschwerdt, E. B., Maggi, R. G., Farmer, P., & Mascarelli, P. E. (2010). Molecular Evidence of Perinatal Transmission Of <i>Bartonella Vinsonii</I> Subsp. <i>berkhoffii</I> And <i>Bartonella Henselae</I> to a Child. *Journal of Clinical Microbiology*, *48*(6), 2289-2293. https://doi.org/10.1128/jcm.00326-10
Breloer, M., & Abraham, D. (2016). <i>Strongyloides</I> Infection in Rodents: Immune Response and Immune Regulation. *Parasitology*, *144*(3), 295-315. https://doi.org/10.1017/s0031182016000111
Brites, D., & Gagneux, S. (2015). Co-evolution of Mycobacterium tuberculosis and Homo sapiens. *Immunol Rev*, *264*(1), 6-24. https://doi.org/10.1111/imr.12264
Brito, L. F., Brune, W., & Stahl, F. R. (2019). Cytomegalovirus (CMV) Pneumonitis: Cell Tropism, Inflammation, and Immunity. *International Journal of Molecular Sciences*, *20*(16), 3865. https://doi.org/10.3390/ijms20163865
Broliden, K. (2001). Parvovirus B19 Infection in Pediatric Solid-organ and Bone Marrow Transplantation. *Pediatric Transplantation*, *5*(5), 320-330. https://doi.org/10.1034/j.1399-3046.2001.00035.x
Bronnimann, D. A., Adam, R. D., Galgiani, J. N., Habib, M. P., Petersen, E., Porter, B., & Bloom, J. W. (1987). Coccidioidomycosis in the Acquired Immunodeficiency Syndrome. *Annals of Internal Medicine*, *106*(3), 372-379. https://doi.org/10.7326/0003-4819-106-3-372
Broussard, G., & Damania, B. (2020). KSHV: Immune Modulation and Immunotherapy. *Frontiers in Immunology*, *10*. https://doi.org/10.3389/fimmu.2019.03084
Brown, M. M., Cartledge, J. D., & Miller, R. F. (2006). Dissemination of <i>Strongyloides Stercoralis</I> as an Immune Restoration Phenomenon in an HIV-1-infected Man on Antiretroviral Therapy. *International Journal of STD & Aids*, *17*(8), 560-561. https://doi.org/10.1258/095646206778145712
Brown-Elliott, B. A., Brown, J. M., Conville, P. S., & Wallace, R. J. (2006). Clinical and Laboratory Features of The<i>Nocardia</I>spp. Based on Current Molecular Taxonomy. *Clinical Microbiology Reviews*, *19*(2), 259-282. https://doi.org/10.1128/cmr.19.2.259-282.2006
Bruce, M. G., & Maaroos, H. I. (2008). Epidemiology of <i>Helicobacter Pylori</I> Infection. *Helicobacter*, *13*(s1), 1-6. https://doi.org/10.1111/j.1523-5378.2008.00631.x
Brumfield, K. D., Chen, A. J., Gangwar, M., Usmani, M., Hasan, N. A., Jutla, A. S., Huq, A., & Colwell, R. R. (2023). Environmental Factors Influencing Occurrence of Vibrio parahaemolyticus and Vibrio vulnificus. *Appl Environ Microbiol*, *89*(6), e0030723. https://doi.org/10.1128/aem.00307-23
Burette, M., & Bonazzi, M. (2020). From Neglected to Dissected: How Technological Advances Are Leading the Way to the Study Of<i>Coxiella Burnetii</I>pathogenesis. *Cellular Microbiology*, *22*(4). https://doi.org/10.1111/cmi.13180

Busselman, R. E., & Hamer, S. A. (2022). Chagas Disease Ecology in the United States: Recent Advances in Understanding<i>Trypanosoma Cruzi</I>Transmission Among Triatomines, Wildlife, and Domestic Animals and a Quantitative Synthesis of Vector–Host Interactions. *Annual Review of Animal Biosciences*, *10*(1), 325-348. https://doi.org/10.1146/annurev-animal-013120-043949

Cafarchia, C., Paradies, R., Figueredo, L. A., Iatta, R., Desantis, S., Antonio Vito Francesco Di, B., Zizzo, N., & Diepeningen, A. D. v. (2019). <i>Fusarium Spp</I>. In Loggerhead Sea Turtles (<i>Caretta Caretta</I>): From Colonization to Infection. *Veterinary Pathology*, *57*(1), 139-146. https://doi.org/10.1177/0300985819880347

Cai, Q., Verma, S. C., Lu, J., & Robertson, E. S. (2010). Molecular Biology of Kaposi's Sarcoma-Associated Herpesvirus and Related Oncogenesis. 87-142. https://doi.org/10.1016/b978-0-12-385032-4.00003-3

Canning, J. S., Laucirica, D. R., Ling, K. M., Nicol, M. P., Stick, S. M., & Kicic, A. (2024). Phage Therapy to Treat Cystic Fibrosis Burkholderia Cepacia Complex Lung Infections: Perspectives and Challenges. *Frontiers in Microbiology*, *15*. https://doi.org/10.3389/fmicb.2024.1476041

Cannon, M. J., Schmid, D. S., & Hyde, T. B. (2010). Review of Cytomegalovirus Seroprevalence and Demographic Characteristics Associated With Infection. *Reviews in Medical Virology*, *20*(4), 202-213. https://doi.org/10.1002/rmv.655

Cao, J. Y., Mansouri, S., & Frappier, L. (2012). Changes in the Nasopharyngeal Carcinoma Nuclear Proteome Induced by the EBNA1 Protein of Epstein-Barr Virus Reveal Potential Roles for EBNA1 in Metastasis and Oxidative Stress Responses. *J Virol*, *86*(1), 382-394. https://doi.org/10.1128/jvi.05648-11

Caraher, E., Reynolds, G., Murphy, P. G., McClean, S., & Callaghan, M. (2007). Comparison of Antibiotic Susceptibility of Burkholderia Cepacia Complex Organisms When Grown Planktonically or as Biofilm in Vitro. *European Journal of Clinical Microbiology & Infectious Diseases*, *26*(3), 213-216. https://doi.org/10.1007/s10096-007-0256-x

Carraro, D. d. S., Carraro, R. M., Campos, S. V., Iuamoto, L. R., Karina Andrighetti de Oliveira, B., Oliveira, L. C. d., Sabino, É. C., Rossi, F., & Pêgo-Fernandes, P. M. (2018). Burkholderia Cepacia, Cystic Fibrosis and Outcomes Following Lung Transplantation: Experiences From a Single Center in Brazil. *Clinics*, *73*, e166. https://doi.org/10.6061/clinics/2018/e166

Carter, R., & Mendis, K. (1992). Transmission Immunity in Malaria: Reflections on the Underlying Immune Mechanisms During Natural Infections and Following Artificial Immunization. *Memórias Do Instituto Oswaldo Cruz*, *87*(suppl 3), 169-173. https://doi.org/10.1590/s0074-02761992000700027

Carvalho, E. M., & Porto, A. D. F. (2004). Epidemiological and Clinical Interaction Between HTLV-1 and <i>Strongyloides Stercoralis</I>. *Parasite Immunology*, *26*(11-12), 487-497. https://doi.org/10.1111/j.0141-9838.2004.00726.x

Caselli, E., D'Accolti, M., Caccuri, F., Soffritti, I., Gentili, V., Bortolotti, D., Rotola, A., Cassai, E., Fiorentini, S., Zani, A., Caruso, A., Rizzo, R., & Luca, D. D. (2020). The U94 Gene of Human Herpesvirus 6: A Narrative Review of Its Role and Potential Functions. *Cells*, *9*(12), 2608. https://doi.org/10.3390/cells9122608

Castelli, M. V., Buitrago, M. J., Bernal-Martínez, L., Gómez-López, A., Rodríguez-Tudela, J. L., & Cuenca-Estrella, M. (2008). Development and Validation of a Quantitative PCR Assay for

Diagnosis of Scedosporiosis. *Journal of Clinical Microbiology*, *46*(10), 3412-3416. https://doi.org/10.1128/jcm.00046-08

Castro, F. A., Domínguez, A., Püschel, K., Wyngard, V. V. D., Snijders, P. J., Franceschi, S., Pawlita, M., & Ferreccio, C. (2014). Serological Prevalence and Persistence of High-Risk Human Papillomavirus Infection Among Women in Santiago, Chile. *BMC Infectious Diseases*, *14*(1). https://doi.org/10.1186/1471-2334-14-361

Castro, G. d., Cervi, M. C., & Martínez, R. (2006). Vertical Transmission of Cryptococcus Neoformans From a Mother Coinfected With Human Immunodeficiency Virus: Case Report. *Revista Da Sociedade Brasileira De Medicina Tropical*, *39*(5), 501-503. https://doi.org/10.1590/s0037-86822006000500016

Catchpole, A., Zabriskie, B. N., Bassett, P., Embley, B., White, D. W., Gale, S. D., & Dawson, W. O. (2023). Association Between Toxoplasma Gondii Infection and Type-1 Diabetes Mellitus: A Systematic Review and Meta-Analysis. *International Journal of Environmental Research and Public Health*, *20*(5), 4436. https://doi.org/10.3390/ijerph20054436

Celina, S. S., & Černý, J. (2022). Coxiella Burnetii in Ticks, Livestock, Pets and Wildlife: A Mini-Review. *Frontiers in Veterinary Science*, *9*. https://doi.org/10.3389/fvets.2022.1068129

Chadsuthi, S., Bicout, D., Wiratsudakul, A., Suwancharoen, D., Petkanchanapong, W., Modchang, C., Triampo, W., Ratanakorn, P., & Chalvet-Monfray, K. (2017). Investigation on Predominant Leptospira Serovars and Its Distribution in Humans and Livestock in Thailand, 2010-2015. *Plos Neglected Tropical Diseases*, *11*(2), e0005228. https://doi.org/10.1371/journal.pntd.0005228

Chadsuthi, S., Chalvet-Monfray, K., Wiratsudakul, A., Suwancharoen, D., & Cappelle, J. (2018). A Remotely Sensed Flooding Indicator Associated With Cattle and Buffalo Leptospirosis Cases in Thailand 2011–2013. *BMC Infectious Diseases*, *18*(1). https://doi.org/10.1186/s12879-018-3537-3

Chai, Y., Ye, C., Zhang, G., & Wang, J. (2015). Histoplasmosis Presenting With Solitary Pulmonary Nodule: Two Cases Mimicking Pulmonary Metastases. *Nigerian Journal of Clinical Practice*, *18*(2), 304. https://doi.org/10.4103/1119-3077.151075

Chan, K., Nasereddin, T., Alter, L., Centurion-Lara, A., Giacani, L., & Parveen, N. (2016). Treponema Pallidum Lipoprotein TP0435 Expressed in Borrelia Burgdorferi Produces Multiple Surface/Periplasmic Isoforms and Mediates Adherence. *Scientific Reports*, *6*(1). https://doi.org/10.1038/srep25593

Chang, Y., & Moore, P. S. (2012). Merkel Cell Carcinoma: A Virus-Induced Human Cancer. *Annual Review of Pathology Mechanisms of Disease*, *7*(1), 123-144. https://doi.org/10.1146/annurev-pathol-011110-130227

Chaparro, C., Maurer, J. R., Cano, C., Krajden, M., Chan, C., Winton, T., Keshavjee, S., Scavuzzo, M., Tullis, E., Hutcheon, M., & Kesten, S. (2001). Infection With <i>Burkholderia Cepacia</I> in Cystic Fibrosis. *American Journal of Respiratory and Critical Care Medicine*, *163*(1), 43-48. https://doi.org/10.1164/ajrccm.163.1.9811076

Charles, R. C., Nakajima, R., Liang, L., Jasinskas, A., Berger, A., Leung, D. T., Kelly, M., Xu, P., Kováč, P., Giffen, S. R., Harbison, J. D., Chowdhury, F., Khan, A. I., Calderwood, S. B., Bhuiyan, T. R., Harris, J. B., Felgner, P. L., Qadri, F., & Ryan, E. T. (2017). Plasma and Mucosal Immunoglobulin M, Immunoglobulin A, and Immunoglobulin G Responses to the Vibrio Cholerae O1 Protein Immunome in Adults With Cholera in Bangladesh. *The Journal of Infectious Diseases*, *216*(1), 125-134. https://doi.org/10.1093/infdis/jix253

Chen, I. C., Li, W.-C., Hong, Y. C., Shie, S. S., Fann, W. C., & Hsiao, C. T. (2011). The Microbiological Profile and Presence of Bloodstream Infection Influence Mortality Rates in Necrotizing Fasciitis. *Critical Care*, *15*(3). https://doi.org/10.1186/cc10278

Chen, J., Du, Y., Guan, C.-H., Fan, H., Liu, Y.-A., Wang, T., Li, X., & Chen, G. (2023). Extracellular Vesicles Derived From <i>Plasmodium</I>-Infected Hosts As Stimuli of "Trained" Innate Immunity. *Current Medicinal Chemistry*, *30*(39), 4450-4465. https://doi.org/10.2174/0929867330666230207115157

Chen, L., Christian, D. A., Kochanowsky, J. A., Phan, A. T., Clark, J. T., Wang, S., Berry, C. T., Oh, J. Y., Chen, X. G., Roos, D. S., Beiting, D. P., Koshy, A. A., & Hunter, C. A. (2020). The <i>Toxoplasma Gondii</I> Virulence Factor ROP16 Acts in Cis and Trans, and Suppresses T Cell Responses. *J Exp Med*, *217*(3). https://doi.org/10.1084/jem.20181757

Chisaka, H., Morita, E., Yaegashi, N., & Sugamura, K. (2003). Parvovirus B19 and the Pathogenesis of Anaemia. *Reviews in Medical Virology*, *13*(6), 347-359. https://doi.org/10.1002/rmv.395

Chisu, V., Mura, L., Foxi, C., & Masala, G. (2021). Coxiellaceae in Ticks From Human, Domestic and Wild Hosts From Sardinia, Italy: High Diversity of Coxiella-Like Endosymbionts. *Acta Parasitologica*, *66*(2), 654-663. https://doi.org/10.1007/s11686-020-00324-w

Cho, J. S., Na, S., Park, M. S., So, Y., Lee, B., Koh, S. O., & Paik, H. C. (2013). A Case Of<i>Burkholderia Cepacia</I>Pneumonia After Lung Transplantation in a Recipient Without Cystic Fibrosis. *The Korean Journal of Critical Care Medicine*, *28*(3), 187. https://doi.org/10.4266/kjccm.2013.28.3.187

Chomel, B. B., & Kasten, R. W. (2010). Bartonellosis, an Increasingly Recognized Zoonosis. *Journal of Applied Microbiology*, *109*(3), 743-750. https://doi.org/10.1111/j.1365-2672.2010.04679.x

Chotirmall, S. H., Al-Alawi, M., Mirković, B., Lavelle, G. M., Logan, P. M., Greene, C. M., & McElvaney, N. G. (2013). <i>Aspergillus</I>-Associated Airway Disease, Inflammation, and the Innate Immune Response. *Biomed Research International*, *2013*, 1-14. https://doi.org/10.1155/2013/723129

Chow, J., Golan, Y., Ruthazer, R., Karchmer, A. W., Carmeli, Y., Lichtenberg, D. A., Chawla, V., Young, J., & Hadley, S. (2008). Factors Associated With Candidemia Caused by Non-<i>albicans Candida</I>Species Versus<i>Candida Albicans</I>in the Intensive Care Unit. *Clinical Infectious Diseases*, *46*(8), 1206-1213. https://doi.org/10.1086/529435

Cimino, P. J., Robirds, D., Tripp, S. R., Pfeifer, J. D., Abel, H., & Duncavage, E. J. (2014). Retinoblastoma Gene Mutations Detected by Whole Exome Sequencing of Merkel Cell Carcinoma. *Modern Pathology*, *27*(8), 1073-1087. https://doi.org/10.1038/modpathol.2013.235

Coenye, T. (2010). Social Interactions in The <i>Burkholderia Cepacia</I> Complex: Biofilms and Quorum Sensing. *Future Microbiology*, *5*(7), 1087-1099. https://doi.org/10.2217/fmb.10.68

Colmenares, M. a., Constant, S. L., Kima, P. E., & McMahon-Pratt, D. (2002). <i>Leishmania Pifanoi</I>Pathogenesis: Selective Lack of a Local Cutaneous Response in the Absence of Circulating Antibody. *Infect Immun*, *70*(12), 6597-6605. https://doi.org/10.1128/iai.70.12.6597-6605.2002

Colmenares, M. a., Kar, S., Goldsmith-Pestana, K., & McMahon-Pratt, D. (2002). Mechanisms of Pathogenesis: Differences Amongst Leishmania Species. *Transactions of the Royal Society of Tropical Medicine and Hygiene*, *96*, S3-S7. https://doi.org/10.1016/s0035-9203(02)90044-1

Colombe, S., Machemba, R., Mtenga, B., Lutonja, P., Kalluvya, S. E., Dood, C. J. d., Hoekstra, P. T., Dam, G. J. v., Paul, L. A. M. C., Urassa, M., Changalucha, J., Todd, J., & Downs, J. A. (2018). Impact of Schistosome Infection on Long-Term HIV/AIDS Outcomes. *Plos Neglected Tropical Diseases*, *12*(7), e0006613. https://doi.org/10.1371/journal.pntd.0006613

Comas, I., Coscolla, M., Luo, T., Borrell, S., Holt, K. E., Kato-Maeda, M., Parkhill, J., Malla, B., Berg, S., Thwaites, G., Yeboah-Manu, D., Bothamley, G., Mei, J., Wei, L., Bentley, S., Harris, S. R., Niemann, S., Diel, R., Aseffa, A.,…Gagneux, S. (2013). Out-of-Africa migration and Neolithic coexpansion of Mycobacterium tuberculosis with modern humans. *Nat Genet*, *45*(10), 1176-1182. https://doi.org/10.1038/ng.2744

Comoli, P., Binggeli, S., Ginevri, F., & Hirsch, H. H. (2006). Polyomavirus-associated Nephropathy: Update on BK Virus-specific Immunity. *Transplant Infectious Disease*, *8*(2), 86-94. https://doi.org/10.1111/j.1399-3062.2006.00167.x

Cook, E. A. J., Glanville, W. A. d., Thomas, L. F., Kariuki, S., Bronsvoort, M., & Fèvre, E. M. (2016). Risk Factors for Leptospirosis Seropositivity in Slaughterhouse Workers in Western Kenya. *Occupational and Environmental Medicine*, *74*(5), 357-365. https://doi.org/10.1136/oemed-2016-103895

Cook, J., Speybroeck, N., Sochanta, T., Heng, S., Sokny, M., Claes, F., Lemmens, K., Theisen, M., Soares, I. S., D'Alessandro, U., Coosemans, M., & Erhart, A. (2012). Sero-Epidemiological Evaluation of Changes in Plasmodium Falciparum and Plasmodium Vivax Transmission Patterns Over the Rainy Season in Cambodia. *Malaria Journal*, *11*(1). https://doi.org/10.1186/1475-2875-11-86

Cooper, C. J., Said, S., Popp, M., Alkhateeb, H., Rodríguez, C., Aguilar, M. P., & Alozie, O. (2014). A Complicated Case of an Immunocompetent Patient With Disseminated Nocardiosis. *Infectious Disease Reports*, *6*(1), 5327. https://doi.org/10.4081/idr.2014.5327

Corcoran, A., & Doyle, S. (2004). Advances in the Biology, Diagnosis and Host–pathogen Interactions of Parvovirus B19. *Journal of Medical Microbiology*, *53*(6), 459-475. https://doi.org/10.1099/jmm.0.05485-0

Cornberg, M., Sheridan, B. S., Saccoccio, F. M., Brehm, M. A., & Selin, L. K. (2007). Protection Against Vaccinia Virus Challenge by CD8 Memory T Cells Resolved by Molecular Mimicry. *J Virol*, *81*(2), 934-944. https://doi.org/10.1128/jvi.01280-06

Coscoy, L. (2007). Immune Evasion by Kaposi's Sarcoma-Associated Herpesvirus. *Nature Reviews Immunology*, *7*(5), 391-401. https://doi.org/10.1038/nri2076

Costa, C. (2012). Polyomavirus-Associated Nephropathy. *World Journal of Transplantation*, *2*(6), 84. https://doi.org/10.5500/wjt.v2.i6.84

Countryman, J., & Miller, G. (1985). Activation of Expression of Latent Epstein-Barr Herpesvirus After Gene Transfer With a Small Cloned Subfragment of Heterogeneous Viral DNA. *Proceedings of the National Academy of Sciences*, *82*(12), 4085-4089. https://doi.org/10.1073/pnas.82.12.4085

Coursaget, P., Samimi, M., Nicol, J., Gardair, C., & Touzé, A. (2013). Human Merkel Cell Polyomavirus: Virological Background and Clinical Implications. *Apmis*, *121*(8), 755-769. https://doi.org/10.1111/apm.12122

Cox, R. A., & Magee, D. M. (2004). Coccidioidomycosis: Host Response and Vaccine Development. *Clinical Microbiology Reviews*, *17*(4), 804-839. https://doi.org/10.1128/cmr.17.4.804-839.2004

Cramer, R. A., Rivera, A., & Hohl, T. M. (2011). Immune Responses Against Aspergillus Fumigatus. *Current Opinion in Infectious Diseases*, *24*(4), 315-322. https://doi.org/10.1097/qco.0b013e328348b159

Crandall, P. G., Ricke, S. C., O'Bryan, C. A., & Parrish, N. (2012). <i>In Vitro</I>effects of Citrus Oils Against<i>Mycobacterium Tuberculosis</I>and Non-Tuberculous<i>Mycobacteria</I>of Clinical Importance. *Journal of Environmental Science and Health Part B*, *47*(7), 736-741. https://doi.org/10.1080/03601234.2012.669331

Cutler, J. E., Corti, M., Lambert, P., Ferris, M. J., & Xin, H. (2011). Horizontal Transmission of Candida Albicans and Evidence of a Vaccine Response in Mice Colonized With the Fungus. *PLoS One*, *6*(7), e22030. https://doi.org/10.1371/journal.pone.0022030

Daccò, V., Claut, L., Piconi, S., Castellazzi, L., Garbarino, F., Teri, A., & Colombo, C. (2019). Successful Ceftazidime-avibactam Treatment of Post-surgery <i>Burkholderia Multivorans</I> Genomovar II Bacteremia and Brain Abscesses in a Young Lung Transplanted Woman With Cystic Fibrosis. *Transplant Infectious Disease*, *21*(3). https://doi.org/10.1111/tid.13082

Dall, A., & Hariharan, S. (2008). BK Virus Nephritis After Renal Transplantation. *Clinical Journal of the American Society of Nephrology*, *3*(Supplement_2), S68-S75. https://doi.org/10.2215/cjn.02770707

Damania, B., & Münz, C. (2019). Immunodeficiencies That Predispose to Pathologies by Human Oncogenic Γ-Herpesviruses. *Fems Microbiology Reviews*, *43*(2), 181-192. https://doi.org/10.1093/femsre/fuy044

Das, P., Lahiri, A., Lahiri, A., & Chakravortty, D. (2010). Modulation of the Arginase Pathway in the Context of Microbial Pathogenesis: A Metabolic Enzyme Moonlighting as an Immune Modulator. *PLOS Pathogens*, *6*(6), e1000899. https://doi.org/10.1371/journal.ppat.1000899

Dauner, J. G., Pan, Y., Hildesheim, A., Harro, C., & Pinto, L. A. (2010). Characterization of the HPV-specific Memory B Cell and Systemic Antibody Responses in Women Receiving an Unadjuvanted HPV16 L1 VLP Vaccine. *Vaccine*, *28*(33), 5407-5413. https://doi.org/10.1016/j.vaccine.2010.06.018

Decker, D. B., Karam, J. A., & Wilcox, D. T. (2009). Pediatric Hemorrhagic Cystitis. *Journal of Pediatric Urology*, *5*(4), 254-264. https://doi.org/10.1016/j.jpurol.2009.02.199

Degrandi, D., Kravets, E., Konermann, C., Beuter-Gunia, C., Klümpers, V., Lahme, S., Rasch, E., Mausberg, A. K., Beer-Hammer, S., & Pfeffer, K. (2012). Murine Guanylate Binding Protein 2 (mGBP2) Controls<i>Toxoplasma Gondii</I>replication. *Proceedings of the National Academy of Sciences*, *110*(1), 294-299. https://doi.org/10.1073/pnas.1205635110

Delbue, S., Ferraresso, M., Ghio, L., Carloni, C., Carluccio, S., Belingheri, M., Edefonti, A., & Ferrante, P. (2013). A Review on JC Virus Infection in Kidney Transplant Recipients. *Clinical and Developmental Immunology*, *2013*, 1-7. https://doi.org/10.1155/2013/926391

Dembinski, J. (2003). Long Term Follow Up of Serostatus After Maternofetal Parvovirus B19 Infection. *Archives of Disease in Childhood*, *88*(3), 219-221. https://doi.org/10.1136/adc.88.3.219

Demir, E. T., Ceyhan, M., Şimşek, M., Gündüz, T., Arlıer, S., Aytaç, R., Aycan, A. E., & Gürbüz, V. (2012). The Prevalence of Different HPV Types in Turkish Women With a Normal Pap Smear. *J Med Virol*, *84*(8), 1242-1247. https://doi.org/10.1002/jmv.23333

Denkers, E. (2003). From Cells to Signaling Cascades: Manipulation of Innate Immunity By<i>Toxoplasma Gondii</I>. *Fems Immunology & Medical Microbiology*, *39*(3), 193-203. https://doi.org/10.1016/s0928-8244(03)00279-7
Denner, J., Bigley, T. M., Phan, T. L., Zimmermann, C., Zhou, X., & Kaufer, B. B. (2019). Comparative Analysis of Roseoloviruses in Humans, Pigs, Mice, and Other Species. *Viruses*, *11*(12), 1108. https://doi.org/10.3390/v11121108
Deorukhkar, S., Saini, S., & Mathew, S. (2014). Non-<i>albicans Candida</I>Infection: An Emerging Threat. *Interdisciplinary Perspectives on Infectious Diseases*, *2014*, 1-7. https://doi.org/10.1155/2014/615958
Dharnidharka, V. R., Abdulnour, H. A., & Araya, C. E. (2011). The BK Virus in Renal Transplant Recipients—review of Pathogenesis, Diagnosis, and Treatment. *Pediatric Nephrology*, *26*(10), 1763-1774. https://doi.org/10.1007/s00467-010-1716-6
Diaz, J. H. (2018). Environmental and Wilderness-Related Risk Factors for Histoplasmosis: More Than Bats in Caves. *Wilderness and Environmental Medicine*, *29*(4), 531-540. https://doi.org/10.1016/j.wem.2018.06.008
Dogra, V., Sinha, A. K., Saxena, R., & Talwar, D. (2016). Aspergillus March: From ABPA to Aspergilloma to Subacute Invasive Aspergillosis. *Allergy Asthma & Clinical Immunology*, *12*(1). https://doi.org/10.1186/s13223-016-0170-9
Doja, A., Bitnun, A., Jones, E., Richardson, S. E., Tellier, R., Petric, M., Heurter, H., & MacGregor, D. (2006). Pediatric Epstein-Barr Virus—Associated Encephalitis: 10-Year Review. *Journal of Child Neurology*, *21*(5), 384-391. https://doi.org/10.1177/08830738060210051101
Dong, K., Jiang, Z., Zhang, J., Qin, H., Chen, J., & Chen, Q. (2022). The Role of SIRT1 in the Process of Toxoplasma Gondii Infection of RAW 264.7 Macrophages. *Frontiers in Microbiology*, *13*. https://doi.org/10.3389/fmicb.2022.1017696
Donovan, M. J., Maciuba, B. Z., Mahan, C. E., & McDowell, M. A. (2009). Leishmania Infection Inhibits Cycloheximide-Induced Macrophage Apoptosis in a Strain-Dependent Manner. *Experimental Parasitology*, *123*(1), 58-64. https://doi.org/10.1016/j.exppara.2009.05.012
Dorn, P. L., Monroy, M. C., & Stevens, L. (2022). Sustainable, Integrated Control of Native Vectors: The Case of Chagas Disease in Central America. *Frontiers in Tropical Diseases*, *3*. https://doi.org/10.3389/fitd.2022.971000
Dropulic, L. K., & Jones, R. J. (2007). Polyomavirus BK Infection in Blood and Marrow Transplant Recipients. *Bone Marrow Transplantation*, *41*(1), 11-18. https://doi.org/10.1038/sj.bmt.1705886
Duan, M. M., Xu, R., Yuan, C. X., Li, Y. Y., Liu, Q., Cheng, G. F., Lin, J., & Feng, X. (2015). SjHSP70, a Recombinant Schistosoma Japonicum Heat Shock Protein 70, Is Immunostimulatory and Induces Protective Immunity Against Cercarial Challenge in Mice. *Parasitology Research*, *114*(9), 3415-3429. https://doi.org/10.1007/s00436-015-4567-z
Eddens, T., & Kolls, J. K. (2014). Pathological and Protective Immunity to Pneumocystis Infection. *Seminars in Immunopathology*, *37*(2), 153-162. https://doi.org/10.1007/s00281-014-0459-z
Elfaki, M. G., Alaidan, A., & Alhokail, A. (2015). Host Response to Brucella Infection: Review and Future Perspective. *The Journal of Infection in Developing Countries*, *9*(07), 697-701. https://doi.org/10.3855/jidc.6625
Elia, F., Villani, S., Ambrogi, F., Signorini, L., Dallari, S., Binda, S., Prímache, V., Pellegrinelli, L., Ferrante, P., & Delbue, S. (2016). JC Virus Infection Is Acquired Very Early in Life: Evidence From

a Longitudinal Serological Study. *Journal of Neurovirology*, *23*(1), 99-105. https://doi.org/10.1007/s13365-016-0477-9
Eliassen, E., Luca, D. D., Rizzo, R., & Barão, I. (2017). The Interplay Between Natural Killer Cells and Human Herpesvirus-6. *Viruses*, *9*(12), 367. https://doi.org/10.3390/v9120367
Elmi, A., Bansal, D., Acharya, A., Skariah, S., Dargham, S. R., Abu-Raddad, L. J., Mohamed-Nady, N., Amuna, P., Althani, A., & Sultan, A. A. (2017). Human Papillomavirus (HPV) Infection: Molecular Epidemiology, Genotyping, Seroprevalence and Associated Risk Factors Among Arab Women in Qatar. *PLoS One*, *12*(1), e0169197. https://doi.org/10.1371/journal.pone.0169197
Enders, M., Weidner, A., Zoellner, I., Searle, K., & Enders, G. (2004). Fetal Morbidity and Mortality After Acute Human Parvovirus B19 Infection in Pregnancy: Prospective Evaluation of 1018 Cases. *Prenatal Diagnosis*, *24*(7), 513-518. https://doi.org/10.1002/pd.940
Engelthaler, D. M., Roe, C. C., Hepp, C. M., Teixeira, M. d. M., Driebe, E. M., Schupp, J. M., Gade, L., Waddell, V., Komatsu, K., Arathoon, E., Logemann, H., Thompson, G. R., Chiller, T., Barker, B. M., Keim, P., & Litvintseva, A. P. (2016). Local Population Structure and Patterns of Western Hemisphere Dispersal For <i>Coccidioides</I> Spp., the Fungal Cause of Valley Fever. *Mbio*, *7*(2). https://doi.org/10.1128/mbio.00550-16
Erol, İ., & Alehan, F. (2006). Refractory Status Epilepticus Owing to Human Parvovirus B19 Encephalitis in a Child. *Journal of Child Neurology*, *21*(9), 820-822. https://doi.org/10.1177/08830738060210092301
Esakkiammal, M., & Rajakumari, D. (2025). Pigeon Droppings as a Cryptococcus Reservoir: A Review. *Uttar Pradesh Journal of Zoology*, *46*(6), 169-177. https://doi.org/10.56557/upjoz/2025/v46i64854
Eskra, L., Canavessi, Á. M. O., Carey, M., & Splitter, G. A. (2001). <i>Brucella Abortus</I> Genes Identified Following Constitutive Growth and Macrophage Infection. *Infect Immun*, *69*(12), 7736-7742. https://doi.org/10.1128/iai.69.12.7736-7742.2001
Fadipe, D. O., Omotuyi, O. I., & Olajubu, F. A. (2024). The Carrier Rate of Treponema Pallidum (Syphilis) Antibody Among Women in Some Selected Locations in Ondo State, Nigeria. *Sokoto Journal of Medical Laboratory Science*, *9*(3), 283-288. https://doi.org/10.4314/sokjmls.v9i3.29
Falkinham, J. O., Norton, C. D., & LeChevallier, M. W. (2001). Factors Influencing Numbers Of <i>Mycobacterium Avium</I> , <i>Mycobacterium Intracellulare</I> , and Other Mycobacteria in Drinking Water Distribution Systems. *Applied and Environmental Microbiology*, *67*(3), 1225-1231. https://doi.org/10.1128/aem.67.3.1225-1231.2001
Fan, S. R., Liu, P., & Li, J. W. (2008). Clinical Characteristics of Vulvovaginal Candidiasis and Antifungal Susceptibilities of <i>Candida</I> Species Isolates Among Patients in Southern China From 2003 to 2006. *Journal of Obstetrics and Gynaecology Research*, *34*(4), 561-566. https://doi.org/10.1111/j.1447-0756.2008.00817.x
Fastenackels, S., Bayard, C., Larsen, M., Magnier, P., Bonnafous, P., Seddiki, N., Appay, V., Gautheret-Dejean, A., & Sauce, D. (2019). Phenotypic and Functional Differences Between Human Herpesvirus 6- And Human Cytomegalovirus-Specific T Cells. *J Virol*, *93*(13). https://doi.org/10.1128/jvi.02321-18
Ferreira, S. d. C., Nóbrega, F. J. F., Araújo, R. C., Santana, R. d. C., Ramalho, L. N. Z., Martinelli, A. L., & Troncon, L. E. A. (2019). P023 Disseminated Histoplasmosis in a Patient With Crohn's Disease Under Tumor Necrosis Factor Blocker Therapy: An Unusual Case. *The American Journal of Gastroenterology*, *114*(1), S6-S7. https://doi.org/10.14309/01.ajg.0000613060.27726.48

Fields, B. S., Benson, R. F., & Besser, R. E. (2002). <i>Legionella</I> and Legionnaires' Disease: 25 Years of Investigation. *Clinical Microbiology Reviews*, *15*(3), 506-526. https://doi.org/10.1128/cmr.15.3.506-526.2002
Finkelstein, R., & Oren, I. (2011). Soft Tissue Infections Caused by Marine Bacterial Pathogens: Epidemiology, Diagnosis, and Management. *Current Infectious Disease Reports*, *13*(5), 470-477. https://doi.org/10.1007/s11908-011-0199-3
Fortes, S. T., Lazéra, M. S., Nishikawa, M. M., Regina Célia Lima de, M., & Wanke, B. (2001). First Isolation of <i>Cryptococcus Neoformans</I> Var. <i>gattii</I> From a Native Jungle Tree in the Brazilian Amazon Rainforest. *Mycoses*, *44*(5), 137-140. https://doi.org/10.1046/j.1439-0507.2001.00651.x
Fowkes, F. J. I., Boeuf, P., & Beeson, J. G. (2016). Immunity to Malaria in an Era of Declining Malaria Transmission. *Parasitology*, *143*(2), 139-153. https://doi.org/10.1017/s0031182015001249
Fowler, K. B., Mucha, J., Neumann, M., Lewandowski, W., Kaczanowska, M., Grys, M., Schmidt, E., Natenshon, A., Talarico, C. A., Buck, P. O., & Diaz-Decaro, J. (2022). A Systematic Literature Review of the Global Seroprevalence of Cytomegalovirus: Possible Implications for Treatment, Screening, and Vaccine Development. *BMC Public Health*, *22*(1). https://doi.org/10.1186/s12889-022-13971-7
Frazer, I. H. (2010). Measuring Serum Antibody to Human Papillomavirus Following Infection or Vaccination. *Gynecologic Oncology*, *118*(1), S8-S11. https://doi.org/10.1016/j.ygyno.2010.04.003
Freedman, M., Jackson, B. R., McCotter, O., & Benedict, K. (2018). Coccidioidomycosis Outbreaks, United States and Worldwide, 1940–2015. *Emerging Infectious Diseases*, *24*(3), 417-423. https://doi.org/10.3201/eid2403.170623
Freitas, E. O. d., Leoratti, F. M. S., Freire-de-Lima, C. G., Morrot, A., & Feijó, D. F. (2016). The Contribution of Immune Evasive Mechanisms to Parasite Persistence in Visceral Leishmaniasis. *Frontiers in Immunology*, *7*. https://doi.org/10.3389/fimmu.2016.00153
Frydman, G. H., Davis, N., Beck, P. L., & Fox, J. G. (2015). <i>Helicobacter Pylori</I> Eradication in Patients With Immune Thrombocytopenic Purpura: A Review and the Role of Biogeography. *Helicobacter*, *20*(4), 239-251. https://doi.org/10.1111/hel.12200
Fujimura, S., Kawamura, T., Kato, S., Tateno, H., & Watanabe, A. (2002). Detection of Helicobacter Pylori in Cow's Milk. *Letters in Applied Microbiology*, *35*(6), 504-507. https://doi.org/10.1046/j.1472-765x.2002.01229.x
Fung, C., Tan, S., Nakajima, M., Skoog, E. C., Camarillo-Guerrero, L. F., Klein, J. A., Lawley, T. D., Solnick, J. V., Fukami, T., & Amieva, M. R. (2019). High-Resolution Mapping Reveals That Microniches in the Gastric Glands Control Helicobacter Pylori Colonization of the Stomach. *Plos Biology*, *17*(5), e3000231. https://doi.org/10.1371/journal.pbio.3000231
Furtado, R., Paul, M., Zhang, J., Sung, J., Karell, P., Kim, R. S., Caillat-Zucman, S., Liang, L., Felgner, P. L., Bauleni, A., Gama, S., Buchwald, A. G., Taylor, T. E., Seydel, K. B., Laufer, M. K., Delahaye, F., Daily, J. P., & Lauvau, G. (2023). Cytolytic Circumsporozoite-Specific Memory CD4+ T Cell Clones Are Expanded During Plasmodium Falciparum Infection. *Nature Communications*, *14*(1). https://doi.org/10.1038/s41467-023-43376-y

G, S. N. B. (2015). Epidemiological Studies and Molecular Characterization of Herpes Simplex Virus Among Urban Population in Chennai, Tamilnadu. *Epidemiology Open Access*, *05*(02). https://doi.org/10.4172/2161-1165.1000187
Gagneux, S. (2012). Host-pathogen coevolution in human tuberculosis. *Philos Trans R Soc Lond B Biol Sci*, *367*(1590), 850-859. https://doi.org/10.1098/rstb.2011.0316
Galgiani, J. N., Ampel, N. M., Catanzaro, A., Johnson, R. H., Stevens, D. A., & Williams, P. (2000). Practice Guidelines for the Treatment of Coccidioidomycosis. *Clinical Infectious Diseases*, *30*(4), 658-661. https://doi.org/10.1086/313747
Gao, G., & Xu, J. (2013). Important Biology Events and Pathways in Brucella Infection and Implications for Novel Antibiotic Drug Targets. *Critical Reviews in Eukaryotic Gene Expression*, *23*(1), 65-76. https://doi.org/10.1615/critreveukargeneexpr.2013006580
Gao, Y., Wu, J., Zhang, M., Hou, M., & Ji, M. (2015). Mice Lack of LRG-47 Display the Attenuated Outcome of Infection With Schistosoma Japonicum. *Parasitology Research*, *115*(3), 1185-1193. https://doi.org/10.1007/s00436-015-4853-9
Garnica, M., Cunha, M. O., Portugal, R., Maiolino, Â., Colombo, A. L., & Nucci, M. (2014). Risk Factors for Invasive Fusariosis in Patients With Acute Myeloid Leukemia and in Hematopoietic Cell Transplant Recipients. *Clinical Infectious Diseases*, *60*(6), 875-880. https://doi.org/10.1093/cid/ciu947
George, R. B., Cartier, Y., Casson, A. G., & Hernandez, P. (2006). Suppurative Mediastinitis Secondary to <i>Burkholderia Cepacia</I> in a Patient With Cystic Fibrosis. *Canadian Respiratory Journal*, *13*(4), 215-218. https://doi.org/10.1155/2006/495720
Gerada, C., Campbell, T. M., Kennedy, J. J., McSharry, B. P., Steain, M., Slobedman, B., & Abendroth, A. (2020). Manipulation of the Innate Immune Response by Varicella Zoster Virus. *Frontiers in Immunology*, *11*. https://doi.org/10.3389/fimmu.2020.00001
Gershon, A. A., Breuer, J., Cohen, J. I., Cohrs, R. J., Gershon, M. D., Gilden, D., Grose, C., Hambleton, S., Kennedy, P. G. E., Oxman, M. N., Seward, J. F., & Yamanishi, K. (2015). Varicella Zoster Virus Infection. *Nature Reviews Disease Primers*, *1*(1). https://doi.org/10.1038/nrdp.2015.16
Giacani, L., Brandt, S., Puray-Chavez, M., Reid, T. B., Godornes, C., Molini, B. J., Benzler, M., Hartig, J. S., Lukehart, S. A., & Centurion-Lara, A. (2012). Comparative Investigation of the Genomic Regions Involved in Antigenic Variation of the TprK Antigen Among Treponemal Species, Subspecies, and Strains. *Journal of Bacteriology*, *194*(16), 4208-4225. https://doi.org/10.1128/jb.00863-12
Gilden, D. H., Cohrs, R. J., & Mahalingam, R. (2003). Clinical and Molecular Pathogenesis of Varicella Virus Infection. *Viral Immunology*, *16*(3), 243-258. https://doi.org/10.1089/088282403322396073
Gilgado, F., Cano, J., Gené, J., & Guarro, J. (2005). Molecular Phylogeny of The <i>Pseudallescheria Boydii</I> Species Complex: Proposal of Two New Species. *Journal of Clinical Microbiology*, *43*(10), 4930-4942. https://doi.org/10.1128/jcm.43.10.4930-4942.2005
Giménez, E., Monzó, C., Albert, E., Fuentes-Trillo, A., Seda, E., Piñana, J. L., Hernández-Boluda, J. C., Solano, C., Chaves, F. J., & Navarro, D. (2020). Diversity and Dynamic Changes of Anelloviruses in Plasma Following Allogeneic Hematopoietic Stem Cell Transplantation. *J Med Virol*, *93*(8), 5167-5172. https://doi.org/10.1002/jmv.26661

Glenn, T. J., Blair, J. E., & Adams, R. H. (2005). Coccidioidomycosis in Hematopoietic Stem Cell Transplant Recipients. *Medical Mycology*, *43*(8), 705-710. https://doi.org/10.1080/13693780500147840

Goepfert, C., Stokar-Regenscheit, N., Schumacher, V., Roos, S., Rossier, C., Baehler, C., Schmitt, S., & Posthaus, H. (2014). <i>Mycobacterium Avium</I>Subsp.<i>avium</I>Infection in Four Veal Calves: Differentiation From Intestinal Tuberculosis. *Biomed Research International*, *2014*, 1-5. https://doi.org/10.1155/2014/715841

Goetting-Minesky, M. P., Godovikova, V., Zheng, W., & Fenno, J. C. (2022). Characterization of Treponema Denticola Major Surface Protein (Msp) by Deletion Analysis and Advanced Molecular Modeling. *Journal of Bacteriology*, *204*(9). https://doi.org/10.1128/jb.00228-22

Goh, J. G., Ravikumar, S., Win, M. S., Cao, Q., Tan, A. L., Lim, J. H. J., Leong, W., Herbrecht, R., Troke, P. F., Kullberg, B. J., Netea, M. G., Chng, W. J., Dan, Y. Y., & Chai, L. Y. A. (2017). Neutrophils Differentially Attenuate Immune Response To<i>Aspergillus</I>infection Through Complement Receptor 3 and Induction of Myeloperoxidase. *Cellular Microbiology*, *20*(3), e12798. https://doi.org/10.1111/cmi.12798

González, J., González, M., Sancho, F. V., Sánchez, M. V. L., Martín-Hernández, R., Tercero-Jaime, J. M., & Olmeda, A. S. (2019). Prevalence of Coxiella Burnetii (Legionellales: Coxiellaceae) Infection Among Wildlife Species and the Tick Hyalomma Lusitanicum (Acari: Ixodidae) in a Meso-Mediterranean Ecosystem. *Journal of Medical Entomology*. https://doi.org/10.1093/jme/tjz169

González-Barrio, D., Carpio, A. J., Sebastián-Pardo, M., Peralbo-Moreno, A., & Ruiz-Fons, F. (2022). The Relevance of the Wild Reservoir in Zoonotic Multi-host Pathogens: The Links Between Iberian Wild Mammals And<i>Coxiella Burnetii</I>. *Transboundary and Emerging Diseases*, *69*(6), 3868-3880. https://doi.org/10.1111/tbed.14758

Goodlet, K. J., Nailor, M. D., Omar, A., Huang, J., LiPuma, J. J., Walia, R., & Tokman, S. (2019). Successful Lung Re-Transplant in a Patient With Cepacia Syndrome Due to Burkholderia Ambifaria. *Journal of Cystic Fibrosis*, *18*(1), e1-e4. https://doi.org/10.1016/j.jcf.2018.08.011

Goyal, S., Castrillón-Betancur, J. C., Klaile, E., & Slevogt, H. (2018). The Interaction of Human Pathogenic Fungi With C-Type Lectin Receptors. *Frontiers in Immunology*, *9*. https://doi.org/10.3389/fimmu.2018.01261

Gravelat, F. N., Beauvais, A., Liu, H., Lee, M. J., Snarr, B. D., Chen, D., Xu, W., Kravtsov, I., Hoareau, C. M., Vanier, G., Urb, M., Campoli, P., Abdallah, Q. A., Lehoux, M., Chabot, J. C., Ouimet, M.-C., Baptista, S. D., Fritz, H., Nierman, W. C.,…Sheppard, D. C. (2013). Aspergillus Galactosaminogalactan Mediates Adherence to Host Constituents and Conceals Hyphal Β-Glucan From the Immune System. *PLOS Pathogens*, *9*(8), e1003575. https://doi.org/10.1371/journal.ppat.1003575

Gray, J. I., Westerhof, L. M., & MacLeod, M. K. L. (2018). The Roles of Resident, Central and Effector Memory <scp>CD</Scp>4 T-cells in Protective Immunity Following Infection or Vaccination. *Immunology*, *154*(4), 574-581. https://doi.org/10.1111/imm.12929

Griffiths, P., Baraniak, I., & Reeves, M. B. (2014). The Pathogenesis of Human Cytomegalovirus. *The Journal of Pathology*, *235*(2), 288-297. https://doi.org/10.1002/path.4437

Grønseth, S., Rogne, T., Hannula, R., Åsvold, B. O., Afset, J. E., & Damås, J. K. (2021). Semiquantitative Real-Time PCR to Distinguish Pneumocystis Pneumonia From Colonization in a

Heterogeneous Population of HIV-Negative Immunocompromised Patients. *Microbiology Spectrum*, *9*(1). https://doi.org/10.1128/spectrum.00026-21
Guatteo, R. R., Beaudeau, F., Joly, A., & Seegers, H. H. (2007). <i>Coxiella Burnetii</I>shedding by Dairy Cows. *Veterinary Research*, *38*(6), 849-860. https://doi.org/10.1051/vetres:2007038
Guérin, E., Poirier, P., Nervo, M., & Terrier, C. L. (2021). Fatal Multiorgan Failure Syndrome in a Strongyloides-HTLV-1 Coinfected Patient, After Treatment With Ivermectin. *Case Reports in Critical Care*, *2021*, 1-5. https://doi.org/10.1155/2021/5554810
Guerra, M. (2013). Leptospirosis: Public Health Perspectives. *Biologicals*, *41*(5), 295-297. https://doi.org/10.1016/j.biologicals.2013.06.010
Guillot, L., Carroll, S. F., Badawy, M., & Qureshi, S. T. (2008). Cryptococcus Neoformans Induces IL-8 Secretion and CXCL1 Expression by Human Bronchial Epithelial Cells. *Respiratory Research*, *9*(1). https://doi.org/10.1186/1465-9921-9-9
Guinea, J. (2014). Global Trends in the Distribution of Candida Species Causing Candidemia. *Clinical Microbiology and Infection*, *20*, 5-10. https://doi.org/10.1111/1469-0691.12539
Gulve, N., Frank, C., Klepsch, M., & Prusty, B. K. (2017). Chromosomal Integration of HHV-6A During Non-Productive Viral Infection. *Scientific Reports*, *7*(1). https://doi.org/10.1038/s41598-017-00658-y
Guptill, L. (2010). Feline Bartonellosis. *Veterinary Clinics of North America Small Animal Practice*, *40*(6), 1073-1090. https://doi.org/10.1016/j.cvsm.2010.07.009
Gurgel-Gonçalves, R., Galvão, C., Costa, J., & Peterson, A. T. (2012). Geographic Distribution of Chagas Disease Vectors in Brazil Based on Ecological Niche Modeling. *Journal of Tropical Medicine*, *2012*, 1-15. https://doi.org/10.1155/2012/705326
Ha, R., Keynan, Y., & Rueda, Z. V. (2022). Increased Susceptibility to Pneumonia Due to Tumour Necrosis Factor Inhibition and Prospective Immune System Rescue via Immunotherapy. *Frontiers in Cellular and Infection Microbiology*, *12*. https://doi.org/10.3389/fcimb.2022.980868
Haag, A. F., Myka, K. K., Arnold, M. F. F., Caro-Hernández, P., & Ferguson, G. P. (2010). Importance of Lipopolysaccharide and Cyclic<i>β</I>-1,2-Glucans In<i>Brucella</I>-Mammalian Infections. *International Journal of Microbiology*, *2010*, 1-12. https://doi.org/10.1155/2010/124509
Hadfield, N. J., Selvendran, S., & Johnston, M. (2019). Fatal Pneumocystis Jirovecii Pneumonia in a Non-Immunocompromised Patient With Alcohol-Related Liver Cirrhosis. *Scottish Medical Journal*, *64*(4), 148-153. https://doi.org/10.1177/0036933019872629
Haeberlein, S., Obieglo, K., Ozir-Fazalalikhan, A., Chayé, M. A. M., Veninga, H., Luciën, E. P. M. v. d. V., Voskamp, A., Boon, L., Joke, M. M. d. H., Westerhof, L. B., Wilbers, R. H. P., Schots, A., Schramm, G., Hokke, C. H., & Smits, H. H. (2017). Schistosome Egg Antigens, Including the Glycoprotein IPSE/alpha-1, Trigger the Development of Regulatory B Cells. *PLOS Pathogens*, *13*(7), e1006539. https://doi.org/10.1371/journal.ppat.1006539
Hajjeh, R. (1995). Disseminated Histoplasmosis in Persons Infected With Human Immunodeficiency Virus. *Clinical Infectious Diseases*, *21*(Supplement_1), S108-S110. https://doi.org/10.1093/clinids/21.supplement_1.s108
Hamasha, A. M. S., Yıldıran, Ş. T., Gönlüm, A., Saraçlı, M. A., & Doğancı, L. (2004). Cryptococcus Neoformans Varieties From Material Under the Canopies of Eucalyptus Trees and Pigeon Dropping Samples From Four Major Cities in Jordan. *Mycopathologia*, *158*(2), 195-199. https://doi.org/10.1023/b:myco.0000041840.34011.23

Hanson, D., Hill, J. A., & Koelle, D. M. (2018). Advances in the Characterization of the T-Cell Response to Human Herpesvirus-6. *Frontiers in Immunology*, *9*. https://doi.org/10.3389/fimmu.2018.01454

Haque, A. R., Sultana, A., Habib, A., & Zakaria, A. S. M. (2014). Prevention &Amp; Management of Herpes Zoster- An Update. *Bangladesh Medical Journal*, *41*(3), 53-56. https://doi.org/10.3329/bmj.v41i3.18961

Harker, K. S., Ueno, N., & Lodoen, M. B. (2015). <i>Toxoplasma Gondii</I> Dissemination: A Parasite's Journey Through the Infected Host. *Parasite Immunology*, *37*(3), 141-149. https://doi.org/10.1111/pim.12163

Harms, A., & Dehio, C. (2012). Intruders Below the Radar: Molecular Pathogenesis of Bartonella SPP. *Clinical Microbiology Reviews*, *25*(1), 42-78. https://doi.org/10.1128/cmr.05009-11

Harms, P. W., Collie, A. M. B., Hovelson, D. H., Cani, A. K., Verhaegen, M., Patel, R. M., Fullen, D. R., Omata, K., Dlugosz, A. A., Tomlins, S. A., & Billings, S. D. (2016). Next Generation Sequencing of Cytokeratin 20-Negative Merkel Cell Carcinoma Reveals Ultraviolet-Signature Mutations and Recurrent TP53 and RB1 Inactivation. *Modern Pathology*, *29*(3), 240-248. https://doi.org/10.1038/modpathol.2015.154

Hauser, N., & Orsini, J. (2015). Cepacia Syndrome in a Non-Cystic Fibrosis Patient. *Case Reports in Infectious Diseases*, *2015*, 1-4. https://doi.org/10.1155/2015/537627

Hays, R., Esterman, A., & McDermott, R. (2015). Type 2 Diabetes Mellitus Is Associated With Strongyloides Stercoralis Treatment Failure in Australian Aboriginals. *Plos Neglected Tropical Diseases*, *9*(8), e0003976. https://doi.org/10.1371/journal.pntd.0003976

Heath, C. H., Slavin, M. A., Sorrell, T. C., Handke, R., Harun, A., Phillips, M., Nguyen, Q., Delhaès, L., Ellis, D., Meyer, W., & Chen, S. C. A. (2009). Population-Based Surveillance for Scedosporiosis in Australia: Epidemiology, Disease Manifestations and Emergence of Scedosporium Aurantiacum Infection. *Clinical Microbiology and Infection*, *15*(7), 689-693. https://doi.org/10.1111/j.1469-0691.2009.02802.x

Hedderwick, S., Lyons, M. J., Liu, M., Vázquez, J. A., & Kauffman, C. A. (2000). Epidemiology of Yeast Colonization in the Intensive Care Unit. *European Journal of Clinical Microbiology & Infectious Diseases*, *19*(9), 663-670. https://doi.org/10.1007/s100960000348

Heegaard, E. D., & Brown, K. (2002). Human Parvovirus B19. *Clinical Microbiology Reviews*, *15*(3), 485-505. https://doi.org/10.1128/cmr.15.3.485-505.2002

Hershberg, R., Lipatov, M., Small, P. M., Sheffer, H., Niemann, S., Homolka, S., Roach, J. C., Kremer, K., Petrov, D. A., Feldman, M. W., & Gagneux, S. (2008). High functional diversity in Mycobacterium tuberculosis driven by genetic drift and human demography. *PLoS Biol*, *6*(12), e311. https://doi.org/10.1371/journal.pbio.0060311

Hidaka, E. (2001). Helicobacter Pylori and Two Ultrastructurally Distinct Layers of Gastric Mucous Cell Mucins in the Surface Mucous Gel Layer. *Gut*, *49*(4), 474-480. https://doi.org/10.1136/gut.49.4.474

Hildreth, J. E. K., & Alcendor, D. J. (2021). JC Polyomavirus and Transplantation: Implications for Virus Reactivation After Immunosuppression in Transplant Patients and the Occurrence of PML Disease. *Transplantology*, *2*(1), 37-48. https://doi.org/10.3390/transplantology2010004

Hill, D. E., Sreekumar, C., & Dubey, J. P. (2005). Biology and Epidemiology Of<i>Toxoplasma Gondii</I>in Man and Animals. *Animal Health Research Reviews*, *6*(1), 41-61. https://doi.org/10.1079/ahr2005100

Hino, M., Ishiko, O., Honda, K., Yamane, T., Ohta, K., Takubo, T., & Tatsumi, N. (2000). Transmission of Symptomatic Parvovirus B19 Infection by Fibrin Sealant Used During Surgery. *British Journal of Haematology*, *108*(1), 194-195. https://doi.org/10.1046/j.1365-2141.2000.01818.x

Hirsch, H. H. (2002). Polyomavirus BK Nephropathy: A (Re-)emerging Complication in Renal Transplantation. *American Journal of Transplantation*, *2*(1), 25-30. https://doi.org/10.1034/j.1600-6143.2002.020106.x

Hirsch, H. H., & Snydman, D. R. (2005). BK Virus: Opportunity Makes a Pathogen. *Clinical Infectious Diseases*, *41*(3), 354-360. https://doi.org/10.1086/431488

Ho, S. S. C., Nashid, N., Waters, V., LiPuma, J. J., Zlosnik, J. E. A., Otley, A., Somers, G. R., Kamath, B. M., & Yau, Y. (2019). Burkholderia Multivorans Septicemia in a Pediatric Liver Transplant Patient. *American Journal of Transplantation*, *19*(3), 933-938. https://doi.org/10.1111/ajt.15065

Hogerwerf, L., Borlée, F., Still, K., Heederik, D., Rotterdam, B. J. v., Bruin, A. d., Nielen, M., & Wouters, I. M. (2012). Detection of Coxiella Burnetii DNA in Inhalable Airborne Dust Samples From Goat Farms After Mandatory Culling. *Applied and Environmental Microbiology*, *78*(15), 5410-5412. https://doi.org/10.1128/aem.00677-12

Hoog, S. d., Marvin-Sikkema, F. D., Lahpoor, G. A., Gottschall, J. C., Prins, R. A., & Guého, E. (1994). Ecology and Physiology of the Emerging Opportunistic Fungi <i>Pseudallescheria Boydii</I> and <i>Scedosporium Prolificans</I>. *Mycoses*, *37*(3-4), 71-78. https://doi.org/10.1111/j.1439-0507.1994.tb00780.x

Hope, R., Aldred, D., & Magan, N. (2005). Comparison of Environmental Profiles for Growth and Deoxynivalenol Production by Fusarium Culmorum and F. Graminearum on Wheat Grain. *Letters in Applied Microbiology*, *40*(4), 295-300. https://doi.org/10.1111/j.1472-765x.2005.01674.x

Horné, D., & Skerrett, S. (2019). Recent Advances in Nontuberculous Mycobacterial Lung Infections. *F1000research*, *8*, 1710. https://doi.org/10.12688/f1000research.20096.1

Houston, S., Schovanek, E., Conway, K. M. E., Mustafa, S., Gomez, A., Ramaswamy, R., Haimour, A., Boulanger, M. J., Reynolds, L. A., & Cameron, C. E. (2022). Identification and Functional Characterization of Peptides With Antimicrobial Activity From the Syphilis Spirochete, Treponema Pallidum. *Frontiers in Microbiology*, *13*. https://doi.org/10.3389/fmicb.2022.888525

Howard, Z. P., & Omsland, A. (2020). Selective Inhibition of Coxiella Burnetii Replication by the Steroid Hormone Progesterone. *Infect Immun*, *88*(12). https://doi.org/10.1128/iai.00894-19

Hsiao, A., Liu, Z., Joelsson, A., & Zhu, J. (2006). <i>Vibrio Cholerae</I>virulence Regulator-Coordinated Evasion of Host Immunity. *Proceedings of the National Academy of Sciences*, *103*(39), 14542-14547. https://doi.org/10.1073/pnas.0604650103

Hughes, W. O. H., & Boomsma, J. J. (2004). Let Your Enemy Do the Work: Within–host Interactions Between Two Fungal Parasites of Leaf–cutting Ants. *Proceedings of the Royal Society B Biological Sciences*, *271*(suppl_3). https://doi.org/10.1098/rsbl.2003.0115

Hunter, C. A., & Sibley, L. D. (2012). Modulation of Innate Immunity by Toxoplasma Gondii Virulence Effectors. *Nature Reviews Microbiology*, *10*(11), 766-778. https://doi.org/10.1038/nrmicro2858

Ikuta, Y., Oba, K., Nai, E., Katori, T., Hashino, M., Inamine, Y., Matsunaga, S., Yamaoka, Y., Sekizuka, T., Ryo, A., & Kuroda, M. (2019). Aseptic Meningitis Caused by Torque Teno Virus in an

Infant: A Case Report. *Journal of Medical Case Reports*, *13*(1). https://doi.org/10.1186/s13256-019-2233-2
Isabella Maria Monteiro de, S., Novaes, R. D., Gonçalves, R. V., Fialho, F. L. B., Carvalho, D. T., Souza, T. B. d., Dias, D. F., Lavorato, S. N., Souza, R. L. M., Marques, M. J., & Castro, A. P. (2022). In Vitro and in Silico Evaluation of the Schistosomicidal Activity of Eugenol Derivatives Using Biochemical, Molecular, and Morphological Tools. *Journal of Venomous Animals and Toxins Including Tropical Diseases*, *28*. https://doi.org/10.1590/1678-9199-jvatitd-2021-0108
Jafri, R., Ali, A., Messonnier, N. E., Tévi-Bénissan, C., Dürrheim, D. N., Eskola, J., Fermon, F., Klugman, K. P., Ramsay, M., Sow, S. O., Shao, Z., Bhutta, Z. A., & Abramson, J. S. (2013). Global Epidemiology of Invasive Meningococcal Disease. *Population Health Metrics*, *11*(1). https://doi.org/10.1186/1478-7954-11-17
James, S. H., & Kimberlin, D. W. (2015). Neonatal Herpes Simplex Virus Infection. *Infectious Disease Clinics of North America*, *29*(3), 391-400. https://doi.org/10.1016/j.idc.2015.05.001
Janković, M., Milićević, O., Todorović, M., Đunić, I., Mihaljević, B., Jovanović, T., & Knežević, A. (2023). Guilty in Absentia – Cytomegalovirus Seropositivity Relates Inversely to Cancer Incidence in Race/Ethnicity Groups and Worldwide: Is Latent Viral Infection Oncopreventive? https://doi.org/10.22541/au.169027024.42199854/v1
Ji, F., Liu, Z., Cao, J., Li, N., Liu, Z., Zuo, J., Yan, C., Wang, X., & Sun, J. (2008). B Cell Response Is Required for Granuloma Formation in the Early Infection of Schistosoma Japonicum. *PLoS One*, *3*(3), e1724. https://doi.org/10.1371/journal.pone.0001724
Ji, J., Sun, H.-K., Smith, J. S., Wang, H., Esser, M. T., Hu, S., Pretorius, R. G., Chen, W., Belinson, J. L., & Qiao, Y. L. (2012). Seroprevalence of Human Papillomavirus Types 6, 11, 16 and 18 in Chinese Women. *BMC Infectious Diseases*, *12*(1). https://doi.org/10.1186/1471-2334-12-137
Jiang, Y., Xu, X., Qing, X., & Pan, W. (2011). Identification and Characterization of Six Novel Tetraspanins From Schistosoma Japonicum. *Parasites & Vectors*, *4*(1). https://doi.org/10.1186/1756-3305-4-190
Jiang, Z.-G., Cohen, J. I., Marshall, L. J., & Major, E. O. (2010). Hexadecyloxypropyl-Cidofovir (CMX001) Suppresses JC Virus Replication in Human Fetal Brain SVG Cell Cultures. *Antimicrobial Agents and Chemotherapy*, *54*(11), 4723-4732. https://doi.org/10.1128/aac.00837-10
Jiao, H., Zhou, Z., Li, B., Xiao, Y., Li, M., Zeng, H., Guo, X., & Gu, G. (2021). The Mechanism of Facultative Intracellular Parasitism of Brucella. *International Journal of Molecular Sciences*, *22*(7), 3673. https://doi.org/10.3390/ijms22073673
João Marcelo Azevedo de Paula, A., & Megid, J. (2013). Brucella Ovis: Invasão, Tráfego, Fatores De Virulência E Resposta Imune. *Semina Ciências Agrárias*, *34*(3). https://doi.org/10.5433/1679-0359.2013v34n3p1301-1312
Johnston, C., Koelle, D. M., & Wald, A. (2011). HSV-2: In Pursuit of a Vaccine. *Journal of Clinical Investigation*, *121*(12), 4600-4609. https://doi.org/10.1172/jci57148
Jong, E. P. d., Haan, T. R. d., Kroes, A. C., Beersma, M. F. C., Oepkes, D., & Walther, F. J. (2006). Parvovirus B19 Infection in Pregnancy. *Journal of Clinical Virology*, *36*(1), 1-7. https://doi.org/10.1016/j.jcv.2006.01.004
Jong, M. F. d., & Tsolis, R. M. (2011). Brucellosis and Type Iv Secretion. *Future Microbiology*, *7*(1), 47-58. https://doi.org/10.2217/fmb.11.136

Jordan, J. A., Huff, D. S., & DeLoia, J. A. (2001). Placental Cellular Immune Response in Women Infected With Human Parvovirus B19 During Pregnancy. *Clinical and Diagnostic Laboratory Immunology*, *8*(2), 288-292. https://doi.org/10.1128/cdli.8.2.288-292.2001

Kaminiów, K., Kiołbasa, M., & Pastuszczak, M. (2024). The Significance of the Cell-Mediated Host Immune Response in Syphilis. *Microorganisms*, *12*(12), 2580. https://doi.org/10.3390/microorganisms12122580

Kant, S., Dasgupta, A., Bagnasco, S. M., & Brennan, D. C. (2022). BK Virus Nephropathy in Kidney Transplantation: A State-of-the-Art Review. *Viruses*, *14*(8), 1616. https://doi.org/10.3390/v14081616

Kantardjiev, T., Levterova, V., Panaiotov, S., Ivanov, I., & Hristozova, E. (2006). Molecular Taxonomy Of<i>Cryptococcus Neoformans</I>Varieties Displaying Phenotypic Similarities. *Biotechnology & Biotechnological Equipment*, *20*(2), 101-103. https://doi.org/10.1080/13102818.2006.10817349

Karatuna, O., Matuschek, E., Åhman, J., Caidi, H., & Kahlmeter, G. (2023). <i>Vibrio</I> Species: Development of EUCAST Susceptibility Testing Methods and MIC and Zone Diameter Distributions on Which to Determine Clinical Breakpoints. *Journal of Antimicrobial Chemotherapy*, *79*(2), 375-382. https://doi.org/10.1093/jac/dkad391

Katzenellenbogen, R. A., Carter, J. J., Stern, J., Kovacic, M. S. B., Mehta, P. A., Sauter, S., Galloway, D. A., & Winer, R. L. (2015). Skin and Mucosal Human Papillomavirus Seroprevalence in Persons With Fanconi Anemia. *Clinical and Vaccine Immunology*, *22*(4), 413-420. https://doi.org/10.1128/cvi.00665-14

Kaye, P. M., Cruz, I., Picado, A., Bocxlaer, K. V., & Croft, S. L. (2020). Leishmaniasis Immunopathology—impact on Design and Use of Vaccines, Diagnostics and Drugs. *Seminars in Immunopathology*, *42*(3), 247-264. https://doi.org/10.1007/s00281-020-00788-y

Khairunnisa, S., Soeroso, N. N., Abdullah, M., Siahaan, L., Eyanoer, P. C., Daulay, E. R., Wahyuningsih, R., & Denning, D. W. (2023). Factors Influencing Histoplasmosis Incidence in Multidrug-Resistant Pulmonary Tuberculosis Patients: A Cross-Sectional Study in Indonesia. *Narra J*, *3*(3), e403. https://doi.org/10.52225/narra.v3i3.403

Khaliq, M. F., Ihle, R., & Perry, J. D. (2018). Immunosuppression With Antitumour Necrosis Factor Therapy Leading to <i>Strongyloides</I> Hyperinfection Syndrome. *Case Reports in Infectious Diseases*, *2018*, 1-4. https://doi.org/10.1155/2018/6341680

Khan, N. A., & Siddiqui, R. (2014). Predator<i>vs</I>aliens: Bacteria Interactions With<i>Acanthamoeba</I>. *Parasitology*, *141*(7), 869-874. https://doi.org/10.1017/s003118201300231x

Khan, Z. U., Guardiola, J., & Salzman, G. (2005). A Case of Mycobacterium Avium Complex Pulmonary Disease in an Immunocompetent Host. *Southern Medical Journal*, *98*(10), 1036-1038. https://doi.org/10.1097/01.smj.0000182516.84423.ea

Kim, E. W., Nadipuram, S. M., Tetlow, A. L., Barshop, W. D., Liu, P. T., Wohlschlegel, J. A., & Bradley, P. J. (2016). The Rhoptry Pseudokinase ROP54 Modulates Toxoplasma Gondii Virulence and Host GBP2 Loading. *Msphere*, *1*(2). https://doi.org/10.1128/msphere.00045-16

Kim, H. J., Ryu, J.-O., Lee, S.-Y., Kim, E.-S., & Kim, H. Y. (2015). Multiplex PCR for Detection of the Vibrio Genus and Five Pathogenic Vibrio Species With Primer Sets Designed Using Comparative Genomics. *BMC Microbiology*, *15*(1). https://doi.org/10.1186/s12866-015-0577-3

Kim, H. S., Kim, Y. E., Yun, E. Y., Ju, J. H., Eun, J., Lee, G. D., Cho, Y. J., Kim, H. C., Lee, J. D., Hwang, Y. S., & Jeong, Y. Y. (2010). A Case of Fatal Hyperinfective Strongyloidiasis With Acute Respiratory Failure and Intestinal Perforation in Lung Cancer Patient. *Tuberculosis and Respiratory Diseases*, *68*(1), 29. https://doi.org/10.4046/trd.2010.68.1.29

Kim, M. A., Oh, J. K., Kim, B. W., Chay, D., Park, D. C., Kim, S. M., Kang, E. S., Kim, J. H., Cho, C. H., Shin, H.-R., & Seo, K. (2012). Prevalence and Seroprevalence of Low-Risk Human Papillomavirus in Korean Women. *Journal of Korean Medical Science*, *27*(8), 922. https://doi.org/10.3346/jkms.2012.27.8.922

Kim, S. H., Ha, Y. E., Youn, J. C., Park, J. S., Sung, H., Kim, M. N., Choi, H., Lee, Y. J., Kang, S. M., Ahn, J. Y., Choi, J. Y., Kim, Y. J., Lee, S. K., Kim, S. J., Peck, K. R., Lee, S. O., Kim, Y. H., Hwang, S., Lee, S. G.,…Han, D. J. (2015). Fatal Scedosporiosis in Multiple Solid Organ Allografts Transmitted From a Nearly-Drowned Donor. *American Journal of Transplantation*, *15*(3), 833-840. https://doi.org/10.1111/ajt.13008

King, A. S., Castro, J. G., & Dow, G. (2008). <i>Nocardia Farcinica</I> Lung Abscess Presenting in the Context of Advanced HIV Infection: Spontaneous Resolution in Response to Highly Active Antiretroviral Therapy Alone. *Canadian Journal of Infectious Diseases and Medical Microbiology*, *20*(3). https://doi.org/10.1155/2009/181750

King, C. C., Jamieson, D. J., Wiener, J., Cu-Uvin, S., Klein, R. S., Rompalo, A., Shah, K. V., & Sobel, J. D. (2011). Bacterial Vaginosis and the Natural History of Human Papillomavirus. *Infectious Diseases in Obstetrics and Gynecology*, *2011*, 1-8. https://doi.org/10.1155/2011/319460

Kireev, F. D., Lopatnikova, J., Alshevskaya, A., & Сенников, С. В. (2025). Role of Tumor Necrosis Factor in Tuberculosis. *Biomolecules*, *15*(5), 709. https://doi.org/10.3390/biom15050709

Kirkham, H., Carnes, S., Lieberman, J. A., Cohen, S., & Reddi, D. (2021). Occult Strongyloides Stercoralis Infection Masquerading as a Drug Reaction With Eosinophilic Pneumonia. *Journal of Clinical Images and Medical Case Reports*, *2*(3). https://doi.org/10.52768/2766-7820/1166

Klenerman, P., & Oxenius, A. (2016). T Cell Responses to Cytomegalovirus. *Nature Reviews Immunology*, *16*(6), 367-377. https://doi.org/10.1038/nri.2016.38

Knöll, A., Louwen, F., Kochanowski, B., Plentz, A., Stüssel, J., Beckenlehner, K., Jilg, W., & Modrow, S. (2002). Parvovirus B19 Infection in Pregnancy: Quantitative Viral DNA Analysis Using a Kinetic Fluorescence Detection System (TaqMan PCR). *J Med Virol*, *67*(2), 259-266. https://doi.org/10.1002/jmv.2216

Knox, B. P., Deng, Q., Rood, M., Eickhoff, J. C., Keller, N. P., & Huttenlocher, A. (2014). Distinct Innate Immune Phagocyte Responses to Aspergillus Fumigatus Conidia and Hyphae in Zebrafish Larvae. *Eukaryotic Cell*, *13*(10), 1266-1277. https://doi.org/10.1128/ec.00080-14

Ko, J., & Splitter, G. A. (2003). Molecular Host-Pathogen Interaction in Brucellosis: Current Understanding and Future Approaches to Vaccine Development for Mice and Humans. *Clinical Microbiology Reviews*, *16*(1), 65-78. https://doi.org/10.1128/cmr.16.1.65-78.2003

Koh, T. H., Tan, A. L., Lo, Y. S., & Oh, H. (2002). <i>Cryptococcus Neoformans</I>var.<i>gattii</I>meningitis in Singapore. *Medical Mycology*, *40*(2), 221-223. https://doi.org/10.1080/mmy.40.2.221.223

Kojima, N., Konda, K. A., & Klausner, J. D. (2022). Notes on Syphilis Vaccine Development. *Frontiers in Immunology*, *13*. https://doi.org/10.3389/fimmu.2022.952284

Kokseng, S. L., & Blair, J. E. (2010). Successful Kidney Transplantation After Coccidioidal Meningitis. *Transplant Infectious Disease*, *13*(3), 285-289. https://doi.org/10.1111/j.1399-3062.2010.00576.x

Kolasa, M., Hagman, S., Verkkoniemi-Ahola, A., Airas, L., Koivisto, K., & Elovaara, I. (2015). Anti-Jc Virus Seroprevalence in a Finnish MS Cohort. *Acta Neurologica Scandinavica*, *133*(5), 391-397. https://doi.org/10.1111/ane.12475

Kolwijck, E., & Frank, L. v. d. V. (2014). The Potential Impact of the Pulmonary Microbiome on Immunopathogenesis of <i>Aspergillus</I>-related Lung Disease. *European Journal of Immunology*, *44*(11), 3156-3165. https://doi.org/10.1002/eji.201344404

Koohi, A. K., Ravanshad, M., Manouchehr, R., Falahi, S., & Baghban, A. (2012). Phylogenetic Analysis of Torque Teno Virus in Hepatitis C Virus Infected Patients in Shiraz. *Hepatitis Monthly*, *12*(7), 437-441. https://doi.org/10.5812/hepatmon.6133

Kosmidis, C., & Denning, D. W. (2014). The Clinical Spectrum of Pulmonary Aspergillosis. *Thorax*, *70*(3), 270-277. https://doi.org/10.1136/thoraxjnl-2014-206291

Kottom, T. J., Carmona, E. M., & Limper, A. H. (2021). Current State of Carbohydrate Recognition and C-Type Lectin Receptors in Pneumocystis Innate Immunity. *Frontiers in Immunology*, *12*. https://doi.org/10.3389/fimmu.2021.798214

Kotton, C. N., & Lattes, R. (2009). Parasitic Infections in Solid Organ Transplant Recipients. *American Journal of Transplantation*, *9*, S234-S251. https://doi.org/10.1111/j.1600-6143.2009.02915.x

Koujah, L., Suryawanshi, R. K., & Shukla, D. (2018). Pathological Processes Activated by Herpes Simplex Virus-1 (HSV-1) Infection in the Cornea. *Cellular and Molecular Life Sciences*, *76*(3), 405-419. https://doi.org/10.1007/s00018-018-2938-1

Kricker, A., Weber, M., Brenner, N., Banks, E., Pawlita, M., Sitas, F., Hodgkinson, V. S., Rahman, B., Kemenade, C. H. v., Armstrong, B. K., & Waterboer, T. (2020). High Ambient Solar UV Correlates With Greater Beta HPV Seropositivity in New South Wales, Australia. *Cancer Epidemiology Biomarkers & Prevention*, *29*(1), 49-56. https://doi.org/10.1158/1055-9965.epi-19-0400

Kristinsdóttir, Í., Visser, L. J., Miellet, W. R., Mariman, R., Pluister, G., Haraldsson, G., Haraldsson, Á., Trzciński, K., & Thors, V. (2023). Meningococcal Carriage in Children and Young Adults: A Cross-Sectional and Longitudinal Study, Iceland, 2019 to 2021. *Eurosurveillance*, *28*(39). https://doi.org/10.2807/1560-7917.es.2023.28.39.2300215

Krzych, U., Zarling, S., & Pichugin, A. (2014). Memory T Cells Maintain Protracted Protection Against Malaria. *Immunology Letters*, *161*(2), 189-195. https://doi.org/10.1016/j.imlet.2014.03.011

Kumar, R., Mickael, C., Kassa, B., Sanders, L., Koyanagi, D., Hernández-Saavedra, D., Freeman, S., Morales-Cano, D., Cogolludo, Á., McKee, A. S., Fontenot, A. P., Butrous, G., Tuder, R. M., & Graham, B. B. (2019). Th2 Cd4 ≪sup>+</Sup> T Cells Are Necessary and Sufficient for ≪i>Schistosoma</I>-Pulmonary Hypertension. *SSRN Electronic Journal*. https://doi.org/10.2139/ssrn.3378685

Kumar, S., Basavraj, G. K., & Papireddy, C. K. R. (2020). Strongyloides Stercoralis Hyper Infection Syndrome. *Indian Journal of Surgery*, *83*(S3), 582-586. https://doi.org/10.1007/s12262-020-02292-x

Kümpers, P., Tiede, A., Kirschner, P. A., Girke, J., Ganser, A., & Peest, D. (2008). Legionnaires' Disease in Immunocompromised Patients: A Case Report of Legionella Longbeachae Pneumonia and Review of the Literature. *Journal of Medical Microbiology*, *57*(3), 384-387. https://doi.org/10.1099/jmm.0.47556-0

Kurniawati, R. D., & Nuryati, S. (2018). The Correlation Between Physical Environmental Factors and the Occurrence of Leptospirosis. *Jurnal Kesehatan Masyarakat*, *14*(2), 223-230. https://doi.org/10.15294/kemas.v14i2.13527

Kuwamoto, S., Higaki, H., Kanai, K., Iwasaki, T., Sano, H., Nagata, K., Kato, K., Kato, M., Murakami, I., Horie, Y., Yamamoto, O., & Hayashi, K. (2011). Association of Merkel Cell Polyomavirus Infection With Morphologic Differences in Merkel Cell Carcinoma. *Human Pathology*, *42*(5), 632-640. https://doi.org/10.1016/j.humpath.2010.09.011

Kwon-Chung, K. J., & Varma, A. K. (2006). Do Major Species Concepts Support One, Two or More Species Within<i>Cryptococcus Neoformans</I>? *Fems Yeast Research*, *6*(4), 574-587. https://doi.org/10.1111/j.1567-1364.2006.00088.x

Lackner, M., & Hoog, S. d. (2011). Parascedosporium and Its Relatives: Phylogeny and Ecological Trends. *Ima Fungus*, *2*(1), 39-48. https://doi.org/10.5598/imafungus.2011.02.01.07

Lagos, D., Vart, R. J., Gratrix, F., Westrop, S. J., Emuss, V., Wong, P. P., Robey, R. C., Imami, N., Bower, M., Gotch, F., & Boshoff, C. (2008). Toll-Like Receptor 4 Mediates Innate Immunity to Kaposi Sarcoma Herpesvirus. *Cell Host & Microbe*, *4*(5), 470-483. https://doi.org/10.1016/j.chom.2008.09.012

Lamarche, C., Orio, J., Collette, S., Senécal, L., Hébert, M. J., Renoult, E., Tibbles, L. A., & Delisle, J. S. (2016). BK Polyomavirus and the Transplanted Kidney. *Transplantation*, *100*(11), 2276-2287. https://doi.org/10.1097/tp.0000000000001333

Lamb, E., Crow, E. T., Lim, K. C., Liang, Y. s., Lewis, F. A., & Davies, S. J. (2007). Conservation of CD4+ T Cell-Dependent Developmental Mechanisms in the Blood Fluke Pathogens of Humans. *International Journal for Parasitology*, *37*(3-4), 405-415. https://doi.org/10.1016/j.ijpara.2006.11.001

Langley, R., Kenna, D., Vandamme, P., Ure, R., & Govan, J. R. W. (2003). Lysogeny and Bacteriophage Host Range Within the Burkholderia Cepacia Complex. *Journal of Medical Microbiology*, *52*(6), 483-490. https://doi.org/10.1099/jmm.0.05099-0

Laniado-Laborín, R. (2007). Expanding Understanding of Epidemiology of Coccidioidomycosis in the Western Hemisphere. *Annals of the New York Academy of Sciences*, *1111*(1), 19-34. https://doi.org/10.1196/annals.1406.004

Larsen, C. N., & Larsen, H. K. (2020). Herpetiform Manifestation of Primary Syphilis: A Case Series. *Acta Dermato Venereologica*, *100*(6), adv00072-00072. https://doi.org/10.2340/00015555-3414

Laskin, B. L., Denburg, M., Furth, S. L., Diorio, D., Goebel, J., Davies, S. M., & Jodele, S. (2013). BK Viremia Precedes Hemorrhagic Cystitis in Children Undergoing Allogeneic Hematopoietic Stem Cell Transplantation. *Biology of Blood and Marrow Transplantation*, *19*(8), 1175-1182. https://doi.org/10.1016/j.bbmt.2013.05.002

Lass-Flörl, C., Roilides, E., Löffler, J., Wilflingseder, D., & Romani, L. (2013). Minireview: Host Defence in Invasive Aspergillosis. *Mycoses*, *56*(4), 403-413. https://doi.org/10.1111/myc.12052

Latgé, J. P., & Chamilos, G. (2019). Aspergillus Fumigatus and Aspergillosis in 2019. *Clinical Microbiology Reviews*, *33*(1). https://doi.org/10.1128/cmr.00140-18

Lau, C. L., Smythe, L. D., Craig, S. B., & Weinstein, P. (2010). Climate Change, Flooding, Urbanisation and Leptospirosis: Fuelling the Fire? *Transactions of the Royal Society of Tropical Medicine and Hygiene*, *104*(10), 631-638. https://doi.org/10.1016/j.trstmh.2010.07.002

Lau, C. L., Watson, C., Lowry, J. H., David, M., Craig, S. B., Wynwood, S. J., Kama, M., & Nilles, E. J. (2016). Human Leptospirosis Infection in Fiji: An Eco-Epidemiological Approach to Identifying Risk Factors and Environmental Drivers for Transmission. *Plos Neglected Tropical Diseases*, *10*(1), e0004405. https://doi.org/10.1371/journal.pntd.0004405

Lee, C.-Y., Thompson, G. R., Hastey, C. J., Hodge, G., Lunetta, J. M., Pappagianis, D., & Heinrich, V. (2015). Coccidioides Endospores and Spherules Draw Strong Chemotactic, Adhesive, and Phagocytic Responses by Individual Human Neutrophils. *PLoS One*, *10*(6), e0129522. https://doi.org/10.1371/journal.pone.0129522

Lee, L., Agwu, A. L., & Hutton, N. (2012). Severe Primary HSV-2 in a Perinatal HIV-Infected Woman With Advanced Immunosuppression. *Case Reports in Medicine*, *2012*, 1-3. https://doi.org/10.1155/2012/346039

Lee, M. J., Liu, H., Barker, B. M., Snarr, B. D., Gravelat, F. N., Abdallah, Q. A., Gavino, C., Baistrocchi, S. R., Ostapska, H., Xiao, T., Ralph, B., Solis, N. V., Lehoux, M., Baptista, S. D., Thammahong, A., Cerone, R. P., Kaminskyj, S. G. W., Guiot, M.-C., Latgé, J. P.,…Sheppard, D. C. (2015). The Fungal Exopolysaccharide Galactosaminogalactan Mediates Virulence by Enhancing Resistance to Neutrophil Extracellular Traps. *PLOS Pathogens*, *11*(10), e1005187. https://doi.org/10.1371/journal.ppat.1005187

Lee, S.-y., Ko, H. C., Kim, S.-I., Lee, Y., & Son, B.-c. (2019). Progressive Multifocal Leukoencephalopathy Diagnosed by Brain Biopsy, Not by the DNA Test for JC Virus. *Asian Journal of Neurosurgery*, *14*(01), 240-244. https://doi.org/10.4103/ajns.ajns_243_17

Lee, S. J., Jeong, H. J., Lee, J. E., Lee, J. S., Xuan, Y. H., Kong, H.-H., Chung, D.-I., Ock, M., & Yu, H. S. (2006). Molecular Characterization of Acanthamoeba Isolated From Amebic Keratitis Related to Orthokeratology Lens Overnight Wear. *The Korean Journal of Parasitology*, *44*(4), 313. https://doi.org/10.3347/kjp.2006.44.4.313

Lefebvre, M. N., & Harty, J. T. (2020). You Shall Not Pass: Memory CD8 T Cells in Liver-Stage Malaria. *Trends in Parasitology*, *36*(2), 147-157. https://doi.org/10.1016/j.pt.2019.11.004

Lei, N., Liu, F., Ren, C., Shen, J., & Liu, M. (2019). An Efficient Schistosoma Japonicum Bivalent Membrane Protein Antigen DNA Vaccine Against Schistosomiasis in Mice. *Medical Science Monitor*, *25*, 9319-9326. https://doi.org/10.12659/msm.919195

Leidal, A. M., Lee, P. W., & McCormick, C. (2012). Viral Subversion of Autophagy Impairs Oncogene-Induced Senescence. *Autophagy*, *8*(7), 1138-1140. https://doi.org/10.4161/auto.20340

Letesson, J. J., Lestrate, P., Delrue, R. M., Danese, I., Bellefontaine, F., Frétin, D., Taminiau, B., Tibor, A., Dricot, A., Deschamps, C., Haine, V., Leonard, S., Laurent, T., Mertens, P., Vandenhaute, J., & Bolle, X. D. (2002). Fun Stories About Brucella: The "Furtive Nasty Bug". *Veterinary Microbiology*, *90*(1-4), 317-328. https://doi.org/10.1016/s0378-1135(02)00208-0

Lewis, R. M., Markowitz, L. E., Panicker, G., & Unger, E. R. (2018). Seroprevalence of Human Papillomavirus 6/11/16/18 Among Self-Identified Gay/Bisexual Men Who Have Sex With Men, Men Who Have Sex With Women, and Females, United States, 2003–2010. *Clinical Infectious Diseases*, *69*(6), 1011-1018. https://doi.org/10.1093/cid/ciy1002

Li, J., Chai, Q., Zhang, Y., Li, B., Wang, J., Qiu, X. B., & Liu, C. H. (2015). <i>Mycobacterium Tuberculosis</I> McE3E Suppresses Host Innate Immune Responses by Targeting ERK1/2 Signaling. *The Journal of Immunology*, *194*(8), 3756-3767. https://doi.org/10.4049/jimmunol.1402679

Li, J., Liu, L., Chen, D., Yin, W., & Huang, J. (2023). Trained Immunity From the Perspective of <i>Plasmodium</I> Infection. *European Journal of Immunology*, *53*(6). https://doi.org/10.1002/eji.202250268

Li, L., Gu, B., Zhou, F., Chi, J., Wang, F., Peng, G., Xie, F., Qing, J., Feng, D., Lu, S., & Yao, K. (2011). Human Herpesvirus 6 Suppresses T Cell Proliferation Through Induction of Cell Cycle Arrest in Infected Cells in the G <sub>2</Sub> /M Phase. *J Virol*, *85*(13), 6774-6783. https://doi.org/10.1128/jvi.02577-10

Li, N., Zhang, C., Li, B., Liu, X., Huang, Y., Xu, S., & Gu, L. (2012). Unique Iron Coordination in Iron-Chelating Molecule Vibriobactin Helps Vibrio Cholerae Evade Mammalian Siderocalin-Mediated Immune Response. *Journal of Biological Chemistry*, *287*(12), 8912-8919. https://doi.org/10.1074/jbc.m111.316034

Liacini, A., Seamone, M. E., Muruve, D. A., & Tibbles, L. A. (2010). Anti-Bk Virus Mechanisms of Sirolimus and Leflunomide Alone and in Combination: Toward a New Therapy for BK Virus Infection. *Transplantation*, *90*(12), 1450-1457. https://doi.org/10.1097/tp.0b013e3182007be2

Libbey, J. E., Cusick, M. F., & Fujinami, R. S. (2013). Role of Pathogens in Multiple Sclerosis. *International Reviews of Immunology*, *33*(4), 266-283. https://doi.org/10.3109/08830185.2013.823422

Lidenge, S. J., Tso, F. Y., Ngalamika, O., Ngowi, J. R., Mortazavi, Y., Kwon, E. H., Shea, D. M., Minhas, V., Mwaiselage, J., Wood, C., & West, J. T. (2018). Similar Immunological Profiles Between African Endemic and Human Immunodeficiency Virus Type 1–Associated Epidemic Kaposi Sarcoma (KS) Patients Reveal the Primary Role of KS-Associated Herpesvirus in KS Pathogenesis. *The Journal of Infectious Diseases*, *219*(8), 1318-1328. https://doi.org/10.1093/infdis/jiy654

Lima, T. S., & Lodoen, M. B. (2019). Mechanisms of Human Innate Immune Evasion by Toxoplasma Gondii. *Frontiers in Cellular and Infection Microbiology*, *9*. https://doi.org/10.3389/fcimb.2019.00103

Lin, X., & Heitman, J. (2006). The Biology of The<i>Cryptococcus Neoformans</I>Species Complex. *Annual Review of Microbiology*, *60*(1), 69-105. https://doi.org/10.1146/annurev.micro.60.080805.142102

Lin, X. a., Sun, A., Ruan, P., Zhang, Z., & Yan, J. (2011). Characterization of Conserved Combined T and B Cell Epitopes in Leptospira interrogansMajor Outer Membrane Proteins OmpL1 and LipL41. *BMC Microbiology*, *11*(1). https://doi.org/10.1186/1471-2180-11-21

Lin, Y., Xiang, Z., & He, Y. (2015). Ontology-Based Representation and Analysis of Host-Brucella Interactions. *Journal of Biomedical Semantics*, *6*(1). https://doi.org/10.1186/s13326-015-0036-y

Lingel, A., Ehlers, E., Wang, Q., Cao, M., Wood, C., Lin, R., & Zhang, L. (2016). Kaposi's Sarcoma-Associated Herpesvirus Reduces Cellular Myeloid Differentiation Primary-Response Gene 88 (MyD88) Expression via Modulation of Its RNA. *J Virol*, *90*(1), 180-188. https://doi.org/10.1128/jvi.02342-15

Linnstaedt, S. D., Gottwein, E., Skalsky, R. L., Luftig, M. A., & Cullen, B. R. (2010). Virally Induced Cellular MicroRNA miR-155 Plays a Key Role in B-Cell Immortalization by Epstein-Barr Virus. *J Virol*, *84*(22), 11670-11678. https://doi.org/10.1128/jvi.01248-10

LiPuma, J. J. (2001). <i>Burkholderia Cepacia</I>complex: A Contraindication to Lung Transplantation in Cystic Fibrosis? *Transplant Infectious Disease*, *3*(3), 149-160. https://doi.org/10.1034/j.1399-3062.2001.003003149.x

LiPuma, J. J. (2005). Update on the Burkholderia Cepacia Complex. *Current Opinion in Pulmonary Medicine*, *11*(6), 528-533. https://doi.org/10.1097/01.mcp.0000181475.85187.ed

Liu, H.-C., Hsu, J. Y., & Chang, K.-M. (2009). Strongyloides Stercoralis Hyperinfection Presenting With Symptoms Mimicking Acute Exacerbation of Chronic Obstructive Pulmonary Disease. *Journal of the Chinese Medical Association*, *72*(8), 442-445. https://doi.org/10.1016/s1726-4901(09)70403-4

Liu, W., MacDonald, M., & You, J. (2016). Merkel Cell Polyomavirus Infection and Merkel Cell Carcinoma. *Current Opinion in Virology*, *20*, 20-27. https://doi.org/10.1016/j.coviro.2016.07.011

Looker, K. J., Elmes, J., Gottlieb, S. L., Schiffer, J. T., Vickerman, P., Turner, K., & Boily, M. C. (2017). Effect of HSV-2 Infection on Subsequent HIV Acquisition: An Updated Systematic Review and Meta-Analysis. *The Lancet Infectious Diseases*, *17*(12), 1303-1316. https://doi.org/10.1016/s1473-3099(17)30405-x

Looker, K. J., Magaret, A., May, M., Turner, K., Vickerman, P., Gottlieb, S. L., & Newman, L. (2015). Global and Regional Estimates of Prevalent and Incident Herpes Simplex Virus Type 1 Infections in 2012. *PLoS One*, *10*(10), e0140765. https://doi.org/10.1371/journal.pone.0140765

Lu, B., Viscidi, R. P., Lee, J. H., Wu, Y., Villa, L. L., Lazcano-Ponce, E., Roberto, J. C. d. S., Baggio, M. L., Quiterio, M., Salmerón, J., Smith, D. C., Abrahamsen, M., Papenfuss, M., Stockwell, H., & Giuliano, A. R. (2011). Human Papillomavirus (HPV) 6, 11, 16, and 18 Seroprevalence Is Associated With Sexual Practice and Age: Results From the Multinational HPV Infection in Men Study (<i>HIM</I> Study). *Cancer Epidemiology Biomarkers & Prevention*, *20*(5), 990-1002. https://doi.org/10.1158/1055-9965.epi-10-1160

Lü, F., Weidmer, A., Liu, C. G., Volinia, S., Croce, C. M., & Lieberman, P. M. (2008). Epstein-Barr Virus-Induced miR-155 Attenuates NF-κB Signaling and Stabilizes Latent Virus Persistence. *J Virol*, *82*(21), 10436-10443. https://doi.org/10.1128/jvi.00752-08

Lundqvist, A., Isa, A., Tolfvenstam, T., Kvist, G., & Broliden, K. (2005). High Frequency of Parvovirus B19 DNA in Bone Marrow Samples From Rheumatic Patients. *Journal of Clinical Virology*, *33*(1), 71-74. https://doi.org/10.1016/j.jcv.2004.11.011

Lupetti, A., Tavanti, A., Davini, P., Ghelardi, E., Corsini, V., Merusi, I., Boldrini, A., Campa, M., & Senesi, S. (2002). Horizontal Transmission Of <i>Candida Parapsilosis</I> Candidemia in a Neonatal Intensive Care Unit. *Journal of Clinical Microbiology*, *40*(7), 2363-2369. https://doi.org/10.1128/jcm.40.7.2363-2369.2002

Luplertlop, N., Muangkaew, W., Pumeesat, P., Suwanmanee, S., & Singkum, P. (2019). Distribution of Scedosporium Species in Soil From Areas With High Human Population Density and Tourist Popularity in Six Geographic Regions in Thailand. *PLoS One*, *14*(1), e0210942. https://doi.org/10.1371/journal.pone.0210942

Ly, A., & Hansen, D. S. (2019). Development of B Cell Memory in Malaria. *Frontiers in Immunology*, *10*. https://doi.org/10.3389/fimmu.2019.00559

Lyon, G. M., Bravo, A. V., Espino, A., Lindsley, M. D., Gutierrez, R. E., Rodríguez, I., Corella, A. N. A., Carrillo, F., McNeil, M. M., Warnock, D. W., & Hajjeh, R. (2004). Histoplasmosis Associated With Exploring a Bat-Inhabited Cave in Costa Rica, 1998–1999. *American Journal of Tropical Medicine and Hygiene*, *70*(4), 438-442. https://doi.org/10.4269/ajtmh.2004.70.438

Macaluso, G., Grippi, F., Bella, S. D., Blanda, V., Gucciardi, F., Torina, A., Guercio, A., & Cannella, V. (2023). A Review on the Immunological Response Against Trypanosoma Cruzi. *Pathogens*, *12*(2), 282. https://doi.org/10.3390/pathogens12020282

Maggi, F., & Bendinelli, M. (2010). Human Anelloviruses and the Central Nervous System. *Reviews in Medical Virology*, *20*(6), 392-407. https://doi.org/10.1002/rmv.668

Maggi, R. G., Ericson, M. E., Mascarelli, P. E., Bradley, J. M., & Breitschwerdt, E. B. (2013). Bartonella Henselae Bacteremia in a Mother and Son Potentially Associated With Tick Exposure. *Parasites & Vectors*, *6*(1). https://doi.org/10.1186/1756-3305-6-101

Maghraby, A. S., Hamed, M. A., & Ali, S. A. (2010). Bioimmunological Responses to Schistosoma Mansoni and Fasciola Gigantica Worm Homogenates Either With or Without Saponin. *The Journal of Infection in Developing Countries*, *4*(05), 334-344. https://doi.org/10.3855/jidc.449

Magnet, A., Galván-Díaz, A. L., Fenoy, S., Izquierdo, F., Rueda, C., Vadillo, C. F., Pérez-Irezábal, J., Bandyopadhyay, K., Visvesvara, G. S., Silva, A. J. d., & Águila, C. d. (2012). Molecular Characterization of Acanthamoeba Isolated in Water Treatment Plants and Comparison With Clinical Isolates. *Parasitology Research*, *111*(1), 383-392. https://doi.org/10.1007/s00436-012-2849-2

Major, E. O. (2011). History and Current Concepts in the Pathogenesis of PML. *Cleveland Clinic Journal of Medicine*, *78*(11 suppl 2), S3-S7. https://doi.org/10.3949/ccjm.78.s2.02

Mak, S., Vélez, N., Castañeda, E., Escandón, P., & Group, C. (2015). The Fungus Among Us: Cryptococcus Neoformans and Cryptococcus Gattii Ecological Modeling for Colombia. *Journal of Fungi*, *1*(3), 332-344. https://doi.org/10.3390/jof1030332

Mancini, M. E., Alessiani, A., Donatiello, A., Didonna, A., D'Attoli, L., Faleo, S., Occhiochiuso, G., Carella, F., Taranto, P. D., Pace, L., Rondinone, V., Damato, A. M., Coppola, R., Pedarra, C., & Goffredo, E. (2023). Systematic Survey of Vibrio Spp. And Salmonella Spp. In Bivalve Shellfish in Apulia Region (Italy): Prevalence and Antimicrobial Resistance. *Microorganisms*, *11*(2), 450. https://doi.org/10.3390/microorganisms11020450

Mani, V., & Hudgins, E. (2021). Balamuthia Mandrillaris Encephalitis in an Uncontrolled Diabetic Patient. *Idcases*, *25*, e01174. https://doi.org/10.1016/j.idcr.2021.e01174

Marascio, N., Loria, M. T., Pavia, G., Peronace, C., Adams, N. J., Campolo, M., Divenuto, F., Lamberti, A. G., Giancotti, A., Barreca, G. S., Mazzitelli, M., Trecarichi, E. M., Torti, C., Perandin, F., Bisoffi, Z., Quirino, A., & Matera, G. (2023). Evaluation of IL-35, as a Possible Biomarker for Follow-Up After Therapy, in Chronic Human Schistosoma Infection. *Vaccines*, *11*(5), 995. https://doi.org/10.3390/vaccines11050995

Marimuthu, S. (2019). Development of a Real-Time PCR Assay for Pneumocystis Jirovecii on the Luminex ARIES® Platform. *Journal of Respiratory Infections*, *3*(1). https://doi.org/10.18297/jri/vol3/iss1/5

Mariya, S., Sumithra, T. G., Anusree, V. N., Amala, P. V., Reshma, K. J., Alex, S., & Sanil, N. K. (2021). Antimicrobial Resistance and Virulence Characteristics of Vibrio Vulnificus, Vibrio Parahaemolyticus and Vibrio Harveyi From Natural Disease Outbreaks of Marine/Estuarine Fishes. *Aquaculture*, *539*, 736608. https://doi.org/10.1016/j.aquaculture.2021.736608

Markowska-Szczupak, A., Wang, K., Rokicka, P., Endo, M., Wei, Z., Ohtani, B., Morawski, A. W., & Kowalska, E. (2015). The Effect of Anatase and Rutile Crystallites Isolated From Titania P25 Photocatalyst on Growth of Selected Mould Fungi. *Journal of Photochemistry and Photobiology B Biology*, *151*, 54-62. https://doi.org/10.1016/j.jphotobiol.2015.07.002

Marques, M., Ferreira, A., & Ribeiro, D. (2018). The Interplay Between Human Cytomegalovirus and Pathogen Recognition Receptor Signaling. *Viruses*, *10*(10), 514. https://doi.org/10.3390/v10100514

Marshall, E. E., Malouli, D., Hansen, S. G., Gilbride, R. M., Hughes, C. M., Ventura, A. B., Ainslie, E., Selseth, A. N., Ford, J. C., Burke, D., Kreklywich, C. N., Womack, J., Legasse, A. W., Axthelm, M. K., Kahl, C., Streblow, D. N., Edlefsen, P. T., Picker, L. J., & Früh, K. (2019). Enhancing Safety of Cytomegalovirus-Based Vaccine Vectors by Engaging Host Intrinsic Immunity. *Science Translational Medicine*, *11*(501). https://doi.org/10.1126/scitranslmed.aaw2603

Matin, A., Siddiqui, R., Jayasekera, S., & Khan, N. A. (2008). Increasing Importance Of <i>Balamuthia Mandrillaris</I>. *Clinical Microbiology Reviews*, *21*(3), 435-448. https://doi.org/10.1128/cmr.00056-07

Matos, A. C., Figueira, L., Matos, M., Pinto, M. d. L., & Coelho, A. C. (2018). Seroprevalence of Mycobacterium Avium Complex in Wild Mammals in the Iberian Peninsula. *Journal of the Hellenic Veterinary Medical Society*, *66*(3), 177. https://doi.org/10.12681/jhvms.15861

Mazzola, J. C., Saito, P. K., Yamakawa, R. H., Watanabe, M. A. E., Silva, W. V. d., Matta, A. C. G., & Borelli, S. D. (2015). Prevalence of Torque Teno Virus in Healthy Donors of Paraná State, Southern Brazil. *Revista Brasileira De Hematologia E Hemoterapia*, *37*(5), 336-340. https://doi.org/10.1016/j.bjhh.2015.07.005

McKinsey, D. S. (2021). Treatment and Prevention of Histoplasmosis in Adults Living With HIV. *Journal of Fungi*, *7*(6), 429. https://doi.org/10.3390/jof7060429

McKinsey, D. S., & McKinsey, J. P. (2011). Pulmonary Histoplasmosis. *Seminars in Respiratory and Critical Care Medicine*, *32*(06), 735-744. https://doi.org/10.1055/s-0031-1295721

McKinsey, D. S., Spiegel, R. A., Hutwagner, L., Stanford, J., Driks, M. R., Brewer, J., Gupta, M., Smith, D. L., O'Connor, M. C., & Dall, L. (1997). Prospective Study of Histoplasmosis in Patients Infected With Human Immunodeficiency Virus: Incidence, Risk Factors, and Pathophysiology. *Clinical Infectious Diseases*, *24*(6), 1195-1203. https://doi.org/10.1086/513653

McManus, D. P., Dunne, D. W., Sacko, M., Utzinger, J., Vennervald, B. J., & Zhou, X. N. (2018). Schistosomiasis. *Nature Reviews Disease Primers*, *4*(1). https://doi.org/10.1038/s41572-018-0013-8

McManus, D. P., & Loukas, A. (2008). Current Status of Vaccines for Schistosomiasis. *Clinical Microbiology Reviews*, *21*(1), 225-242. https://doi.org/10.1128/cmr.00046-07

Melo, T. T. d., Araújo, J. M. d., Durães, F. d. V., Caliari, M. V., Oliveira, S. C., Coelho, P. M. Z., & Fonseca, C. T. (2010). Immunization With Newly Transformed <i>Schistosoma Mansoni</I> Schistosomula Tegument Elicits Tegument Damage, Reduction in Egg and Parasite Burden. *Parasite Immunology*, *32*(11-12), 749-759. https://doi.org/10.1111/j.1365-3024.2010.01244.x

Mendoza, N., Noël, P., & Blair, J. E. (2015). Diagnosis, Treatment, and Outcomes of Coccidioidomycosis in Allogeneic Stem Cell Transplantation. *Transplant Infectious Disease*, *17*(3), 380-388. https://doi.org/10.1111/tid.12372

Michael, K. M., Waterboer, T., Pfister, H., Gariglio, M., Majewski, S., Fãvre, M., & Pawlita, M. (2010). Seroreactivity of 38 Human Papillomavirus Types in Epidermodysplasia Verruciformis

Patients, Relatives, and Controls. *Journal of Investigative Dermatology*, *130*(3), 841-848. https://doi.org/10.1038/jid.2009.356

Miller, A. C., Arakkal, A. T., Koeneman, S. H., Cavanaugh, J., Thompson, G. R., Baddley, J. W., & Polgreen, P. M. (2022). Frequency and Duration Of, and Risk Factors For, Diagnostic Delays Associated With Histoplasmosis. *Journal of Fungi*, *8*(5), 438. https://doi.org/10.3390/jof8050438

Miller, M. B., Hendren, R., & Gilligan, P. H. (2004). Posttransplantation Disseminated Coccidioidomycosis Acquired From Donor Lungs. *Journal of Clinical Microbiology*, *42*(5), 2347-2349. https://doi.org/10.1128/jcm.42.5.2347-2349.2004

Minero, M. V., Marín, M., Cercenado, E., Martín-Rabadán, P., Bouza, E., & Muñóz, P. (2009). Nocardiosis at the Turn of the Century. *Medicine*, *88*(4), 250-261. https://doi.org/10.1097/md.0b013e3181afa1c8

Mitchell, A. B., & Glanville, A. R. (2021). The Impact of Resistant Bacterial Pathogens Including Pseudomonas Aeruginosa and Burkholderia on Lung Transplant Outcomes. *Seminars in Respiratory and Critical Care Medicine*, *42*(03), 436-448. https://doi.org/10.1055/s-0041-1728797

Mizuno, T., Debnath, A. K., & Miyoshi, S. i. (2020). Hemolysin Of<i>Vibrio</I>Species. https://doi.org/10.5772/intechopen.88920

Mohandas, V., & Ballal, M. (2011). Distribution of Candida Species in Different Clinical Samples and Their Virulence: Biofilm Formation, Proteinase and Phospholipase Production: A Study on Hospitalized Patients in Southern India. *Journal of Global Infectious Diseases*, *3*(1), 4. https://doi.org/10.4103/0974-777x.77288

Mooij, S. H., Klis, F. v. d., Marianne, A. B. v. d. S., Schepp, R. M., Speksnijder, A., Bogaards, J. A., Melker, H. E. d., Vries, H. J. d., Snijders, P. J., & Maarten, F. S. v. d. L. (2013). Seroepidemiology of High-Risk HPV in HIV-Negative and HIV-Infected MSM: The H2M Study. *Cancer Epidemiology Biomarkers & Prevention*, *22*(10), 1698-1708. https://doi.org/10.1158/1055-9965.epi-13-0460

Mooney, J. P., Lee, S. J., Lokken, K. L., Nanton, M. R., Nuccio, S. P., McSorley, S. J., & Tsolis, R. M. (2015). Transient Loss of Protection Afforded by a Live Attenuated Non-Typhoidal Salmonella Vaccine in Mice Co-Infected With Malaria. *Plos Neglected Tropical Diseases*, *9*(9), e0004027. https://doi.org/10.1371/journal.pntd.0004027

Moore, P. S., & Chang, Y. (2003). Kaposi's Sarcoma–Associated Herpesvirus Immunoevasion and Tumorigenesis: Two Sides of the Same Coin? *Annual Review of Microbiology*, *57*(1), 609-639. https://doi.org/10.1146/annurev.micro.57.030502.090824

Moormann, A. M. (2009). How Might Infant and Paediatric Immune Responses Influence Malaria Vaccine Efficacy? *Parasite Immunology*, *31*(9), 547-559. https://doi.org/10.1111/j.1365-3024.2009.01137.x

Moradimotlagh, A., Chen, S., Koohbor, S., Moon, K. M., Foster, L. J., Reiner, N. E., & Nandan, D. (2023). Leishmania Infection Upregulates and Engages Host Macrophage Argonaute 1, and System-Wide Proteomics Reveals Argonaute 1-Dependent Host Response. *Frontiers in Immunology*, *14*. https://doi.org/10.3389/fimmu.2023.1287539

Morales-Suárez-Varela, M., Peraita-Costa, I., & González, A. L. (2019). Supplementation With Micronutrients and Schistosomiasis: Systematic Review and Meta-Analysis. *Pathogens and Global Health*, *113*(3), 101-108. https://doi.org/10.1080/20477724.2019.1603902

Moran-Gilad, J., Mentasti, M., Lazarovitch, T., Huberman, Z., Stocki, T., Sadik, C., Shahar, T., Anis, E., Valinsky, L., Harrison, T. G., & Grotto, I. (2014). Molecular Epidemiology of Legionnaires‘ Disease in Israel. *Clinical Microbiology and Infection*, *20*(7), 690-696. https://doi.org/10.1111/1469-0691.12425

Morenikeji, O. A., Adeleye, O., Omoruyi, E. C., & Oyeyemi, O. T. (2016). Anti-<i>Schistosoma</I>IgG Responses In<i>Schistosoma Haematobium</I>single and Concomitant Infection With Malaria Parasites. *Pathogens and Global Health*, *110*(2), 74-78. https://doi.org/10.1080/20477724.2016.1174499

Moseley, P. T., Klenerman, P., & Kadambari, S. (2022). Indirect Effects of Cytomegalovirus Infection: Implications for Vaccine Development. *Reviews in Medical Virology*, *33*(1). https://doi.org/10.1002/rmv.2405

Mouhajir, A., Poirier, W., Angebault, C., Rahal, E., Bouabid, R., Bougnoux, M. E., Kobi, A., Zouhair, R., Bouchara, J.-P., & Giraud, S. (2020). Scedosporium Species in Soils From Various Biomes in Northwestern Morocco. *PLoS One*, *15*(2), e0228897. https://doi.org/10.1371/journal.pone.0228897

Mseddi, F., Sellami, A., Jarboui, M. A., Sellami, H., Makni, F., & Ayadi, A. (2011). First Environmental Isolations of Cryptococcus Neoformans and Cryptococcus Gattii in Tunisia and Review of Published Studies on Environmental Isolations in Africa. *Mycopathologia*, *171*(5), 355-360. https://doi.org/10.1007/s11046-010-9381-7

Mura, M., Ruffié, C., Combredet, C., Aliprandini, E., Formaglio, P., Chitnis, C. E., Amino, R., & Tangy, F. (2019). Recombinant Measles Vaccine Expressing Malaria Antigens Induces Long-Term Memory and Protection in Mice. *NPJ Vaccines*, *4*(1). https://doi.org/10.1038/s41541-019-0106-8

Mwachui, M. A., Crump, L., Hartskeerl, R. A., Zinsstag, J., & Hattendorf, J. (2015). Environmental and Behavioural Determinants of Leptospirosis Transmission: A Systematic Review. *Plos Neglected Tropical Diseases*, *9*(9), e0003843. https://doi.org/10.1371/journal.pntd.0003843

Mwenda, J. M., Soda, E., Weldegebriel, G., Katsande, R., Biey, J. N., Traore, T., Gouveia, L. d., Plessis, M. d., Gottberg, A. v., Antonio, M., Kwambana-Adams, B., Worwui, A., Gierke, R., Schwartz, S., Beneden, C. V., Cohen, A. L., Serhan, F., & Lessa, F. C. (2019). Pediatric Bacterial Meningitis Surveillance in the World Health Organization African Region Using the Invasive Bacterial Vaccine-Preventable Disease Surveillance Network, 2011–2016. *Clinical Infectious Diseases*, *69*(Supplement_2), S49-S57. https://doi.org/10.1093/cid/ciz472

Nakamura, T., Sato, Y., Watanabe, D., Ito, H., Shimonohara, N., Tsuji, T., Nakajima, N., Suzuki, Y., Matsuo, K., Nakagawa, H., Sata, T., & Katano, H. (2010). Nuclear Localization of Merkel Cell Polyomavirus Large T Antigen in Merkel Cell Carcinoma. *Virology*, *398*(2), 273-279. https://doi.org/10.1016/j.virol.2009.12.024

Nanakaly, H. T., & Hussen, B. M. (2019). Seroprevalence of Cytomegalovirus Among Voluntary Blood Donors in Erbil Province, North Iraq. *Zanco Journal of Pure and Applied Sciences*, *31*(3). https://doi.org/10.21271/zjpas.31.3.1

Nash, E. F., Coonar, A. S., Kremer, R., Tullis, E., Hutcheon, M., Singer, L. G., Keshavjee, S., & Chaparro, C. (2010). Survival of Burkholderia Cepacia Sepsis Following Lung Transplantation in Recipients With Cystic Fibrosis. *Transplant Infectious Disease*, *12*(6), 551-554. https://doi.org/10.1111/j.1399-3062.2010.00525.x

Nasrallah, G. K., Dargham, S. R., Harfouche, M., & Abu-Raddad, L. J. (2020). Seroprevalence of Herpes Simplex Virus Types 1 and 2 in Indian and Filipino Migrant Populations in Qatar: A Cross-Sectional Survey. *Eastern Mediterranean Health Journal*, *26*(5), 609-615. https://doi.org/10.26719/2020.26.5.609

Nastke, M.-D., Becerra, A., Yin, L., Domínguez-Amorocho, O., Gibson, L., Stern, L. J., & Calvo-Calle, J. M. (2012). Human CD4<sup>+</sup>T Cell Response to Human Herpesvirus 6. *J Virol*, *86*(9), 4776-4792. https://doi.org/10.1128/jvi.06573-11

Natarajan, P., & Narayanan, S. (2011). Mitogen-Activated Protein Kinases Mediate the Production of B-Cell Lymphoma 2 Protein by Mycobacterium Tuberculosis in Monocytes. *Biochemistry (Moscow)*, *76*(8), 938-950. https://doi.org/10.1134/s0006297911080104

Navarro-Navarro, M., Ruiz-Bustos, P., Valencia, D., Robles-Zepeda, R. E., Ruíz-Bustos, E., Virués, C., Hernández, J., Domínguez, Z., & Velázquez, C. (2013). Antibacterial Activity of Sonoran Propolis and Some of Its Constituents Against Clinically Significant<i>Vibrio</I>Species. *Foodborne Pathogens and Disease*, *10*(2), 150-158. https://doi.org/10.1089/fpd.2012.1318

Nefzi, F., Lambert, C., Gautheret-Dejean, A., Fisson, S., Khebizi, Q., Khélif, A., Agut, H., & Aouni, M. (2016). Cytokine and Cellular Responses to Human Herpesvirus-6B in Patients With B-acute Lymphoblastic Leukemia. *Microbiology and Immunology*, *60*(11), 770-777. https://doi.org/10.1111/1348-0421.12452

Negi, M., Sepolia, N., Panwar, S., Kumar, M., Singla, J., & Aggarwal, R. (2019). Prevalence of Oral Parameters in Smokeless Tobacco-Associated Precancer. *Journal of Family Medicine and Primary Care*, *8*(12), 3956. https://doi.org/10.4103/jfmpc.jfmpc_667_19

Netea, M. G., Warris, A., Jos, W. M. v. d. M., Fenton, M. J., Verver-Janssen, T. J. G., Jacobs, L., Andresen, T. K., Verweij, P. E., & Kullberg, B. J. (2003). <i>Aspergillus Fumigatus</I>Evades Immune Recognition During Germination Through Loss of Toll-Like Receptor-4–Mediated Signal Transduction. *The Journal of Infectious Diseases*, *188*(2), 320-326. https://doi.org/10.1086/376456

Nevez, G., Chabé, M., Rabodonirina, M., Virmaux, M., Dei-Cas, E., Hauser, P. M., & Totet, A. (2008). Nosocomial<i>Pneumocystis Jirovecii</I>infections. *Parasite*, *15*(3), 359-365. https://doi.org/10.1051/parasite/2008153359

Ngalamika, O., & Munsaka, S. (2020). Cells of the Innate and Adaptive Immune Systems in Kaposi's Sarcoma. *Journal of Immunology Research*, *2020*, 1-8. https://doi.org/10.1155/2020/8852221

Nicoli, F., Mantelli, B., Gallerani, E., Telatin, V., Bonazzi, I., Marconi, P., Gavioli, R., Gabrielli, L., Lazzarotto, T., Barzon, L., Palù, G., & Caputo, A. (2020). HPV-Specific Systemic Antibody Responses and Memory B Cells Are Independently Maintained Up to 6 Years and in a Vaccine-Specific Manner Following Immunization With Cervarix and Gardasil in Adolescent and Young Adult Women in Vaccination Programs in Italy. *Vaccines*, *8*(1), 26. https://doi.org/10.3390/vaccines8010026

Niedelman, W., Gold, D., Rosowski, E. E., Sprokholt, J. K., Lim, D., Arenas, A. F., Melo, M. B., Spooner, E., Yaffe, M. B., & Saeij, J. P. J. (2012). The Rhoptry Proteins ROP18 and ROP5 Mediate Toxoplasma Gondii Evasion of the Murine, but Not the Human, Interferon-Gamma Response. *PLOS Pathogens*, *8*(6), e1002784. https://doi.org/10.1371/journal.ppat.1002784

Nielsen, K., Obaldia, A. L. D., & Heitman, J. (2007). <i>Cryptococcus Neoformans</I> Mates on Pigeon Guano: Implications for the Realized Ecological Niche and Globalization. *Eukaryotic Cell*, *6*(6), 949-959. https://doi.org/10.1128/ec.00097-07

Niknam, N., Malhotra, P., Kim, A., & Koenig, S. (2017). Disseminated Histoplasmosis Presenting as Diabetic Keto-Acidosis in an Immunocompetent Patient. *BMJ Case Reports*, *2017*, bcr2016217915. https://doi.org/10.1136/bcr-2016-217915

Nisbet, A. I., Omuse, G., Revathi, G., & Adam, R. D. (2018). Seroprevalence Data at a Private Teaching Hospital in Kenya: An Examination of Toxoplasma Gondii, Cytomegalovirus, Rubella, Hepatitis A, and Entamoeba Histolytica. *PLoS One*, *13*(10), e0204867. https://doi.org/10.1371/journal.pone.0204867

Niu, S., & Zhao, L. (2022). Metagenomic Next-Generation Sequencing Clinches the Diagnosis of Legionella Pneumonia in a Patient With Acute Myeloid Leukemia: A Case Report and Literature Review. *Frontiers in Cellular and Infection Microbiology*, *12*. https://doi.org/10.3389/fcimb.2022.924597

Niyyati, M., Lorenzo-Morales, J., Rezaeian, M., Martín-Navarro, C. M., Haghi, A. M., Maciver, S. K., & Valladares, B. (2009). Isolation of Balamuthia Mandrillaris From Urban Dust, Free of Known Infectious Involvement. *Parasitology Research*, *106*(1), 279-281. https://doi.org/10.1007/s00436-009-1592-9

Njeru, J., Henning, K., Pletz, M. W., Heller, R., & Neubauer, H. (2016). Q Fever Is an Old and Neglected Zoonotic Disease in Kenya: A Systematic Review. *BMC Public Health*, *16*(1). https://doi.org/10.1186/s12889-016-2929-9

Nnaoma, C., Chika-Nwosuh, O., & Engell, C. (2019). The Worm That Clogs the Lungs: Strongyloides Hyper-Infection Leading to Fatal Acute Respiratory Distress Syndrome (ARDS). *American Journal of Case Reports*, *20*, 377-380. https://doi.org/10.12659/ajcr.914640

Nogaro, S., Hafalla, J. C. R., Walther, B., Remarque, E. J., Tetteh, K. K. A., Conway, D. J., Riley, E. M., & Walther, M. (2011). The Breadth, but Not the Magnitude, of Circulating Memory B Cell Responses to P. Falciparum Increases With Age/Exposure in an Area of Low Transmission. *PLoS One*, *6*(10), e25582. https://doi.org/10.1371/journal.pone.0025582

Norberg, A. N., Paulo Roberto Blanco Moreira, N., Manhães, F. C., Mangiavacchi, B. M., Faial, L. C. M., Filho, R. M. F., Borges, T. R. B., & Claudia Caixeta Franco Andrade, C. (2023). Steroid Therapy in COVID-19 as a Triggering Factor for Strongyloides Stercoralis Hyperinfection. https://doi.org/10.56238/innovhealthknow-037

Novosel, D., Lipej, Z., Čubrić-Ćurik, V., & Jungić, A. (2012). Presence of Torque Teno Sus Virus in Porcine Circovirus Type 2-associated Disease in Croatia. *Veterinary Record*, *171*(21), 529-529. https://doi.org/10.1136/vr.100887

Nucci, M., & Anaissie, E. (2007). <i>Fusarium</I> Infections in Immunocompromised Patients. *Clinical Microbiology Reviews*, *20*(4), 695-704. https://doi.org/10.1128/cmr.00014-07

Nukaga, S., Murakami, H., Yagi, K., Satomi, R., Oyama, T., Maeshima, A., & Oyamada, Y. (2021). A Solitary Pulmonary Nodule Caused by Mycobacterium Avium With Pleural Effusion and Pleuritis After Transbronchial Biopsy: A Case Report. *Journal of Medical Case Reports*, *15*(1). https://doi.org/10.1186/s13256-021-02929-9

O'Bryan, T. (2005). Pseudallescheriasis in the 21st Century. *Expert Review of Anti-Infective Therapy*, *3*(5), 765-773. https://doi.org/10.1586/14787210.3.5.765

O'Sullivan, M. G., Anderson, D. C., Fikes, J. D., Bain, F. T., Carlson, C. S., Green, S. W., Young, N. S., & Brown, K. (1994). Identification of a Novel Simian Parvovirus in Cynomolgus Monkeys With Severe Anemia. A Paradigm of Human B19 Parvovirus Infection. *Journal of Clinical Investigation*, *93*(4), 1571-1576. https://doi.org/10.1172/jci117136

Obeagu, E. I., & Obeagu, G. U. (2024). The Impact of Maternal Malaria on Adaptive Immune Responses in Offspring. *International Journal of Medical Sciences and Pharma Research*, *10*(2), 10-17. https://doi.org/10.22270/ijmspr.v10i2.93

Okamoto, H. (2009). History of Discoveries and Pathogenicity of TT Viruses. 1-20. https://doi.org/10.1007/978-3-540-70972-5_1

Okwor, I., & Uzonna, J. E. (2013). The Immunology of Leishmania/Hiv Co-Infection. *Immunol Res*, *56*(1), 163-171. https://doi.org/10.1007/s12026-013-8389-8

Oliveira, I., Torrico, F., Muñóz, J., & Gascón, J. (2010). Congenital Transmission of Chagas Disease: A Clinical Approach. *Expert Review of Anti-Infective Therapy*, *8*(8), 945-956. https://doi.org/10.1586/eri.10.74

Olivier, M., Badaró, R., Medrano, F. J., & Moreno, J. (2003). The Pathogenesis Of<i>Leishmania</I>/Hiv Co-Infection: Cellular and Immunological Mechanisms. *Annals of Tropical Medicine and Parasitology*, *97*(sup1), 79-98. https://doi.org/10.1179/000349803225002561

Olleros, M. L., Chávez-Galán, L., Segueni, N., Bourigault, M. L., Vesin, D., Kruglov, A., Drutskaya, M. S., Bisig, R., Ehlers, S., Aly, S., Walter, K., Kuprash, D. V., Chouchkova, M., Kozlov, S., Erard, F., Ryffel, B., Quesniaux, V., Nedospasov, S. A., & García, I. (2015). Control of Mycobacterial Infections in Mice Expressing Human Tumor Necrosis Factor (TNF) but Not Mouse TNF. *Infect Immun*, *83*(9), 3612-3623. https://doi.org/10.1128/iai.00743-15

Omosigho, O., Izevbuwa, O. E., Ibrahim, O., Adetunji, O., & Otojareri, K. (2022). Seroprevalence of Cytomegalovirus Among Human Immunodeficiency Virus (HIV) Postiev Patients Attending HAART Clinic in Kawar State, North-Central Nigeria. *Microbes and Infectious Diseases*, *0*(0), 0-0. https://doi.org/10.21608/mid.2022.151771.1355

Onaiwu, C. O., Velagapudi, M., Sarsam, L., Utley, L., Bricker, L., Bendi, V. S., & Vivekanandan, R. (2017). Rare Multidrug-Resistant Pulmonary Nocardiosis in AIDS. *Cureus*. https://doi.org/10.7759/cureus.1839

Ongrádi, J., Ablashi, D. V., Yoshikawa, T., Stercz, B., & Ogata, M. (2016). Roseolovirus-Associated Encephalitis in Immunocompetent and Immunocompromised Individuals. *Journal of Neurovirology*, *23*(1), 1-19. https://doi.org/10.1007/s13365-016-0473-0

Ordaya, E. E., Johnson, J. R., Drekonja, D., Niehans, G. E., & Kaka, A. S. (2021). Aspergillus Osteomyelitis Secondary to Chronic Necrotizing Pulmonary Aspergillosis in a Patient With Rheumatoid Arthritis. *Cureus*. https://doi.org/10.7759/cureus.17774

Orosz, E., & Posta, K. (2020). Genotyping of Acantamoeba Spp. From Rhisophere in Hungary. *Acta Microbiologica Et Immunologica Hungarica*, *67*(3), 171-175. https://doi.org/10.1556/030.66.2019.041

Ortíz, A. P., Tortolero-Luna, G., Romaguera, J., Pérez, C. M., González, D., Muñoz, C., González, L., Marrero, E., Suárez, E., Palefsky, J. M., Panicker, G., & Unger, E. R. (2018). Seroprevalence of HPV 6, 11, 16 and 18 and Correlates of Exposure in Unvaccinated Women Aged 16–64 Years in Puerto Rico. *Papillomavirus Research*, *5*, 109-113. https://doi.org/10.1016/j.pvr.2018.03.006

Ostermann, E., Pawletko, K., Indenbirken, D., Schumacher, U., & Brune, W. (2015). Stepwise Adaptation of Murine Cytomegalovirus to Cells of a Foreign Host for Identification of Host Range Determinants. *Medical Microbiology and Immunology*, *204*(3), 461-469. https://doi.org/10.1007/s00430-015-0400-7

Ota, M., Serada, S., Naka, T., & Mori, Y. (2014). MHC Class I Molecules Are Incorporated Into Human Herpesvirus-6 Viral Particles and Released Into the Extracellular Environment. *Microbiology and Immunology*, *58*(2), 119-125. https://doi.org/10.1111/1348-0421.12121

Ouwendijk, W. J. D., & Georges, M. G. M. V. (2014). Pathogenesis of Varicelloviruses in Primates. *The Journal of Pathology*, *235*(2), 298-311. https://doi.org/10.1002/path.4451

Pakzad, I., Kalani, B. S., Asadollahi, P., Sadeghifard, N., & Kazemian, H. (2023). In Silico Study of the Proteins Involved in the Persistence of Brucella Spp. *Current Drug Discovery Technologies*, *20*(1). https://doi.org/10.2174/1570163819666220805161821

Parikh, S. R., Campbell, H., Bettinger, J. A., Harrison, L. H., Marshall, H., Martinón-Torres, F., Sáfadi, M. A. P., Shao, Z., Zhu, B., Gottberg, A. v., Borrow, R., Ramsay, M., & Ladhani, S. (2020). The Everchanging Epidemiology of Meningococcal Disease Worldwide and the Potential for Prevention Through Vaccination. *Journal of Infection*, *81*(4), 483-498. https://doi.org/10.1016/j.jinf.2020.05.079

Parish, J. M., & Blair, J. E. (2008). Coccidioidomycosis. *Mayo Clinic Proceedings*, *83*(3), 343-349. https://doi.org/10.4065/83.3.343

Park, S., Cheong, J., Kyi, K., Aranez, J. L., Abu-Farsakh, S., Whitney-Miller, C. L., Al-Judaibi, B., & Laryea, M. (2020). Cholestasis and Disseminated Histoplasmosis in a Psoriatic Patient on Infliximab: Case Report and Review of Literature. *BMC Gastroenterology*, *20*(1). https://doi.org/10.1186/s12876-020-01290-3

Park, S. J., & Mehrad, B. (2009). Innate Immunity To<i>Aspergillus</I>Species. *Clinical Microbiology Reviews*, *22*(4), 535-551. https://doi.org/10.1128/cmr.00014-09

Parsonnet, J. (1999). Fecal and Oral Shedding of ≪EMPH TYPE="ITAL">Helicobacter Pylori</EMPH> From Healthy Infected Adults. *JAMA*, *282*(23), 2240. https://doi.org/10.1001/jama.282.23.2240

Pattabiraman, C., Prasad, P., Sudarshan, S., George, A. K., Sreenivas, D., Rasheed, R., Ghosh, A., Pal, A., Hameed, S. K. S., Bandyopadhyay, B., Desai, A., & Ravi, V. (2022). Identification and Genomic Characterization of Parvovirus B19V Genotype 3 Viruses From Cases of Meningoencephalitis in West Bengal, India. *Microbiology Spectrum*, *10*(2). https://doi.org/10.1128/spectrum.02251-21

Paudel, A., Patel, N. S., Quang, J. Q., Casella, C., Sigal, A., Parajuli, P. B., Oladunjoye, O., Oke, I. O., Khanal, S., & Bhattarai, K. (2021). Rapidly Progressive Granulomatous Amoebic Encephalitis in a Diabetic Individual. *Cureus*. https://doi.org/10.7759/cureus.19336

Pearce, E. J., & MacDonald, A. S. (2002). The Immunobiology of Schistosomiasis. *Nature Reviews Immunology*, *2*(7), 499-511. https://doi.org/10.1038/nri843

Pedroso, R. d. S., Ferreira, J. C., & Cândido, R. C. (2009). The Isolation and Characterization of Virulence Factors of Cryptococcus Spp. From Saprophytic Sources in the City of Ribeirão Preto, São Paulo, Brazil. *Microbiological Research*, *164*(2), 221-227. https://doi.org/10.1016/j.micres.2007.01.002

Perio, M. A. d., Benedict, K., Williams, S., Niemeier-Walsh, C., Green, B. J., Coffey, C. C., Giuseppe, M. D., Toda, M., Park, J. H., Bailey, R. L., & Nett, R. J. (2021). Occupational Histoplasmosis:

Epidemiology and Prevention Measures. *Journal of Fungi*, *7*(7), 510. https://doi.org/10.3390/jof7070510

Pessenda, G., & Silva, J. S. (2020). Arginase and Its Mechanisms in <i>Leishmania</I> Persistence. *Parasite Immunology*, *42*(7). https://doi.org/10.1111/pim.12722

Pfaller, M. A., Jones, R. N., Doern, G. V., Sader, H. S., Messer, S. A., Houston, A., Coffman, S. L., & Hollis, R. J. (2000). Bloodstream Infections Due To <i>Candida</I> Species: SENTRY Antimicrobial Surveillance Program in North America and Latin America, 1997-1998. *Antimicrobial Agents and Chemotherapy*, *44*(3), 747-751. https://doi.org/10.1128/aac.44.3.747-751.2000

Pinho, J. M. R., Cardoso, F. C., Lopes, D. d. O., Pinheiro, C. S., Caliari, M. V., Oliveira, F. M. S., Leite, L. C. C., & Oliveira, S. C. (2009). Immunization With SmIg, a Novel Tegument Protein From<i>Schistosoma Mansoni</I>, Fails to Induce Protection in Mice but Reduces Liver Pathology. *Parasitology*, *137*(7), 1079-1088. https://doi.org/10.1017/s0031182009991387

Pissarra, J., Pagniez, J., Petitdidier, E., Séveno, M., Vigy, O., Bras-Gonçalves, R., Lemesre, J.-L., & Holzmüller, P. (2021). Proteomic Analysis of the Promastigote Secretome of Seven <i>Leishmania</I> Species. *Journal of Proteome Research*, *21*(1), 30-48. https://doi.org/10.1021/acs.jproteome.1c00244

Pitassi, L. H. U., Pedro Paulo Vissotto de Paiva, D., Scorpio, D. G., Drummond, M. R., Lania, B. G., Barjas-Castro, M. L., Gilioli, R., Colombo, S., Sowy, S., Breitschwerdt, E. B., Nicholson, W. L., & Paulo Eduardo Neves Ferreira, V. (2015). Bartonella Spp. Bacteremia in Blood Donors From Campinas, Brazil. *Plos Neglected Tropical Diseases*, *9*(1), e0003467. https://doi.org/10.1371/journal.pntd.0003467

Poester, F. P., Samartino, L., & Santos, R. L. (2013). Pathogenesis and Pathobiology of Brucellosis in Livestock. *Revue Scientifique Et Technique De L Oie*, *32*(1), 105-115. https://doi.org/10.20506/rst.32.1.2193

Poppe, L. K., Kankasa, C., Wood, C., & West, J. T. (2020). Relationships Between Maternal Antibody Responses and Early Childhood Infection With Kaposi Sarcoma-Associated Herpesvirus. *The Journal of Infectious Diseases*, *222*(10), 1723-1730. https://doi.org/10.1093/infdis/jiaa288

Potter, A., Banazis, M., Yang, R., Reid, S., & Fenwick, S. (2011). Prevalence of Coxiella Burnetii in Western Grey Kangaroos (Macropus Fuliginosus) in Western Australia. *Journal of Wildlife Diseases*, *47*(4), 821-828. https://doi.org/10.7589/0090-3558-47.4.821

Potter, A., Stephens, D., & Keulenaer, B. d. (2003). <i>Strongyloides</I>hyper-Infection: A Case for Awareness. *Annals of Tropical Medicine and Parasitology*, *97*(8), 855-860. https://doi.org/10.1179/000349803225002453

Pulgarin, J. A. H., Piedrahíta, J. A. A., Gómez, G. A., Palacio, J. F. S., Ordóñez, K., & Ramos, D. A. (2021). Closing Gaps in Histoplasmosis: Clinical Characteristics and Factors Associated With Probable/Histoplasmosis in HIV/AIDS Hospitalized Patients, a Retrospective Cross-Sectional Study in Two Tertiary Centers in Pereira, Colombia. *Aids Research and Therapy*, *18*(1). https://doi.org/10.1186/s12981-021-00377-5

Qin, Y., Zhou, G., Jiao, F., Cheng, C., Meng, C., Wang, L., Wu, S., Fan, C., Li, J., Zhou, B., Chu, Y. I., & Jiao, H. (2024). Brucella Mediates Autophagy, Inflammation, and Apoptosis to Escape Host Killing. *Frontiers in Cellular and Infection Microbiology*, *14*. https://doi.org/10.3389/fcimb.2024.1408407

Qiu, J., Söderlund-Venermo, M., & Young, N. S. (2017). Human Parvoviruses. *Clinical Microbiology Reviews*, *30*(1), 43-113. https://doi.org/10.1128/cmr.00040-16

Quesniaux, V., Jacobs, M., Allie, N., Grivennikov, S. I., Nedospasov, S. A., García, I., Olleros, M. L., Shebzukhov, Y. V., Kuprash, D. V., Vasseur, V., Rose, S., Court, N., Vacher, R., & Ryffel, B. (2010). TNF in Host Resistance to Tuberculosis Infection. 157-179. https://doi.org/10.1159/000289204

Radolf, J. D., Deka, R. K., Anand, A., Šmajs, D., Norgard, M. V., & Yang, X. (2016). Treponema Pallidum, the Syphilis Spirochete: Making a Living as a Stealth Pathogen. *Nature Reviews Microbiology*, *14*(12), 744-759. https://doi.org/10.1038/nrmicro.2016.141

Rafael, D., Tomer, E., & Kobiler, O. (2022). A Single Herpes Simplex Virus 1 Genome Reactivates From Individual Cells. *Microbiology Spectrum*, *10*(4). https://doi.org/10.1128/spectrum.01144-22

Rahman, S., Pierce, C. M., Waterboer, T., Rollison, D. E., Ingles, D. J., Torres, B. N., Michel, A., Sudenga, S. L., Pawlita, M., Villa, L. L., Lazcano-Ponce, E., Borenstein, A. R., Wang, W., & Giuliano, A. R. (2016). Seroprevalence of Cutaneous Human Papillomaviruses (HPVs) Among Men in the Multinational HPV Infection in Men Study. *Journal of General Virology*, *97*(12), 3291-3301. https://doi.org/10.1099/jgv.0.000620

Raju, R. S., Vinayak, K. N., Bapat, V. M., Balkisanji, A. P., & Chandrakant, P. S. (2013). Acute Human Parvovirus B19 Infection: Cytologic Diagnosis. *Indian Journal of Hematology and Blood Transfusion*, *30*(S1), 133-134. https://doi.org/10.1007/s12288-013-0287-7

Ramos, K. J., Quon, B. S., Psoter, K. J., Lease, E. D., Mayer-Hamblett, N., Aitken, M. L., & Goss, C. H. (2016). Predictors of Non-Referral of Patients With Cystic Fibrosisfor Lung Transplant Evaluation in the United States. *Journal of Cystic Fibrosis*, *15*(2), 196-203. https://doi.org/10.1016/j.jcf.2015.11.005

Randhawa, P., Zygmunt, D., Shapiro, R., Vats, A., Weck, K. E., Swalsky, P., & Finkelstein, S. (2003). Viral Regulatory Region Sequence Variations in Kidney Tissue Obtained From Patients With BK Virus Nephropathy. *Kidney International*, *64*(2), 743-747. https://doi.org/10.1046/j.1523-1755.2003.00132.x

Reinhold, I., Quiblier, C., Blaser, F., Bögeholz, J., Imkamp, F., Schuurmans, M. M., Soyka, M. B., Zbinden, R., & Mueller, N. J. (2024). Detection of <i>Scedosporium</I> Spp.: Colonizer or Pathogen? A Retrospective Analysis of Clinical Significance and Management in a Large Tertiary Center. *Medical Mycology*, *62*(2). https://doi.org/10.1093/mmy/myae002

Rengifo-Correa, L., Stephens, C. R., Morrone, J. J., Téllez-Rendón, J. L., & González-Salazar, C. (2017). Understanding Transmissibility Patterns of Chagas Disease Through Complex Vector–host Networks. *Parasitology*, *144*(6), 760-772. https://doi.org/10.1017/s0031182016002468

Renzette, N., Pokalyuk, C., Gibson, L., Bhattacharjee, B., Schleiss, M. R., Hamprecht, K., Yamamoto, A. Y., Mussi-Pinhata, M. M., Britt, W. J., Jensen, J. D., & Kowalik, T. F. (2015). Limits and Patterns of Cytomegalovirus Genomic Diversity in Humans. *Proceedings of the National Academy of Sciences*, *112*(30). https://doi.org/10.1073/pnas.1501880112

Reoma, L., Trindade, C., Monaco, M. C., Solis, J., García-Montojo, M., Vu, P. T., Johnson, K. R., Beck, E., Nair, G., Khan, O., Quezado, M., Hewitt, S. M., Reich, D. S., Childs, R., & Nath, A. (2019). Fatal Encephalopathy With Wild-type JC Virus and Ruxolitinib Therapy. *Annals of Neurology*, *86*(6), 878-884. https://doi.org/10.1002/ana.25608

Richards, G. P., Watson, M. A., & Parveen, S. (2005). Development of a Simple and Rapid Fluorogenic Procedure for Identification Of <i>Vibrionaceae</I> Family Members. *Applied and Environmental Microbiology*, *71*(7), 3524-3527. https://doi.org/10.1128/aem.71.7.3524-3527.2005

Richter, S. S., Galask, R. P., Messer, S. A., Hollis, R. J., Diekema, D. J., & Pfaller, M. A. (2005). Antifungal Susceptibilities Of <i>Candida</I> Species Causing Vulvovaginitis and Epidemiology of Recurrent Cases. *Journal of Clinical Microbiology*, *43*(5), 2155-2162. https://doi.org/10.1128/jcm.43.5.2155-2162.2005

Riley, T. P., Hellman, L. M., Gee, M. H., Mendoza, J. L., Alonso, J. A., Foley, K. C., Nishimura, M. I., Kooi, C. W. V., García, K. C., & Baker, B. M. (2018). T Cell Receptor Cross-Reactivity Expanded by Dramatic Peptide–MHC Adaptability. *Nature Chemical Biology*, *14*(10), 934-942. https://doi.org/10.1038/s41589-018-0130-4

Rist, M. J., Hibbert, K. M., Croft, N. P., Smith, C., Neller, M. A., Burrows, J. M., Miles, J. J., Purcell, A. W., Rossjohn, J., Gras, S., & Burrows, S. R. (2015). T Cell Cross-Reactivity Between a Highly Immunogenic EBV Epitope and a Self-Peptide Naturally Presented by HLA-B*18:01+ Cells. *The Journal of Immunology*, *194*(10), 4668-4675. https://doi.org/10.4049/jimmunol.1500233

Rizzo, R., & Luca, D. D. (2018). Human Herpesvirus 6A and 6B and NK Cells. *Acta Microbiologica Et Immunologica Hungarica*, *65*(2), 119-125. https://doi.org/10.1556/030.65.2018.010

Roca, F. J., Whitworth, L., Prag, H. A., Murphy, M. P., & Ramakrishnan, L. (2022). Tumor Necrosis Factor Induces Pathogenic Mitochondrial ROS in Tuberculosis Through Reverse Electron Transport. *Science*, *376*(6600). https://doi.org/10.1126/science.abh2841

Rodrigues, C. F., Rodrigues, M. E., Silva, S., & Henriques, M. (2017). Candida Glabrata Biofilms: How Far Have We Come? *Journal of Fungi*, *3*(1), 11. https://doi.org/10.3390/jof3010011

Roest, H.-J., Gelderen, B. v., Dinkla, A., Frangoulidis, D., Zijderveld, F. v., Rebel, J. M. J., & Keulen, L. v. (2012). Q Fever in Pregnant Goats: Pathogenesis and Excretion of Coxiella Burnetii. *PLoS One*, *7*(11), e48949. https://doi.org/10.1371/journal.pone.0048949

Rommereim, L. M., Bellini, V., Fox, B. A., Pètre, G., Rak, C., Touquet, B., Aldebert, D., Dubremetz, J. F., Cesbron-Delauw, M. F., Mercier, C., & Bzik, D. J. (2016). Phenotypes Associated With Knockouts of Eight Dense Granule Gene Loci (GRA2-9) in Virulent Toxoplasma Gondii. *PLoS One*, *11*(7), e0159306. https://doi.org/10.1371/journal.pone.0159306

Rönnberg, C., Lugaajju, A., Nyman, A., Hammar, U., Bottai, M., Lautenbach, M. J., Sundling, C., Kironde, F., & Persson, K. E. M. (2021). A Longitudinal Study of Plasma BAFF Levels in Mothers and Their Infants in Uganda, and Correlations With Subsets of B Cells. *PLoS One*, *16*(1), e0245431. https://doi.org/10.1371/journal.pone.0245431

Rose, C. E., Paciullo, C. A., Kelly, D. R., Dougherty, M. J., & Fleckenstein, L. (2009). Fatal Outcome of Disseminated Strongyloidiasis Despite Detectable Plasma and Cerebrospinal Levels of Orally Administered Ivermectin. *Journal of Parasitology Research*, *2009*, 1-4. https://doi.org/10.1155/2009/818296

Rosowski, E. E., Raffa, N., Knox, B. P., Golenberg, N., Keller, N. P., & Huttenlocher, A. (2018). Macrophages Inhibit Aspergillus Fumigatus Germination and Neutrophil-Mediated Fungal Killing. *PLOS Pathogens*, *14*(8), e1007229. https://doi.org/10.1371/journal.ppat.1007229

Ross, S. A., Michaels, M. G., Ahmed, A., Palmer, A. L., Sánchez, P. J., Bernstein, D. I., Feja, K. N., Stewart, A., Boppana, S. B., & Fowler, K. B. (2018). Contribution of Breastfeeding to False-

Positive Saliva Polymerase Chain Reaction for Newborn Congenital Cytomegalovirus Screening. *The Journal of Infectious Diseases*, *217*(10), 1612-1615. https://doi.org/10.1093/infdis/jiy057

Rougeron, A., Giraud, S., Alastruey-Izquierdo, A., Cano, J., Rainer, J., Mouhajir, A., Gal, S. L., Nevez, G., Meyer, W., & Bouchara, J.-P. (2017). Ecology of Scedosporium Species: Present Knowledge and Future Research. *Mycopathologia*, *183*(1), 185-200. https://doi.org/10.1007/s11046-017-0200-2

Ruan, Y., Wen, Z., Chen, K., Xi, J., Wu, B., Xu, Z., Jiang, M.-Z., Zhang, J., Chen, Y., & Liu, Q. (2023). Exogenous Interleukin-37 Alleviates Hepatitis With Reduced Dendritic Cells and Induced Regulatory T Cells in Acute Murine Cytomegalovirus Infection. *Journal of Immunology Research*, *2023*, 1-13. https://doi.org/10.1155/2023/1462048

Sakyi, S. A., Danquah, K. O., Ephraim, R. D., Enimil, A., Frimpong, V., Fondjo, L. A., & Darkoh, E. L. (2018). Evaluating the Contribution of <i>Nocardia</I> Spp. And <i>Mycobacterium Tuberculosis</I> to Pulmonary Infections Among HIV and Non-Hiv Patients at the Komfo Anokye Teaching Hospital, Ghana. *Canadian Journal of Infectious Diseases and Medical Microbiology*, *2018*, 1-7. https://doi.org/10.1155/2018/2910198

Saling, C., Gea-Banacloche, J., Trickett, J. S., & Blair, J. E. (2021). Coccidioidomycosis in Allogeneic Stem Cell Transplant Recipients: Case Series and Review of the Literature. *Journal of Fungi*, *7*(5), 339. https://doi.org/10.3390/jof7050339

Santos, C. d. S., & Brodskyn, C. u. I. (2014). The Role of CD4 and CD8â€‰T Cells in Human Cutaneous Leishmaniasis. *Frontiers in Public Health*, *2*. https://doi.org/10.3389/fpubh.2014.00165

Santos-Juanes, J., Fernández-Vega, I., Fuentes, N., Galache, C., Coto-Segura, P., Vivanco, B., Astudillo, A., & Martínez-Camblor, P. (2015). Merkel Cell Carcinoma and Merkel Cell Polyomavirus: A Systematic Review and Meta-Analysis. *British Journal of Dermatology*, *173*(1), 42-49. https://doi.org/10.1111/bjd.13870

Sarıgüzel, F. M., Berk, E., Koç, A. N., Sav, H., & Demir, G. (2015). Investigation of the Relationship Between Virulence Factors and Genotype of Candida Spp. Isolated From Blood Cultures. *The Journal of Infection in Developing Countries*, *9*(08), 857-864. https://doi.org/10.3855/jidc.5359

Schachtner, T., Müller, K., Stein, M., Diezemann, C., Sefrin, A., Babel, N., & Reinke, P. (2011). BK Virus-Specific Immunity Kinetics: A Predictor of Recovery From Polyomavirus BK-Associated Nephropathy. *American Journal of Transplantation*, *11*(11), 2443-2452. https://doi.org/10.1111/j.1600-6143.2011.03693.x

Schenk, T., Enders, M., Pollak, S., Hahn, R., & Huzly, D. (2009). High Prevalence of Human Parvovirus B19 DNA in Myocardial Autopsy Samples From Subjects Without Myocarditis or Dilative Cardiomyopathy. *Journal of Clinical Microbiology*, *47*(1), 106-110. https://doi.org/10.1128/jcm.01672-08

Scherer, E. M., Smith, R. A., Gallego, D. F., Carter, J. J., Wipf, G. C., Hoyos, M., Stern, M. E., Thurston, T., Trinklein, N. D., Wald, A., & Galloway, D. A. (2016). A Single Human Papillomavirus Vaccine Dose Improves B Cell Memory in Previously Infected Subjects. *Ebiomedicine*, *10*, 55-64. https://doi.org/10.1016/j.ebiom.2016.06.042

Scherpenisse, M., Mollers, M., Schepp, R. M., Boot, H. J., Meijer, C. J., Berbers, G. A. M., Klis, F. v. d., & Melker, H. E. d. (2012). Changes in Antibody Seroprevalence of Seven High-Risk HPV Types Between Nationwide Surveillance Studies From 1995–96 and 2006–07 in the Netherlands. *PLoS One*, *7*(11), e48807. https://doi.org/10.1371/journal.pone.0048807

Scheurer, M. E., Pritchett, J. C., Amirian, E. S., Zemke, N. R., Lusso, P., & Ljungman, P. (2012). HHV-6 Encephalitis in Umbilical Cord Blood Transplantation: A Systematic Review and Meta-Analysis. *Bone Marrow Transplantation*, *48*(4), 574-580. https://doi.org/10.1038/bmt.2012.180
Schiffer, J. T., Wald, A., Selke, S., Corey, L., & Magaret, A. (2011). The Kinetics of Mucosal Herpes Simplex Virus–2 Infection in Humans: Evidence for Rapid Viral-Host Interactions. *The Journal of Infectious Diseases*, *204*(4), 554-561. https://doi.org/10.1093/infdis/jir314
Schneider, C. L., & Hudson, A. W. (2011). The Human Herpesvirus-7 (HHV-7) U21 Immunoevasin Subverts NK-Mediated Cytoxicity Through Modulation of MICA and MICB. *PLOS Pathogens*, *7*(11), e1002362. https://doi.org/10.1371/journal.ppat.1002362
Schuster, F. L., Guglielmo, B. J., & Visvesvara, G. S. (2006). In-Vitro Activity of Miltefosine and Voriconazole on Clinical Isolates of Free-Living Amebas: <i>Balamuthia Mandrillaris, Acanthamoeba</I> Spp., and <i>Naegleria Fowleri</I>. *Journal of Eukaryotic Microbiology*, *53*(2), 121-126. https://doi.org/10.1111/j.1550-7408.2005.00082.x
Schwartz, B. S., & Mawhorter, S. (2013). Parasitic Infections in Solid Organ Transplantation. *American Journal of Transplantation*, *13*, 280-303. https://doi.org/10.1111/ajt.12120
Schwartz, E., Rosenthal, E., & Bank, I. (2014). Gamma Delta T Cells in Non-immune Patients During Primary Schistosomal Infection. *Immunity Inflammation and Disease*, *2*(1), 56-61. https://doi.org/10.1002/iid3.18
Schweininger, J., Scherer, M., Rothemund, F., Schilling, E.-M., Wörz, S., Stamminger, T., & Muller, Y. A. (2021). Cytomegalovirus Immediate-Early 1 Proteins Form a Structurally Distinct Protein Class With Adaptations Determining Cross-Species Barriers. *PLOS Pathogens*, *17*(8), e1009863. https://doi.org/10.1371/journal.ppat.1009863
Seo, M.-G., Lee, S. H., VanBik, D., Ouh, I.-O., Yun, S.-H., Choi, E., Park, Y., Lee, S.-E., Kim, J. W., Cho, G. J., Kwon, O. D., & Kwak, D. (2016). Detection and Genotyping of Coxiella Burnetii and Coxiella-Like Bacteria in Horses in South Korea. *PLoS One*, *11*(5), e0156710. https://doi.org/10.1371/journal.pone.0156710
Sgouras, D. N., Trang, T. T. H., & Yamaoka, Y. (2015). Pathogenesis of <i>Helicobacter Pylori</I> Infection. *Helicobacter*, *20*(S1), 8-16. https://doi.org/10.1111/hel.12251
Shannon, J. G., & Heinzen, R. A. (2008). Infection of Human Monocyte-Derived Macrophages With Coxiella Burnetii. 189-200. https://doi.org/10.1007/978-1-60327-032-8_15
Shao, C., Qu, J., He, L., Zhang, Y., Wang, J., Wang, Y., Zhou, H., & Liu, X. (2005). Transient Overexpression of Γ Interferon Promotes<i>Aspergillus</I>clearance in Invasive Pulmonary Aspergillosis. *Clinical & Experimental Immunology*, *142*(2), 233-241. https://doi.org/10.1111/j.1365-2249.2005.02828.x
Shigemura, K., Osawa, K., Jikimoto, T., Yoshida, H., Hayama, B., Ohji, G., Iwata, K., Fujisawa, M., & Arakawa, S. (2014). Comparison of the Clinical Risk Factors Between Candida Albicans and Candida Non-Albicans Species for Bloodstream Infection. *The Journal of Antibiotics*, *67*(4), 311-314. https://doi.org/10.1038/ja.2013.141
Shikanai-Yasuda, M. A. (2016). Chagas Disease and HIV. 1-10. https://doi.org/10.1007/978-1-4614-9610-6_408-1
Shikoska, I., Djadjovski, I., Krstevski, K., Hristovska, Z. P., Janevsкi, A., Rashikj, L., Cvetkovikj, A., & Cvetkovikj, I. (2023). First Report of <i>Bartonella Henselae</I> Infection in a Cat in North Macedonia. *Macedonian Veterinary Review*, *46*(2), 207-213. https://doi.org/10.2478/macvetrev-2023-0028

Shimizu, Y., Sunaga, N., Dobashi, K., Fueki, M., Fueki, N., Makino, S., & Mori, M. (2009). Serum Markers in Interstitial Pneumonia With and Without Pneumocystis Jiroveciicolonization: A Prospective Study. *BMC Infectious Diseases*, *9*(1). https://doi.org/10.1186/1471-2334-9-47

Shimoda, M., Inagaki, T., Davis, R. R., Merleev, A. A., Tepper, C. G., Maverakis, E., & Izumiya, Y. (2023). Virally Encoded Interleukin-6 Facilitates KSHV Replication in Monocytes and Induction of Dysfunctional Macrophages. *PLOS Pathogens*, *19*(10), e1011703. https://doi.org/10.1371/journal.ppat.1011703

Shorman, M., & Al-Tawfiq, J. A. (2009). Strongyloides Stercoralis Hyperinfection Presenting as Acute Respiratory Failure and Gram-Negative Sepsis in a Patient With Astrocytoma. *International Journal of Infectious Diseases*, *13*(5), e288-e291. https://doi.org/10.1016/j.ijid.2008.11.019

Sia, I. G. (2012). Mycobacteria. 126-137. https://doi.org/10.1093/med/9780199827626.003.0011

Siena, E., Bodini, M., & Medini, D. (2018). Interplay Between Virulence and Variability Factors as a Potential Driver of Invasive Meningococcal Disease. *Computational and Structural Biotechnology Journal*, *16*, 61-69. https://doi.org/10.1016/j.csbj.2018.02.002

Siewert, L. K., Korotaev, A., Sędzicki, J., Fromm, K., Pinschewer, D. D., & Dehio, C. (2022). Identification of The <i>Bartonella</I> Autotransporter CFA as a Protective Antigen and Hypervariable Target of Neutralizing Antibodies in Mice. *Proceedings of the National Academy of Sciences*, *119*(25). https://doi.org/10.1073/pnas.2202059119

Silva, S., Negri, M., Henriques, M., Oliveira, R., Williams, D. W., & Azeredo, J. (2012). <i>Candida Glabrata, Candida Parapsilosis</I>and<i>Candida Tropicalis</I>: Biology, Epidemiology, Pathogenicity and Antifungal Resistance. *Fems Microbiology Reviews*, *36*(2), 288-305. https://doi.org/10.1111/j.1574-6976.2011.00278.x

Silva-Vergara, M. L., Eduardo Rodrigues da Cunha, C., Vissotto, E. D. F., Ana Cristina Araújo Lemos, S., Chica, J. E. L., Etchebehere, R. M., & Adad, S. J. (2007). Disseminated Balamuthia Mandrillaris Amoeba Infection in an AIDS Patient From Brazil. *American Journal of Tropical Medicine and Hygiene*, *77*(6), 1096-1098. https://doi.org/10.4269/ajtmh.2007.77.1096

Singh, V. R., Smith, D. K., Lawerence, J., Kelly, P. C., Thomas, A., Spitz, B., & Sarosi, G. A. (1996). Coccidioidomycosis in Patients Infected With Human Immunodeficiency Virus: Review of 91 Cases at a Single Institution. *Clinical Infectious Diseases*, *23*(3), 563-568. https://doi.org/10.1093/clinids/23.3.563

Smith, D. F., Peacock, C. S., & Cruz, Â. K. (2007). Comparative Genomics: From Genotype to Disease Phenotype in the Leishmaniases. *International Journal for Parasitology*, *37*(11), 1173-1186. https://doi.org/10.1016/j.ijpara.2007.05.015

Smith, S. I., Fowora, M., & Pellicano, R. (2019). Infections With <i>Helicobacter Pylori</I> and Challenges Encountered in Africa. *World Journal of Gastroenterology*, *25*(25), 3183-3195. https://doi.org/10.3748/wjg.v25.i25.3183

Sobel, J. D., Zervos, M., Reed, B. D., Hooton, T. M., Soper, D. E., Nyirjesy, P., Heine, M. W., Willems, J. I. A., & Panzer, H. (2003). Fluconazole Susceptibility of Vaginal Isolates Obtained From Women With Complicated <i>Candida</I> Vaginitis: Clinical Implications. *Antimicrobial Agents and Chemotherapy*, *47*(1), 34-38. https://doi.org/10.1128/aac.47.1.34-38.2003

Soyza, A. D., & Corris, P. A. (2003). Lung Transplantation and the Burkholderia Cepacia Complex. *The Journal of Heart and Lung Transplantation*, *22*(9), 954-958. https://doi.org/10.1016/s1053-2498(03)00024-x

Spandole-Dinu, S., Cimponeriu, D., Toma, M., Radu, I., & Ion, D. A. (2013). Rapid Detection of Human Torque Teno Viruses Using High-Resolution Melting Analysis. *Balkan Journal of Medical Genetics*, *16*(1), 55-62. https://doi.org/10.2478/bjmg-2013-0018

Speth, C., Rambach, G., Lass-Flörl, C., Howell, P. L., & Sheppard, D. C. (2019). Galactosaminogalactan (GAG) and Its Multiple Roles in <i>Aspergillus</I> Pathogenesis. *Virulence*, *10*(1), 976-983. https://doi.org/10.1080/21505594.2019.1568174

Splitter, G. A., & Banai, M. (2006). Microarray Analysis of Brucella Melitensis Pathogenesis. https://doi.org/10.32747/2006.7709884.bard

Sprong, H., Tijsse-Klasen, E., Langelaar, M., Bruin, A. d., Fonville, M., Gassner, F., Takken, W., Wieren, S. v., Nijhof, A. M., Jongejan, F., Maassen, C. B. M., Scholte, E. J., Hovius, J. W., Hovius, K. E., Špitalská, E., & Duynhoven, Y. T. v. (2011). Prevalence of <i>Coxiella Burnetii</I> in Ticks After a Large Outbreak of Q Fever. *Zoonoses and Public Health*, *59*(1), 69-75. https://doi.org/10.1111/j.1863-2378.2011.01421.x

Srivastava, A., Srivastava, A., & Singh, R. K. (2023). Insight Into the Epigenetics of Kaposi's Sarcoma-Associated Herpesvirus. *International Journal of Molecular Sciences*, *24*(19), 14955. https://doi.org/10.3390/ijms241914955

St-Germain, G., Laverdière, M., Pelletier, R., Bourgault, A. M., Libman, M., Lemieux, C., & Noel, G. J. (2001). Prevalence and Antifungal Susceptibility of 442 <i>Candida</I> Isolates From Blood and Other Normally Sterile Sites: Results of a 2-Year (1996 to 1998) Multicenter Surveillance Study in Quebec, Canada. *Journal of Clinical Microbiology*, *39*(3), 949-953. https://doi.org/10.1128/jcm.39.3.949-953.2001

Staerck, C., Yaakoub, H., Vandeputte, P., Tabiasco, J., Godon, C., Gastebois, A., Giraud, S., Guillemette, T., Calenda, A., Delneste, Y., Fleury, M., & Bouchara, J. P. (2021). The Glycosylphosphatidylinositol-Anchored Superoxide Dismutase of Scedosporium Apiospermum Protects the Conidia From Oxidative Stress. *Journal of Fungi*, *7*(7), 575. https://doi.org/10.3390/jof7070575

Šťastná-Marková, M., Hamšíková, E., Hainz, P., Hubáček, P., Kroutilová, M., Kryštofová, J., Ludvíková, V., Musil, J., Pecherková, P., Saláková, M., Šroller, V., Vydra, J., & Němečková, Š. (2021). Pretransplant BK Virus-Specific T-Cell-Mediated Immunity and Serotype Specific Antibodies May Have Utility in Identifying Patients at Risk of BK Virus-Associated Haemorrhagic Cystitis After Allogeneic HSCT. *Vaccines*, *9*(11), 1226. https://doi.org/10.3390/vaccines9111226

Stern-Ginossar, N., Weisburd, B., Michalski, A., Le-Trilling, V. T. K., Hein, M. Y., Huang, S. X., Ma, M., Shen, B., Qian, S. B., Hengel, H., Mann, M., Ingolia, N. T., & Weissman, J. S. (2012). Decoding Human Cytomegalovirus. *Science*, *338*(6110), 1088-1093. https://doi.org/10.1126/science.1227919

Stevenson, M. M., Ing, R., Berretta, F., & Miu, J. (2011). Regulating the Adaptive Immune Response to Blood-Stage Malaria: Role of Dendritic Cells and CD4<sup>+</sup>Foxp3<sup>+</sup> Regulatory T Cells. *International Journal of Biological Sciences*, *7*(9), 1311-1322. https://doi.org/10.7150/ijbs.7.1311

Stewart, E., & Thompson, G. R. (2016). Update on the Epidemiology of Coccidioidomycosis. *Current Fungal Infection Reports*, *10*(4), 141-146. https://doi.org/10.1007/s12281-016-0266-1

Strenger, V., Kayser, S., Witte, K., Laßner, D., Schwinger, W., Jahn, G., Urban, C., & Feuchtinger, T. (2016). Individuals With Inherited Chromosomally Integrated Human Herpes Virus 6 (ciHHV-6) Have Functionally Active HHV-6 Specific T-Cell Immunity. *Clinical Microbiology and Infection*, *22*(2), 209.e205-209.e208. https://doi.org/10.1016/j.cmi.2015.10.006

Suganuma, M., Yamaguchi, K., Ono, Y., Matsumoto, H., Hayashi, T., Ogawa, T., Imai, K., Kuzuhara, T., Nishizono, A., & Fujiki, H. (2008). TNF-α-inducing Protein, a Carcinogenic Factor Secreted From <i>H. Pylori</I>, Enters Gastric Cancer Cells. *International Journal of Cancer*, *123*(1), 117-122. https://doi.org/10.1002/ijc.23484

Sun, R., Lin, S. F., Gradoville, L., Yuan, Y., Zhu, F., & Miller, G. (1998). A Viral Gene That Activates Lytic Cycle Expression of Kaposi's Sarcoma-Associated Herpesvirus. *Proceedings of the National Academy of Sciences*, *95*(18), 10866-10871. https://doi.org/10.1073/pnas.95.18.10866

Sun, Y., Khan, M. A., Mangiola, S., & Barrow, A. D. (2023). IL17RB and IL17REL Expression Are Associated With Improved Prognosis in HPV-Infected Head and Neck Squamous Cell Carcinomas. *Pathogens*, *12*(4), 572. https://doi.org/10.3390/pathogens12040572

Tabtieng, T., Degterev, A., & Gaglia, M. (2018). Caspase-Dependent Suppression of Type I Interferon Signaling Promotes Kaposi's Sarcoma-Associated Herpesvirus Lytic Replication. *J Virol*, *92*(10). https://doi.org/10.1128/jvi.00078-18

Tamm, M., Malouf, M. A., & Glanville, A. R. (2001). Pulmonary Scedosporium Infection Following Lung Transplantation. *Transplant Infectious Disease*, *3*(4), 189-194. https://doi.org/10.1034/j.1399-3062.2001.30402.x

Tartz, S., Deschermeier, C., Retzlaff, S., Heussler, V. T., Šebo, P., Fleischer, B., & Jacobs, T. (2013). <i>Plasmodium Berghei</I> Sporozoite Challenge of Vaccinated <scp>BALB</Scp>/C Mice Leads to the Induction of Humoral Immunity and Improved Function of <scp>CD</Scp>8<sup>+</Sup> Memory <scp>T</Scp> cells. *European Journal of Immunology*, *43*(3), 693-704. https://doi.org/10.1002/eji.201142262

Tavanti, A., Lupetti, A., Ghelardi, E., Corsini, V., Davini, P., Filipponi, F., Boggi, U., Biancofiore, G., Campa, M., & Senesi, S. (2001). Molecular Monitoring of Candida Albicans Infections in Liver Transplant Recipients. *European Journal of Clinical Microbiology & Infectious Diseases*, *20*(8), 544-553. https://doi.org/10.1007/s100960100551

Tesini, B., Wright, T. W., Malone, J., Haidaris, C. G., Harber, M., Sant, A. J., Nayak, J. L., & Gigliotti, F. (2017). Immunization With Pneumocystis Cross-Reactive Antigen 1 (Pca1) Protects Mice Against Pneumocystis Pneumonia and Generates Antibody to Pneumocystis Jirovecii. *Infect Immun*, *85*(4). https://doi.org/10.1128/iai.00850-16

Thakker, S., & Verma, S. C. (2016). Co-Infections and Pathogenesis of KSHV-Associated Malignancies. *Frontiers in Microbiology*, *7*. https://doi.org/10.3389/fmicb.2016.00151

Thompson, G. E. (2011). Pulmonary Coccidioidomycosis. *Seminars in Respiratory and Critical Care Medicine*, *32*(06), 754-763. https://doi.org/10.1055/s-0031-1295723

Thurlow, C. M., Joseph, S. J., Ganova-Raeva, L., Katz, S. S., Pereira, L., Chen, C., Debra, A., Vilfort, K., Workowski, K., Cohen, S. E., Reno, H., Sun, Y., Burroughs, M., Sheth, M., Chi, K.-H., Danavall, D., Philip, S., Cao, W., Kersh, E. N., & Pillay, A. (2022). Selective Whole-Genome Amplification as a Tool to Enrich Specimens With Low Treponema Pallidum Genomic DNA Copies for Whole-Genome Sequencing. *Msphere*, *7*(3). https://doi.org/10.1128/msphere.00009-22

Tian, Y., Pan, S., Zhang, Y., Pei, J., Liu, J., Xie, Y., & Feng, X. (2020). Occurrence and Quantification of Anelloviruses and Herpesviruses in Gingival Tissue in Chinese Shanghai Sub-Population. *BMC Oral Health*, *20*(1). https://doi.org/10.1186/s12903-020-01188-2

Tiggelaar, S. M., Lin, M., Viscidi, R. P., Ji, J., & Smith, J. S. (2012). Age-Specific Human Papillomavirus Antibody and Deoxyribonucleic Acid Prevalence: A Global Review. *Journal of Adolescent Health*, *50*(2), 110-131. https://doi.org/10.1016/j.jadohealth.2011.10.010

Tissot-Dupont, H., Amadei, M.-A., Nezri, M., & Raoult, D. (2004). Wind in November, Q Fever in December. *Emerging Infectious Diseases*, *10*(7), 1264-1269. https://doi.org/10.3201/eid1007.030724

To, K., Cao, R., Yegiazaryan, A., Owens, J., & Venketaraman, V. (2020). General Overview of Nontuberculous Mycobacteria Opportunistic Pathogens: Mycobacterium Avium and Mycobacterium Abscessus. *Journal of Clinical Medicine*, *9*(8), 2541. https://doi.org/10.3390/jcm9082541

Toussi, S. S., Pan, N., Walters, H., & Walsh, T. J. (2013). Infections in Children and Adolescents With Juvenile Idiopathic Arthritis and Inflammatory Bowel Disease Treated With Tumor Necrosis Factor-  Inhibitors: Systematic Review of the Literature. *Clinical Infectious Diseases*, *57*(9), 1318-1330. https://doi.org/10.1093/cid/cit489

Tripet, F. (2009). Ecological Immunology of Mosquito-Malaria Interactions: Of Non-Natural Versus Natural Model Systems and Their Inferences. *Parasitology*, *136*(14), 1935-1942. https://doi.org/10.1017/s0031182009006234

Tse, S. W., Radtke, A. J., Espinosa, D. A., Cockburn, I. A., & Zavala, F. (2014). The Chemokine Receptor CXCR6 Is Required for the Maintenance of Liver Memory CD8+ T Cells Specific for Infectious Pathogens. *The Journal of Infectious Diseases*, *210*(9), 1508-1516. https://doi.org/10.1093/infdis/jiu281

Tsiouma, G. K. (2023). Chronic Cough an Unlikely Complication of Pharyngolaryngeal Herpes Zoster. *Journal of Clinical and Medical Images Case Reports*, *3*(3). https://doi.org/10.55920/2771-019x/1437

Tu, M., Liu, F., Chen, S., Wang, M., & Cheng, A. (2015). Role of Capsid Proteins in Parvoviruses Infection. *Virology Journal*, *12*(1). https://doi.org/10.1186/s12985-015-0344-y

Tu, W., & Rao, S. (2016). Mechanisms Underlying T Cell Immunosenescence: Aging and Cytomegalovirus Infection. *Frontiers in Microbiology*, *7*. https://doi.org/10.3389/fmicb.2016.02111

Tuddenham, L., Jung, J., Chane-Woon-Ming, B., Dölken, L., & Pfeffer, S. (2012). Small RNA Deep Sequencing Identifies MicroRNAs and Other Small Noncoding RNAs From Human Herpesvirus 6B. *J Virol*, *86*(3), 1638-1649. https://doi.org/10.1128/jvi.05911-11

Tyschik, E. A., Shcherbakova, S. M., Ибрагимов, P. P., & Rebrikov, D. V. (2017). Transplacental Transmission of Torque Teno Virus. *Virology Journal*, *14*(1). https://doi.org/10.1186/s12985-017-0762-0

Ullah, H., Ikhtiar, Z., & Ikhtiar, H. (2024). A Rare Presentation of Herpes Simplex Virus Encephalitis With Ischemic Stroke and Associated Acyclovir-Induced Acute Kidney Injury. *Cureus*. https://doi.org/10.7759/cureus.71978

Vaccarella, S., Franceschi, S., Clifford, G. M., Touzé, A., Hsu, C. C., Sanjosé, S. d., Anh, P. T. H., Hieu, N. T., Matos, E., Shin, H. R., Sukvirach, S., Thomas, J. O., Boursaghin, L., Gaitan, J., Snijders, P. J., Meijer, C. J., Muñóz, N., Herrero, R., & Coursaget, P. (2010). Seroprevalence of Antibodies

Against Human Papillomavirus (HPV) Types 16 and 18 in Four Continents: The International Agency for Research on Cancer HPV Prevalence Surveys. *Cancer Epidemiology Biomarkers & Prevention*, *19*(9), 2379-2388. https://doi.org/10.1158/1055-9965.epi-10-0336
Valigurová, A. B., & Kolářová, I. (2023). Unrevealing the Mystery of Latent Leishmaniasis: What Cells Can Host Leishmania? *Pathogens*, *12*(2), 246. https://doi.org/10.3390/pathogens12020246
Vanhove, M., Beale, M. A., Rhodes, J., Chanda, D., Lakhi, S., Kwenda, G., Molloy, S. F., Karunaharan, N., Stone, N., Harrison, T. S., Bicanic, T., & Fisher, M. C. (2016). Genomic Epidemiology of <i>Cryptococcus</I> Yeasts Identifies Adaptation to Environmental Niches Underpinning Infection Across an African <scp>HIV</Scp>/<scp>AIDS</Scp> Cohort. *Molecular Ecology*, *26*(7), 1991-2005. https://doi.org/10.1111/mec.13891
Vazirani, J., Crowhurst, T., Morrissey, C. O., & Snell, G. I. (2021). Management of Multidrug Resistant Infections in Lung Transplant Recipients With Cystic Fibrosis. *Infect Drug Resist*, *Volume 14*, 5293-5301. https://doi.org/10.2147/idr.s301153
Vermis, K. (2002). Evaluation of Restriction Fragment Length Polymorphism Analysis of 16S rDNA as a Tool for Genomovar Characterisation Within the Burkholderia Cepacia Complex. *Fems Microbiology Letters*, *214*(1), 1-5. https://doi.org/10.1016/s0378-1097(02)00845-5
Vermis, K., Vandekerckhove, C., Nelis, H. J., & Vandamme, P. (2002). Evaluation of Restriction Fragment Length Polymorphism Analysis of 16S rDNA as a Tool for Genomovar Characterisation Within The<i>Burkholderia Cepacia</I>complex. *Fems Microbiology Letters*, *214*(1), 1-5. https://doi.org/10.1111/j.1574-6968.2002.tb11316.x
Veugen, J. M. J., Savelkoul, P. H. M., Nuijts, R. M., Dickman, M. M., & Wolffs, P. (2025). Enhancing <i>Acanthamoeba</I> Diagnostics: Rapid Detection of Viable <i>Acanthamoeba</I> Trophozoites and Cysts Using Viability PCR Assay. *Microbiology Spectrum*, *13*(3). https://doi.org/10.1128/spectrum.01811-24
Viana, E. C., Silva, A. L. d., Lucas Coutinho Tuma da, P., Mota, L. O., Silva, E. S. S., Santos, C. N. B., Santos, M. C. F., Valença, J. G. S., & Bichara, C. N. C. (2022). Prevalence of the Transmission Type of Chagas Disease in the Brazilian Amazon. *International Journal of Health Science*, *2*(15), 2-5. https://doi.org/10.22533/at.ed.1592152215032
Virgin, H. W., Latreille, P., Wamsley, P., Hallsworth, K., Weck, K. E., Canto, A. J. D., & Speck, S. H. (1997). Complete Sequence and Genomic Analysis of Murine Gammaherpesvirus 68. *J Virol*, *71*(8), 5894-5904. https://doi.org/10.1128/jvi.71.8.5894-5904.1997
Viscuse, P. V., & Mohabbat, A. B. (2019). 69-Year-Old Woman With Fatigue, Dyspnea, and Lower Extremity Pain. *Mayo Clinic Proceedings*, *94*(1), 149-154. https://doi.org/10.1016/j.mayocp.2018.04.028
Visvesvara, G. S., Moura, H., & Schuster, F. L. (2007). Pathogenic and Opportunistic Free-Living Amoebae:<i>Acanthamoeba</I>spp.,<i>Balamuthia Mandrillaris</I>,<i>Naegleria Fowleri</I>, And<i>Sappinia Diploidea</I>. *Fems Immunology & Medical Microbiology*, *50*(1), 1-26. https://doi.org/10.1111/j.1574-695x.2007.00232.x
Vojdani, A., Almulla, A. F., Zhou, B., Al-Hakeim, H. K., & Maes, M. (2024). Reactivation of Herpesvirus Type 6 and IgA/IgM-mediated Responses to Activin-a Underpin Long COVID, Including Affective Symptoms and Chronic Fatigue Syndrome. *Acta Neuropsychiatrica*, *36*(3), 172-184. https://doi.org/10.1017/neu.2024.10

Voth, D. E., & Heinzen, R. A. (2009). Coxiella Type IV Secretion and Cellular Microbiology. *Current Opinion in Microbiology*, *12*(1), 74-80. https://doi.org/10.1016/j.mib.2008.11.005
Walzer, P. D., Djawe, K., Levin, L., Daly, K. R., Koch, J., Kingsley, L., Witt, M. D., Golub, E. T., Bream, J. H., Taiwo, B., & Morris, A. (2009). Long-Term Serologic Responses to The<i>Pneumocystis Jirovecii</I>Major Surface Glycoprotein in HIV-Positive Individuals With and Without<i>P. Jirovecii</I>Infection. *The Journal of Infectious Diseases*, *199*(9), 1335-1344. https://doi.org/10.1086/597803
Wang, C.-h., Sun, Q., Yan, J., Liao, X., Long, S., Zheng, M., Zhang, Y., Yang, X., Shi, G., Zhao, Y., Wang, G., & Pan, J. (2023). The Species Distribution and Antimicrobial Resistance Profiles of Nocardia Species in China: A Systematic Review and Meta-Analysis. *Plos Neglected Tropical Diseases*, *17*(7), e0011432. https://doi.org/10.1371/journal.pntd.0011432
Wang, D., Hu, M., Liu, W., Zhao, Y., & Xu, Y. (2024). Bacteremia Caused by Nocardia Farcinica: A Case Report and Literature Review. *BMC Infectious Diseases*, *24*(1). https://doi.org/10.1186/s12879-024-09230-2
Wang, H., Wu, Y., Ojcius, D. M., Yang, X., Zhang, C., Ding, S.-b., Lin, X. a., & Yan, J. (2012). Leptospiral Hemolysins Induce Proinflammatory Cytokines Through Toll-Like Receptor 2-and 4-Mediated JNK and NF-κB Signaling Pathways. *PLoS One*, *7*(8), e42266. https://doi.org/10.1371/journal.pone.0042266
Wang, H., Zhu, Y., Cui, Q., Wu, W., Li, G., Chen, D., Xiang, L., Qu, J., Shi, D., & Lu, B. (2022). Epidemiology and Antimicrobial Resistance Profiles of The <i>Nocardia</I> Species in China, 2009 to 2021. *Microbiology Spectrum*, *10*(2). https://doi.org/10.1128/spectrum.01560-21
Wang, Q., Zhang, Y., Yang, Q., Fu, S., Qu, B., & Defoirdt, T. (2021). One Health Pathogen Surveillance Demonstrated the Dissemination of Gut Pathogens Within the Two Coastal Regions Associated With Intensive Farming. *Gut Pathogens*, *13*(1). https://doi.org/10.1186/s13099-021-00442-4
Wang, W., Wang, X., Yang, L., Fu, W., Pan, D., Liu, J., Ye, J., Zhao, Q., Zhu, H., Cheng, T., & Xia, N. (2016). Modulation of Host CD59 Expression by Varicella-Zoster Virus in Human Xenografts in Vivo. *Virology*, *491*, 96-105. https://doi.org/10.1016/j.virol.2016.01.019
Warnke, C., Adams, O., Gold, R., Hartung, H. P., Hohlfeld, R., Wiendl, H., & Kieseier, B. C. (2011). Progressive Multifokale Leukenzephalopathie Unter Natalizumab. *Der Nervenarzt*, *82*(4), 475-480. https://doi.org/10.1007/s00115-010-3091-8
Warren-Gash, C., Forbes, H., & Breuer, J. (2017). Varicella and Herpes Zoster Vaccine Development: Lessons Learned. *Expert Review of Vaccines*, *16*(12), 1191-1201. https://doi.org/10.1080/14760584.2017.1394843
Weiss, G. E., Traoré, B., Kayentao, K., Ongoïba, A., Doumbo, S., Doumtabé, D., Koné, Y., Dia, S., Guindo, A., Traoré, A., Huang, C. Y., Miura, K., Mircetic, M., Li, S., Baughman, A., Narum, D. L., Miller, L. H., Doumbo, O. K., Pierce, S. K., & Crompton, P. D. (2010). The Plasmodium Falciparum-Specific Human Memory B Cell Compartment Expands Gradually With Repeated Malaria Infections. *PLOS Pathogens*, *6*(5), e1000912. https://doi.org/10.1371/journal.ppat.1000912
Werner, J., Ishida, K., Wisler, J., Karbowski, C. M., Kalanzi, J., Bussiere, J. L., & Monticello, T. M. (2020). Phosphatidylinositol 3-Kinase Δ Inhibitor-Induced Immunomodulation and Secondary Opportunistic Infection in the Cynomolgus Monkey (<i>Macaca Fascicularis</I>). *Toxicologic Pathology*, *48*(8), 949-964. https://doi.org/10.1177/0192623320966238

Wheat, J. (1997). Histoplasmosis: Experience During Outbreaks in Indianapolis and Review of the Literature. *Medicine*, *76*(5), 339-354. https://doi.org/10.1097/00005792-199709000-00002
Wheat, L. J. (1982). Risk Factors for Disseminated or Fatal Histoplasmosis. *Annals of Internal Medicine*, *96*(2), 159-163. https://doi.org/10.7326/0003-4819-96-2-159
Williamson, E., Speers, D., Arthur, I., Harnett, G. B., Ryan, G., & Inglis, T. (2001). Molecular Epidemiology Of <i>Scedosporium Apiospermum</I> Infection Determined by PCR Amplification of Ribosomal Intergenic Spacer Sequences in Patients With Chronic Lung Disease. *Journal of Clinical Microbiology*, *39*(1), 47-50. https://doi.org/10.1128/jcm.39.1.47-50.2001
Wilson, J. (2012). Nocardiosis: Updates and Clinical Overview. *Mayo Clinic Proceedings*, *87*(4), 403-407. https://doi.org/10.1016/j.mayocp.2011.11.016
Wilson, L. E., Pawlita, M., Castle, P. E., Waterboer, T., Sahasrabuddhe, V. V., Gravitt, P. E., Schiffman, M., & Wentzensen, N. (2013). Natural Immune Responses Against Eight Oncogenic Human Papillomaviruses in the ASCUS-LSIL Triage Study. *International Journal of Cancer*, *133*(9), 2172-2181. https://doi.org/10.1002/ijc.28215
Wilson, M. S., Mentink-Kane, M., Pesce, J., Ramalingam, T. R., Thompson, R. W., & Wynn, T. A. (2006). Immunopathology of Schistosomiasis. *Immunol Cell Biol*, *85*(2), 148-154. https://doi.org/10.1038/sj.icb.7100014
Wipasa, J., Suphavilai, C., Okell, L., Cook, J., Corran, P. H., Thaikla, K., Liewsaree, W., Riley, E. M., & Hafalla, J. C. R. (2010). Long-Lived Antibody and B Cell Memory Responses to the Human Malaria Parasites, Plasmodium Falciparum and Plasmodium Vivax. *PLOS Pathogens*, *6*(2), e1000770. https://doi.org/10.1371/journal.ppat.1000770
Wong, Y. Y., Lee, C. W., Bong, C. W., Lim, J. H., Ng, C. C., Narayanan, K., Sim, E. U., & Wang, A. (2024). Environmental Factors That Regulate Vibrio Spp. Abundance and Community Structure in Tropical Waters. *Anthropocene Coasts*, *7*(1). https://doi.org/10.1007/s44218-024-00054-w
Wu, T.-T., Usherwood, E. J., Stewart, J. P., Nash, A. A., & Sun, R. (2000). Rta of Murine Gammaherpesvirus 68 Reactivates the Complete Lytic Cycle From Latency. *J Virol*, *74*(8), 3659-3667. https://doi.org/10.1128/jvi.74.8.3659-3667.2000
Xavier, M. N., Paíxão, T. A., Hartigh, A. B. d., Tsolis, R. M., & Santos, R. L. (2010). Pathogenesis of Brucella Spp. *The Open Veterinary Science Journal*, *4*(1), 109-118. https://doi.org/10.2174/1874318801004010109
Xi, Y., Li, X., Liu, L. Z., Xiu, F., Yi, X., Chen, H., & You, X. (2024). Sneaky Tactics: Ingenious Immune Evasion Mechanisms Of <i>Bartonella</I>. *Virulence*, *15*(1). https://doi.org/10.1080/21505594.2024.2322961
Xiao, Y., Li, M., Guo, X., Zeng, H., Shuai, X., Guo, J., Huang, Q., Chu, Y., Zhou, B., Wen, J. J., Liu, J., & Jiao, H. (2022). Inflammatory Mechanism of Brucella Infection in Placental Trophoblast Cells. *International Journal of Molecular Sciences*, *23*(21), 13417. https://doi.org/10.3390/ijms232113417
Xin, H., Dziadek, S., Bundle, D. R., & Cutler, J. E. (2008). Synthetic Glycopeptide Vaccines Combining Β-Mannan and Peptide Epitopes Induce Protection Against Candidiasis. *Proceedings of the National Academy of Sciences*, *105*(36), 13526-13531. https://doi.org/10.1073/pnas.0803195105
Yamamoto, M., Okuyama, M., Su, J., Kimura, T., Kamiyama, N., Saiga, H., Ohshima, J., Sasai, M., Kayama, H., Okamoto, T., Huang, D. C., Soldati-Favre, D., Horie, K., Takeda, J., & Takeda, K. (2012). A Cluster of Interferon-Γ-Inducible P65 GTPases Plays a Critical Role in Host Defense

Against Toxoplasma Gondii. *Immunity*, *37*(2), 302-313. https://doi.org/10.1016/j.immuni.2012.06.009

Yamamoto, Y., Morooka, M., Hashimoto, S., Ihra, M., & Yoshikawa, T. (2013). Analysis of the Shedding of Three Β-herpesviruses in Urine and Saliva of Children With Renal Disease. *J Med Virol*, *86*(3), 505-511. https://doi.org/10.1002/jmv.23782

Yang, F., Gritsenko, V., Lu, H., Zhen, C., Gao, L., Berman, J., & Jiang, Y. (2021). Adaptation to Fluconazole via Aneuploidy Enables Cross-Adaptation to Amphotericin B and Flucytosine in Cryptococcus Neoformans. *Microbiology Spectrum*, *9*(2). https://doi.org/10.1128/spectrum.00723-21

Yang, N., Tong, Z., Wang, Z., Xu, M., Zhang, Q., Zhang, H., Wang, Y., Yi, J., Sun, T., Cui, B., & Chen, C. (2020). Brucella Outer Membrane Proteins to Control Maturation and Antigen Presentation of Mouse Dendritic Cells. https://doi.org/10.21203/rs.3.rs-57350/v1

Yao, Y.-F., Jia, F., Liu, J., Chen, C.-F., Yu, B., & Hu, X. (2021). Disseminated Infection by <i>Fusarium Solani </I>in Acute Lymphocytic Leukemia: A Case Report. *World Journal of Clinical Cases*, *9*(22), 6443-6449. https://doi.org/10.12998/wjcc.v9.i22.6443

Ye, F., Lei, X., & Gao, S. J. (2011). Mechanisms of Kaposi's Sarcoma-Associated Herpesvirus Latency and Reactivation. *Advances in Virology*, *2011*, 1-19. https://doi.org/10.1155/2011/193860

Ye, L., Taylor, G. P., & Rosadas, C. (2022). Human T-Cell Lymphotropic Virus Type 1 and Strongyloides Stercoralis Co-Infection: A Systematic Review and Meta-Analysis. *Frontiers in Medicine*, *9*. https://doi.org/10.3389/fmed.2022.832430

Yi, D., Xu, Q. Y., He, Y., Zheng, X. Q., Yang, T. C., & Lin, Y. (2024). <i>Treponema Pallidum</I> Protein Tp47 Induced Prostaglandin <scp>E2</Scp> to Inhibit the Phagocytosis in Human Macrophages. *Journal of the European Academy of Dermatology and Venereology*, *38*(6), 1166-1178. https://doi.org/10.1111/jdv.19809

Yi, J. (2019). LRRK2 Kinase Plays an Important Role in the Intracellular Survival of Brucella Abortus 2308 in Murine Macrophages and in a Mouse Infection Model. *Pakistan Veterinary Journal*, *39*(04), 534-540. https://doi.org/10.29261/pakvetj/2019.050

Yohannan, B., & Feldman, M. (2019). Fatal<i>Balamuthia Mandrillaris</I>Encephalitis. *Case Reports in Infectious Diseases*, *2019*, 1-5. https://doi.org/10.1155/2019/9315756

Yu, S., Wang, J., Fang, Q., Zhang, J., & Yan, F. (2016). Specific Clinical Manifestations of Nocardia: A Case Report and Literature Review. *Experimental and Therapeutic Medicine*, *12*(4), 2021-2026. https://doi.org/10.3892/etm.2016.3571

Yu, V. L., Plouffe, J. F., Pastoris, M. C., Stout, J. E., Schousboe, M., Widmer, A. F., Summersgill, J. T., File, T. M., Heath, C. H., Paterson, D. L., & Chereshsky, A. (2002). Distribution Of<i>Legionella</I>Species and Serogroups Isolated by Culture in Patients With Sporadic Community-Acquired Legionellosis: An International Collaborative Survey. *The Journal of Infectious Diseases*, *186*(1), 127-128. https://doi.org/10.1086/341087

Zalani, S., Holley-Guthrie, E., & Kenney, S. C. (1996). Epstein-Barr Viral Latency Is Disrupted by the Immediate-Early BRLF1 Protein Through a Cell-Specific Mechanism. *Proceedings of the National Academy of Sciences*, *93*(17), 9194-9199. https://doi.org/10.1073/pnas.93.17.9194

Zhang, J., Wang, Z., Chen, Y., Zhou, Z., Yang, Q., Fu, Y., Zhao, F., Li, X., Chen, Q., Li, F., Jiang, Y., & Yu, Y. (2022). Antifungal Susceptibility and Molecular Characteristics of Cryptococcus Spp. Based on Whole-Genome Sequencing in Zhejiang Province, China. *Frontiers in Microbiology*, *13*. https://doi.org/10.3389/fmicb.2022.991703

Zheng, Y., Zhao, Y., Wang, Y., & Rao, J. (2021). A Multiplex Real-Time PCR Quantitation of Human Herpesvirus-6, 7, 8 Viruses: Application in Blood Transfusions. *Virology Journal*, *18*(1). https://doi.org/10.1186/s12985-021-01510-6

Zheng, Y. W., Zheng, X. Q., Guo, Y., Xie, J.-W., Wang, M., Xu, Q. Y., Zhu, X. Z., & Lin, L. R. (2025). Characteristics of Immune Response and Pathogen Tissue Dissemination During Progressive <i>Treponema Pallidum</I> Infection: Insights From Humanized Mice. *Scandinavian Journal of Immunology*, *101*(3). https://doi.org/10.1111/sji.70005

Zheng, Z., Chen, F., Qin, L., Lu, A., Xu, H., Zhao, M., & Zhao, Y. (2024). Application of Ventriculoscopy in Granulomatous Amoebic Encephalitis: A Case Report in China and Literature Review. *Frontiers in Medicine*, *11*. https://doi.org/10.3389/fmed.2024.1431225

Zhou, X., Chen, X., Zheng, X., & Xia, X. (2023). Curative Effect of Helicobacter Pylori Radical Treatment on Gastric Cancer Patients With Helicobacter Pylori Infection. *Academic Journal of Medicine & Health Sciences*, *4*(6). https://doi.org/10.25236/ajmhs.2023.040614

Zhu, A., Ma, Q., & Liu, Z. (2024). Omadacycline for the Treatment of Patients With Legionella Pneumophila Pneumonia After Experiencing Liver Dysfunction: Case Series. *Frontiers in Microbiology*, *15*. https://doi.org/10.3389/fmicb.2024.1408443

Zhu, S., & Viejo-Borbolla, A. (2021). Pathogenesis and Virulence of Herpes Simplex Virus. *Virulence*, *12*(1), 2670-2702. https://doi.org/10.1080/21505594.2021.1982373

Zou, Z., Meng, Z., Ma, C., Liang, D., Sun, R., & Lan, K. (2017). Guanylate-Binding Protein 1 Inhibits Nuclear Delivery of Kaposi's Sarcoma-Associated Herpesvirus Virions by Disrupting Formation of Actin Filament. *J Virol*, *91*(16). https://doi.org/10.1128/jvi.00632-17

Zuhair, M., Smit, G. S. A., Wallis, G., Jabbar, F., Smith, C., Devleesschauwer, B., & Griffiths, P. (2019). Estimation of the Worldwide Seroprevalence of Cytomegalovirus: A Systematic Review and Meta-analysis. *Reviews in Medical Virology*, *29*(3). https://doi.org/10.1002/rmv.2034
